# Genetic architecture of the developing forelegs of *Drosophila prolongata*; an exaggerated weapon and ornament

**DOI:** 10.1101/2025.10.09.681446

**Authors:** Tyler Audet, Jhoniel Perdigón Ferreira, Abhishek Meena, Mariam Abass, Arteen Torabi-Marashi, John Yeom, Fatima Zaghloul, Nour Zaghloul, Arie Mizrahi, Julia Novikov, Emma Xi, Stefan Lüpold, Ian Dworkin

## Abstract

Extreme secondary sexual traits are some of the most striking phenotypes in nature. Studies on the genetics of these phenotypes have largely focused on within-species functional analyses of signalling pathways. Although useful, these do not provide insight into the evolutionary mechanisms that occur during the evolution of trait exaggeration. *Drosophila prolongata* offers an exceptional opportunity to explore the evolution of trait exaggeration, as it is the only species in the melanogaster species group with male-specific foreleg size exaggeration under both intra- and intersexual selection. Here, we used sex-specific RNA-seq from fore- and midleg tissues during early development and after initiation of sexually dimorphic growth between these tissues in *D. prolongata*. We also sampled the same developmental stages in *D. carrolli* (∼4MYA divergence) and *D. melanogaster* (∼20MYA). Using comparisons of gene expression between sexes, species, tissues, and developmental stages, we found a positive relationship between the number, but not the magnitude of differential expression of sex-biased genes, with the extent of phenotypic dimorphism. One gene with a large effect, *grain*, caused *D. prolongata*-like leg size phenotypes in *D. melanogaster* legs when knocked down. We further found only modest changes to magnitude and direction of expression differences in signalling pathways previously implicated in sexually dimorphic evolution. This suggests that these pathways regulating trait expression and dimorphism but may not be primary drivers of their phenotypic evolution.

**Significance statement:** How sexes evolve distinct forms while developing from a shared genome continues to be incompletely resolved. We explore a sexually dimorphic exaggerated trait, enlarged forelegs in *Drosophila prolongata*, and compare changes in sex-biased gene expression between tissues and developmental stages in *D. prolongata* and two closely related species without foreleg exaggeration. We show that major developmental pathways don’t appear to change substantially in their direction or magnitude of expression between species. We further show a transcription factor, *grain*, that when knocked-down, induces *D. prolongata* leg-like phenotypes in *D. melanogaster* at low penetrance. Our results suggest that despite large morphological differences and patterns of dimorphism, gene expression changes between the developing tissues may be more modest than previously thought.

## Introduction

Elaborate or exaggerated secondary sexual traits are among the most striking features in the animal kingdom. These traits fall along a spectrum for two functions, from weapons for fighting or intimidating conspecifics of the same sex to ornaments used for attracting mates of the opposite sex (McCullough et al. 2016). Exaggerated weapons typically evolve when ecological conditions allow their bearer to secure disproportionate mating success in direct contests (Emlen 2008). By contrast, ornaments generally evolve as a consequence of female preferences (Darwin 1871), for example driven by indirect benefits via good genes (Kirkpatrick and Ryan 1991; Kirkpatrick 1996), processes like Fisherian runaway selection (Fisher 1930), among other mechanism. These secondary sexual traits can constitute a substantial proportion of overall “size” relative to an individual’s body (Kodric-Brown et al. 2006; Voje 2016). Consequently, understanding their genetic architecture is of great interest to evolutionary biologists, as it helps assess the plausibility of different evolutionary models for sexual selection and reveal how inter-sex genetic correlations can either facilitate adaptive evolution or create sexual conflict by hindering the sexes from reaching their specific optima.

Early models of Fisherian runaway and good genes processes often used a population genetic framework, assuming simple genetic architectures (Fisher 1930). However, not all predictions of these models are supported under biologically plausible, highly polygenic genetic architectures for ornaments (Pomiankowski and Møller 1997). One example is the lek paradox, where genetic variation for ornaments is predicted to deplete rapidly under some forms of good genes models (Kirkpatrick and Ryan 1991). Yet, empirically, these traits often have as much, if not greater genetic variances than traits associated with other fitness components (Pomiankowski and Møller 1997). Resolutions to this “paradox” have been proposed, for example via the evolutionary co-option of mechanisms influencing organismal condition (Rowe and Houle 1996). However, if the trait that ultimately becomes the target of sexual selection is polygenic from the outset (either with an infinitesimal or omnigenic-like distribution of allelic effects), the paradox disappears. Even strong and continued sexual selection would not deplete genetic variance as substantial phenotypic response could occur via modest shifts in genome-wide allele frequencies, accompanied by the replenishment of variation through new mutations (Rowe and Houle 1996). If the genetic architecture of exaggerated trait variation relies on few genes of large phenotypic effect, however, this fundamentally alters predictions.

The genetic mechanisms through which sexual dimorphism is achieved is still an open area of inquiry. This involves understanding the sex-specific mode-of-action of allelic effects (i.e. genotype-by-sex interactions) (Testa and Dworkin 2016; Zhu et al. 2023). While some sexually selected traits, like the sex combs among species of *Drosophila,* are sex-limited, many others represent sex-specific elaborations or exaggerations (Kopp 2011). Knowing the relative contribution of sex-limited genetic effects versus sex-specific amplification genetic effects, is crucial for our understanding of how sexual dimorphism evolves.

One mechanism implicated in generating sexual dimorphism and potentially resolving sexual conflict from a largely shared genome, is sex-biased gene expression during development (Perry et al. 2014). Sex differences in gene expression have been evaluated across many species, tissues, and time points (Ellegren and Parsch 2007; Parsch and Ellegren 2013; Perry et al. 2014; Ingleby et al. 2015; Mank 2017). Whole-body studies often suggest moderate to substantial sex-biased expression in a large fraction of the genome, whilst more targeted, tissue-specific studies (particularly during trait development) have generally identified few, though still several hundred genes with substantial sex-differential expression, a number that typically increases in adult tissues (ref. 18 in *Drosophila* as an example). While distinguishing cause from effect in these studies is challenging, several broad inferences have been drawn. Among sexually-dimorphic (but not sex-limited) morphological traits, there is evidence for an increase in number of sex-biased genes and often the magnitude of expression of sex-biased genes in the associated tissues during development (Zinna et al. 2018; Toubiana et al. 2021). When sexual dimorphism involves size exaggeration, genes related to growth control are commonly found (Moczek and Rose 2009; Wilkinson et al. 2013; Cox et al. 2017; Zinna et al. 2018; Toubiana et al. 2021). Intriguingly, even for traits with male-biased size dimorphism, the relevant tissues can show both male- and female-biased genes (Toubiana et al. 2021). The implication of these studies is that sexual dimorphism is likely mediated by numerous genes that collectively influence its extent.

In contrast to transcriptional studies, functional genetic studies directly test the role of genes in development, thereby revealing not only sex differences in gene expression, but also differences in gene-specific sensitivity to perturbation. As with expression profiling, genes associated with growth signalling have been implicated in the development of sexually exaggerated traits. Due to the common pattern of condition dependence of these structures, primary candidates for exaggerated trait growth are insulin and Target of Rapamycin (TOR) signaling pathways due to their central role in regulating cell proliferation, growth, and metabolic processes (Warren et al. 2013). In *Drosophila melanogaster*, disrupting insulin signalling disproportionately reduces the size of the larger sex, thereby diminishing sexual dimorphism (Shingleton et al. 2005; Rideout et al. 2015; Sawala and Gould 2017; McDonald et al. 2021; Millington, Brownrigg, Basner-Collins, et al. 2021; Millington, Brownrigg, Chao, et al. 2021). These previous results, as well as their potential to explain condition-dependent signalling, make the insulin signalling pathway a strong candidate for pathways underlying sexually dimorphic evolution. Genetic knockdowns of *insulin receptor* (*InR*) using RNAi in the rhinoceros beetle *Trypoxylus dichotomus* reduces or eliminates horns (Emlen et al. 2012), with similar results observed for sexually dimorphic horn development in dung beetles (Casasa and Moczek 2018; Rohner et al. 2023). Insulin signalling genes show similar responses during development of exaggerated tissues in *T. dichotomus* (Zinna et al. 2018). Finally, condition-dependent expression and phenotypic responses to knocking down insulin-like peptide has also been shown to influence exaggerated weapon (mandible) size in the flour beetle *Gnatocerus cornutus* (Okada et al. 2019).

Somatic sexual differentiation in holometabolous insects is largely cell-autonomous, as observed in gynandromorphic individuals (Dobzhansky 1931; J Patterson 1938; Jahner et al. 2015). In *Drosophila*, this is mediated by the core sex-determination pathway initiated by the X-linked, and dosage-dependent *Sex-lethal* (*Sxl*) gene, resulting in a pattern of sex-specific production of downstream gene isoforms such as *transformer* (*tra*) and *doublesex (dsx)*, and ultimately sex-biased gene expression within and between tissues (Gowen and Fung 1957; Baker and Ridge 1980; Yan and Perrimon 2015; Hérault et al. 2024). This differential expression influences sex-specific growth patterns via hormonal modulation of metabolism and cell-autonomous mechanisms (Rideout et al. 2015; Mathews et al. 2017; Millington, Brownrigg, Chao, et al. 2021; Wat et al. 2021). *dsx* also mediates trait expression for the male-limited sex combs in *D. melanogaster* (Tanaka et al. 2011; Rice et al. 2019). Knockdowns of *dsx* reduces male horn size in the dung beetle *Onthophagous taurus* and *O. sagittarius* (Kijimoto et al. 2012; Ledón-Rettig et al. 2017), and the exaggeration of male mandibles in the golden stag beetle C*yclommatus metallifer*, but can induce rudimentary horns or mandible growth in females (Gotoh et al. 2014; Gotoh et al. 2016). Beyond these core pathways, perturbing genes involved in axis specification and limb patterning can also reduce sex-specific trait exaggeration, such as the Hox gene *ultrabithorax* (*ubx*) in water strider legs (Khila et al. 2009; Refki et al. 2014; Crumière and Khila 2019) or *sex-combs reduced* (*scr*) in *O. nigriventris* horn size (Wasik et al. 2010).

Interestingly, genes that mediate limb patterning and growth, such as *distal-less* (*dll*), have also been implicated in the development of beetle horns (Moczek and Rose 2009). The perturbation of these genes provides important information on which genes are essential for trait expression, but they do not necessarily pinpoint evolutionary changes responsible for the origin and exaggeration or of a trait itself.

While trait-specific sexual size dimorphism is common across the ∼177 species of the *Drosophila melanogaster* subgroup (Kopp 2006), it is typically modest (less than 15% differences in size) and female-biased, likely due to fecundity selection. *Drosophila prolongata*, a member of the Southeast Asian rhopaloa clade (Singh 1977), is a notable exception for its male-biased body size dimorphism among Drosophilids (Rohner et al. 2018) and exaggerated male forelegs (Figure 1). In *D. prolongata*, larger forelegs benefit males in contests and, thus, increase mating success, indicating a role as weapons (Toyoshima and Matsuo 2023). These legs are also patterned in alternating black and white bands and are used in novel courtship displays, such as arm waving and female stimulation by leg vibration, suggesting a dual function as ornaments subject to female preference (Setoguchi et al. 2014). These traits are absent in the close relative *D. carrolli* (Gompel and Kopp 2018), despite a relatively recent divergence of ∼4 million years (Setoguchi et al. 2014). The natural histories of *D. prolongata* and *D. carrolli* are largely unknown, and therefore the evolutionary context of this trait exaggeration is an outstanding question. However, the evolution of trait exaggeration in just one lineage within this clade makes *D. prolongata* an exceptional model for investigating the evolutionary mechanisms underlying the origin and exaggeration of a sexually selected trait.

**Figure 1:**
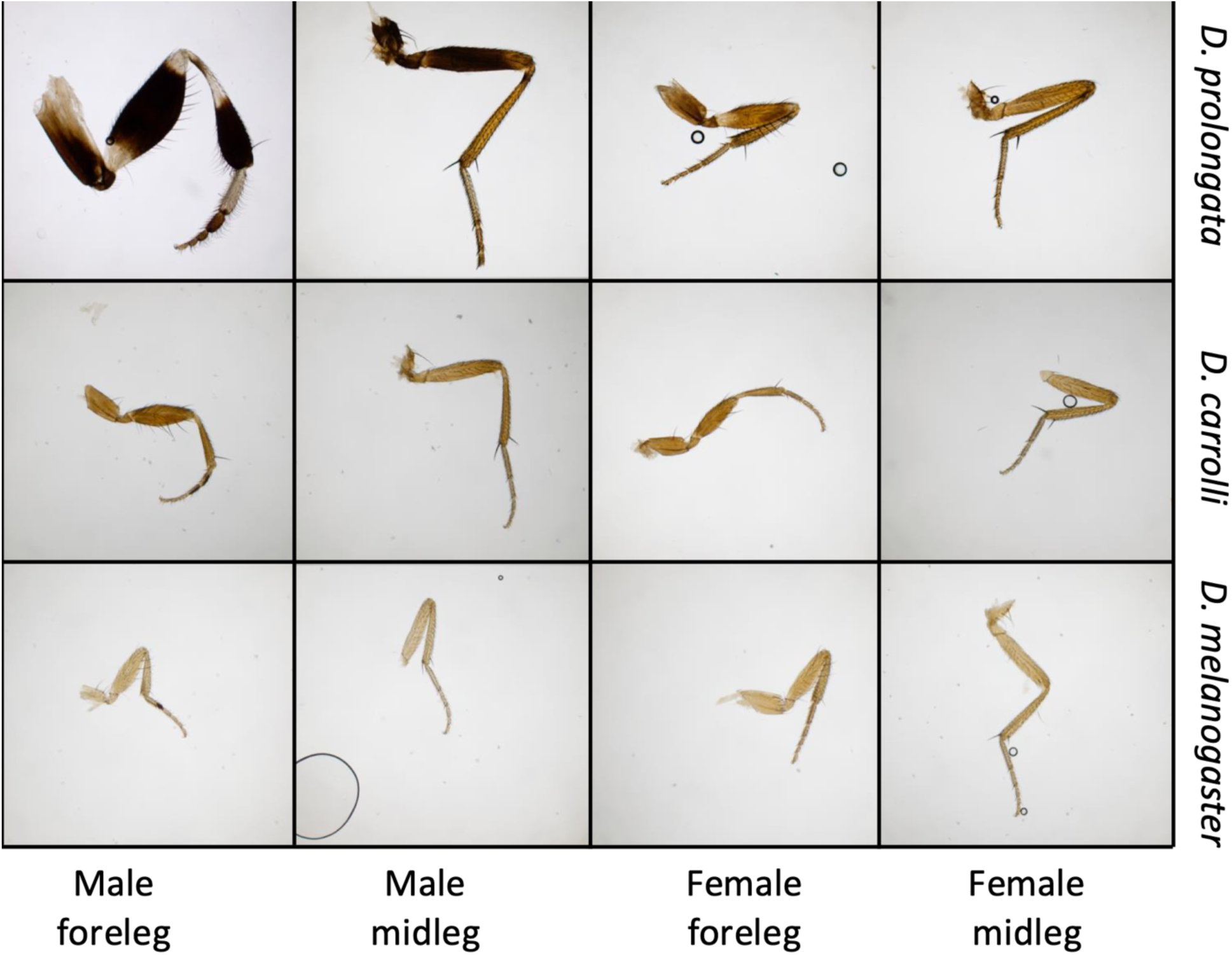
Images of adult fore- and midleg of each species and sex. All were sampled at juvenile stages (i.e., as imaginal discs) for RNA sequencing. All images taken at 40x magnification.

Understanding the evolution of sexually dimorphic traits requires examining how sex-biased gene expression (SBGE) changes throughout development, as both the number and direction of biased genes shift over time (Ingleby et al. 2015). In *D. prolongata* males, exaggerated forelegs originate from imaginal discs containing on average 16% more cells than midleg discs by the end of larval development. This difference is absent in *D. prolongata* females and in either sex of *D. carrolli* (Luecke and Kopp 2019). The recent availability of the genome of both species (Kim et al. 2021; Luecke et al. 2024) now permits probing the genetic basis of this trait. Previous work on adult sex-specific pheromones in these two species revealed that out of 526 male-biased genes, 53 showed such bias in only *D. prolongata* (Luo et al. 2025), fewer than predicted by models such as genic capture. This SBGE was potentially driven by a transposable element (‘honghaier’) that moved regulatory binding sites for *dsx* and *bric-a-brac 1* (*bab1*) to new loci in *D. prolongata* in front of a group of five cuticular hydrocarbon genes (Luo et al. 2025).

We profiled SBGE in the developing leg imaginal discs in *D. prolongata*, *D. carrolli*, and *D. melanogaster*. We focused on two time points in the final larval instar: an early stage (18 hours before wandering) immediately before the initiation of dimorphic cell proliferation, and a late stage (8 hours before wandering) after the difference is established (Luecke and Kopp 2019). Within and between species we compared sex-specific imaginal disc expression profiles between both fore-and midlegs. We found that 38.1% of mapped genes show sex-biased gene expression in the foreleg of *D. prolongata*, with 283 (0.03% of genes) exhibiting at least a log_2_ fold change, including genes previously associated with sex-biased body size and leg development in *D. melanogaster*. We functionally evaluated candidate genes via knockdown in *D. melanogaster* and found one candidate, *grain* (*grn*), that partially phenocopied the *D. prolongata* leg size phenotype. We discuss our results in the context of how temporal changes in SBGE contribute to the evolution of sexual dimorphism.

## Methods

### Drosophila husbandry

The *D. prolongata* population for this experiment was derived from ∼200 individuals collected by JPF in the Sa Pa region of Vietnam in 2018. This population had been maintained under laboratory conditions (∼50 individuals/bottle across multiple bottles, and offspring mixing in each generation to minimize inbreeding, 14:10hr light-dark cycles at 18°C and 60% relative humidity), with overlapping generations at a census number well above the 200 founder individuals to preserve genetic diversity (Perdigón Ferreira and Lüpold 2022). The *D. carrolli* population was obtained from A. Kopp and O. Barmina who collected the founder flies in Brunei in 2003. This population was also maintained and expanded at 18°C for over two generations in our laboratory prior to the experiment. The *D. melanogaster* was Canton-S, normally maintained at 25°C but also reared at 18°C for two generations before the for experiment for consistency with the other two species.

### Larval staging, sampling and dissection

For sample collection and staging, each species was kept in plastic cages (∼30cm^3^) at high adult density with yeast paste on orange-juice agar in Petri dishes as oviposition substrate. To stage larvae from their hatching time, plates were removed from cages and yeast paste sieved through a fine mesh. Eggs (but not larvae) were subsequently placed on grape-juice agar plates for collection. From these egg collections, freshly hatched first-instar larvae were collected hourly, for three hours. Larvae were considered newly hatched ±30 minutes of each collection time, and were allowed to develop in cornmeal-yeast-agar media (for recipe see ref <u>1)</u> until sampling for dissection.

Larval dissections for leg imaginal discs occurred at two developmental time points during the third larval instar: an early stage (18 hours before larval wandering; 6 days + 6 hours after hatching) and a late stage (8 hours before larval wandering; 6 days + 16 hours after hatching). These timepoints were selected based on Luecke and Kopp (Luecke and Kopp 2019), who demonstrated that sexually dimorphic cell proliferation in *D. prolongata* foreleg discs begins in the latter half of the third (final) instar. The early timepoint captured the stage immediately before the onset of quantifiable dimorphism, when signals are active but differences in cell number are still minimal. The late timepoint occurred just after dimorphic proliferation had begun, when a clear male bias in the cell number of the foreleg disc is detectable. By comparing these two timepoints, we aimed to profile gene expression both at the initiation of sex-biased development and during active sexual differentiation.

Larval sex was determined in the morning of dissections based on the presence of a larger gonad in male larvae (large transparent “disc” visible in the larval fat body), after which larvae were allowed to re-enter food to minimize stress caused by removal from the substrate. Larvae were dissected in chilled 1× PBS solution to reduce induction of a transcriptional stress response. During dissection, the anterior portion of the cuticle was removed before inverting the larval body and extracting the brain with the leg imaginal discs attached. Foreleg and midleg imaginal discs were then detached from the brain and stored separately in 200μL of RNAlater (Thermo Fisher Scientific) at -80°C until extraction.

As leg imaginal discs are very small tissues, ∼30 discs from ∼15-20 individuals were used for each unique biological sample to ensure sufficient RNA for library preparation. For each species/sex/stage/tissue we collected five independent biological replicates for individual library preparation and sequencing. We opted for “bulk” RNA sequencing due to both biological relevance and practical considerations. As shown in Figure 1, sexual size dimorphism in adult *D. prolongata* legs is visible across most leg segments even if most pronounced in the femur and tibia. As the larval imaginal discs sampled contain the precursor cells for all these future adult structures (Kojima 2004; Schubiger et al. 2012), bulk sampling captures the transcriptome of the entire developing organ where these differences originate. Additionally, single-cell RNA sequencing remains cost-prohibitive for a study of this scale and presents ongoing analytical challenges for robustly quantifying expression differences.

Unlike for *D. melanogaster* and *D. carrolli*, *D. prolongata* males lack a sex comb. The development of this sex-limited structure in other species occurs during pupal stages, after the larval periods we examined (Barmina and Kopp 2007; Atallah et al. 2009; Tanaka et al. 2009; Tanaka et al. 2011), which ensures that expression changes are not due to differences in sex comb development between legs or species. Gene expression differences between leg segments at these early stages are known to be modest (Barmina et al. 2005), confirming that bulk sampling is suitable for detecting the early, broad-scale changes in gene expression that underlie the initial growth differences we aimed to study, suggesting that bulk sampling is suitable for detecting the early, broad-scale changes in gene expression that underlie the initial growth differences we aimed to study.

### RNA extraction, library preparation and sequencing

RNA extractions were done using Qiagen RNeasy spin columns (CAT# 74106) with DNase treatment (CAT# 79256). To remove RNAlater, we added 1mL of cold 1× PBS to the 1.5mL micro-centrifuge tubes containing 200μL of RNAlater and centrifuged (4°C) to separate discs from solution. Supernatant was pipetted off to leave the imaginal discs and as little RNAlater as possible, and the recommended protocol for the Qiagen RNeasy kit was followed from that point. Following assessment of samples for quantity and purity, total RNA was shipped to the Centre d’expertise et de services, Génome Québec, for additional quality checks (bio-Analyzer), library preparation and sequencing. NEB Stranded mRNA libraries with Nextera adaptor sequences were prepared for each biological sample and sequenced with an Illumina NovaSeq6000 to an average of ∼50 million clusters/sample (100bp paired-end reads). To achieve this sample depth, each library was sequenced twice, split between two runs for which all (multiplexed) samples were done jointly on a single lane within a run.

### Functional test of candidate genes in D. melanogaster using RNAi knockdown

To explore potential effects of candidate gene expression on leg morphology, we used the bipartite UAS-RNAi Gal4 system in *D. melanogaster* to titrate gene expression in the developing legs. Before crossing our candidate UAS-RNAi to the *Pen^NP6333^-Gal4*, we crossed *Pen^NP6333^-Gal4* to a *UAS-GFP.NLS* strain and observed moderate GFP expression in third-instar leg discs, suggesting moderate GAL4 expression during development in our target tissues. *Pen^NP6333^-Gal4* (introgressed into the Samarkand wild-tyoe background marked with *w^-^*), UAS-GFP (BDSC# 4775), *CG30457* UAS-RNAi (BDSC# 62960), *CG13285* UAS-RNAi (BDSC# 53680), *dysfusion* (*dysf)* UAS-RNAi (BDSC# 35010), *grain* (*grn*) UAS-RNAi (BDSC# 33746), *bric-a-brac1* (*bab1*) UAS-RNAi (BDSC# 57410), *Sox box protein 15* (*Sox15*) UAS-RNAi (BDSC# 57264), and *CG9896* UAS-RNAi (BDSC# 42587) were allowed to lay in vials for 48 hours to keep larval density low. All RNAi lines were part of the *Drosophila* Transgenic RNAi project (TRiP; Perkins et al. 2015). Unmated females and males were collected from vials for 6 days, and then RNAi males were crossed to *Pen^NP6333^-Gal4* females (all crosses were performed reciprocally). Females in these crosses were allowed to lay eggs for 48 hours before being flipped into backup vials for another 48 hours of oviposition. Vials were incubated at both 25^°^C, or 28^°^C, as the UAS-Gal4 allows for temperature-mediated titration of expression.

Offspring from each cross were collected after eclosion and cuticle sclerotization and stored in 70% ethanol until imaging. Flies were imaged using a Leica MZ7.5 scope using a Leica IC90.E camera. For imaging, flies were dissected in mounting solution (70% glycerol in PBS, with a small amount of phenol as a bacteriostatic). Adult fore- and midlegs were removed and imaged separately, and thorax was imaged laterally and measured from behind the tip of the scutellum to the notch created from the margin of the humeral callus and the prescutum. Each leg had length and width measured for the femur, as well as thorax length. Images were measured using ImageJ (Rueden et al. 2017).

### Bioinformatic and statistical analysis

After initial quality control with fastQC (version and flags for all software available in Table S1), pairs of raw sequence reads were trimmed of adaptor sequences and low-quality positions using BBduk (Bushnell 2021). First and second runs for the same samples were concatenated prior to alignment. Reads were then indexed and mapped within each species to assembled annotations:

*D. prolongata* (GCA_036346975.1) and *D. carrolli* (GCA_018152295.1) from Luecke et al. (Luecke et al. 2024), and *D. melanogaster* (assembly version 6.23) from FlyBase.org. Reads were counted at the gene level using STAR, excluding multi-mapped reads (Dobin and Gingeras 2016), before importing them into R (version 4.4.2; R Core Team 2023) for analysis. A principal component analysis (PCA) was performed both within and between species for the count data (with gene lists filtered for genes orthologous in all three species) using the ‘rlog’ variance stabilization in DESeq2. These patterns were consistent even when using smaller subsets of varying genes (from 500 to 2000 in 250 gene increments).

Counts were modelled gene-by-gene within each species using a generalised linear mixed model (GLMM) with a negative binomial distribution (quadratic dispersion parameterization), as implemented in glmmTMB (Brooks et al. 2017). The full model was specified with gene counts as a function of developmental stage (early vs. late), leg tissue (fore-vs. midleg imaginal disc), and sex and their interactions as our fixed effects. We modelled day of sampling effects nested within stage as a random effect. Log transformed sample specific size factors from the DESeq2 function ‘estimateSizeFactors’ were used as model offsets (Love et al. 2014). Expression estimates and contrasts were computed using emmeans (Lenth et al. 2018), focusing primarily on sex differences and their interactions with species, time or tissue. Estimated contrasts were regularized using the ashr library (Stephens et al. 2016). Counts modelling using DESeq2 confirmed that model estimates (but not standard errors on estimates) were similar to those from the mixed models, but the mixed model had moderated fold changes (Figure S1; S2).

### Multivariate examination of changes in sex-biased gene expression

To assess the relative degrees of similarity in SBGE in forelegs compared to midlegs within species, as well as the similarity of SBGE in forelegs between species, we conducted an analysis of vector correlation and ratio of magnitudes of expression. This analysis was based on adapted scripts from both Zinna et al. (Zinna et al. 2018) and Scott et al. (Scott et al. 2022), which allowed us to quantify differences in the direction or magnitude of SBGE for sets of genes, even if individual genes did not ‘significantly’ differ. This approach is informative when exploring expression changes for biological variables such as pathways involved with growth. Vector correlation was calculated as 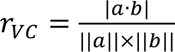 where vectors ***a*** and ***b*** each represent vectors of male-female contrasts with regularization. *r_VC_* values closer to 1 suggest similar direction of sex-biased expression changes, while values near 0 suggest little association. For our comparison of vector magnitudes, we calculated the ratio of magnitudes (ℓ^2^ norm) of vectors of male-female SBGE contrasts using 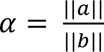. Where higher *α* values suggest greater magnitudes of SBGE in *a* relative to *b* and an *α* of 1 would mean that the magnitude of SBGE is similar. For this study we had two primary analyses: 1) *a* = species 1 foreleg SBGE and *b* = species 2 foreleg SBGE, and 2) *a* = species 1 foreleg SBGE and *b* = species 1 midleg SBGE.

### Comparison of sexual dimorphism to sex-biased gene expression

We examined the relationship between sexual size dimorphism calculated from log transformed morphological measures as (male femur width – female femur width) + (male femur length – female femur length) for both foreleg and midleg separately, in all three species, and sex-biased gene expression. We estimated the relationship between this SSD value and the magnitude of SBGE for all genes, as well as for genes with an estimated expression with a log_2_ fold change with 95% confidence intervals non-overlapping with the log_2_ fold change threshold in *D. prolongata*. The magnitude for SBGs was calculated as ℓ^2^-norms of relevant vectors of effect sizes. To assess uncertainty in these estimates, we performed non-parametric bootstrapping, resampling with replacement 10,000 times and calculating the 95% quantiles of the distribution to generate standard percentile intervals. We also compared these SD values with the number of SBGs that had confidence intervals not overlapping zero after ashr regularization, as well as a comparison to our genes that showed a log_2_ fold (confidence on our estimates non-overlapping with +1 or -1) change after ashr regularization.

### Molecular evolution of grain

Genomes of *D. prolongata* (GCA_036346975.1), *D. carrolli* (GCA_018152295.1), and *D. rhopaloa* (GCA_018152115.1) were downloaded from NCBI to both explore sequence changes in *grn*, as well as to look for the previously identified transposable element honghaier (Luo et al. 2025) up- or downstream of any of our candidates. The annotated *grn* sequence from *D. melanogaster* was reciprocally blasted to all three genomes to identify the *grn* location, and that corresponding sequence was extracted from each genome using bedtools getfasta (Quinlan and Hall 2010) and aligned using MUSCLE (Edgar 2004a; Edgar 2004b). This was done with each exon of *grn* individually. To search for changes in the predicted enhancer of *grn* between species, sequences were first filtered for tandem repeats using tandem repeat finder (Benson 1999) with flags recommended for SCRMshaw inputs, and then predictions were made using SCRMshaw (Asma et al. 2024). SCRMshaw was trained on all available *Drosophila* enhancers found at: https://github.com/HalfonLab/dmel_training_sets. SCRMshaw was run using default settings on the *D. prolongata*, *D. carrolli*, and *D. rhopaloa* genomes, and predicted enhancer outputs were searched for the appearance of *grn* as the nearest downstream gene, using UNIX grep. To identify downstream targets of *grn*, we extracted the nearest genes to identified CHIPseq targets of grain reported by the mod-encode project (ENCSR909QHH), which were then used as SBGE vectors for our *r_VC_* and *α* analyses above.

The honghaier transposable element has been recently shown to contain both *dsx* and *bab1* binding sites (Luo et al. 2025), and likely influences species-specific changes in expression in *D. prolongata*. To search for the honghaier transposable element near genes of interest, we extracted the CDS region using the coordinates in the annotations of *D. prolongata*, *D. carrolli*, and *D. rhopaloa*, including 10,000bp up- and downstream using bedtools getfasta, and blasted the honghaier sequence against each of these loci to identify potential insertion sites.

## Results

### Expression profiles show species and sex-specific clustering, with little evidence for clustering based on specific leg imaginal disc, or developmental stage

First, we determined if the unique foreleg morphology of male *D. prolongata* (Figure 1) correlated with broad-scale changes in the transcriptional profile of male foreleg imaginal discs relative to females, other species, or midlegs with their largely absent trait exaggeration. In a principal component analysis on all samples, 8008 genes were unambiguously orthologous between all three species and thus suitable for potential inclusion. Using the 1000 most variable genes as features, PC1 accounted for 48% of variance in gene expression and clearly separated all three species. PC2 separated *D. carrolli* out from *D. melanogaster* and *D. prolongata* and accounted for 25% of the variance (Figure 2). Sex differences were associated with a combination of PC3 (5%) and PC4 (2%) (Figure 2). We further evaluated the transcriptome-wide expression profiles within species (Figure 3, Figure S3). While the effects of sex and developmental stage appeared to group together in the PCA, we saw only modest evidence of broad-scale differences between the first and second leg imaginal discs, even in *D. prolongata* males.

**Figure 2:**
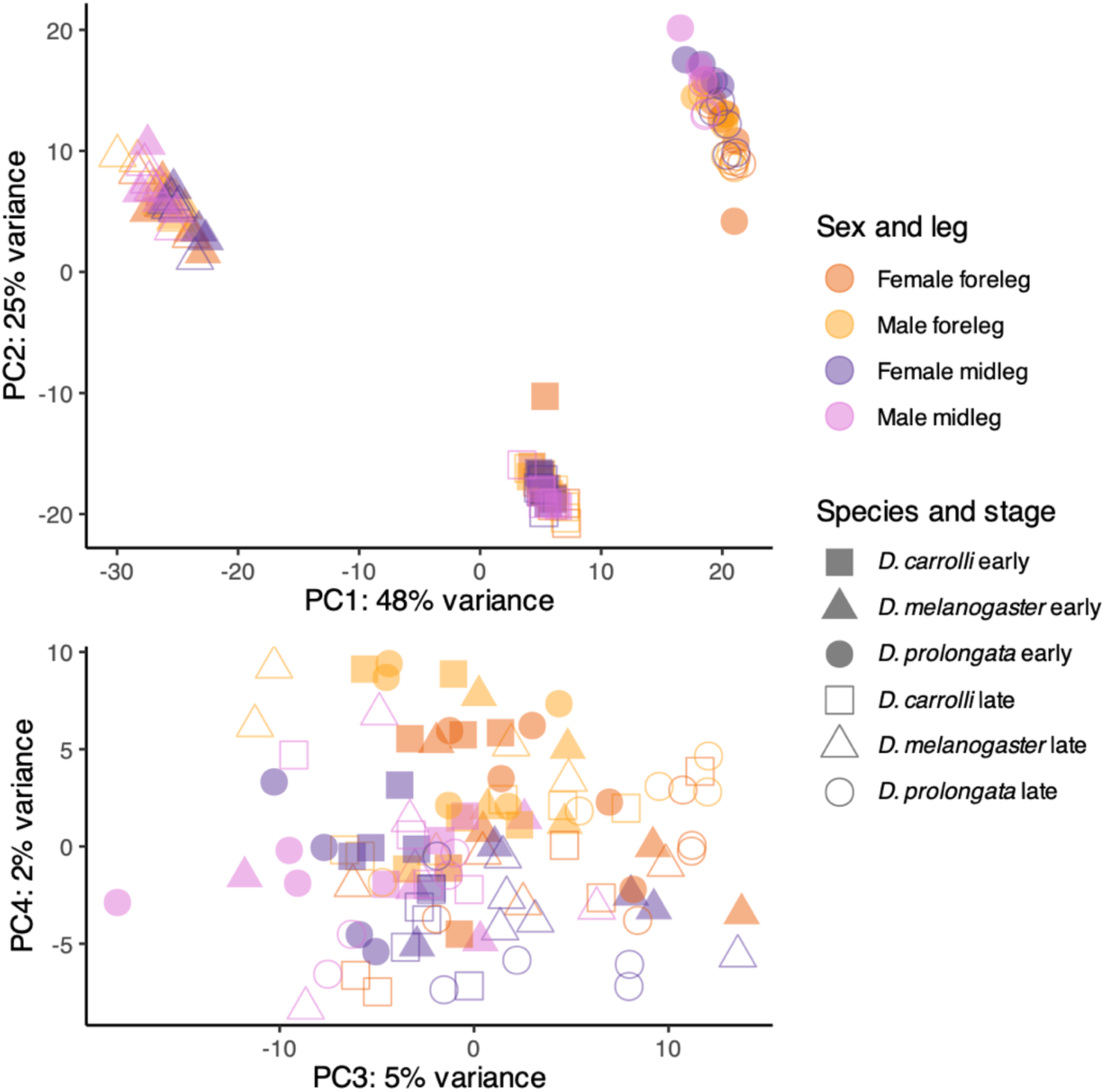
PCA projection of developmental stage, leg, sex, and species for the top 1000 most variable genes in *D. prolongata*, *D. carrolli*, and *D. melanogaster.* A regularized log (rlog) was applied to the count matrices prior to computing the PCs.

**Figure 3:**
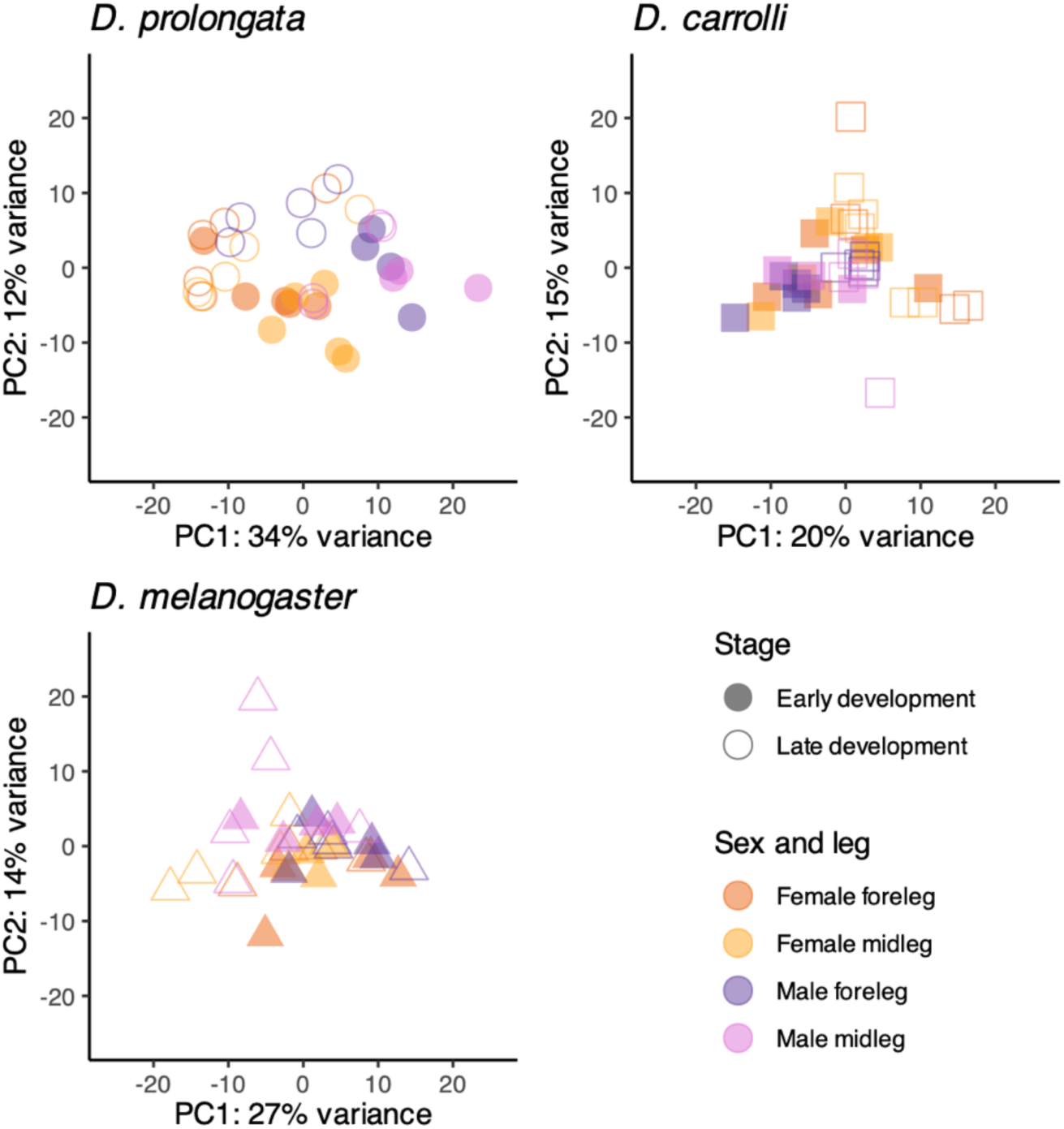
PCA projection of developmental stage, leg, and sex within species for the top 1000 most variable genes in *D. prolongata*, *D. carrolli*, and *D. melanogaster* with a regularized log transformation applied.

### Sex-biased gene expression in D. prolongata forelegs is species-specific

Sex-biased genes were more numerous in *D. prolongata* than in either *D. carrolli* or *D. melanogaster* (Table 1; Figure 4 left; S4; S5; Supplemental File 1). When gene were filtered for at least a minimum of a two-fold expression difference (minimum based on 95% confidence intervals of contrasts) between the sexes in the foreleg but not in the midleg, *D. prolongata* showed the greatest number of genes that were either male-(58 compared to 7 *D. carrolli* and 9 *D. melanogaster*) or female-biased (129 compared to 34 *D. carrolli* and 17 *D. melanogaster*).

**Figure 4:**
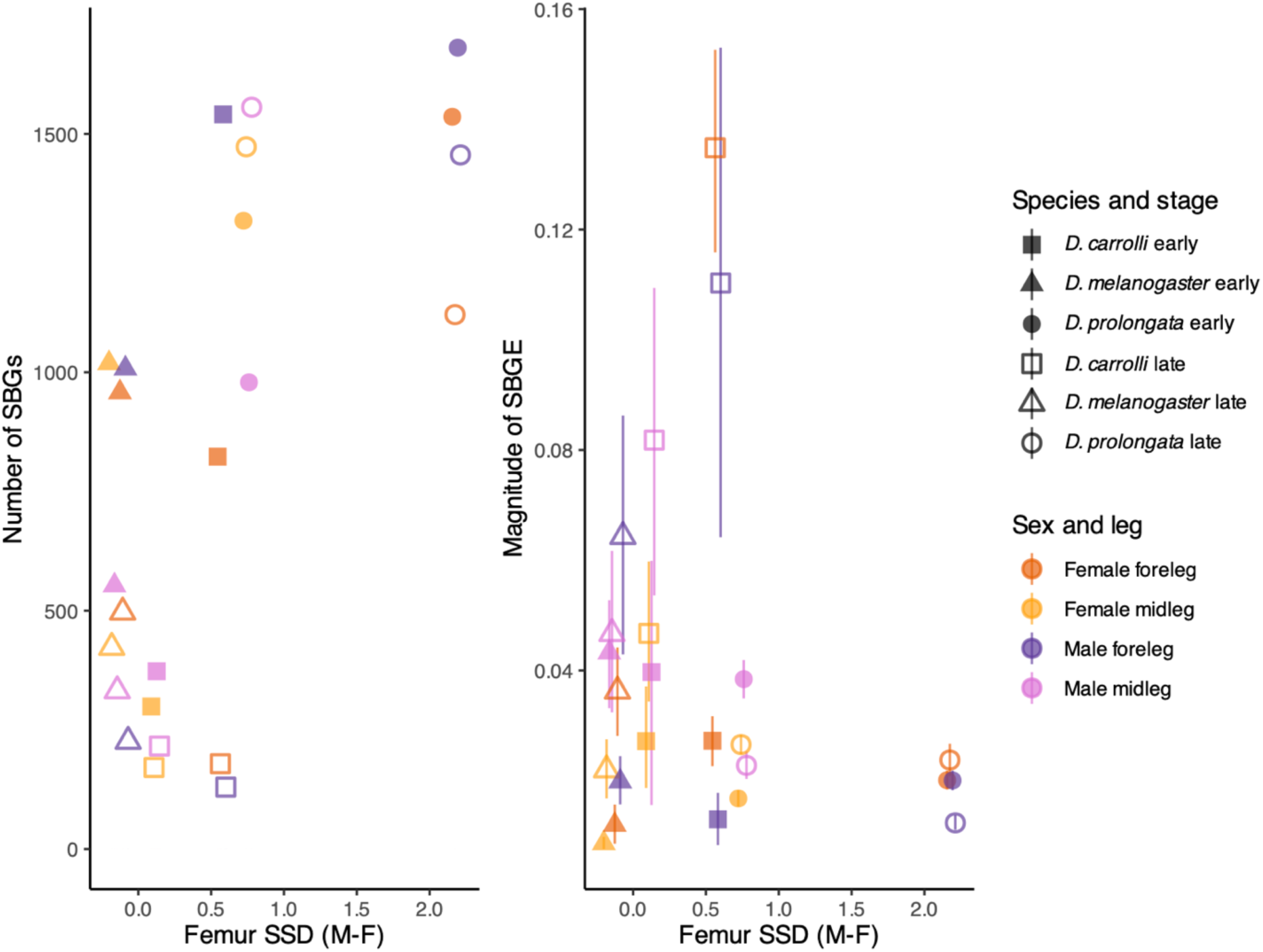
Relationship of the magnitude of expression (SBGE) changes (left) or number (right) for sex-biased genes (SBG) and degree of adult sexual size dimorphism, measured as female – male log transformed trait size. Magnitude and number of SBGs represent the number (or magnitude) of male (female) biased genes at each developmental stage in each species.

**Table 1:** Number of genes with a minimum two-fold change in each species for each sex (Log_2_ cut-off based on upper- or lower-bound of the 95% confidence interval). Foreleg genes are those that are not log_2_ fold changed in midleg as well. Numbers in parentheses represent sex-biased genes with regularized 95% CIs that do not overlap zero.

| Species | Female-biased | Male-biased | Female-biased<br>foreleg | Male-biased<br>foreleg |
| --- | --- | --- | --- | --- |
| *D. prolongata* | 110 (2820) | 173 (2845) | 129 (1920) | 58 (2248) |
| *D. carrolli* | 35 (1062) | 19 (1745) | 34 (930) | 7 (1590) |
| *D. melanogaster* | 21 (1559) | 39 (1412) | 17 (1189) | 9 (1123) |

Using the genes identified as showing SBGE in *D. prolongata*, we evaluated to what degree the number and overall magnitude of expression differences in the developing leg imaginal discs is associated with the degree of phenotypic sexual size dimorphism (SSD) in the adult legs within and between species (Figure 4). We did not observe a strong relationship between magnitude of expression differences in SBGs and the most sexually dimorphic structures. For instance, male *D. prolongata* femurs, the trait with the greatest SSD, exhibited similar magnitudes of SBGE as most other tissues, with the highest magnitudes being in *D. carrolli* in both sexes during late development (Figure 4 right). However, the absolute number of genes that showed SBGs was positively related to phenotypic SSD, and this pattern held for both the full list of SBGs (Figure 4 left) and a subset of the genes that showed a minimum two-fold difference between the sexes (Figure S6 left).

The differentially expressed genes of primary interest for our study were those with sex-biased gene expression only in the foreleg imaginal discs of *D. prolongata*, and without a parallel change either in the corresponding midlegs or in any leg of the other two species. Using custom interaction contrasts, we identified a set of genes with these attributes (Supplemental File 1). We observed sets of genes with both male- and female-biased directions of expression differences in the foreleg imaginal discs of *D. prolongata*, including several genes whose orthologs influence organ-specific size and shape.

### RNAi knockdowns of a subset of our candidates shows changes to femur width and length as well as a qualitatively D. prolongata-like phenotype in D. melanogaster femurs

We identified all SBGs with a minimum two-fold expression difference in *D. prolongata* forelegs without concomitant magnitude of differences in midlegs (Supplemental file 2). We refined the list of candidate genes using FlyBase for known or presumed roles in organ size or sex-specific effects. We knocked down expression of orthologs of these genes in the leg imaginal discs of *D. melanogaster* during larval development and examined changes in lengths and widths of adult legs. We chose to knock down *grn*, *dysf*, *CG30457*, *CG13285*, *sox15*, *bab1*, and *otp* based on their SBGE patterns (Figure 5). Among these, we observed two *grn* knockdown individuals (<1% of total number of flies) with highly enlarged femur widths, qualitatively resembling legs of *D. prolongata* (Figure 6). This phenocopy was not observed in any other individuals from other crosses (More than 1000 individuals phenotyped).

**Figure 5:**
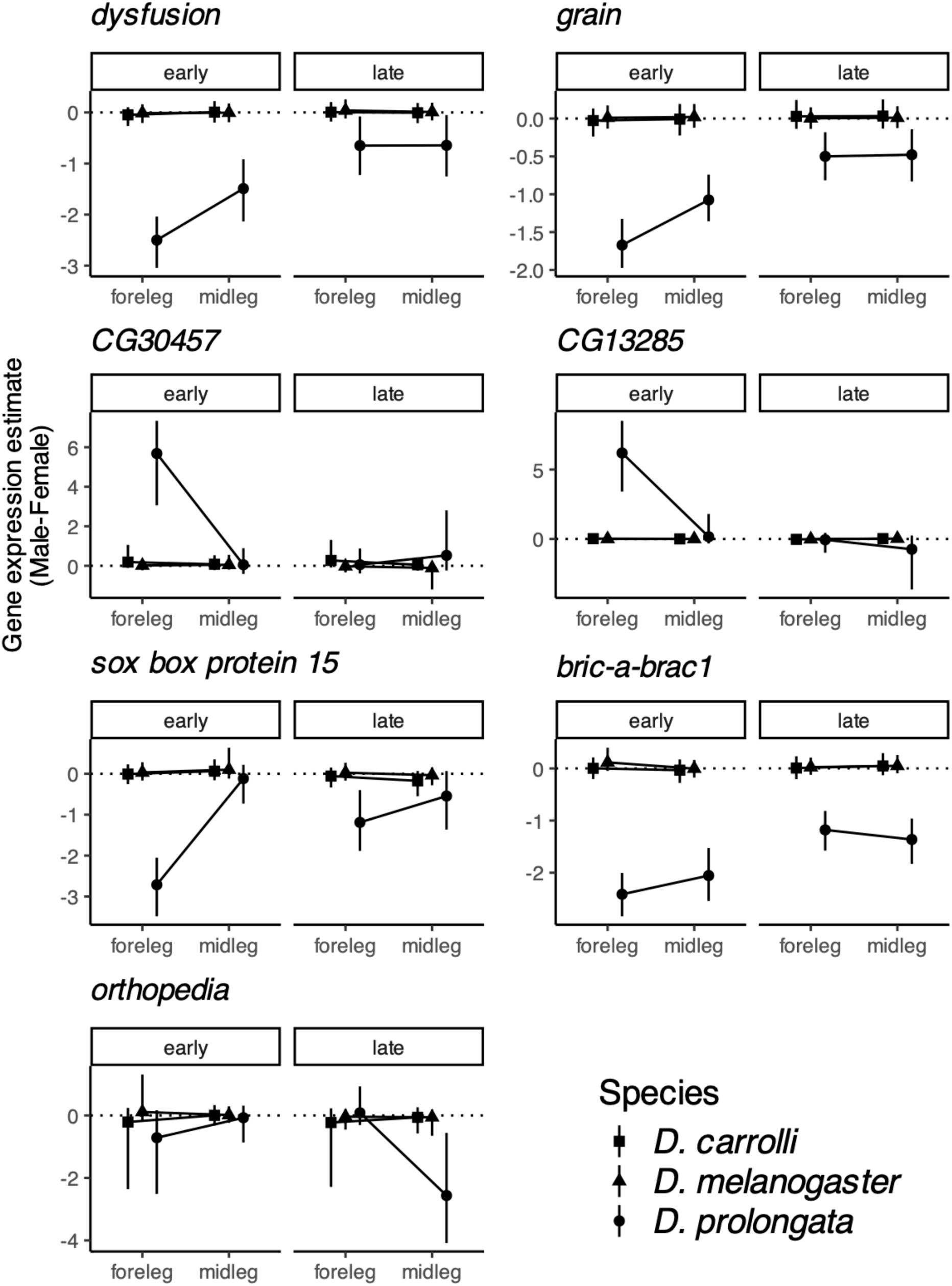
Interaction plots of sex-biased gene expression for candidate genes chosen for RNAi-mediated gene knockdown. Male-female contrast (regularized estimates) of gene expression (*log_2_*) on the y-axis and both stage and leg are shown separate for each gene and species.

**Figure 6:**
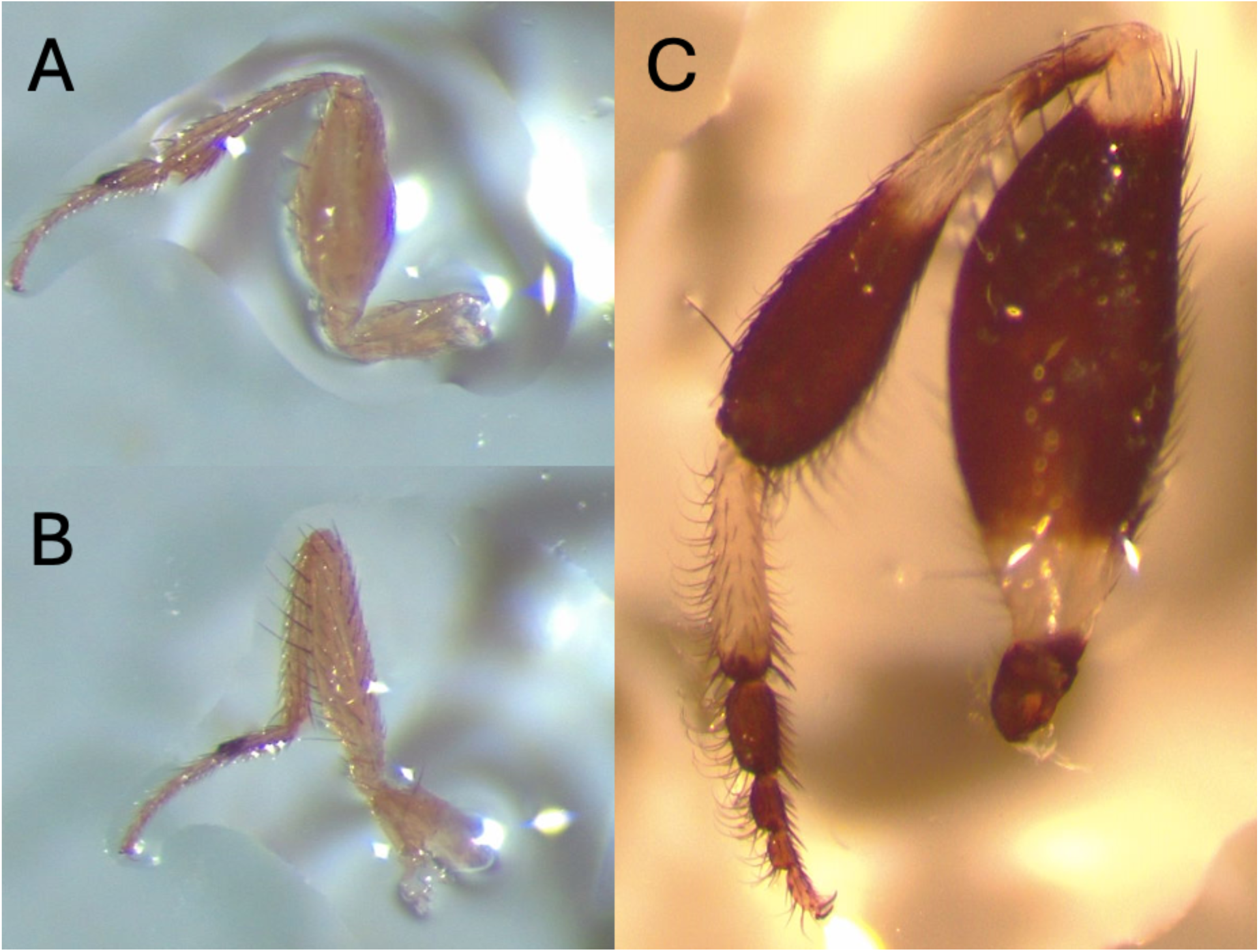
Male foreleg in A) *grn* knockdown *D. melanogaster* B) control *D. melanogaster*. C) wild-type *D. prolongata.* All images are at taken at 50x magnification.

Here, we focus on the 28°C treatment throughout our results as expression knock-down in these treatments is stronger, but we saw similar trends at 25°C (Figure 7). Forefemur length decreased in male knockdowns of *grn*, and *sox15* (Table S2; Figure S7) and in female knockdowns of *bab1*, *grn*, *otp*, and *sox15* (Table S2; Figure S7). Likewise, forefemur width decreased in *dysf* males and *CG13285* females, but it increased in *bab1* and *sox15* females (Table S2; Figure S7).

**Figure 7:**
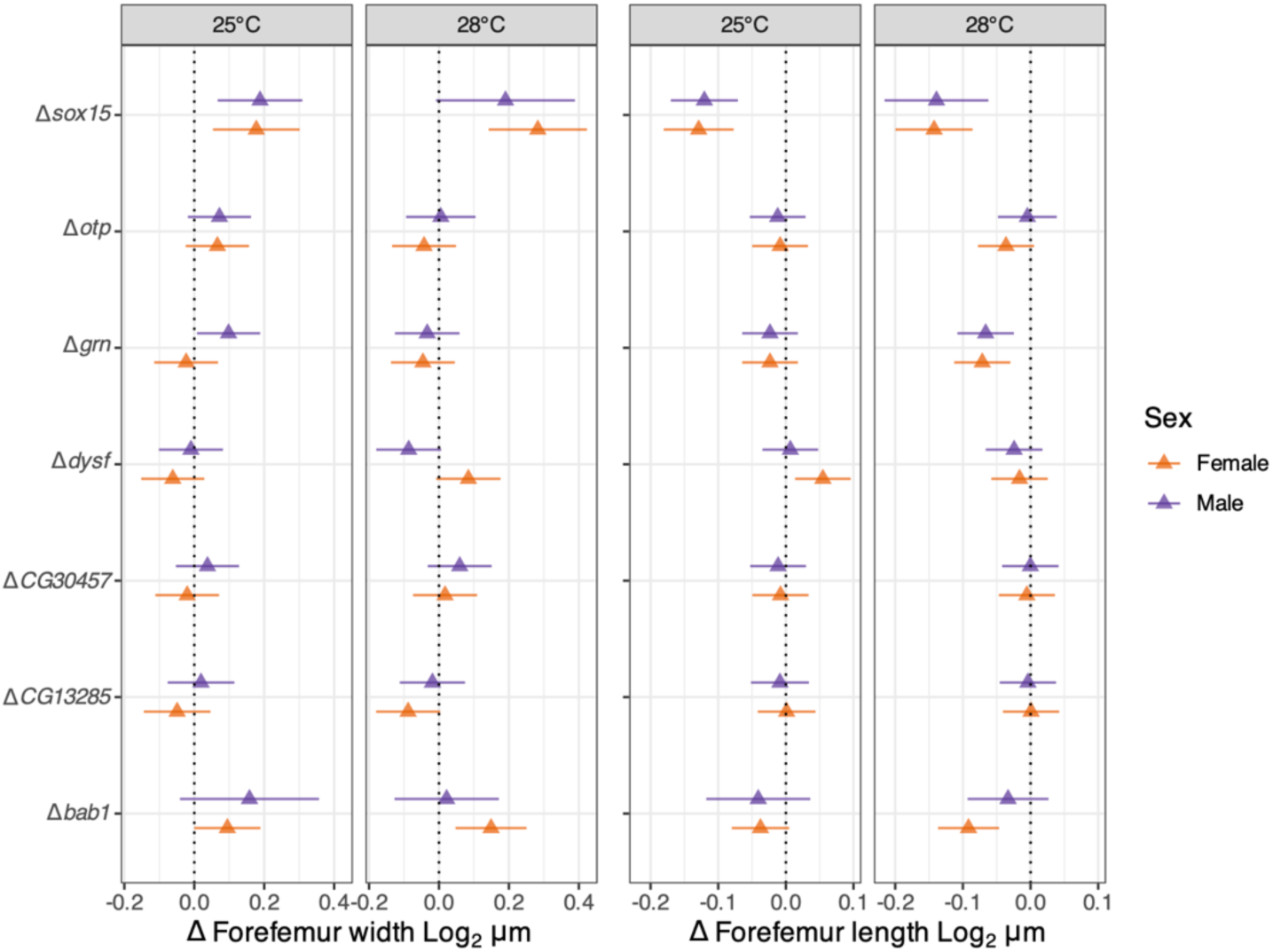
Changes to leg traits size relative adjusted for thorax length for UAS-RNAi strains crossed to NP3666-GAL4 contrasted with control crosses from experiments performed at 25°C and 28°C. Contrasts calculated from model estimates.

We next explored leg size changes while accounting for body (thorax) size. We observed some reduction in thorax length relative to controls in the UAS-RNAi offspring of *bric-a-brac1* (*bab1*), and *orthopedia* (*otp*) in males, as well as in *bab1*, *CG13285*, *dysfusion* (*dysf*), and *sox box protein 15* (*sox15*) in females (Table S2; Figure S7). Adjusting for body size, forefemur length decreased in both sexes in *sox15* and *grn* knockdowns, and in just females in *bab1* and *otp* (Figure 7; Table S3). Forefemur width was reduced in males and increased in females when *dysf* was knocked down but increased in both sexes when *sox15* was knocked down (Figure 7; Table S4).

### Grain is well conserved in the closest relatives of D. prolongata

Given our observations of knock-downs with *grain*, a GATA transcription factor involved in growth and metabolism (Brown and Castelli-Gair Hombría 2000; Kokki et al. 2021), we examined *grn* for modifications of its protein coding region that could be linked to changes in gene function. Exon alignment of *grn* between *D. prolongata*, *D. carrolli*, *D. rhopaloa,* and *D. melanogaster* show high levels of conservation. The exon 4 GATA binding site had 100% amino acid conservation, and the exon 5 GATA binding site had only a few synonymous substitutions present in *D. rhopaloa*, *D. carrolli*, and *D. prolongata* compared to *D. melanogaster*. Changes outside the binding-domain were also uncommon between *D. prolongata* and *D. carrolli* (amino acid changes: 1/169 exon 1, 0/95 exon 2, 4/276 exon 3, 0/47 exon 4, 0/47 exon 5, 0/62 exon 6, 0/62 exon 7). Cis-regulatory module predictions from SCRIMshaw revealed no modules that were unique to *D. prolongata* that were also predicted to regulate *grn*. When predicted modules were searched for the gene identifier *grn*, all predicted modules were identified in *D. carrolli* or *D. rhopaloa* as well.

### The transposable element honghaier occurs up or down stream in most genes of interest in multiple species, with seven being exclusive to D. prolongata

Looking 10,000bp up- and downstream, we identified seven of the candidate genes of interest that had a blast hit for the honghaier transposable element exclusively in *D. prolongata*. These genes were *smydA*, *rst*, *CG11378*, *CG14075*, *CG14356*, *CG32564*, and *CG9411*, all of which showed SBGE surpassing our minimum two-fold change threshold (Supplemental file 2) but did not stand out relative to our other genes of interest.

### Signalling pathways and candidate sex-biased genes appear to be expressed in similar direction and magnitude between all three species

As the observed differential expression involving many developmental and growth genes, we examined changes across tissues and species in both direction and ratio of magnitudes for vectors of sex-biased expression (i.e. contrast of expression differences between males and females).

Changes in the overall ratio of magnitudes of expression for our minimum two-fold candidate SBGs between fore- and midleg imaginal discs within species was highest in *D. prolongata* early (1.6×) but only slightly so compared to *D. melanogaster* early in development (0.94×), although within the upper bounds of our 95% intervals for magnitude ratios of randomly drawn genes (Figure 8). During the later developmental time-period, the relative increase in magnitude remained modest, but greater than 95% random intervals in *D. prolongata* (1.3×; Figure 8).

**Figure 8:**
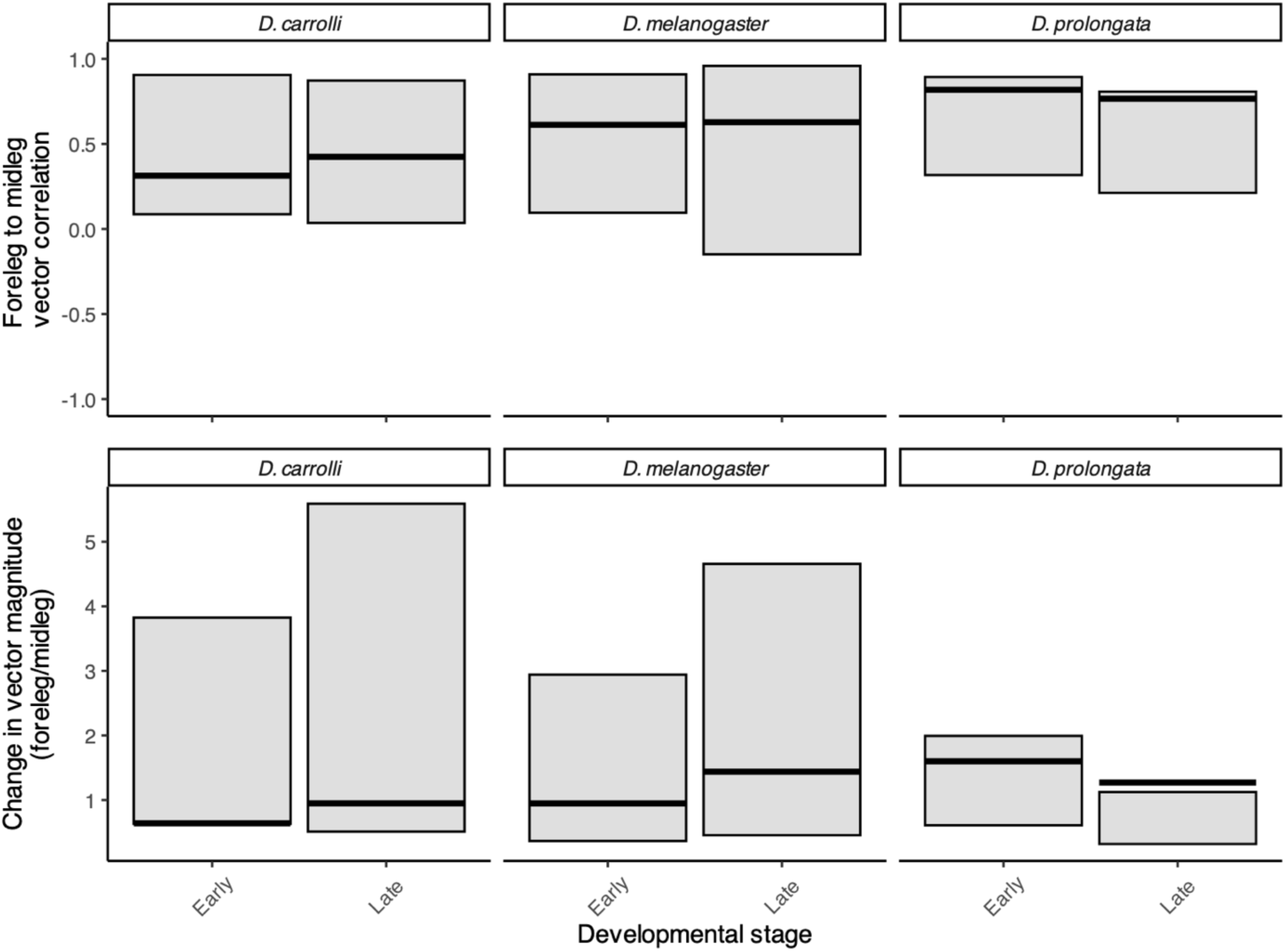
Changes in direction and magnitude of expression for vectors of SBGs, between foreleg and midleg imaginal discs. Based on ∼110 SBGs with a minimum two-fold difference (one *log_2_* unit) in *D. prolongata,* that also share orthologs in both *D. carrolli* and *D. melanogaster*. Top row shows correlations of vectors (degree of shared direction of effects) for SBGE in foreleg compared to midleg within all three species. Bottom row is the relative change in magnitude of SBGE in foreleg relative to midleg imaginal discs within each species for the same set of genes.

When we partitioned the genes with a minimum two-fold change between the sexes into male- and female-biased genes, we observed a striking pattern (Figure 9). We observed a negative vector correlations (for SBGE between foreleg and midleg) for the male-biased genes early in development (r_VC_ = -0.39), but positive ones later (r_VC_ = 0.73). This pattern was not observed in the other species, and below our randomly drawn vector range (Figure 9). We did not see this in female-biased gene vector correlations in any of the three species (Figure S8).

**Figure 9:**
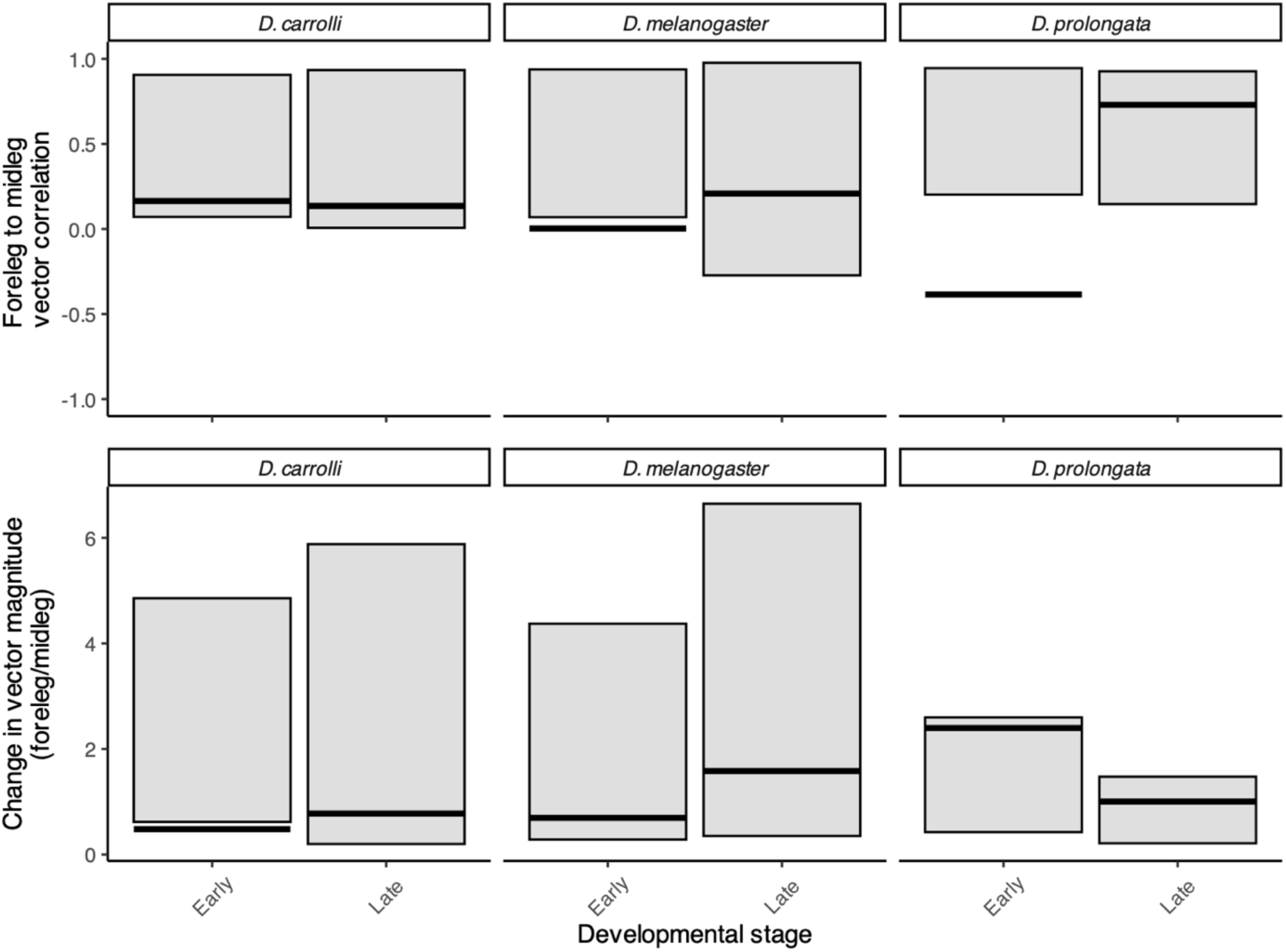
Vector correlation and change in magnitude of male-biased genes showing at minimum two-fold expression change in *D. prolongata* forelegs, but not midlegs or in either of the other species, for a total of 36 genes. Top row shows direction of SBGE in foreleg compared to midleg within all three species. Bottom row is the magnitude of SBGE in foreleg relative to midleg within each species.

Between species, vectors for sex-biased genes with a minimum of two-fold change were only weakly to moderately correlated with each other (*D. prolongata* / *D. carrolli* early r_VC_ = 0.08, late r_VC_ = 0.31; *D. prolongata* / *D. melanogaster* early r_VC_ = -0.12, late r_VC_ = -0.10). The correlations were somewhat similar between forelegs in all species in both stages and were within the range of randomly drawn vectors. As expected, the change in the magnitude of the vector of SBGE for our candidate genes is highest in *D. prolongata* forelegs compared to either *D. carrolli* (8.85× greater early, and 5.78× greater late) and *D. melanogaster* (6.71× greater early, and 7.59× greater late) forelegs, while the expression magnitudes in forelegs were similar between *D. melanogaster* and *D. carrolli* (Figure 10). When we explored male- and female-biased genes separately between species, male-biased genes showed an 11.34· magnitiude of expression in *D. prolongata* early compared to *D. carrolli*, and 23.31× increase compared to *D. melanogaster*, but not later in development (Figure S9). Female-biased genes also showed a difference in magnitude when comparing *D. prolongata* to *D. carrolli* both early (7.71×) and late (6.83×) in development, and when *D. prolongata* is compared to *D. melanogaster* late in development (7.72×). Female-biased genes also showed a negative correlation in the foreleg early in development compared to *D. carrolli* (r_VC_ = -0.22) compared to randomly drawn genes (Figure S10). For *D. carrolli* female-biased genes, this represents a change in direction of the correlation, with later developmental genes becoming positively correlated with *D. prolongata* (r_VC_ = 0.24).

**Figure 10:**
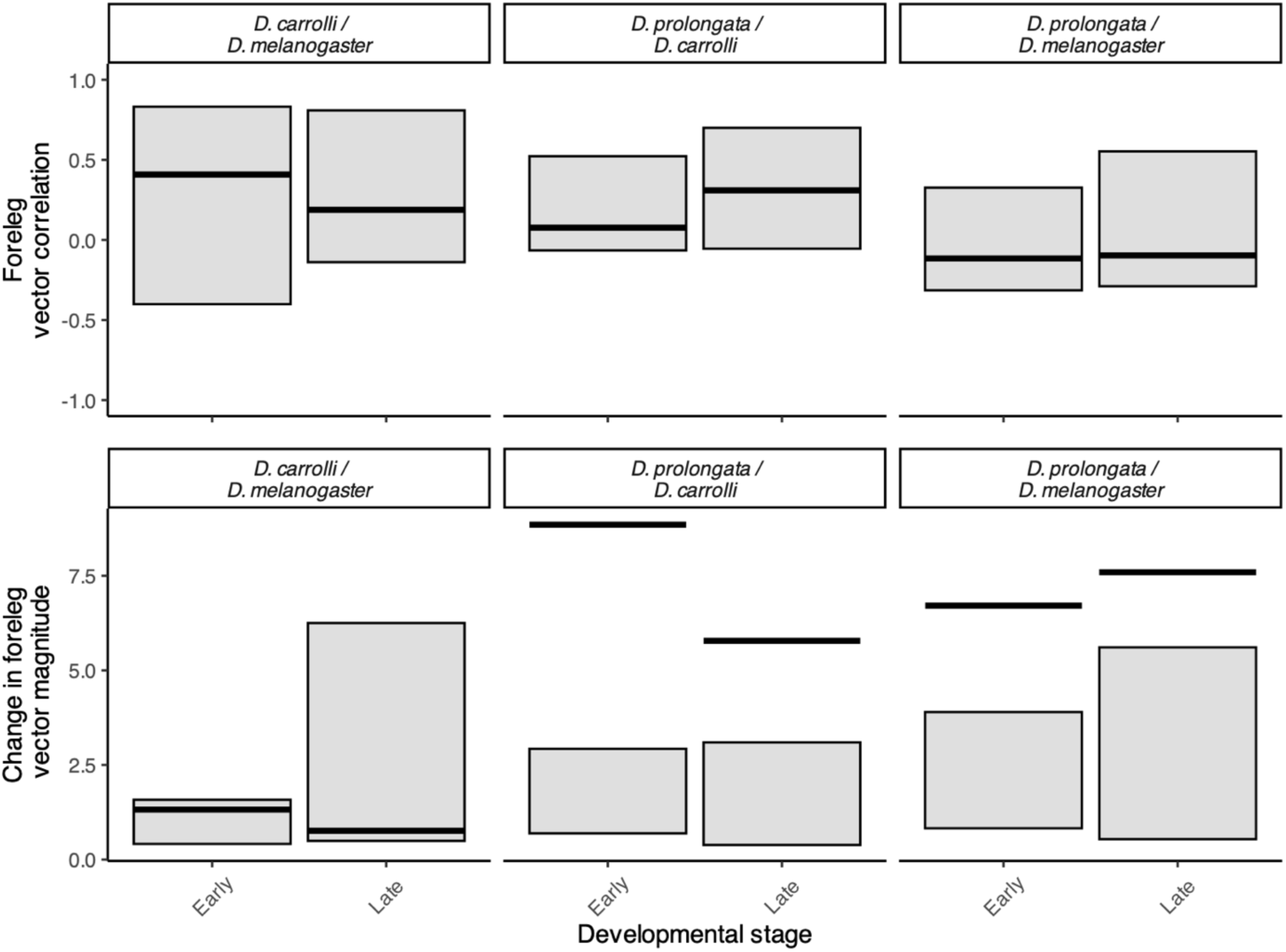
Degree of shared direction and changes in magnitude of vectors for sex-biased genes between species in the foreleg imaginal disc. Based on ∼80 SBGs with a minimum two-fold difference (one *log_2_* unit) in *D. prolongata,* with orthologs in *D. carrolli* and *D. melanogaster*. Top row shows degree of shared direction (expressed as a correlation) of SBGE in foreleg between *D. prolongata* and the other species, the first column shows *D. carrolli* compared to *D. melanogaster* as a point of reference. Bottom row is the change in magnitude of SBGE in foreleg between each species. Values greater than one represent a relative increase in magnitude of SBGE for this set of genes in *D. prolongata* relative to the other species.

To verify that this magnitude difference was not explained by selecting genes based on their expression differences in *D. prolongata*, we repeated the above analysis using the two-fold differentially expressed *D. melanogaster* genes based on the same thresholds as for *D. prolongata* candidates. This yielded many fewer genes with a two-fold change in expression (10 genes meeting this threshold in *D. melanogaster*), with magnitudes closer to or less than one between the species (*D. prolongata* / *D. carrolli* early = 1.32× greater, late =1.28× greater; *D. prolongata* / *D. melanogaster* early = 0.55×, late = 0.40×; Figure S11).

### Sex-biased expression in key signalling pathways are broadly similar in direction and magnitude between all three species

We also examined changes among genes within key signaling pathways underlying tissue growth within and between species. Since the insulin signaling pathway (InS) has previously been implicated with exaggerated trait growth (Emlen et al. 2006; Emlen et al. 2012), we specifically explored this pathway both between tissues within species, and between species in the foreleg (Figure S12). Within species, vector correlation was similar between foreleg and midleg both early (r_VC_ = 0.63) and late (r_VC_ = 0.74) in *D. prolongata*, and magnitudes were within our 95% random vector intervals. The one exception is in *D. carrolli*, where there is a clear negative correlation between fore- and midlegs for InS (r_VC_ = -0.32). In *D. prolongata*, InS magnitude was 1.45× greater in forelegs compared with midlegs early and 0.74× later in development. These magnitudes were similar or lower than *D. melanogaster* and *D. carrolli,* and within our distribution of random gene magnitudes (Figure S12). For other signaling pathways, vector correlations between foreleg and midleg within species were consistently high and outside our 95% interval of random gene draws, while the ratios of magnitudes between foreleg and midleg imaginal discs were generally close to 1, meaning magnitude of SBGE between legs was approximately equivalent (Figure S12-S18).

Comparing SBGE in the InS pathway in forelegs between species, we further observed weak to moderate positive vector correlations between *D. prolongata* and *D. carrolli* (early r_VC_ = 0.11; late r_VC_ = 0.65) and between *D. prolongata* and *D. melanogaster* (early r_VC_ = 0.05; late r_VC_ = 0.16), as well as magnitudes around 1 between *D. prolongata* and both *D. carrolli* (early = 1.1×; late = 0.85×) and *D. melanogaster* (early = 1.52×; late = 1.99×; Figure S19). For all other signalling pathways we explored, SBGE r_VC_ were positively correlated between species, and magnitude was around 1 (Figure S20-S25).

As Grn is a transcription factor, we explored its putative targets (defined as the nearest gene to reported Grn binding sites reported by the mod-encode project ENCSR909QHH). The magnitude of sex-biased gene expression for Grn targets was greater in *D. prolongata* (1.63× greater than early *D. carrolli*; 1.06× greater than late *D. carrolli*; 2.43× greater than early *D. melanogaster*; 1.34× greater in late *D. carrolli*), exceededing our 95% random gene vectors in early *D. melanogaster* (Figure S26). Changes in the magnitude of SBGE for putative Grn targets in *D. carrolli* is lower than *D. melanogaster* both early (0.67×) and late (0.78×) in development.

## Discussion

The evolution of sexual dimorphism and its relationship to changes in SBGE has been of interest, and a topic with many open questions (Parsch and Ellegren 2013; Ingleby et al. 2015; Grath and Parsch 2016). A simple ‘good genes’ model would suggest a potentially small number of genes under selection in sexually selected traits, whereas various polygenic models, including the genic capture model, would suggest a substantial fraction of the genome mediating evolutionary changes for these traits (Rowe and Houle 1996; Pomiankowski and Møller 1997). While an indirect assessment, our results are most consistent with some genetic architecture intermediate to these extremes. If the sexual dimorphism was mediated by small expression changes in a substantial fraction of the genome, we would predict that the male *D. prolongata* foreleg would have separated out clearly in the PCA, which it did not (Figure 2, 3). At the same time, we do observe a small set of genes with very large expression differences, exceeding patterns observed in the other two species (Figure 5, 10), including *grain* whose knockdown in *D. melanogaster* (with very low penetrance) qualitatively recapitulates the *D. prolongata* foreleg (Figure 6). In previous studies on sexually dimorphic tissue samples (as opposed to whole-body sampling), small proportions of genes were observed to differentially expressed (Wilkinson et al. 2013; Zinna et al. 2018; Toubiana et al. 2021), although statistical power remains an issue for many such studies. We found a positive relationship between the number of SBGs and degree of phenotypic dimorphism between species, but no strong association between magnitude of expression differences and phenotypic dimorphism in the adult (Figure 4). This correlation between number of genes and dimorphism despite such a small proportion of highly differentially expressed genes between sexes and tissues suggests a role of regulatory genes in dimorphic evolution, as co-option of regulatory genes may contribute to substantial phenotypic change with a modest number of genes. Indeed, our analyses revealed changes in relatively few transcription factors (TF) that may be at least partially responsible for the maintenance of the sexually exaggerated forelegs in *D. prolongata*, such as *grn*.

Little is known about developmental pathways through which *grn* might modulate adult morphology, as many mutations are homozygous lethal. Previous work has demonstrated that tissue-specific loss of function of *grn* results in shorter, wider femurs (Brown and Castelli-Gairm Hombría 2000), and our functional results of *grn* knockdowns, that can resemble *D. prolongata* support this finding. *grn* has been implicated in spiracle formation (Brown and Castelli-Gair Hombría 2000), sugar metabolism (Kokki et al. 2021), and neuron specification (Garces and Thor 2006), suggesting involvement of this transcription factor across multiple developmental processes. In the closest relatives of *D. prolongata*, *D. carrolli* and *D. rhopaloa*, we found no major changes in the protein sequence of *grn* and 100% conservation in the GATA binding domains between all three species and *D. melanogaster*. We also identified a strong transcriptional female bias for *dysf* and *sox15*, with corresponding phenotypic changes to femur width or length when knocked down in *D. melanogaster* (Figure 7). *Grn*, *dysf*, and *sox15* have all been demonstrated to be regulated via notch signaling (Garces and Thor 2006; Miller et al. 2009; Córdoba and Estella 2020), a pathway known to influence leg morphology and joint formation in *D. melanogaster* (Rauskolb and Irvine 1999). However, our results do not indicate large changes to the magnitude of Notch SBGE in *D. prolongata* forelegs relative to those of *D. carrolli* and *D. melanogaster* (Figure S13; S20). We saw changes in the expression magnitude of potential targets of Grn in the forelegs of *D. prolongata* compared to *D. carrolli* and *D. melanogaster* (Figure S26), but the large number (3407) of putative Grn targets from the mod-encode data, may include many false positives, and needs to be further evaluated.

Beyond size differences, *D. prolongata* legs also demonstrate dimorphic colouration (Setoguchi et al. 2014). Previous work has implicated the two *bric-a-brac* paralogs (*bab1* and *bab2*) as regulators of sex-specific pigmentation in the *D. melanogaster* thorax (Kopp et al. 2000; Williams et al. 2008). These studies have further found that *bab1/2* is under control of the female isoform of *dsx* (Kopp et al. 2000), as well as responsible for leg development and tarsal specification (Godt et al. 1993). *dsx* has also been implicated as a regulator of sexually dimorphic tibia growth in the gazelle dung beetle *Digitonthophagus gazella* (Rohner et al. 2021). We found female biased expression in both *dsx* and *bab1* greater in *D. prolongata* than in *D. melanogaster* or *D. carrolli*, and knockdowns of *bab1* in *D. melanogaster* resulted in sex-specific changes in both femur width and length in *D. melanogaster* (Figure 7), suggesting a potential role of the *dsx*/*bab1* gene cascade in the legs of *D. prolongata*. We caveat this result by the fact that this requires further functional work such as immunofluorescence.

We also explored broad changes in gene pathway correlations for both direction and magnitude of sets of SBGs. We found male-biased genes demonstrate a direction of SBGE between the legs that was negatively correlated, and distinct from randomly drawn genes in *D. prolongata* but not the other species (Figure 9). We also observed a much higher magnitude of SBGE in foreleg imaginal discs of *D. prolongata* relative to *D. carrolli* and *D. melanogaster* (Figure 10). This again suggests that the extent of SBGE in the forelegs may be important for sexually dimorphic morphology in *D. prolongata*. Despite these results, the overall magnitude of SBGE did not seem to relate to the degree of morphological sexual dimorphism, unlike the absolute number of genes (Figure 5). Understanding the functional relationship between magnitude of SBGE as it relates to sexual dimorphism remains an open and critical question.

In other systems with sexually exaggerated traits, growth pathway differences have been associated with sexual dimorphism; specifically in insulin signaling (reviewed in ref (Emlen et al. 2006)). We explored changes in insulin signaling (and other pathways) between legs within species, and in forelegs between species. We found surprisingly little difference in direction and magnitude of the SBGs (Figure 10; S15; S20). Our sampling occurred shortly before, and shortly after, the initiation of sexually dimorphic cells proliferation in *D. prolongata* forelegs (Luecke and Kopp 2019). However, whether this particular period of growth showed sex-specific responses to insulin signaling remains unknown. Another potential reason for the lack of transcriptional difference for these growth signals, despite differences in proliferation may reflect the specific traits we examined. Many of the studies implicating growth signaling look at novel traits (horns), whereas *D. prolongata* forelegs are existing traits that have been exaggerated, which may influence the extent to which important growth signalling pathways can be altered.

Our work adds to our growing knowledge of how sex-specific expression of a largely shared genome can modulate sexual dimorphism. *Drosophila prolongata* is an ideal model for studying the evolution of sexual dimorphism and addressing outstanding questions in sexual selection due to its ease of rearing large samples, its expression of a phylogenetically novel trait, and the genetic tools available in the closely related *D. melanogaster*.

## Acknowledgements

We would like to thank Artyom Kopp for *D. carrolli* and *D. rhopaloa*, David Luecke, Yige Luo, Artyom Kopp for draft genome annotations of *D. carrolli* and *D. prolongata*. We would also like to thank the undergraduate volunteers who aided in imaging of flies. This work was funded by a National Science and Engineering Research Council Discovery grant to ID, a Swiss National Science Foundation (grant 310030_197651) to SL, a URPP Evolution in Action: From Genomes to Ecosystems to AM, JPF, SL, and an Ontario Graduate Scholarship to TA.

## References

Alexander Dobin, Thomas R. Gingeras. 2016. Optimizing RNA-Seq mapping with STAR. Methods Mol. Biol. 1415:245–262.

Asma H, Liu L, Halfon MS. 2024. SCRMshaw: Supervised cis-regulatory module prediction for insect genomes. PLOS ONE 19:e0311752.

Atallah J, Liu NH, Dennis P, Hon A, Godt D, Larsen EW. 2009. Cell dynamics and developmental bias in the ontogeny of a complex sexually dimorphic trait in *Drosophila melanogaster*. Evol. Dev. 11:191–204.

Baker BS, Ridge KA. 1980. Sex and the single cell. I. On the action of major loci affecting sex determination in Drosophila melanogaster. Genetics 94:383–423.

Barmina O, Gonzalo M, McIntyre LM, Kopp A. 2005. Sex- and segment-specific modulation of gene expression profiles in *Drosophila*. Dev. Biol. 288:528–544.

Barmina O, Kopp A. 2007. Sex-specific expression of a HOX gene associated with rapid morphological evolution. Dev. Biol. 311:277–286.

Benson G. 1999. Tandem repeats finder: a program to analyze DNA sequences. Nucleic Acids Res. 27:573–580.

Brooks M E, Kristensen K, Benthem K J,van, Magnusson A, Berg C W, Nielsen A, Skaug H J, Mächler M, Bolker B M. 2017. glmmTMB balances speed and flexibility among packages for zero-inflated generalized linear mixed modeling. R J. 9:378–400.

Brown S, Castelli-Gair Hombría J. 2000. *Drosophila grain* encodes a GATA transcription factor required for cell rearrangement during morphogenesis. Development 127:4867–4876.

Bushnell B. 2021.BBTools: BBMap short read aligner, and other bioinformatic tools.

Casasa S, Moczek AP. 2018. Insulin signalling’s role in mediating tissue-specific nutritional plasticity and robustness in the horn-polyphenic beetle Onthophagus taurus. Proc. R. Soc. B Biol. Sci. 285:20181631.

Córdoba S, Estella C. 2020. Role of notch signaling in leg development in *Drosophila melanogaster*. Notch Signaling in Embryology and Cancer: Notch Signaling in Embryology. p. 103–127. 10.1007/978-3-030-34436-8_7

Cox RM, Cox CL, McGlothlin JW, Card DC, Andrew AL, Castoe TA. 2017. Hormonally mediated increases in sex-biased gene expression accompany the breakdown of between-sex genetic correlations in a sexually dimorphic lizard. Am. Nat. 189:315– 332.

Crumière AJJ, Khila A. 2019. Hox genes mediate the escalation of sexually antagonistic traits in water striders. Biol. Lett. 15:20180720.

Darwin C. 1871. The descent of man, and selection in relation to sex. D. Appleton

Dobzhansky T. 1931. Interaction between female and male parts in gynandromorphs of *Drosophila simulans*. Wilhelm Roux Arch. Für Entwicklungsmechanik Org. 123:719– 746.

Edgar RC. 2004a. MUSCLE: multiple sequence alignment with high accuracy and high throughput. Nucleic Acids Res. 32:1792–1797.

Edgar RC. 2004b. MUSCLE: a multiple sequence alignment method with reduced time and space complexity. BMC Bioinformatics 5:113.

Ellegren H, Parsch J. 2007. The evolution of sex-biased genes and sex-biased gene expression. Nat. Rev. Genet. 8:689–698.

Emlen DJ. 2008. The evolution of animal weapons. Annu. Rev. Ecol. Evol. Syst. 39:387–413.

Emlen DJ, Szafran Q, Corley LS, Dworkin I. 2006.Insulin signaling and limb-patterning: candidate pathways for the origin and evolutionary diversification of beetle ‘horns.’ *Heredity* 97:179–191.

Emlen DJ, Warren IA, Johns A, Dworkin I, Lavine LC. 2012. A mechanism of extreme growth and reliable signaling in sexually selected ornaments and weapons. Science 337:860–864.

Fisher RA. 1930. The genetical theory of natural selection. Oxford, England: Clarendon Press

Garces A, Thor S. 2006. Specification of *Drosophila* aCC motoneuron identity by a genetic cascade involving *even-skipped*, *grain* and *zfh1*. Development 133:1445–1455.

Godt D, Couderc J-L, Cramton SE, Laski FA. 1993. Pattern formation in the limbs of *Drosophila* : *bric à brac* is expressed in both a gradient and a wave-like pattern and is required for specification and proper segmentation of the tarsus. Development 119:799–812.

Gompel N, Kopp A. 2018. *Drosophila (Sophophora) carrolli* n. sp., a new species from Brunei, closely related to *Drosophila (Sophophora) rhopaloa* (Diptera: Drosophilidae). Zootaxa 4434:502–510.

Gotoh H, Miyakawa H, Ishikawa A, Ishikawa Y, Sugime Y, Emlen DJ, Lavine LC, Miura T. 2014. Developmental link between sex and nutrition; *doublesex* regulates sex-specific mandible growth via juvenile hormone signaling in stag beetles. PLOS Genet. 10:e1004098.

Gotoh H, Zinna RA, Warren I, DeNieu M, Niimi T, Dworkin I, Emlen DJ, Miura T, Lavine LC. 2016. Identification and functional analyses of sex determination genes in the sexually dimorphic stag beetle *Cyclommatus metallifer*. BMC Genomics 17:250.

Gowen JW, Fung S-TC. 1957. Determination of sex through genes in a major sex locus in *Drosophila Melanogaster*. Heredity 11:397–402.

Grath S, Parsch J. 2016. Sex-biased gene expression. Annu. Rev. Genet. 50:29–44.

Hérault C, Pihl T, Hudry B. 2024. Cellular sex throughout the organism underlies somatic sexual differentiation. Nat. Commun. 15:6925.

Ingleby FC, Flis I, Morrow EH. 2015. Sex-biased gene expression and sexual conflict throughout development. Exp. Des.

J Patterson. 1938. Aberrant forms in *Drosophila* and sex fifferentiation. Am. Nat. 72:193– 206.

Jahner JP, Lucas LK, Wilson JS, Forister ML. 2015. Morphological outcomes of gynandromorphism in Lycaeides butterflies (Lepidoptera: Lycaenidae). J. Insect Sci. 15:38.

Khila A, Abouheif E, Rowe L. 2009. Evolution of a novel appendage ground plan in water striders is driven by changes in the Hox gene *Ultrabithorax*. PLOS Genet. 5:e1000583.

Kijimoto T, Moczek AP, Andrews J. 2012. Diversification of *doublesex* function underlies morph-, sex-, and species-specific development of beetle horns. Proc. Natl. Acad. Sci. 109:20526–20531.

Kim BY, Wang JR, Miller DE, Barmina O, Delaney E, Thompson A, Comeault AA, Peede D, D’Agostino ER, Pelaez J, et al. 2021. Highly contiguous assemblies of 101 drosophilid genomes. eLife 10:e66405.

Kirkpatrick M. 1996. Good genes and direct selection in the evolution of mating preferences. Evolution 50:2125–2140.

Kirkpatrick M, Ryan MJ. 1991. The evolution of mating preferences and the paradox of the lek. Nature 350:33–38.

Kodric-Brown A, Sibly RM, Brown JH. 2006. The allometry of ornaments and weapons. Proc. Natl. Acad. Sci. 103:8733–8738.

Kojima T. 2004. The mechanism of Drosophila leg development along the proximodistal axis. Dev. Growth DiUer. 46:115–129.

Kokki K, Lamichane N, Nieminen AI, Ruhanen H, Morikka J, Robciuc M, Rovenko BM, Havula E, Käkelä R, Hietakangas V. 2021. Metabolic gene regulation by *Drosophila* GATA transcription factor Grain. PLOS Genet. 17:e1009855.

Kopp A. 2006. Basal relationships in the *Drosophila melanogaster* species group. Mol. Phylogenet. Evol. 39:787–798.

Kopp A. 2011. *Drosophila* sex combs as a model of evolutionary innovations. Evol. Dev. 13:504–522.

Kopp A, Duncan I, Carroll SB. 2000. Genetic control and evolution of sexually dimorphic characters in *Drosophila*. Nature 408:553–559.

Ledón-Rettig CC, Zattara EE, Moczek AP. 2017. Asymmetric interactions between doublesex and tissue- and sex-specific target genes mediate sexual dimorphism in beetles. Nat. Commun. 8:14593.

Lenth R, Singmann H, Love J, Buerkner P. 2018.emmeans: Estimated marginal means, aka Least-Squares Means.

Love MI, Huber W, Anders S. 2014. Moderated estimation of fold change and dispersion for RNA-seq data with DESeq2. Genome Biol. 15:550.

Luecke D, Luo Y, Krzystek H, Jones C, Kopp A. 2024. Highly contiguous genome assembly of *Drosophila prolongata* - a model for evolution of sexual dimorphism and male-specific innovations. G3 GenesGenomesGenetics. 14.10:jkae155.

Luecke DM, Kopp A. 2019. Sex-specific evolution of relative leg size in Drosophila prolongata results from changes in the intersegmental coordination of tissue growth. Evolution 73:2281–2294.

Luo Y, Takau A, Li J, Fan T, Hopkins BR, Le Y, Ramirez SR, Matsuo T, Kopp A. 2025. Regulatory changes in the fatty acid elongase eloF underlie the evolution of sex-specific pheromone profiles in *Drosophila prolongata*. BMC Biol. 23:117.

Mank JE. 2017. The transcriptional architecture of phenotypic dimorphism. *Nat*. Ecol. Evol. 1:0006.

Mathews KW, Cavegn M, Zwicky M. 2017. Sexual dimorphism of body size is controlled by dosage of the X-Chromosomal gene *Myc* and by the sex-determining gene *tra* in *Drosophila*. Genetics 205:1215–1228.

McCullough EL, Miller CW, Emlen DJ. 2016. Why sexually selected weapons are not ornaments. Trends Ecol. Evol. 31:742–751.

McDonald JMC, Nabili P, Thorsen L, Jeon S, Shingleton AW. 2021. Sex-specific plasticity and the nutritional geometry of insulin-signaling gene expression in *Drosophila melanogaster*. EvoDevo 12:6.

Miller SW, Avidor-Reiss T, Polyanovsky A, Posakony JW. 2009. Complex interplay of three transcription factors in controlling the tormogen differentiation program of *Drosophila* mechanoreceptors. Dev. Biol. 329:386–399.

Millington JW, Brownrigg GP, Basner-Collins PJ, Sun Z, Rideout EJ. 2021. Genetic manipulation of insulin/insulin-like growth factor signaling pathway activity has sex-biased effects on *Drosophila* body size. G3 GenesGenomesGenetics 11:jkaa067.

Millington JW, Brownrigg GP, Chao C, Sun Z, Basner-Collins PJ, Wat LW, Hudry B, Miguel-Aliaga I, Rideout EJ. 2021. Female-biased upregulation of insulin pathway activity mediates the sex difference in *Drosophila* body size plasticity. eLife 10:e58341.

Moczek AP, Rose DJ. 2009. Differential recruitment of limb patterning genes during development and diversification of beetle horns. Proc. Natl. Acad. Sci. 106:8992– 8997.

Okada Y, Katsuki M, Okamoto N, Fujioka H, Okada K. 2019. A specific type of insulin-like peptide regulates the conditional growth of a beetle weapon. PLOS Biol. 17:e3000541.

Parsch J, Ellegren H. 2013. The evolutionary causes and consequences of sex-biased gene expression. Nat. Rev. Genet. 14:83–87.

Perdigón Ferreira J, Lüpold S. 2022. Condition-and context-dependent alternative reproductive tactic in *Drosophila prolongata*. Behav. Ecol.

Perkins LA, Holderbaum L, Tao R, Hu Y, Sopko R, McCall K, Yang-Zhou D, Flockhart I, Binari R, Shim H-S, et al. 2015. The Transgenic RNAi Project at Harvard Medical School: Resources and Validation. Genetics 201:843–852.

Perry JC, Harrison PW, Mank JE. 2014. The ontogeny and evolution of sex-biased gene expression in Drosophila melanogaster. Mol. Biol. Evol. 31:1206–1219.

Pomiankowski A, Møller AP. 1997. A resolution of the lek paradox. Proc. R. Soc. Lond. B Biol. Sci. 260:21–29.

Quinlan AR, Hall IM. 2010. BEDTools: a flexible suite of utilities for comparing genomic features. Bioinformatics 26:841–842.

Rauskolb C, Irvine KD. 1999. Notch-mediated segmentation and growth control of the *Drosophila* leg. Dev. Biol. 210:339–350.

Refki PN, Armisén D, Crumière AJJ, Viala S, Khila A. 2014. Emergence of tissue sensitivity to Hox protein levels underlies the evolution of an adaptive morphological trait. Dev. Biol. 392:441–453.

Rice GR, Barmina O, Luecke D, Hu K, Arbeitman M, Kopp A. 2019. Modular tissue-specific regulation of doublesex underpins sexually dimorphic development in *Drosophila*. Development 146:dev178285.

Rideout EJ, Narsaiya MS, Grewal SS. 2015. The sex determination gene *transformer* regulates male-female differences in *Drosophila* Body Size. PLOS Genet. 11:e1005683.

Rohner PT, Casasa S, Moczek AP. 2023. Assessing the evolutionary lability of insulin signalling in the regulation of nutritional plasticity across traits and species of horned dung beetles. J. Evol. Biol. 36:1641–1648.

Rohner PT, Linz DM, Moczek AP. 2021. Doublesex mediates species-, sex-, environment- and trait-specific exaggeration of size and shape. Proc. R. Soc. B Biol. Sci. 288:20210241.

Rohner PT, Pitnick S, Blanckenhorn WU, Snook RR, Bächli G, Lüpold S. 2018. Interrelations of global macroecological patterns in wing and thorax size, sexual size dimorphism, and range size of the Drosophilidae. Ecography 41:1707–1717.

Rowe L, Houle D. 1996. The lek paradox and the capture of genetic variance by condition dependent traits. Proc. R. Soc. Lond. B Biol. Sci. 263:1415–1421.

Rueden CT, Schindelin J, Hiner MC, DeZonia BE, Walter AE, Arena ET, Eliceiri KW. 2017. ImageJ2: ImageJ for the next generation of scientific image data. BMC Bioinformatics 18:529.

Sawala A, Gould AP. 2017. The sex of specific neurons controls female body growth in *Drosophila*. PLOS Biol. 15:e2002252.

Schubiger G, Schubiger M, Sustar A. 2012. The three leg imaginal discs of *Drosophila*: “Vive la différence.” Dev. Biol. 369:76–90.

Scott AM, Dworkin I, Dukas R. 2022. Evolution of sociability by artificial selection*. Evolution 76:541–553.

Setoguchi S, Takamori H, Aotsuka T, Sese J, Ishikawa Y, Matsuo T. 2014. Sexual dimorphism and courtship behavior in *Drosophila prolongata*. J. Ethol. 32:91–102.

Shingleton AW, Das J, Vinicius L, Stern DL. 2005. The temporal requirements for insulin signaling during development in *Drosophila*. PLOS Biol. 3:e289.

Singh B. 1977. Two new and two unrecorded species of the genus Drosophila Fallen (Diptera: Drosophilidae) from Shillong, Meghalaya, India. Proc Zool Soc 30:31.

Stephens M, Carbonetto P, Gerard D, Lu M, Sun L, Willwerscheid J, Xiao N. 2016. ashr: methods for adaptive shrinkage, using empirical Bayes. :2.2–63. Available from: https://CRAN.R-project.org/package=ashr

Tanaka K, Barmina O, Kopp A. 2009. Distinct developmental mechanisms underlie the evolutionary diversification of *Drosophila* sex combs. Proc. Natl. Acad. Sci. 106:4764–4769.

Tanaka K, Barmina O, Sanders LE, Arbeitman MN, Kopp A. 2011. Evolution of sex-specific traits through changes in HOX-dependent *doublesex* Expression. PLoS Biol. 9:e1001131.

Testa ND, Dworkin I. 2016. The sex-limited effects of mutations in the EGFR and TGF-β signaling pathways on shape and size sexual dimorphism and allometry in the *Drosophila* wing. Dev. Genes Evol. 226:159–171.

Toubiana W, Armisén D, Dechaud C, Arbore R, Khila A. 2021. Impact of male trait exaggeration on sex-biased gene expression and genome architecture in a water strider. BMC Biol. 19:89.

Toyoshima N, Matsuo T. 2023. Fight outcome influences male mating success in *Drosophila prolongata*. J. Ethol. 41:119–27

Voje KL. 2016. Scaling of morphological characters across trait type, sex, and environment. Am. Nat. 187:89–98.

Warren IA, Gotoh H, Dworkin IM, Emlen DJ, Lavine LC. 2013. A general mechanism for conditional expression of exaggerated sexually-selected traits. BioEssays 35:889– 899.

Wasik BR, Rose DJ, Moczek AP. 2010. Beetle horns are regulated by the Hox gene, Sex combs reduced, in a species-and sex-specific manner. Evol. Dev. 12:353–362.

Wat LW, Chowdhury ZS, Millington JW, Biswas P, Rideout EJ. 2021. Sex determination gene *transformer* regulates the male-female difference in Drosophila fat storage via the adipokinetic hormone pathway. eLife 10:e72350.

Wilkinson GS, Johns PM, Metheny JD, Baker RH. 2013. Sex-biased gene expression during head development in a sexually dimorphic stalk-eyed fly. PLOS ONE 8:e59826.

Williams TM, Selegue JE, Werner T, Gompel N, Kopp A, Carroll SB. 2008. The regulation and evolution of a genetic switch controlling sexually dimorphic traits in *Drosophila*. Cell 134:610–623.

Yan D, Perrimon N. 2015. *spenito* is required for sex determination in *Drosophila melanogaster*. Proc. Natl. Acad. Sci. 112:11606–11611.

Zhu C, Ming MJ, Cole JM, Edge MD, Kirkpatrick M, Harpak A. 2023. Amplification is the primary mode of gene-by-sex interaction in complex human traits. Cell Genomics 3:100297.

Zinna R, Emlen D, Lavine LC, Johns A, Gotoh H, Niimi T, Dworkin I. 2018. Sexual dimorphism and heightened conditional expression in a sexually selected weapon in the Asian rhinoceros beetle. Mol. Ecol. 27:5049–5072.

