## Supplemental Figures for "Genetic architecture of the developing forelegs of *Drosophila prolongata*; an exaggerated weapon and ornament"

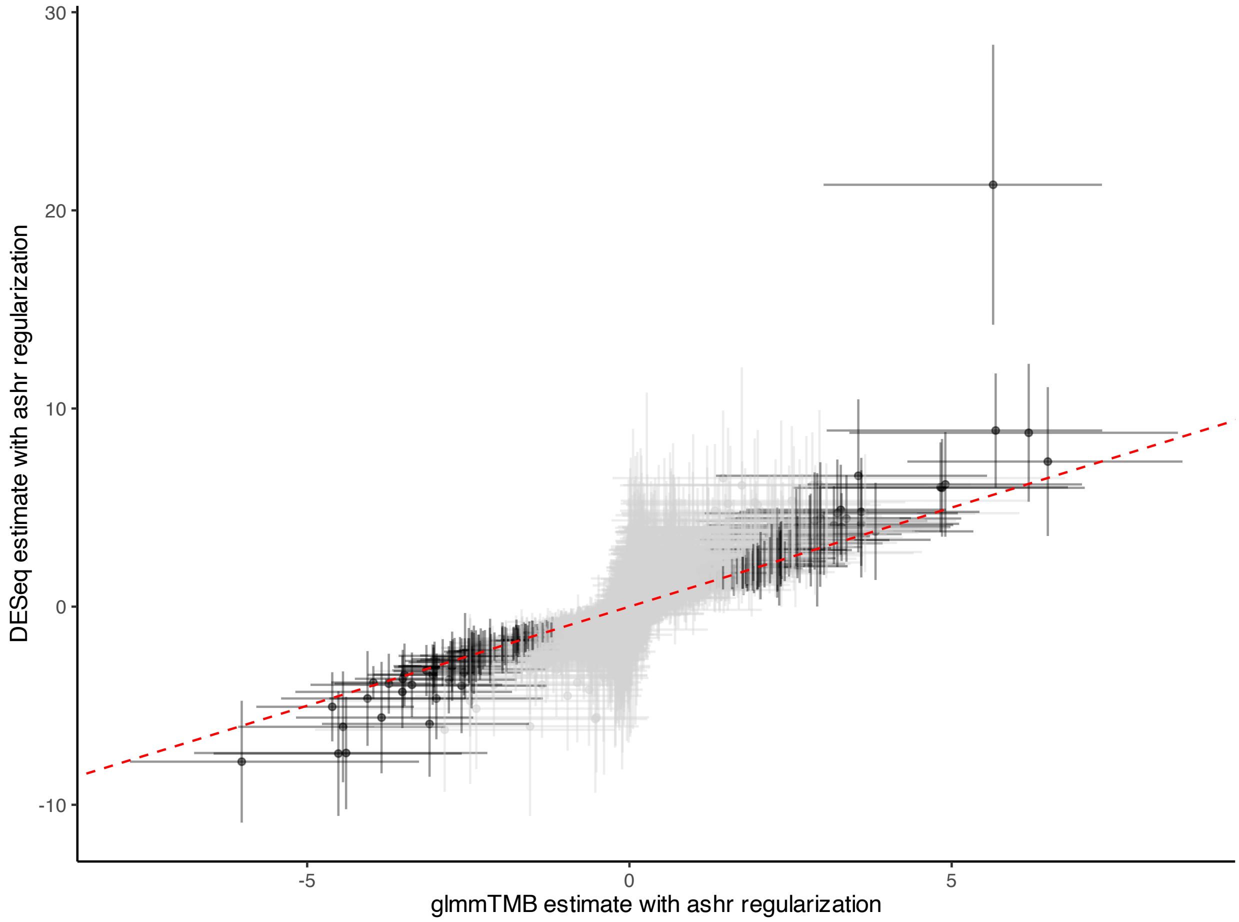

Figure S1: DeSeq2 compared glmmTMB estimate for $log_2$ fold change foreleg genes in *D. prolongata* early in development comparison. glmmTMB estimates regularized with ashr and adjusted 95% confidence intervals, DESeq estimate intervals are 2*SE. Red dotted line at a slope of 1 and intercept of zero. DESeq ashr regularization was done using the lfcshrink function. The high DESeq2 estimate gene is LOC108145159_1.

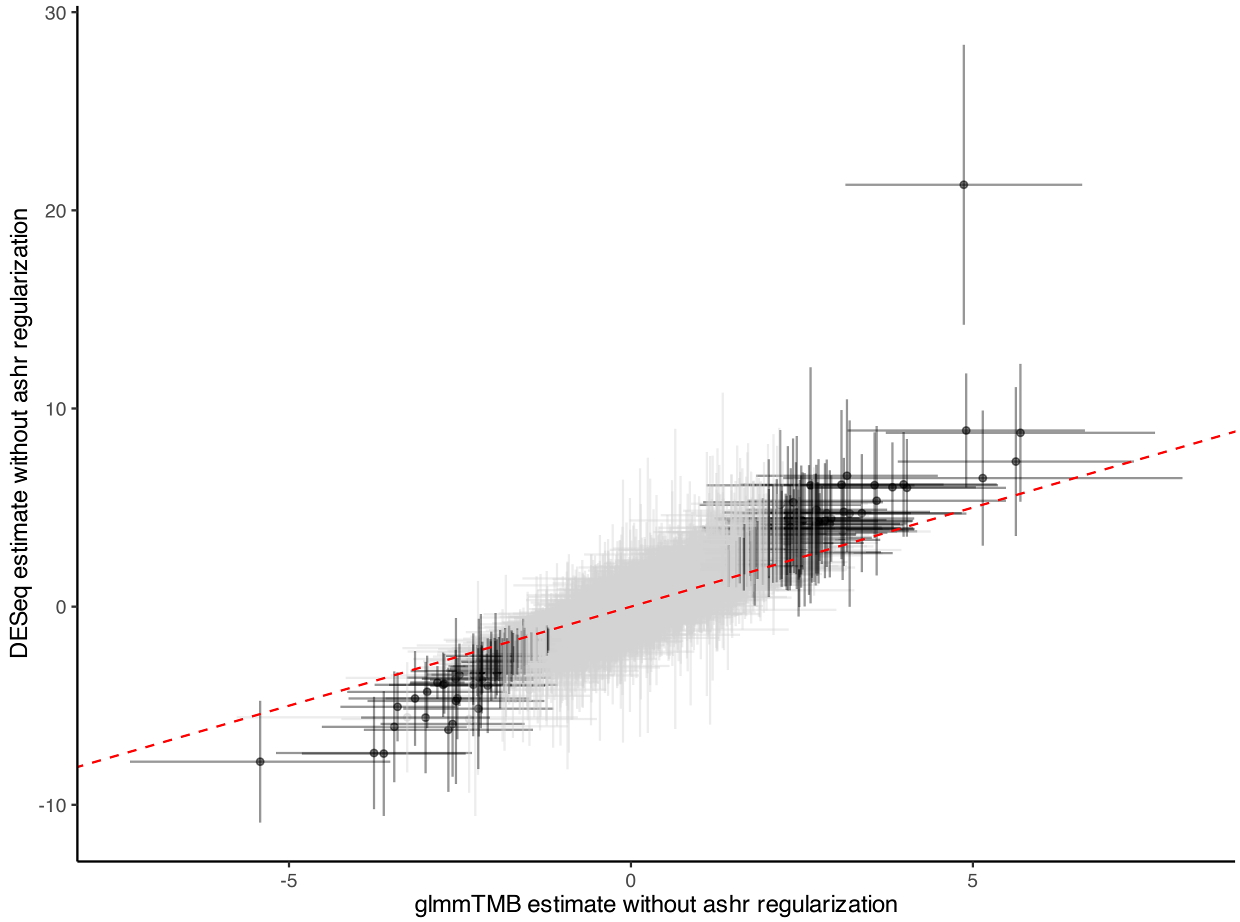

Figure S2: DeSeq2 compared glmmTMB estimate for $\log_2$ fold change foreleg genes in *D. prolongata* early in development comparison, without regularization. DESeq estimate intervals are 2*SE. Red dotted line at a slope of 1 and intercept of zero. The high DESeq2 estimate is LOC108145159_1.

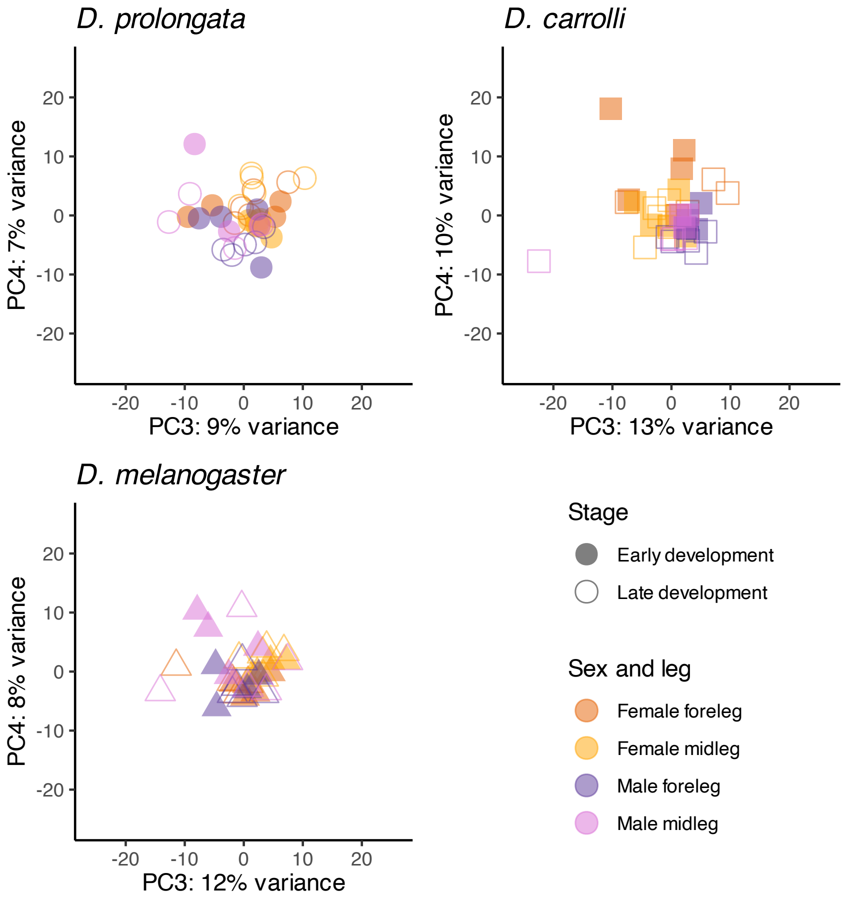

Figure S3: PC3 and PC4 within species with the top 1000 most variables genes for sex, stage, and leg.

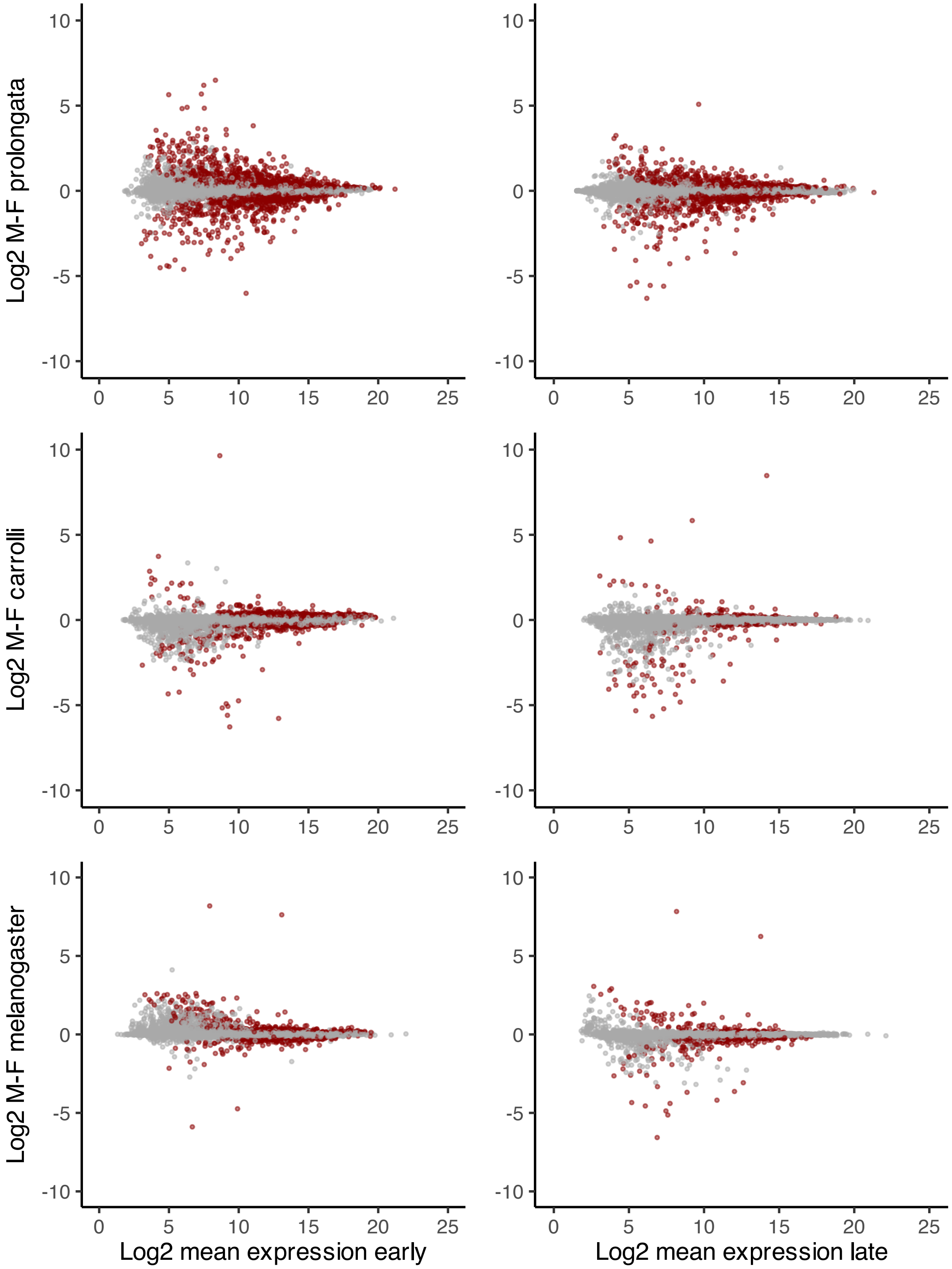

Figure S4: MA plot of sex-biased gene expression in the foreleg imaginal disc for each species. Estimated fold changes (computed with emmeans) was regularized with ashr. Red dots represent genes with ashr corrected 95% CIs that do not overlap 0.

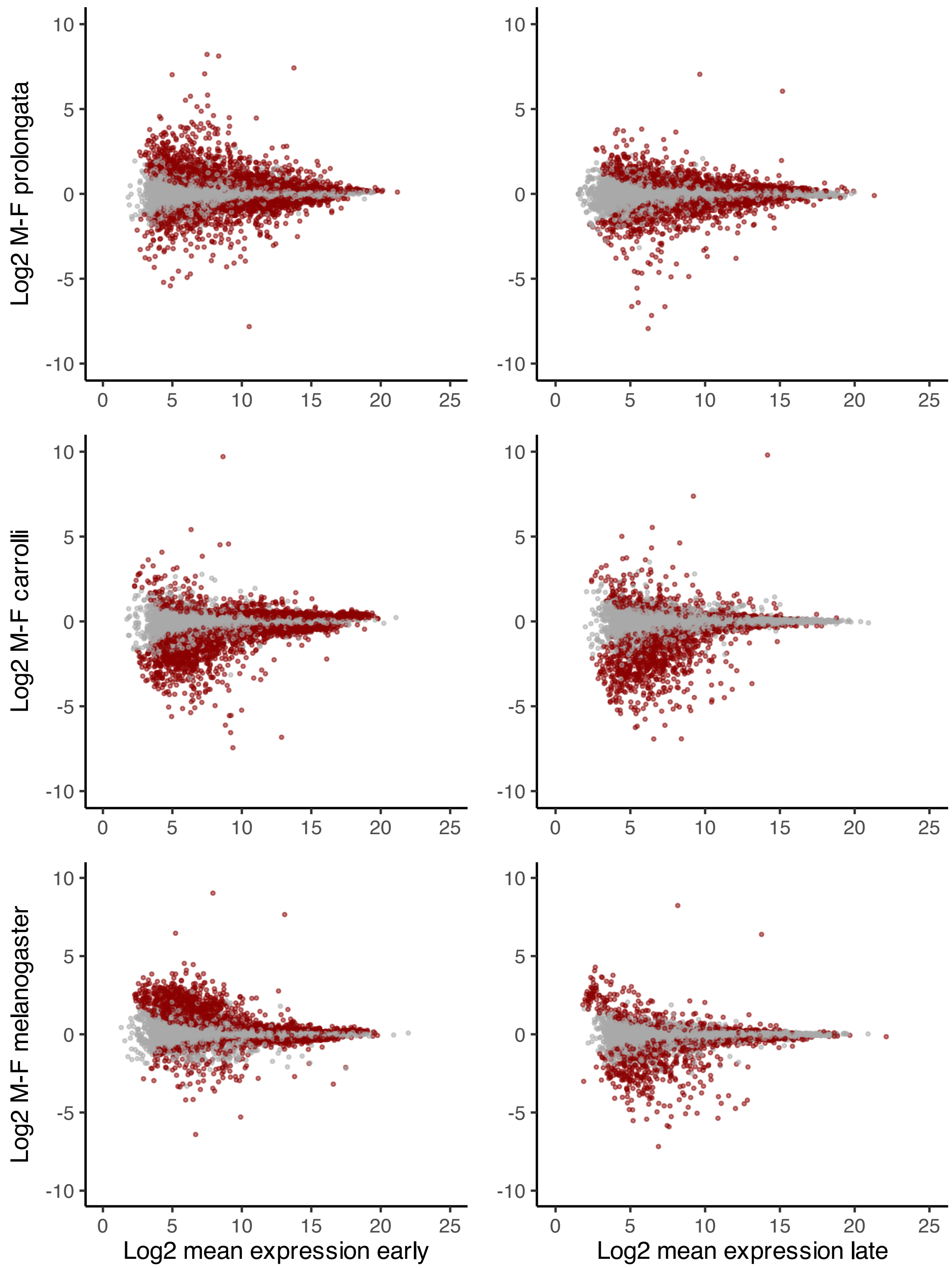

Figure S5: Figure S4: MA plot of sex-biased gene expression in the foreleg imaginal disc for each species. Estimated fold changes (computed with emmeans) was not regularized for comparison to Figure S4. Red dots represent genes whose non-regularized 95% CIs do not overlap 0, which highlights more genes than those highlighted in Figure S4.

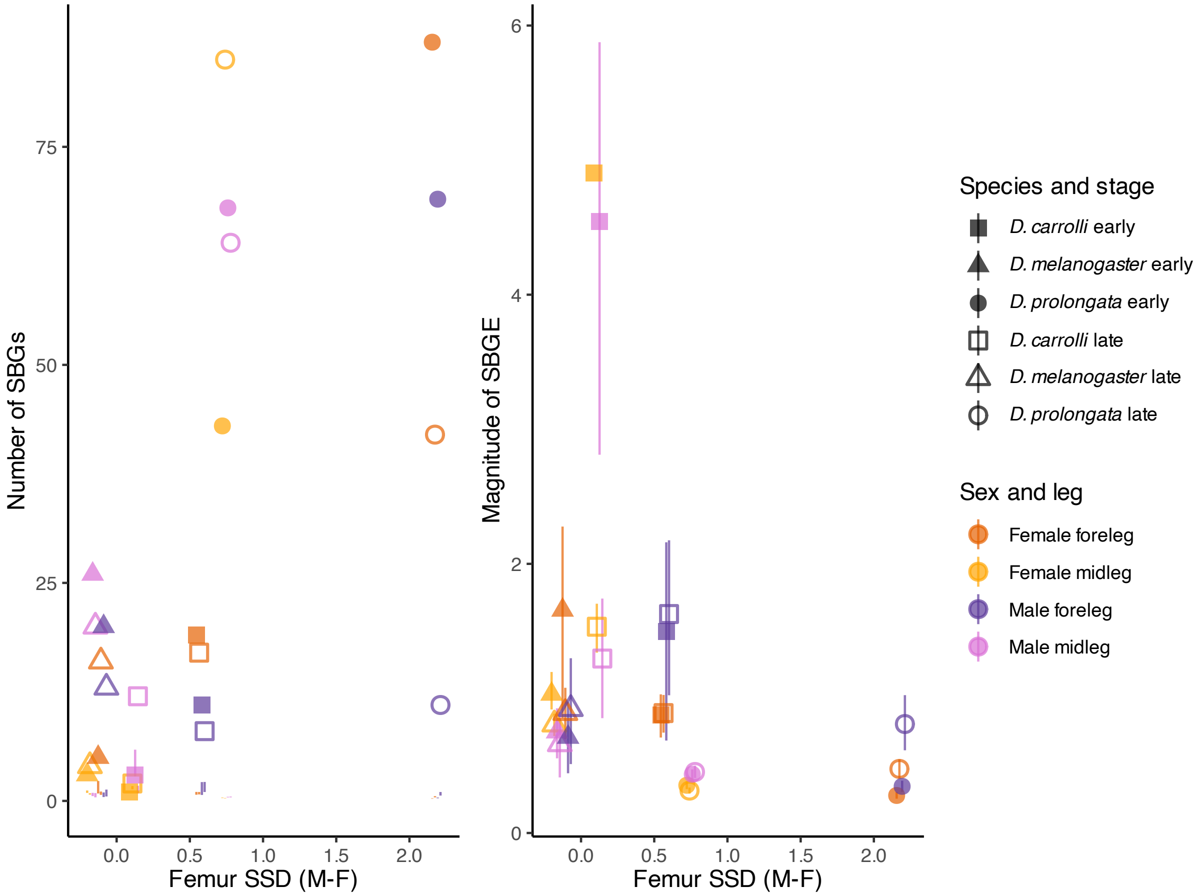

Figure S6: Relationship of the magnitude of expression (SBGE) changes (left) or number (right) for sex-biased genes (SBG) that meet a $\log_2$ fold expression change threshold (number of genes for each in Table 1) and degree of adult sexual size dimorphism, measured as female – male trait size. Magnitude and number of SBGs represent the number (or magnitude) of male (female) biased genes at each developmental stage in each species.

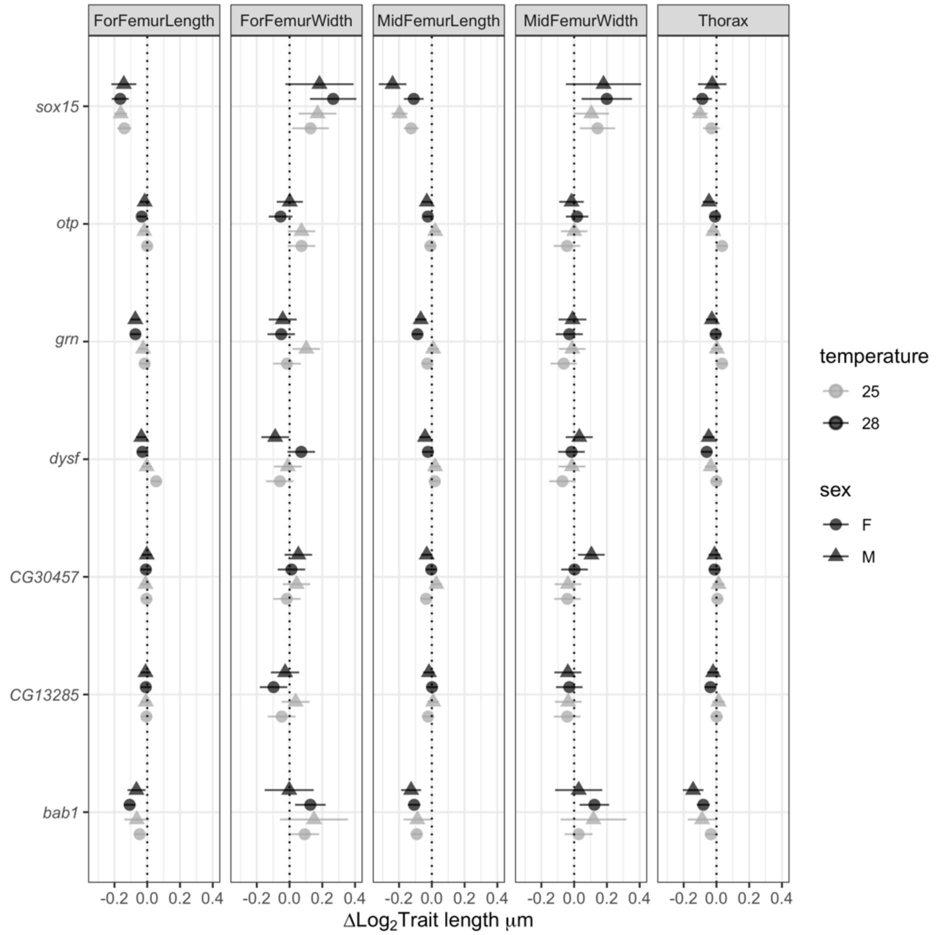

Figure S7: *pen*-Gal4 (NP6333-Gal4) knockdowns of candidate genes and their change relative to control crosses.

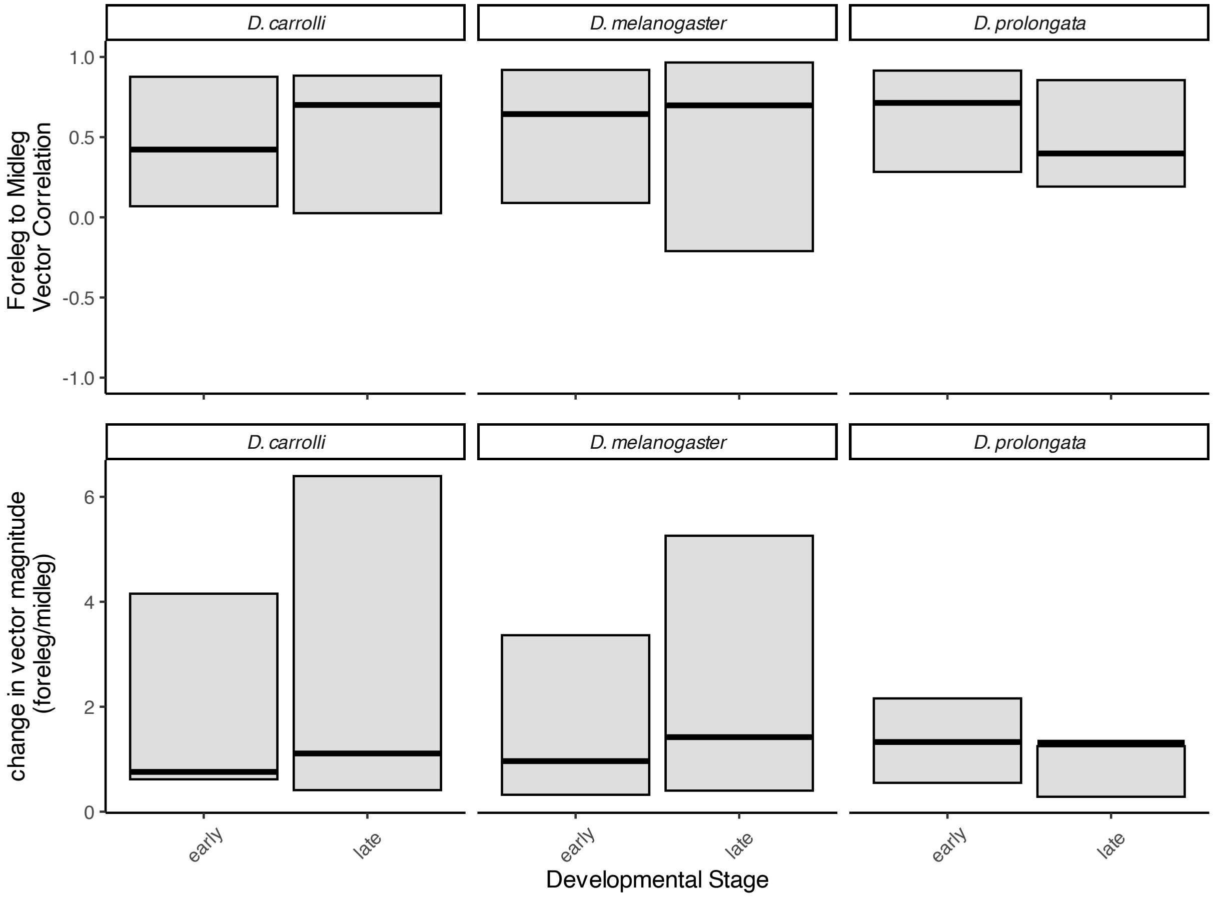

Figure S8: Vector direction and magnitude of $log_2$ fold biased genes within species between foreleg and midleg. Based on 75 female-biased $log_2$ fold change genes that overlap in all species. Top row shows direction of SBGE in foreleg compared to midleg within all three species. Bottom row is the magnitude of SBGE in foreleg relative to midleg within each species.

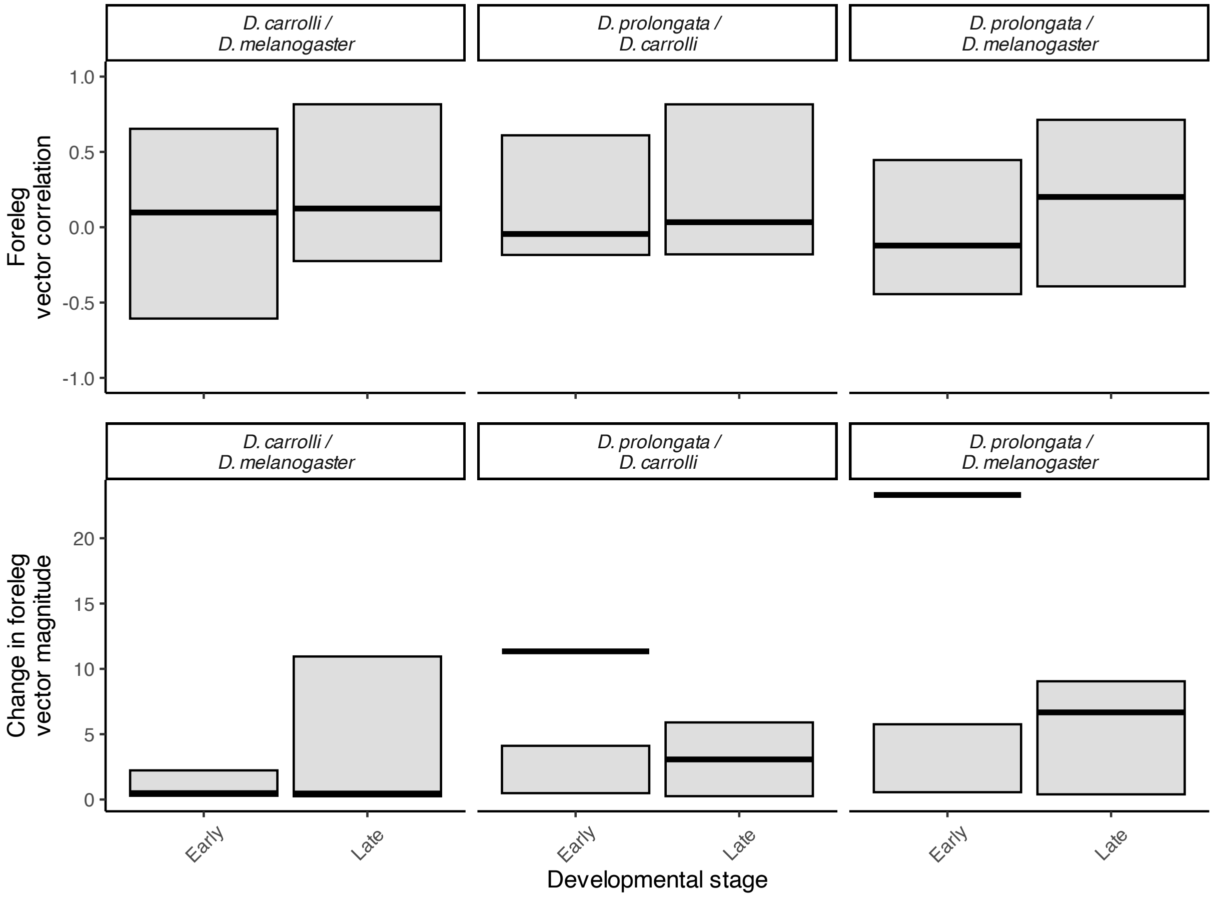

Figure S9: Direction and magnitude of male-biased $log_2$ fold biased genes between species in the foreleg. Based on 36 genes that overlap in all species. Top row shows direction of SBGE in foreleg compared between D. prolongata and the other species, the first column shows *D. carrolli* compared to *D. melanogaster* as a point of reference. Bottom row is the magnitude of SBGE in foreleg between each species.

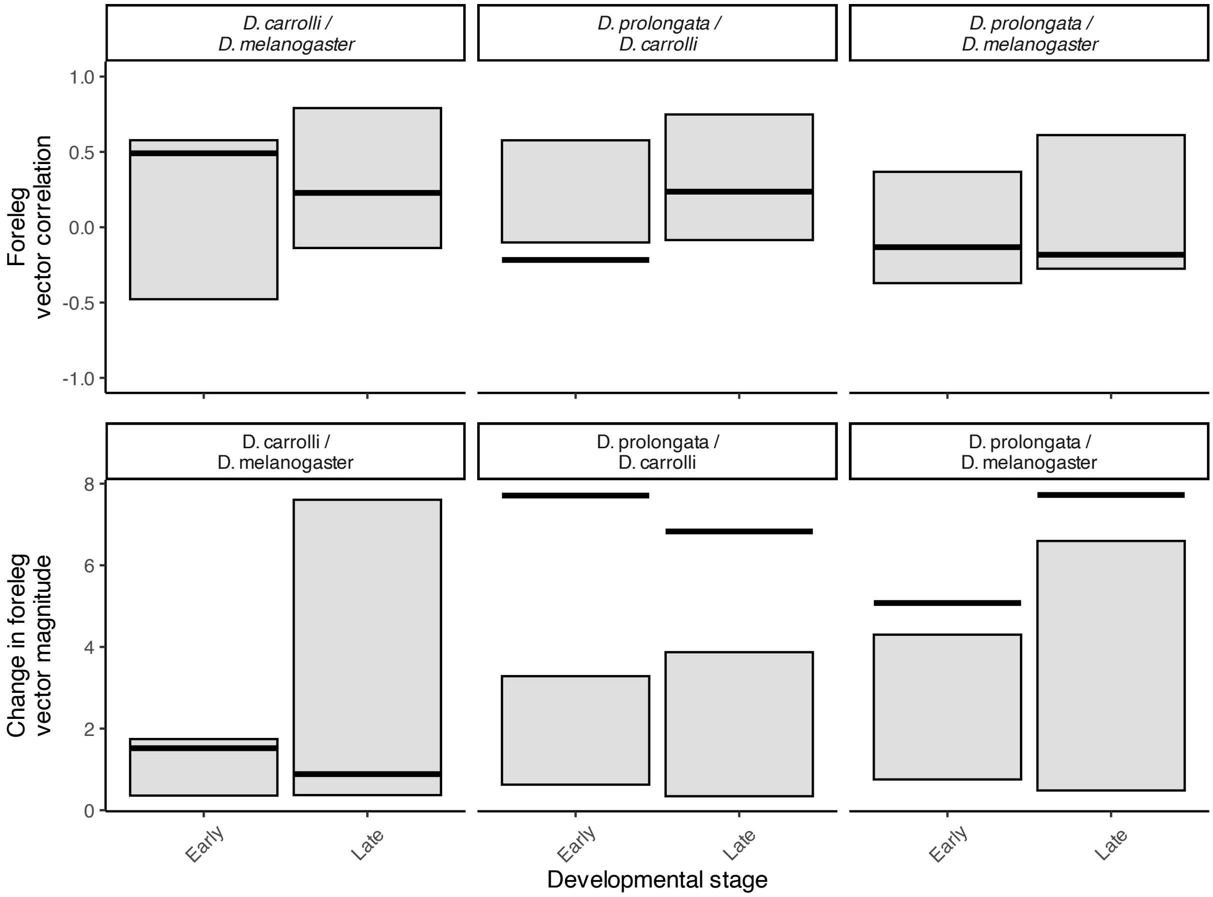

Figure S10: Direction and magnitude of female-biased $\log_2$ fold biased genes between species in the foreleg. Based on 75 genes that overlap in all species. Top row shows direction of SBGE in foreleg compared between D. prolongata and the other species, the first column shows *D. carrolli* compared to *D. melanogaster* as a point of reference. Bottom row is the magnitude of SBGE in foreleg between each species.

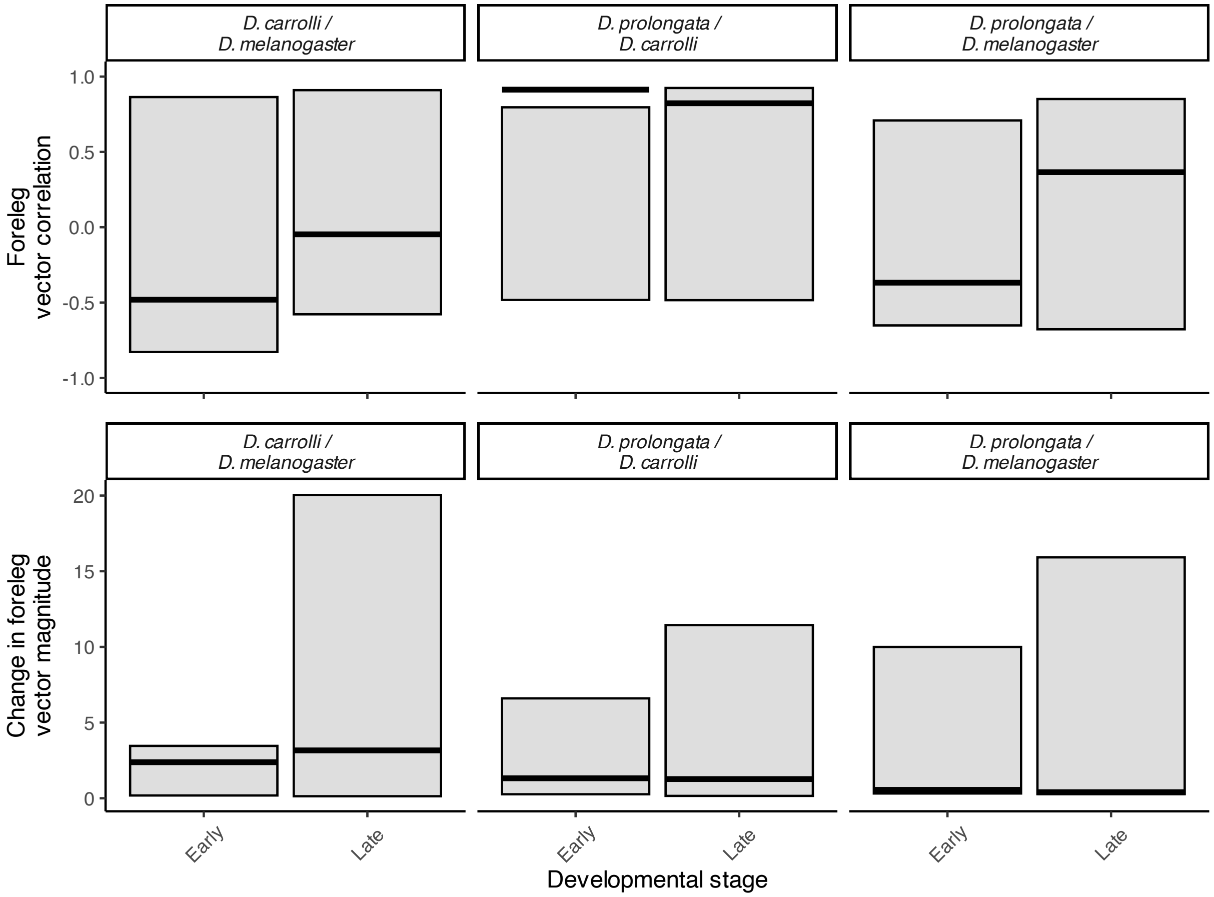

Figure S11: Direction and magnitude of *D. melanogaster* $\log_2$ fold biased genes between species in the foreleg. Based on 110 genes that overlap in all species. Top row shows direction of SBGE in foreleg compared between D. prolongata and the other species, the first column shows *D. carrolli* compared to *D. melanogaster* as a point of reference. Bottom row is the magnitude of SBGE in foreleg between each species.

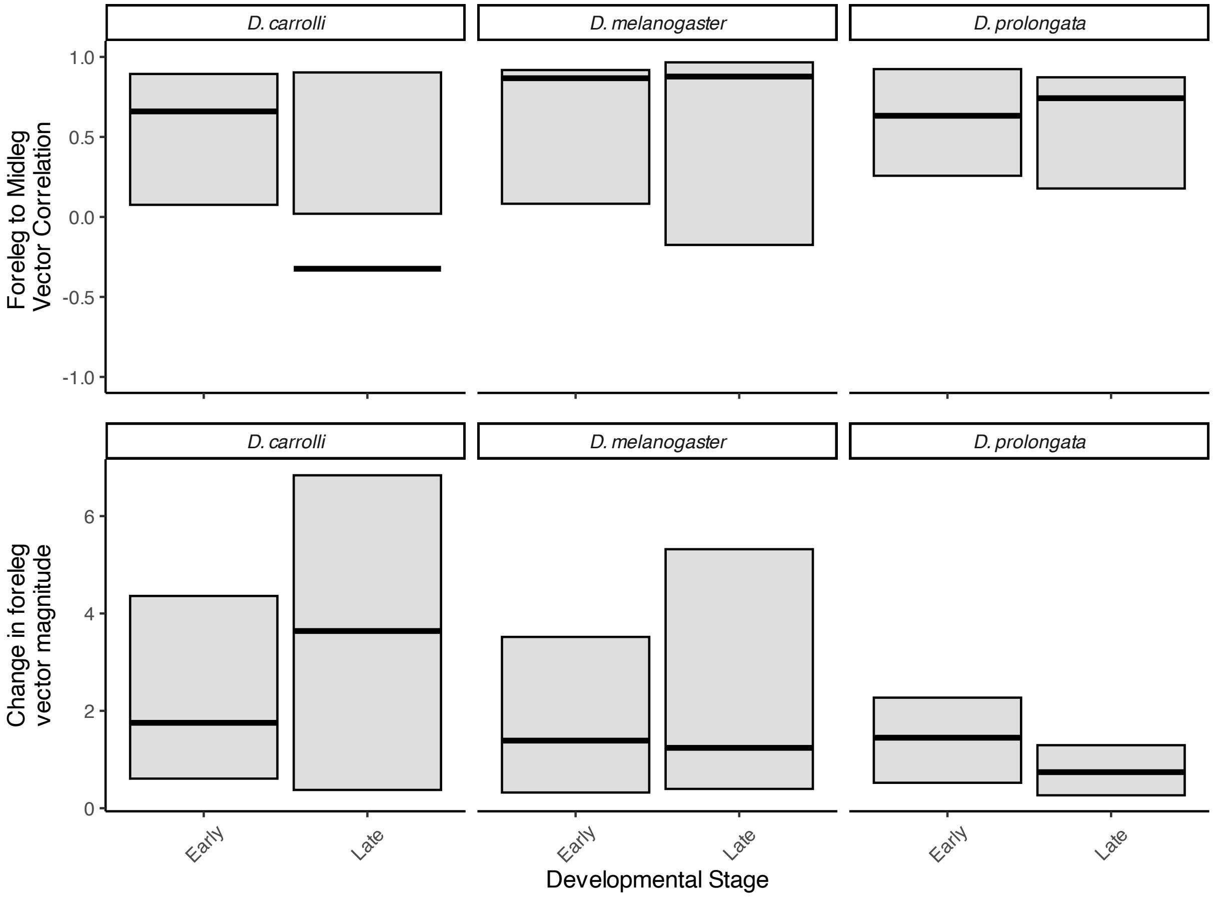

Figure S12: Vector direction and magnitude of Insulin signalling genes within species between foreleg and midleg. Based on genes that overlap in all species. Top row shows direction of SBGE in foreleg compared to midleg within all three species. Bottom row is the magnitude of SBGE in foreleg relative to midleg within each species.

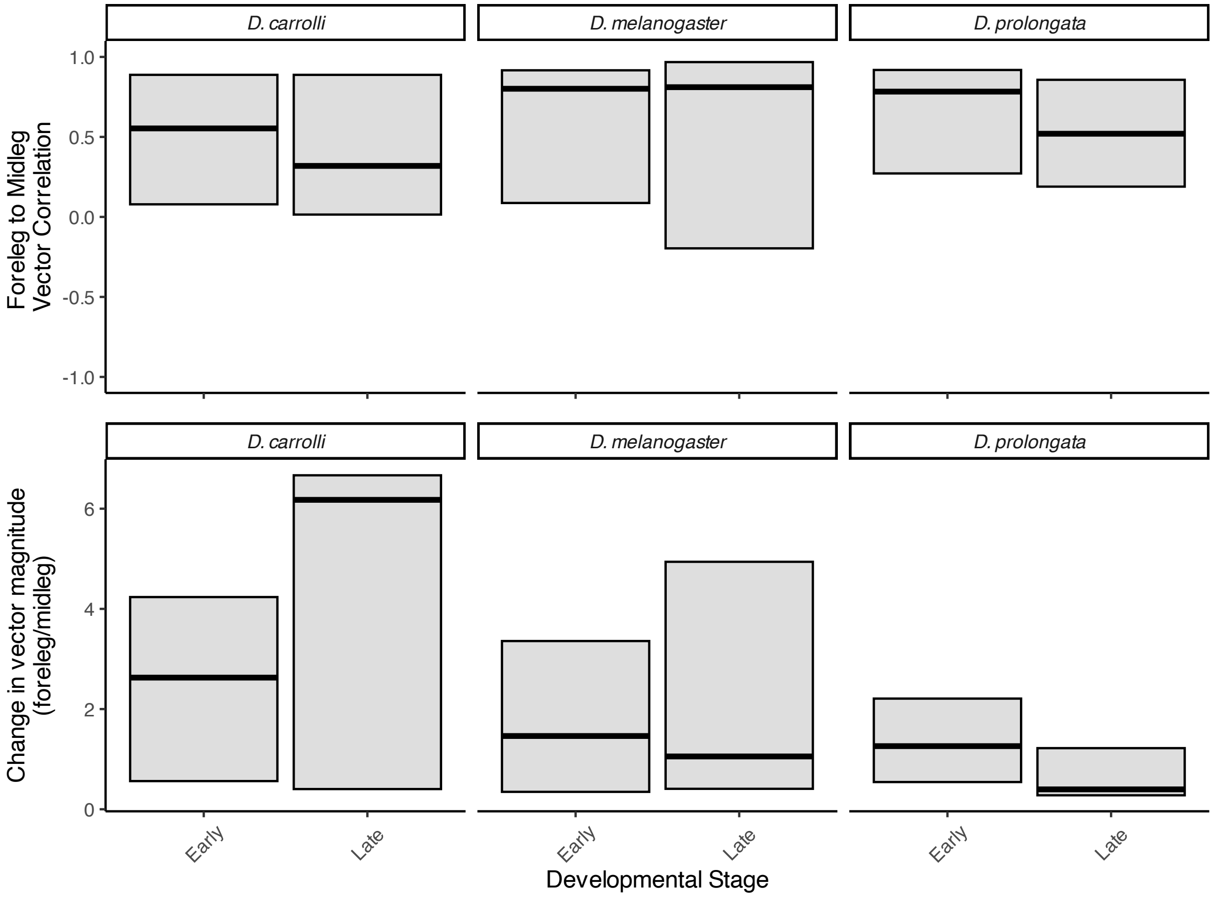

Figure S13: Vector direction and magnitude of Notch signalling genes within species between foreleg and midleg. Based on genes that overlap in all species. Top row shows direction of SBGE in foreleg compared to midleg within all three species. Bottom row is the magnitude of SBGE in foreleg relative to midleg within each species.

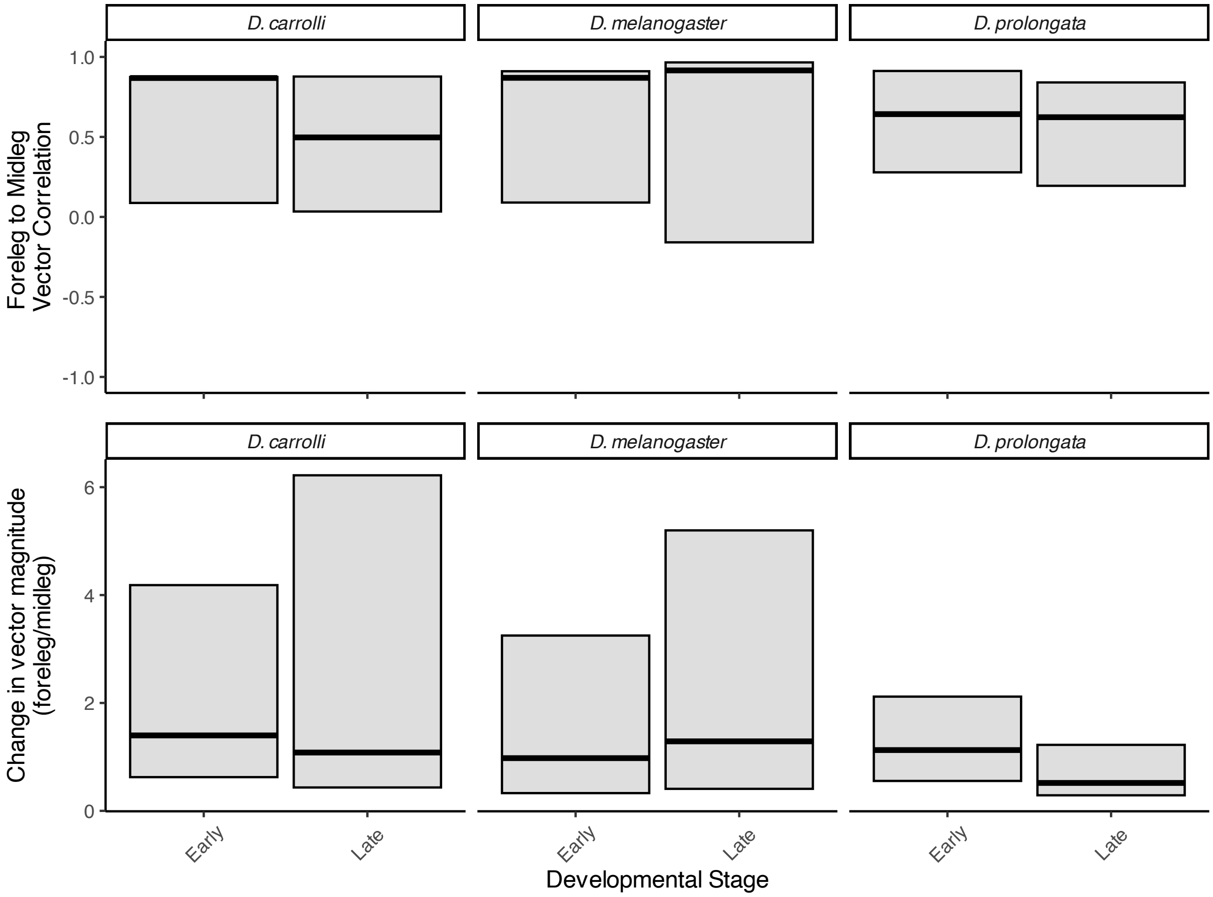

Figure S14: Vector direction and magnitude of Hippo signalling genes within species between foreleg and midleg. Based on genes that overlap in all species. Top row shows direction of SBGE in foreleg compared to midleg within all three species. Bottom row is the magnitude of SBGE in foreleg relative to midleg within each species.

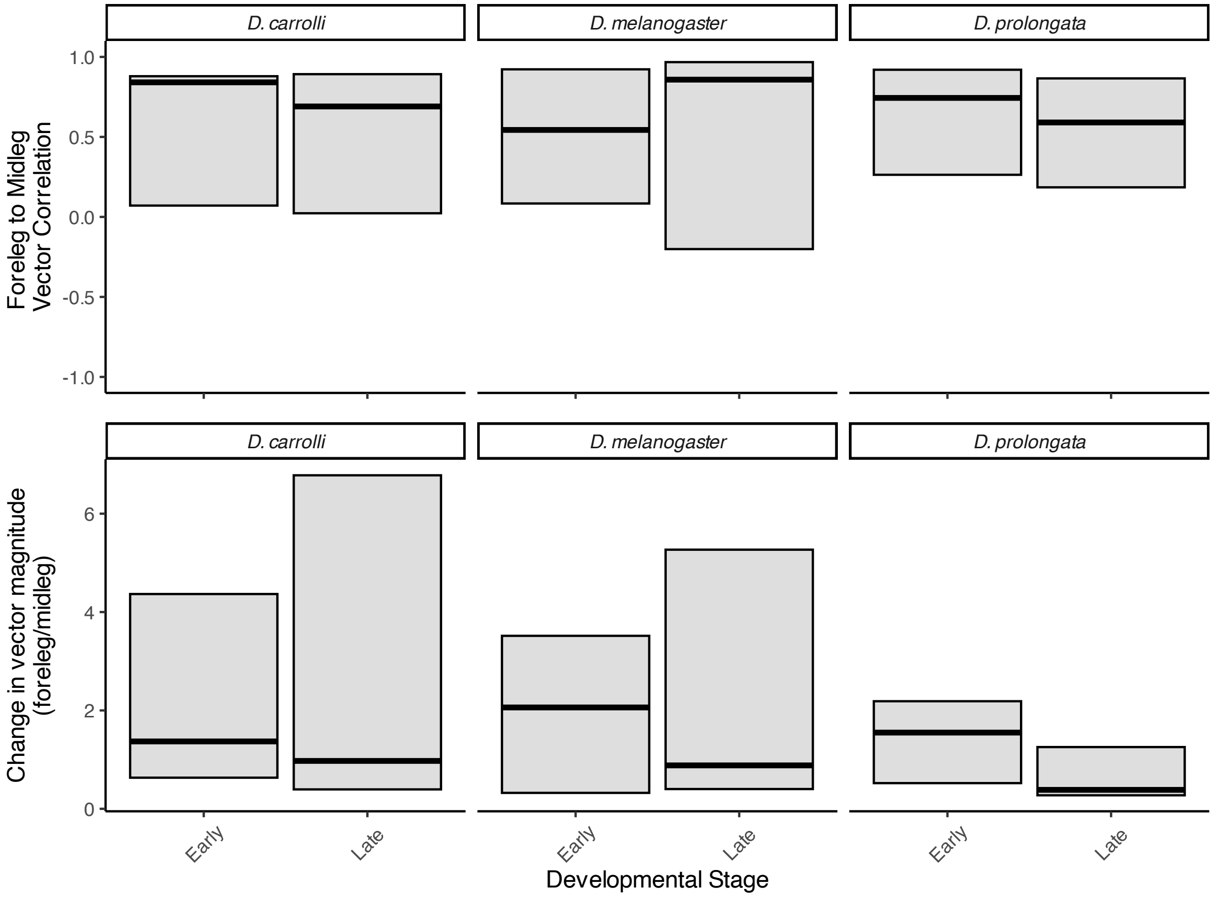

Figure S15: Vector direction and magnitude of EGFR signalling genes within species between foreleg and midleg. Based on genes that overlap in all species. Top row shows direction of SBGE in foreleg compared to midleg within all three species. Bottom row is the magnitude of SBGE in foreleg relative to midleg within each species.

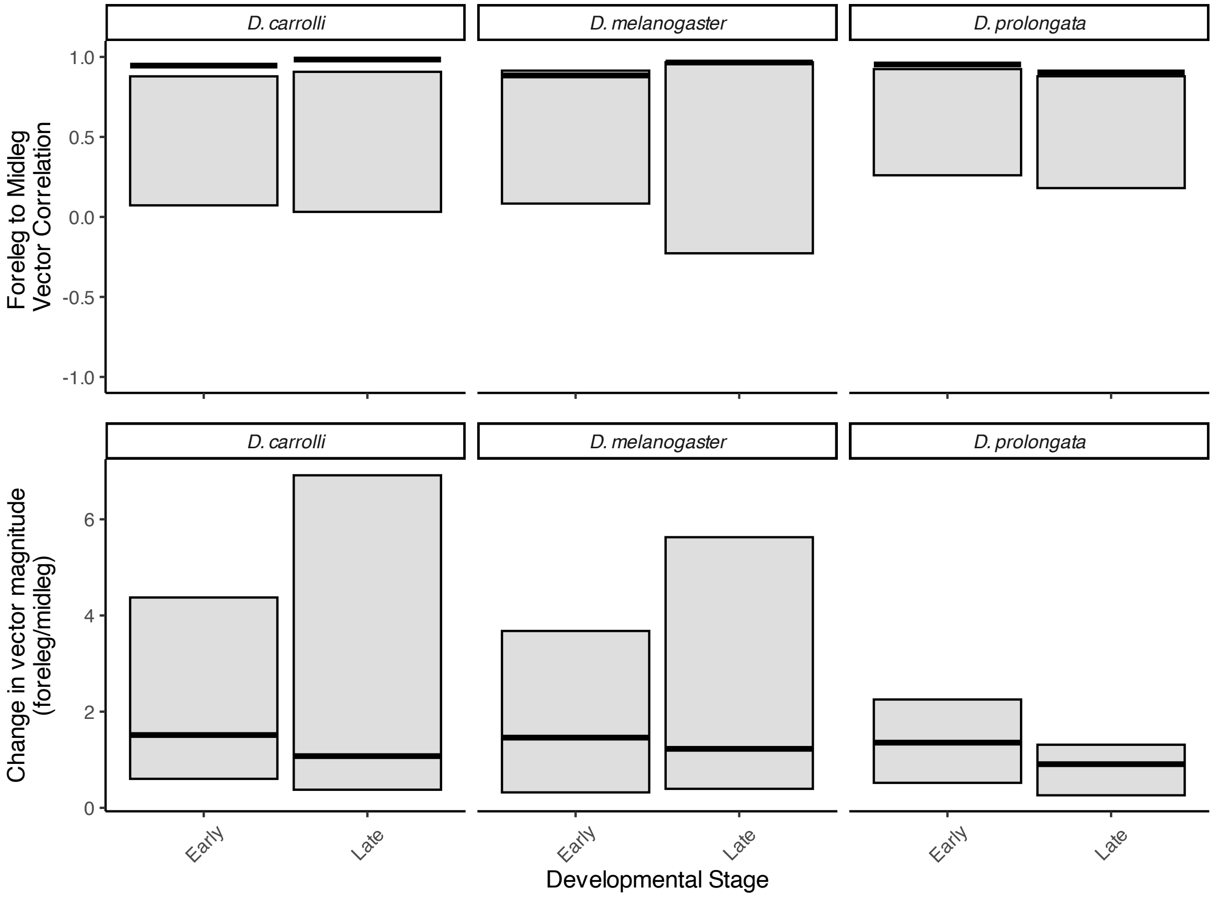

Figure S16: Vector direction and magnitude of BMP signalling genes within species between foreleg and midleg. Based on genes that overlap in all species. Top row shows direction of SBGE in foreleg compared to midleg within all three species. Bottom row is the magnitude of SBGE in foreleg relative to midleg within each species.

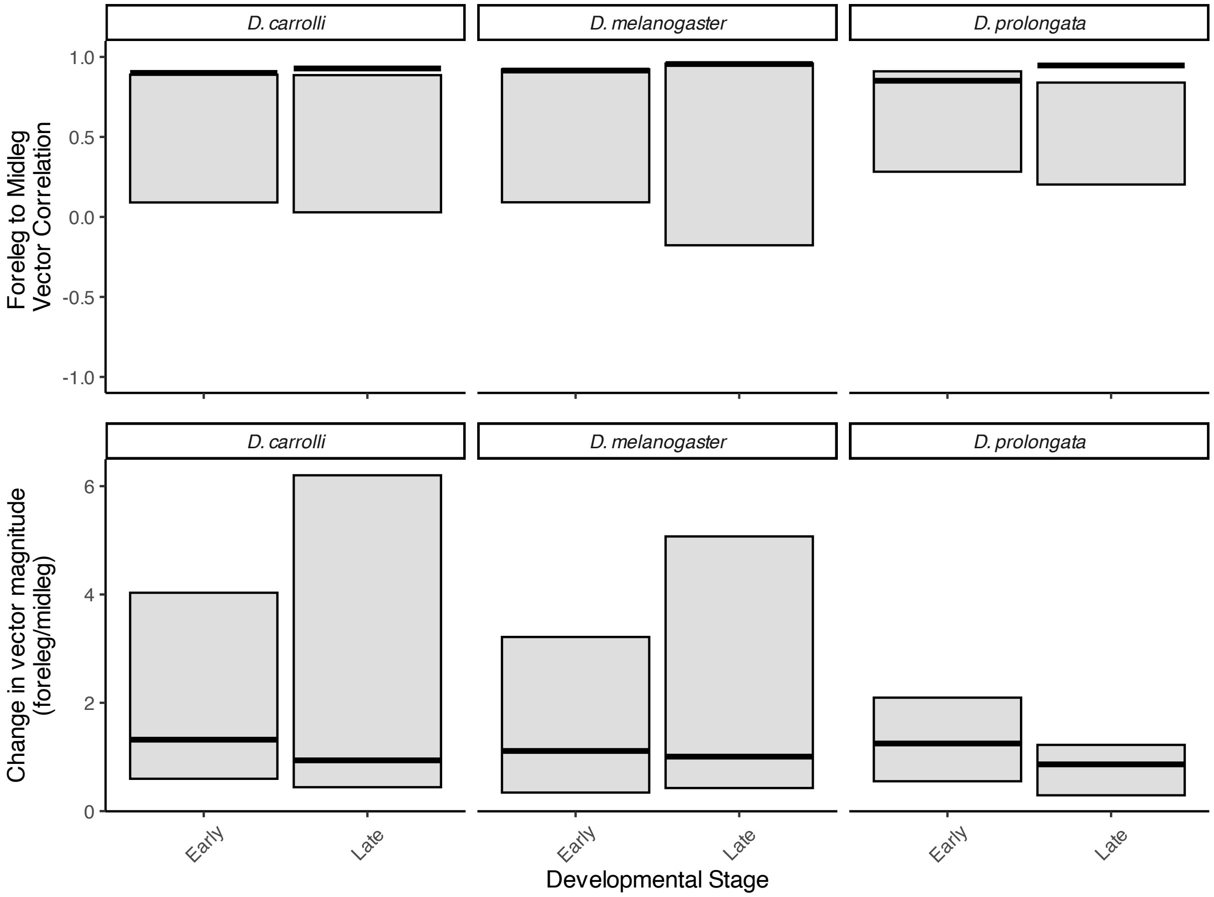

Figure S17: Vector direction and magnitude of Hedgehog signalling genes within species between foreleg and midleg. Based on genes that overlap in all species. Top row shows direction of SBGE in foreleg compared to midleg within all three species. Bottom row is the magnitude of SBGE in foreleg relative to midleg within each species.

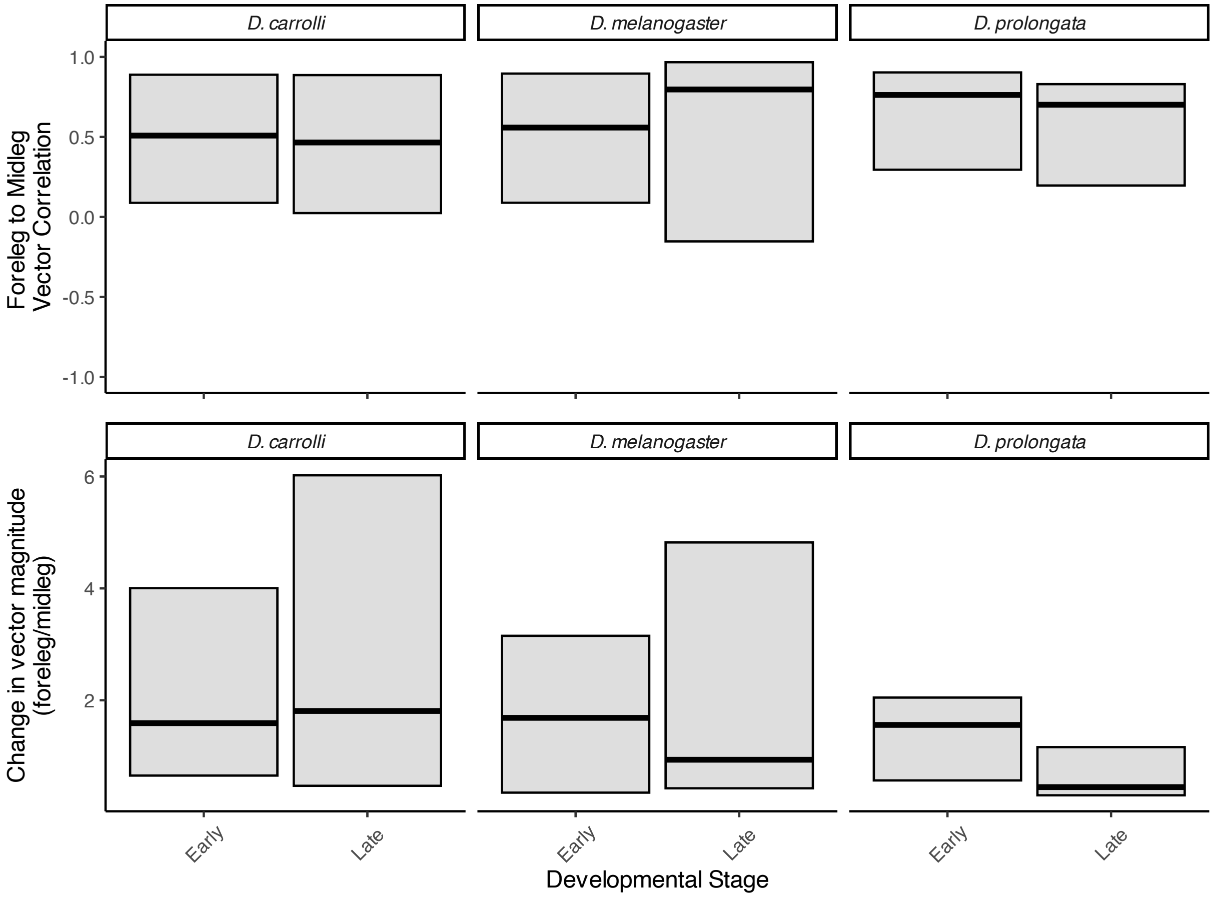

Figure S18: Vector direction and magnitude of WNT signalling genes within species between foreleg and midleg. Based on genes that overlap in all species. Top row shows direction of SBGE in foreleg compared to midleg within all three species. Bottom row is the magnitude of SBGE in foreleg relative to midleg within each species.

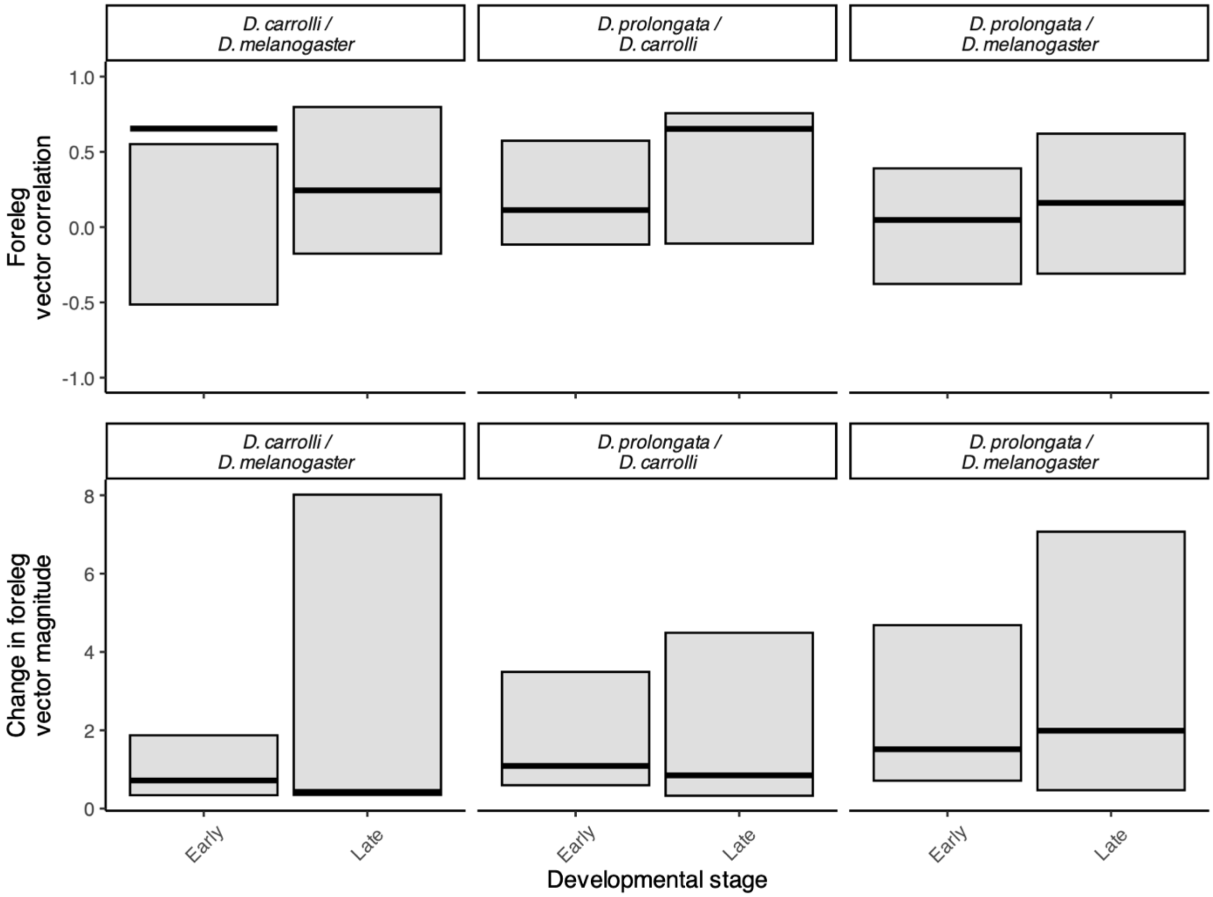

Figure S19: Degree of shared direction, and changes in magnitude for Insulin signalling genes between species in the foreleg. Top row shows direction of SBGE in foreleg compared between *D. prolongata* and the other species, and the first column showing *D. carrolli* compared to *D. melanogaster* as a point of reference. Bottom row is the magnitude of SBGE in foreleg between each species.

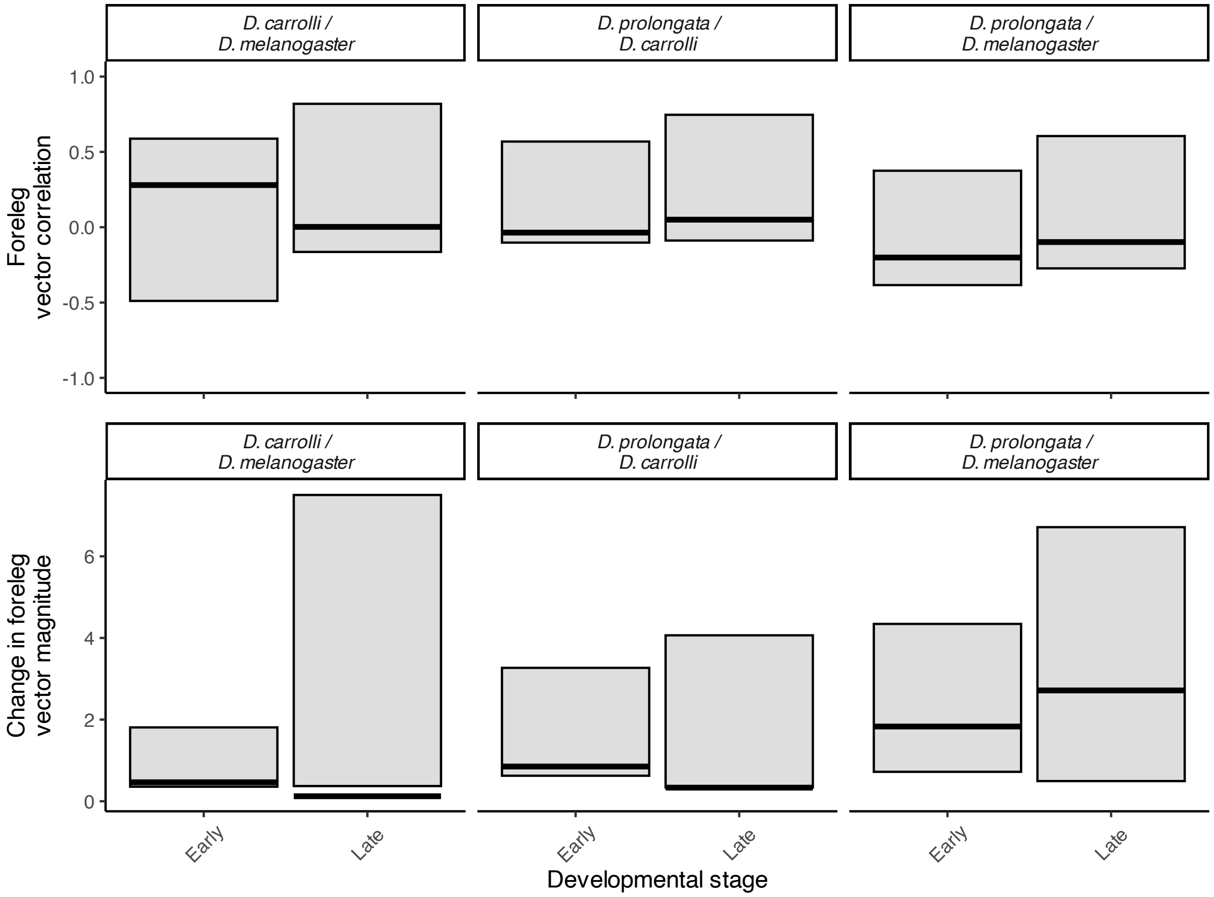

Figure S20: Direction and magnitude Notch signalling genes between species in the foreleg. Top row shows direction of SBGE in foreleg compared between *D. prolongata* and the other species, with the last column being the comparison of *D. carrolli* compared to *D. melanogaster* as a point of reference. Bottom row is the magnitude of SBGE in foreleg between each species.

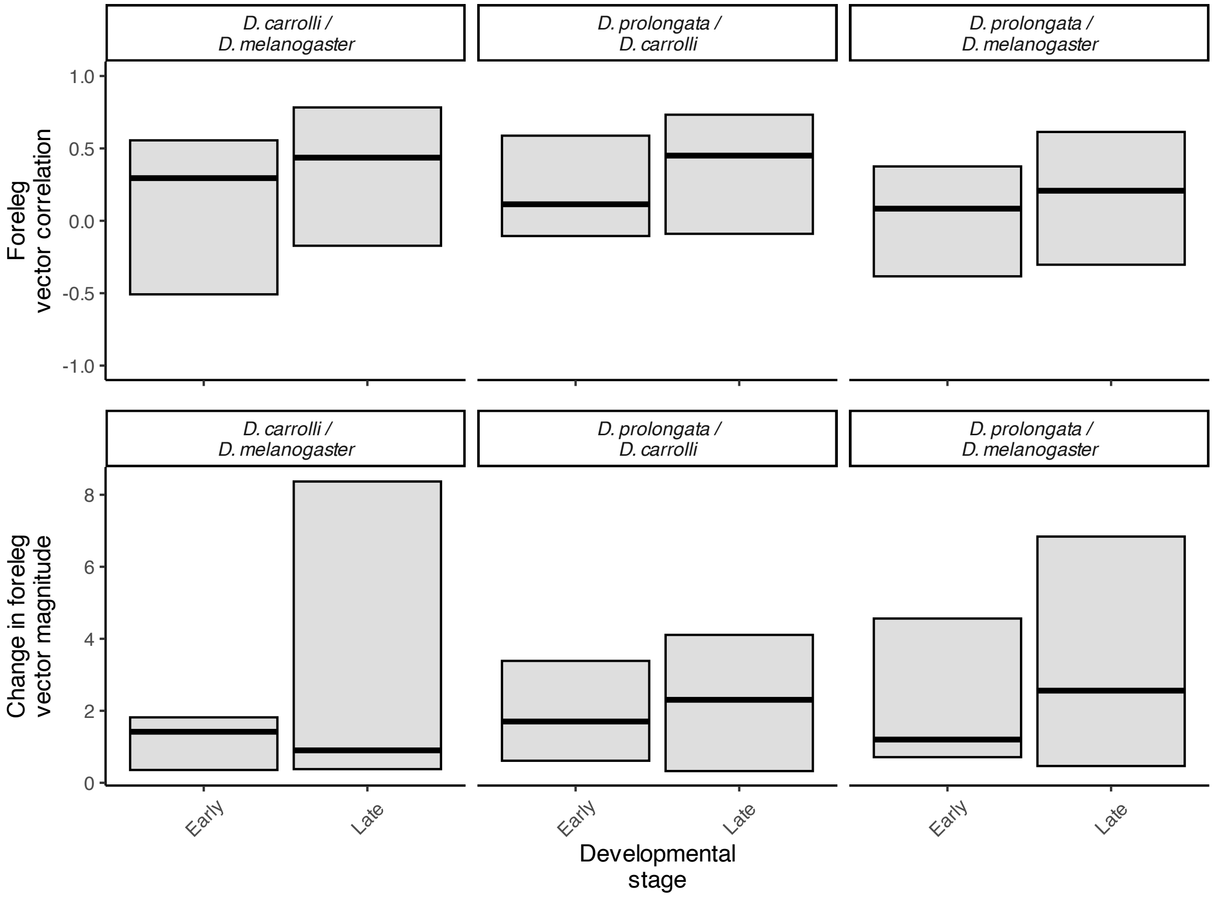

Figure S21: Direction and magnitude of EGFR signalling genes between species in the foreleg. Top row shows direction of SBGE in foreleg compared between *D. prolongata* and the other species, with the last column being the comparison of *D. carrolli* compared to *D. melanogaster* as a point of reference. Bottom row is the magnitude of SBGE in foreleg between each species.

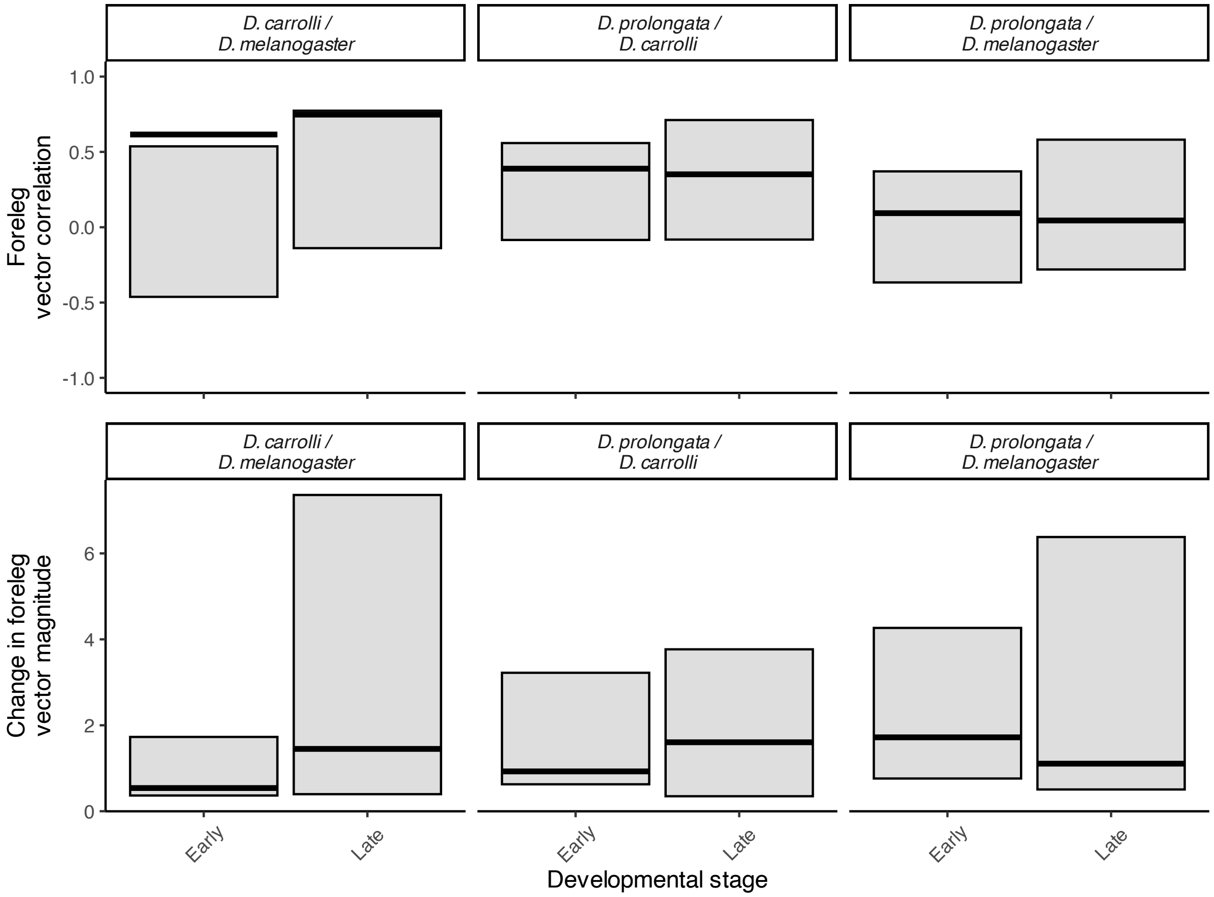

Figure S22: Direction and magnitude of Hippo signalling genes between species in the foreleg. Top row shows direction of SBGE in foreleg compared between *D. prolongata* and the other species, with the last column being the comparison of *D. carrolli* compared to *D. melanogaster* as a point of reference. Bottom row is the magnitude of SBGE in foreleg between each species.

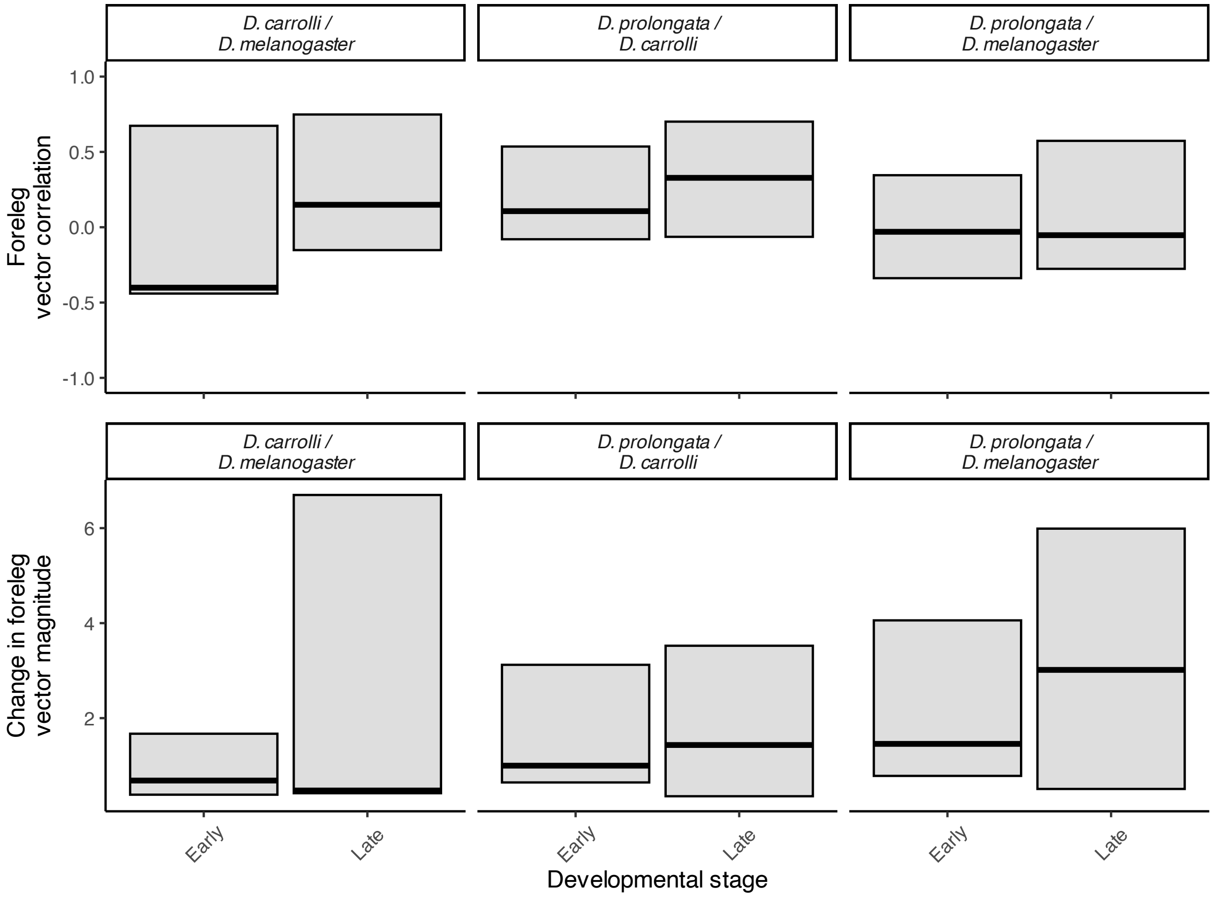

Figure S23: Direction and magnitude of WNT signalling genes between species in the foreleg. Top row shows direction of SBGE in foreleg compared between *D. prolongata* and the other species, with the last column being the comparison of *D. carrolli* compared to *D. melanogaster* as a point of reference. Bottom row is the magnitude of SBGE in foreleg between each species.

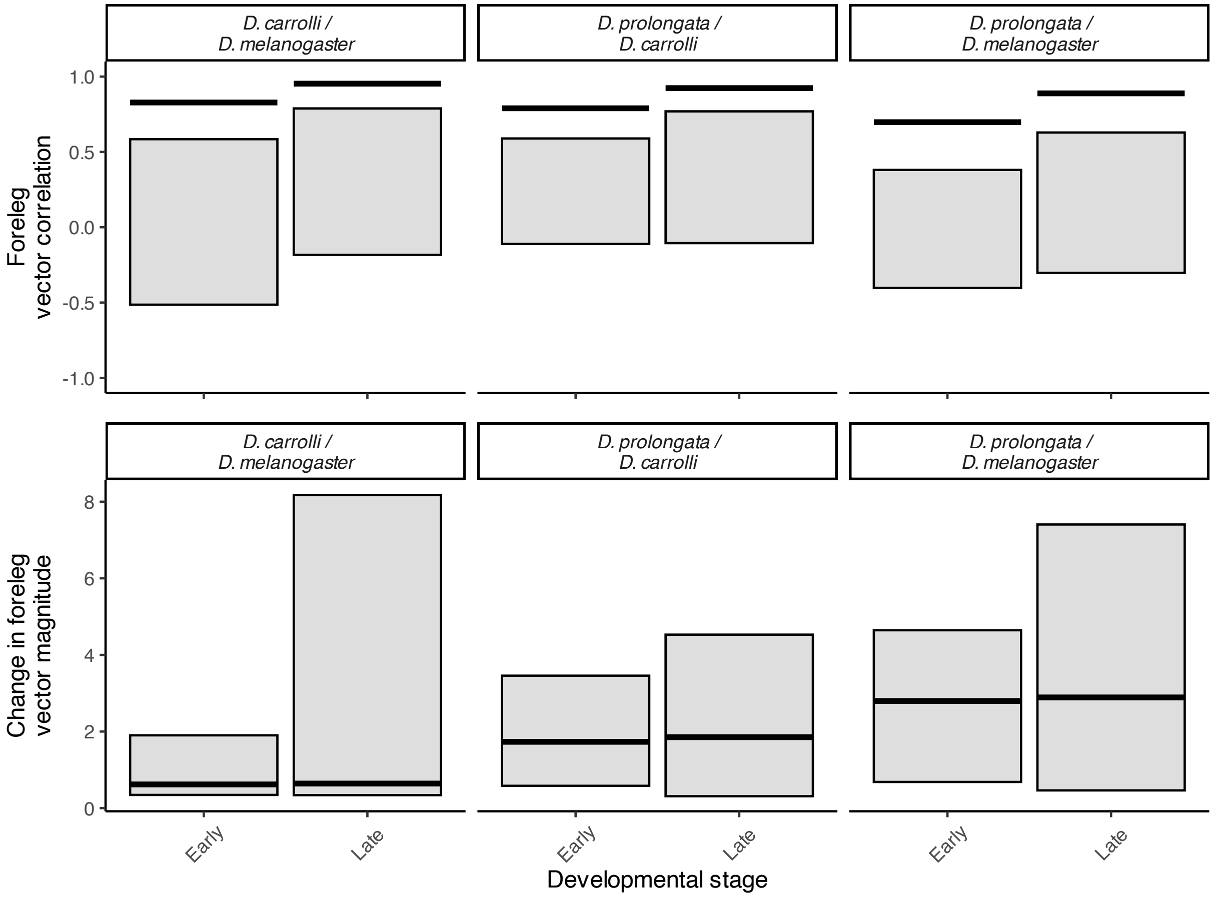

Figure S24: Direction and magnitude of BMP signalling genes between species in the foreleg. Top row shows direction of SBGE in foreleg compared between *D. prolongata* and the other species, with the last column being the comparison of *D. carrolli* compared to *D. melanogaster* as a point of reference. Bottom row is the magnitude of SBGE in foreleg between each species.

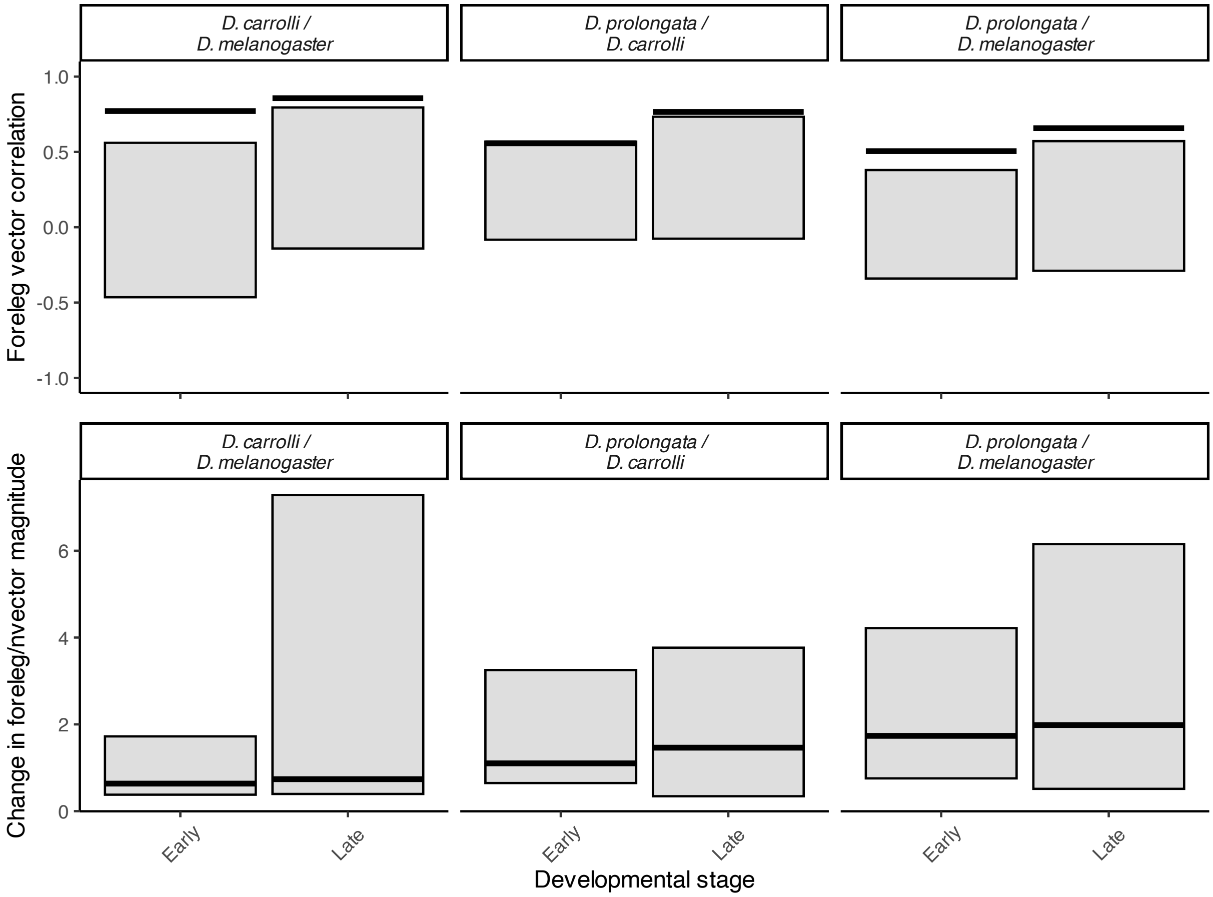

Figure S25: Direction and magnitude of Hedgehog signalling genes between species in the foreleg. Top row shows direction of SBGE in foreleg compared between *D. prolongata* and the other species, with the last column being the comparison of *D. carrolli* compared to *D. melanogaster* as a point of reference. Bottom row is the magnitude of SBGE in foreleg between each species.

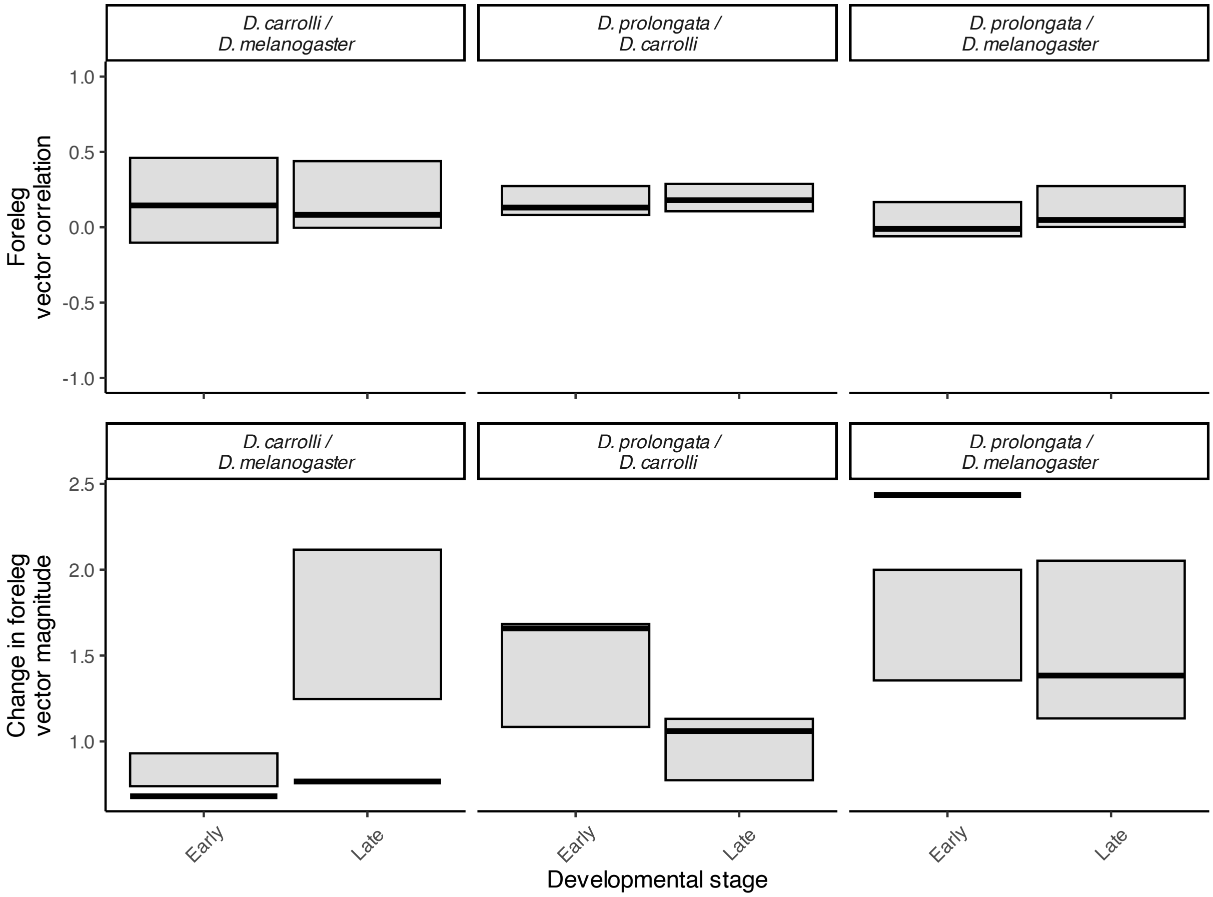

Figure S26: Direction and magnitude of putative Grn targets between species in the foreleg. Top row shows direction of SBGE in foreleg compared between *D. prolongata* and the other species, the first column shows *D. carrolli* compared to *D. melanogaster* as a point of reference. Bottom row is the magnitude of SBGE in foreleg between each species.

Table S1: Summary of software used, versions, and any non-default flags or parameters

| software | version | flags |
|---|---|---|
| BBduk | 39.06 | threads=32 ftr=100 ktrim=r k=21 mink=10 hdist=1 tpe tbo qtrim=rl trimq=15 minlength=36 rcomp=t |
| agat | 0.9.2 | convert_sp_gff2gtf.pl |
| STAR | 2.7.11b | --quantMode TranscriptomeSAM GeneCounts --outSAMtype BAM SortedByCoordinate --sjdbFileChrStartEnd |
| glmmTMB | 1.1.10 | the_counts ~ (sex+leg+stage)^3 + diag(1 \| stage:replicate) offset = normFactors family = nbinom2() |
| emmeans | 1.10.5 | |
| DESeq2 | 1.46.0 | lfcShrink(res, type = "ashr") |
| blast+ | 2.14.1 | -outfmt "6 stitle sstart send sseqid" -sorthits 4 -max_target_seqs 1 |
| bedtools getfasta | 2.31 | |
| MUSCLE | 3.8.1551 | |
| Tandem Repeat Finder | 4.09.1 | 2 7 7 80 10 50 500 -m -h |
| SCRMshaw | 1.1 | --imm |
| tidyverse | 2.0.0 | |
| ashr | 2.2-63 | |
| data.table | 1.16.2 | |

Table S2: model estimates for all traits contrasted between UAS-RNAi genes and *Pen*$^{NP3666}$-*Gal4* in the multivariate model with thorax modelled.

| trait | temp | sex | code_trt.vs.ctrl1 | estimate | SE | df | t.ratio | p.value |
|---|---|---|---|---|---|---|---|---|
| ForFemurLength | 25 | F | bab1 - control | -0.05 | 0.01 | 5803 | -3.79 | 3.60e-04 |
| ForFemurLength | 25 | F | CG13285 - control | -5.05e-03 | 0.01 | 5803 | -0.43 | 0.78 |
| ForFemurLength | 25 | F | CG30457 - control | -4.88e-03 | 0.01 | 5803 | -0.42 | 0.78 |
| ForFemurLength | 25 | F | dysf - control | 0.05 | 0.01 | 5803 | 4.73 | 8.07e-06 |
| ForFemurLength | 25 | F | grn - control | -0.02 | 0.01 | 5803 | -1.36 | 0.30 |
| ForFemurLength | 25 | F | otp - control | -4.88e-04 | 0.01 | 5803 | -0.04 | 0.97 |
| ForFemurLength | 25 | F | Sox15 - control | -0.14 | 0.02 | 5803 | -8.84 | 8.75e-18 |
| ForFemurLength | 28 | F | bab1 - control | -0.11 | 0.01 | 5803 | -8.21 | 9.82e-16 |
| ForFemurLength | 28 | F | CG13285 - control | -8.57e-03 | 0.01 | 5803 | -0.74 | 0.49 |
| ForFemurLength | 28 | F | CG30457 - control | -7.97e-03 | 0.01 | 5803 | -0.69 | 0.49 |
| ForFemurLength | 28 | F | dysf - control | -0.03 | 0.01 | 5803 | -2.53 | 0.02 |
| ForFemurLength | 28 | F | grn - control | -0.07 | 0.01 | 5803 | -6.21 | 1.32e-09 |
| ForFemurLength | 28 | F | otp - control | -0.03 | 0.01 | 5803 | -3.25 | 2.02e-03 |
| ForFemurLength | 28 | F | Sox15 - control | -0.16 | 0.02 | 5803 | -8.53 | 1.31e-16 |
| ForFemurLength | 25 | M | bab1 - control | -0.06 | 0.03 | 5803 | -2.25 | 0.07 |
| ForFemurLength | 25 | M | CG13285 - control | -8.19e-03 | 0.01 | 5803 | -0.69 | 0.57 |

| | | | | | | | | |
|---|---|---|---|---|---|---|---|---|
| ForFemurLength | 25 | M | CG30457 - control | -0.01 | 0.01 | 5803 | -0.94 | 0.49 |
| ForFemurLength | 25 | M | dysf - control | -3.02e-03 | 0.01 | 5803 | -0.26 | 0.79 |
| ForFemurLength | 25 | M | grn - control | -0.03 | 0.01 | 5803 | -2.17 | 0.07 |
| ForFemurLength | 25 | M | otp - control | -0.02 | 0.01 | 5803 | -1.57 | 0.21 |
| ForFemurLength | 25 | M | Sox15 - control | -0.16 | 0.01 | 5803 | -11.17 | 7.45e-28 |
| ForFemurLength | 28 | M | bab1 - control | -0.07 | 0.02 | 5803 | -3.22 | 3.01e-03 |
| ForFemurLength | 28 | M | CG13285 - control | -0.01 | 0.01 | 5803 | -0.89 | 0.44 |
| ForFemurLength | 28 | M | CG30457 - control | -2.71e-03 | 0.01 | 5803 | -0.23 | 0.82 |
| ForFemurLength | 28 | M | dysf - control | -0.04 | 0.01 | 5803 | -3.11 | 3.33e-03 |
| ForFemurLength | 28 | M | grn - control | -0.07 | 0.01 | 5803 | -6.20 | 4.31e-09 |
| ForFemurLength | 28 | M | otp - control | -0.02 | 0.01 | 5803 | -1.42 | 0.22 |
| ForFemurLength | 28 | M | Sox15 - control | -0.14 | 0.03 | 5803 | -5.05 | 1.56e-06 |
| ForFemurWidth | 25 | F | bab1 - control | 0.09 | 0.03 | 5803 | 2.80 | 0.02 |
| ForFemurWidth | 25 | F | CG13285 - control | -0.05 | 0.03 | 5803 | -1.52 | 0.18 |
| ForFemurWidth | 25 | F | CG30457 - control | -0.02 | 0.03 | 5803 | -0.53 | 0.62 |
| ForFemurWidth | 25 | F | dysf - control | -0.06 | 0.03 | 5803 | -1.90 | 0.10 |
| ForFemurWidth | 25 | F | grn - control | -0.02 | 0.03 | 5803 | -0.50 | 0.62 |
| ForFemurWidth | 25 | F | otp - control | 0.07 | 0.03 | 5803 | 2.36 | 0.04 |
| ForFemurWidth | 25 | F | Sox15 - control | 0.13 | 0.04 | 5803 | 3.07 | 0.01 |
| ForFemurWidth | 28 | F | bab1 - control | 0.13 | 0.03 | 5803 | 3.65 | 9.10e-04 |
| ForFemurWidth | 28 | F | CG13285 - control | -0.10 | 0.03 | 5803 | -3.13 | 4.08e-03 |
| ForFemurWidth | 28 | F | CG30457 - control | 0.01 | 0.03 | 5803 | 0.37 | 0.71 |
| ForFemurWidth | 28 | F | dysf - control | 0.07 | 0.03 | 5803 | 2.29 | 0.04 |
| ForFemurWidth | 28 | F | grn - control | -0.05 | 0.03 | 5803 | -1.63 | 0.12 |
| ForFemurWidth | 28 | F | otp - control | -0.05 | 0.03 | 5803 | -2.04 | 0.06 |
| ForFemurWidth | 28 | F | Sox15 - control | 0.27 | 0.05 | 5803 | 5.07 | 2.94e-06 |
| ForFemurWidth | 25 | M | bab1 - control | 0.15 | 0.08 | 5803 | 1.94 | 0.09 |
| ForFemurWidth | 25 | M | CG13285 - control | 0.04 | 0.03 | 5803 | 1.17 | 0.28 |
| ForFemurWidth | 25 | M | CG30457 - control | 0.04 | 0.03 | 5803 | 1.39 | 0.23 |
| ForFemurWidth | 25 | M | dysf - control | -0.01 | 0.03 | 5803 | -0.33 | 0.74 |
| ForFemurWidth | 25 | M | grn - control | 0.10 | 0.03 | 5803 | 3.28 | 3.63e-03 |
| ForFemurWidth | 25 | M | otp - control | 0.07 | 0.03 | 5803 | 2.35 | 0.04 |
| ForFemurWidth | 25 | M | Sox15 - control | 0.17 | 0.04 | 5803 | 4.00 | 4.55e-04 |
| ForFemurWidth | 28 | M | bab1 - control | -1.96e-03 | 0.06 | 5803 | -0.04 | 0.97 |
| ForFemurWidth | 28 | M | CG13285 - control | -0.03 | 0.03 | 5803 | -0.85 | 0.56 |
| ForFemurWidth | 28 | M | CG30457 - control | 0.05 | 0.03 | 5803 | 1.72 | 0.20 |
| ForFemurWidth | 28 | M | dysf - control | -0.09 | 0.03 | 5803 | -2.81 | 0.03 |
| ForFemurWidth | 28 | M | grn - control | -0.04 | 0.03 | 5803 | -1.31 | 0.33 |
| ForFemurWidth | 28 | M | otp - control | 1.46e-03 | 0.03 | 5803 | 0.05 | 0.97 |
| ForFemurWidth | 28 | M | Sox15 - control | 0.18 | 0.08 | 5803 | 2.36 | 0.06 |
| MidFemurLength | 25 | F | bab1 - control | -0.09 | 0.01 | 5803 | -6.84 | 3.13e-11 |
| MidFemurLength | 25 | F | CG13285 - control | -0.02 | 0.01 | 5803 | -1.79 | 0.10 |
| MidFemurLength | 25 | F | CG30457 - control | -0.04 | 0.01 | 5803 | -2.76 | 0.01 |
| MidFemurLength | 25 | F | dysf - control | 0.02 | 0.01 | 5803 | 1.39 | 0.19 |
| MidFemurLength | 25 | F | grn - control | -0.03 | 0.01 | 5803 | -2.20 | 0.05 |
| MidFemurLength | 25 | F | otp - control | -8.63e-03 | 0.01 | 5803 | -0.68 | 0.50 |
| MidFemurLength | 25 | F | Sox15 - control | -0.13 | 0.02 | 5803 | -7.43 | 9.01e-13 |
| MidFemurLength | 28 | F | bab1 - control | -0.11 | 0.01 | 5803 | -7.63 | 1.96e-13 |
| MidFemurLength | 28 | F | CG13285 - control | 1.03e-03 | 0.01 | 5803 | 0.08 | 0.94 |

| | | | | | | | | |
|---|---|---|---|---|---|---|---|---|
| MidFemurLength | 28 | F | CG30457 - control | -3.42e-03 | 0.01 | 5803 | -0.27 | 0.92 |
| MidFemurLength | 28 | F | dysf - control | -0.02 | 0.01 | 5803 | -1.87 | 0.09 |
| MidFemurLength | 28 | F | grn - control | -0.09 | 0.01 | 5803 | -6.87 | 2.55e-11 |
| MidFemurLength | 28 | F | otp - control | -0.02 | 0.01 | 5803 | -2.23 | 0.05 |
| MidFemurLength | 28 | F | Sox15 - control | -0.11 | 0.02 | 5803 | -4.90 | 2.34e-06 |
| MidFemurLength | 25 | M | bab1 - control | -0.09 | 0.03 | 5803 | -2.80 | 0.02 |
| MidFemurLength | 25 | M | CG13285 - control | 8.07e-03 | 0.01 | 5803 | 0.62 | 0.53 |
| MidFemurLength | 25 | M | CG30457 - control | 0.03 | 0.01 | 5803 | 2.14 | 0.08 |
| MidFemurLength | 25 | M | dysf - control | 0.02 | 0.01 | 5803 | 1.54 | 0.17 |
| MidFemurLength | 25 | M | grn - control | 8.29e-03 | 0.01 | 5803 | 0.65 | 0.53 |
| MidFemurLength | 25 | M | otp - control | 0.02 | 0.01 | 5803 | 1.59 | 0.17 |
| MidFemurLength | 25 | M | Sox15 - control | -0.20 | 0.02 | 5803 | -11.00 | 5.15e-27 |
| MidFemurLength | 28 | M | bab1 - control | -0.13 | 0.02 | 5803 | -5.59 | 8.11e-08 |
| MidFemurLength | 28 | M | CG13285 - control | -0.02 | 0.01 | 5803 | -1.36 | 0.18 |
| MidFemurLength | 28 | M | CG30457 - control | -0.03 | 0.01 | 5803 | -2.44 | 0.02 |
| MidFemurLength | 28 | M | dysf - control | -0.04 | 0.01 | 5803 | -3.15 | 2.91e-03 |
| MidFemurLength | 28 | M | grn - control | -0.07 | 0.01 | 5803 | -5.22 | 4.34e-07 |
| MidFemurLength | 28 | M | otp - control | -0.03 | 0.01 | 5803 | -2.55 | 0.01 |
| MidFemurLength | 28 | M | Sox15 - control | -0.24 | 0.03 | 5803 | -7.64 | 1.74e-13 |
| MidFemurWidth | 25 | F | bab1 - control | 0.03 | 0.03 | 5803 | 0.86 | 0.39 |
| MidFemurWidth | 25 | F | CG13285 - control | -0.04 | 0.03 | 5803 | -1.40 | 0.19 |
| MidFemurWidth | 25 | F | CG30457 - control | -0.04 | 0.03 | 5803 | -1.39 | 0.19 |
| MidFemurWidth | 25 | F | dysf - control | -0.07 | 0.03 | 5803 | -2.39 | 0.06 |
| MidFemurWidth | 25 | F | grn - control | -0.06 | 0.03 | 5803 | -2.12 | 0.08 |
| MidFemurWidth | 25 | F | otp - control | -0.04 | 0.03 | 5803 | -1.47 | 0.19 |
| MidFemurWidth | 25 | F | Sox15 - control | 0.14 | 0.04 | 5803 | 3.57 | 2.54e-03 |
| MidFemurWidth | 28 | F | bab1 - control | 0.12 | 0.03 | 5803 | 3.68 | 1.65e-03 |
| MidFemurWidth | 28 | F | CG13285 - control | -0.03 | 0.03 | 5803 | -0.98 | 0.58 |
| MidFemurWidth | 28 | F | CG30457 - control | 2.24e-03 | 0.03 | 5803 | 0.07 | 0.94 |
| MidFemurWidth | 28 | F | dysf - control | -0.02 | 0.03 | 5803 | -0.53 | 0.70 |
| MidFemurWidth | 28 | F | grn - control | -0.03 | 0.03 | 5803 | -0.97 | 0.58 |
| MidFemurWidth | 28 | F | otp - control | 0.02 | 0.03 | 5803 | 0.71 | 0.67 |
| MidFemurWidth | 28 | F | Sox15 - control | 0.20 | 0.06 | 5803 | 3.50 | 1.65e-03 |
| MidFemurWidth | 25 | M | bab1 - control | 0.12 | 0.07 | 5803 | 1.59 | 0.39 |
| MidFemurWidth | 25 | M | CG13285 - control | -0.04 | 0.03 | 5803 | -1.19 | 0.41 |
| MidFemurWidth | 25 | M | CG30457 - control | -0.04 | 0.03 | 5803 | -1.25 | 0.41 |
| MidFemurWidth | 25 | M | dysf - control | -0.01 | 0.03 | 5803 | -0.42 | 0.81 |
| MidFemurWidth | 25 | M | grn - control | -0.01 | 0.03 | 5803 | -0.39 | 0.81 |
| MidFemurWidth | 25 | M | otp - control | 8.61e-04 | 0.03 | 5803 | 0.03 | 0.98 |
| MidFemurWidth | 25 | M | Sox15 - control | 0.11 | 0.04 | 5803 | 2.62 | 0.06 |
| MidFemurWidth | 28 | M | bab1 - control | 0.03 | 0.05 | 5803 | 0.52 | 0.70 |
| MidFemurWidth | 28 | M | CG13285 - control | -0.04 | 0.03 | 5803 | -1.26 | 0.49 |
| MidFemurWidth | 28 | M | CG30457 - control | 0.11 | 0.03 | 5803 | 3.48 | 3.48e-03 |
| MidFemurWidth | 28 | M | dysf - control | 0.03 | 0.03 | 5803 | 1.03 | 0.53 |
| MidFemurWidth | 28 | M | grn - control | -9.90e-03 | 0.03 | 5803 | -0.31 | 0.75 |
| MidFemurWidth | 28 | M | otp - control | -0.02 | 0.03 | 5803 | -0.58 | 0.70 |
| MidFemurWidth | 28 | M | Sox15 - control | 0.18 | 0.09 | 5803 | 2.10 | 0.13 |
| Thorax | 25 | F | bab1 - control | -0.03 | 0.01 | 5803 | -2.44 | 0.03 |
| Thorax | 25 | F | CG13285 - control | 9.17e-04 | 0.01 | 5803 | 0.07 | 0.97 |

| | | | | | | | | |
|---|---|---|---|---|---|---|---|---|
| Thorax | 25 | F | CG30457 - control | 5.55e-03 | 0.01 | 5803 | 0.42 | 0.94 |
| Thorax | 25 | F | dysf - control | -5.59e-04 | 0.01 | 5803 | -0.04 | 0.97 |
| Thorax | 25 | F | grn - control | 0.03 | 0.01 | 5803 | 2.63 | 0.03 |
| Thorax | 25 | F | otp - control | 0.03 | 0.01 | 5803 | 2.59 | 0.03 |
| Thorax | 25 | F | Sox15 - control | -0.03 | 0.02 | 5803 | -1.58 | 0.20 |
| Thorax | 28 | F | bab1 - control | -0.08 | 0.01 | 5803 | -5.44 | 3.94e-07 |
| Thorax | 28 | F | CG13285 - control | -0.04 | 0.01 | 5803 | -2.76 | 0.01 |
| Thorax | 28 | F | CG30457 - control | -0.01 | 0.01 | 5803 | -0.84 | 0.56 |
| Thorax | 28 | F | dysf - control | -0.06 | 0.01 | 5803 | -4.47 | 2.74e-05 |
| Thorax | 28 | F | grn - control | -4.58e-03 | 0.01 | 5803 | -0.34 | 0.73 |
| Thorax | 28 | F | otp - control | -8.72e-03 | 0.01 | 5803 | -0.69 | 0.57 |
| Thorax | 28 | F | Sox15 - control | -0.09 | 0.02 | 5803 | -3.91 | 2.19e-04 |
| Thorax | 25 | M | bab1 - control | -0.09 | 0.03 | 5803 | -2.71 | 0.02 |
| Thorax | 25 | M | CG13285 - control | 0.01 | 0.01 | 5803 | 0.91 | 0.42 |
| Thorax | 25 | M | CG30457 - control | 0.01 | 0.01 | 5803 | 0.97 | 0.42 |
| Thorax | 25 | M | dysf - control | -0.03 | 0.01 | 5803 | -2.62 | 0.02 |
| Thorax | 25 | M | grn - control | 2.70e-03 | 0.01 | 5803 | 0.20 | 0.84 |
| Thorax | 25 | M | otp - control | -0.02 | 0.01 | 5803 | -1.42 | 0.27 |
| Thorax | 25 | M | Sox15 - control | -0.10 | 0.02 | 5803 | -5.76 | 6.27e-08 |
| Thorax | 28 | M | bab1 - control | -0.14 | 0.02 | 5803 | -6.09 | 8.20e-09 |
| Thorax | 28 | M | CG13285 - control | -0.02 | 0.01 | 5803 | -1.54 | 0.17 |
| Thorax | 28 | M | CG30457 - control | -0.01 | 0.01 | 5803 | -0.89 | 0.43 |
| Thorax | 28 | M | dysf - control | -0.05 | 0.01 | 5803 | -3.49 | 1.69e-03 |
| Thorax | 28 | M | grn - control | -0.03 | 0.01 | 5803 | -2.09 | 0.06 |
| Thorax | 28 | M | otp - control | -0.05 | 0.01 | 5803 | -3.28 | 2.43e-03 |
| Thorax | 28 | M | Sox15 - control | -0.03 | 0.03 | 5803 | -0.79 | 0.43 |

Table S3: Model estimates for forefemur length contrasted between UAS-RNAi genes and *Pen*$^{NP3666}$*-Gal4*. Contrasts performed using emmeans.

| **contrast** | **temp** | **sex** | **estimate** | **SE** | **df** | **Lower.CL** | **upper.CL** |
|---|---|---|---|---|---|---|---|
| bab1 - control | 25 | F | -0.0375389876 | 0.01613063 | 1083 | -0.08000523 | 0.004927259 |
| CG13285 - control | 25 | F | 0.0011269694 | 0.01624377 | 1083 | -0.04163713 | 0.043891070 |
| CG30457 - control | 25 | F | -0.0078946882 | 0.01574506 | 1083 | -0.04934585 | 0.033556471 |
| dysf - control | 25 | F | 0.0547771303 | 0.01564277 | 1083 | 0.01359527 | 0.095958990 |
| grn - control | 25 | F | -0.0234483297 | 0.01566639 | 1083 | -0.06469239 | 0.017795735 |
| otp - control | 25 | F | -0.0084377792 | 0.01566576 | 1083 | -0.04968016 | 0.032804605 |
| Sox15 - control | 25 | F | -0.1288173762 | 0.01960156 | 1083 | -0.18042134 | -0.077213412 |
| bab1 - control | 28 | F | -0.0917693423 | 0.01713551 | 1083 | -0.13688106 | -0.046657628 |
| CG13285 - control | 28 | F | 0.0007136154 | 0.01582474 | 1083 | -0.04094733 | 0.042374557 |
| CG30457 - control | 28 | F | -0.0055084783 | 0.01577260 | 1083 | -0.04703214 | 0.036015184 |
| dysf - control | 28 | F | -0.0162827001 | 0.01584253 | 1083 | -0.05799046 | 0.025425062 |
| grn - control | 28 | F | -0.0714064010 | 0.01577033 | 1083 | -0.11292409 | -0.029888715 |

| | | | | | | | |
|---|---|---|---|---|---|---|---|
| otp - control | 28 | F | -0.0363353894 | 0.01577209 | 1083 | -0.07785770 | 0.005186923 |
| Sox15 - control | 28 | F | -0.1426575817 | 0.02167124 | 1083 | -0.19971029 | -0.085604878 |
| bab1 - control | 25 | M | -0.0408074119 | 0.02918856 | 1083 | -0.11765055 | 0.036035728 |
| CG13285 - control | 25 | M | -0.0087280910 | 0.01625184 | 1083 | -0.05151343 | 0.034057252 |
| CG30457 - control | 25 | M | -0.0114694707 | 0.01565165 | 1083 | -0.05267472 | 0.029735778 |
| dysf - control | 25 | M | 0.0066387459 | 0.01567410 | 1083 | -0.03462560 | 0.047903094 |
| grn - control | 25 | M | -0.0234330708 | 0.01564839 | 1083 | -0.06462973 | 0.017763588 |
| otp - control | 25 | M | -0.0120650896 | 0.01565619 | 1083 | -0.05328228 | 0.029152102 |
| Sox15 - control | 25 | M | -0.1204625608 | 0.01886189 | 1083 | -0.17011924 | -0.070805884 |
| bab1 - control | 28 | M | -0.0332852146 | 0.02272880 | 1083 | -0.09312210 | 0.026551670 |
| CG13285 - control | 28 | M | -0.0039771831 | 0.01579660 | 1083 | -0.04556403 | 0.037609667 |
| CG30457 - control | 28 | M | -0.0003061673 | 0.01586645 | 1083 | -0.04207691 | 0.041464578 |
| dysf - control | 28 | M | -0.0243400768 | 0.01590747 | 1083 | -0.06621881 | 0.017538660 |
| grn - control | 28 | M | -0.0664262469 | 0.01587354 | 1083 | -0.10821567 | -0.024636828 |
| otp - control | 28 | M | -0.0047311627 | 0.01651417 | 1083 | -0.04820711 | 0.038744784 |
| Sox15 - control | 28 | M | -0.1391134595 | 0.02916397 | 1083 | -0.21589186 | -0.062335061 |

Table S4: Model estimates for forefemur width contrasted between UAS-RNAi genes and *Pen$^{NP3666}$-Gal4*. Contrasts performed using emmeans.

| contrast | temp | sex | estimate | SE | df | Lower.CL | upper.CL |
|---|---|---|---|---|---|---|---|
| bab1 - control | 25 | F | 0.094611369 | 0.03588458 | 1074 | 0.0001386695 | 0.189084068 |
| CG13285 - control | 25 | F | -0.049012888 | 0.03624164 | 1074 | -0.1444255952 | 0.046399820 |
| CG30457 - control | 25 | F | -0.020289078 | 0.03463077 | 1074 | -0.1114608905 | 0.070882734 |
| dysf - control | 25 | F | -0.061504692 | 0.03428173 | 1074 | -0.1517575890 | 0.028748205 |
| grn - control | 25 | F | -0.023262941 | 0.03470732 | 1074 | -0.1146362762 | 0.068110395 |
| otp - control | 25 | F | 0.066107797 | 0.03435827 | 1074 | -0.0243466072 | 0.156562200 |
| Sox15 - control | 25 | F | 0.177377395 | 0.04708123 | 1074 | 0.0534274748 | 0.301327316 |
| bab1 - control | 28 | F | 0.148925771 | 0.03861650 | 1074 | 0.0472608125 | 0.250590729 |
| CG13285 - control | 28 | F | -0.088202287 | 0.03480112 | 1074 | -0.1798225607 | 0.003417988 |
| CG30457 - control | 28 | F | 0.017442770 | 0.03472036 | 1074 | -0.0739648918 | 0.108850432 |
| dysf - control | 28 | F | 0.084116145 | 0.03495065 | 1074 | -0.0078978100 | 0.176130100 |
| grn - control | 28 | F | -0.045919208 | 0.03471467 | 1074 | -0.1373118884 | 0.045473471 |
| otp - control | 28 | F | -0.042676080 | 0.03471697 | 1074 | -0.1340748238 | 0.048722664 |
| Sox15 - control | 28 | F | 0.282723118 | 0.05341550 | 1074 | 0.1420970846 | 0.423349151 |
| bab1 - control | 25 | M | 0.157870801 | 0.07554723 | 1074 | -0.0410210413 | 0.356762644 |
| CG13285 - control | 25 | M | 0.019276816 | 0.03627292 | 1074 | -0.0762182428 | 0.114771874 |
| CG30457 - control | 25 | M | 0.037675970 | 0.03431590 | 1074 | -0.0526668937 | 0.128018834 |

| | | | | | | | |
|---|---|---|---|---|---|---|---|
| dysf - control | 25 | M | -0.009630205 | 0.03473338 | 1074 | -0.1010721412 | 0.081811732 |
| grn - control | 25 | M | 0.098036301 | 0.03430513 | 1074 | 0.0077218148 | 0.188350787 |
| otp - control | 25 | M | 0.071957663 | 0.03433067 | 1074 | -0.0184240648 | 0.162339390 |
| Sox15 - control | 25 | M | 0.188016790 | 0.04601418 | 1074 | 0.0668760753 | 0.309157505 |
| bab1 - control | 28 | M | 0.021891240 | 0.05664375 | 1074 | -0.1272337466 | 0.171016227 |
| CG13285 - control | 28 | M | -0.018731631 | 0.03545567 | 1074 | -0.1120751388 | 0.074611877 |
| CG30457 - control | 28 | M | 0.059363936 | 0.03472619 | 1074 | -0.0320590852 | 0.150786957 |
| dysf - control | 28 | M | -0.086570653 | 0.03522299 | 1074 | -0.1793015733 | 0.006160266 |
| grn - control | 28 | M | -0.033613239 | 0.03510253 | 1074 | -0.1260270511 | 0.058800573 |
| otp - control | 28 | M | 0.005257614 | 0.03774336 | 1074 | -0.0941086575 | 0.104623886 |
| Sox15 - control | 28 | M | 0.190262820 | 0.07550569 | 1074 | -0.0085196511 | 0.389045290 |
