## Supplemental Figures2 for "Genetic architecture of the developing forelegs of *Drosophila prolongata*; an exaggerated weapon and ornament"

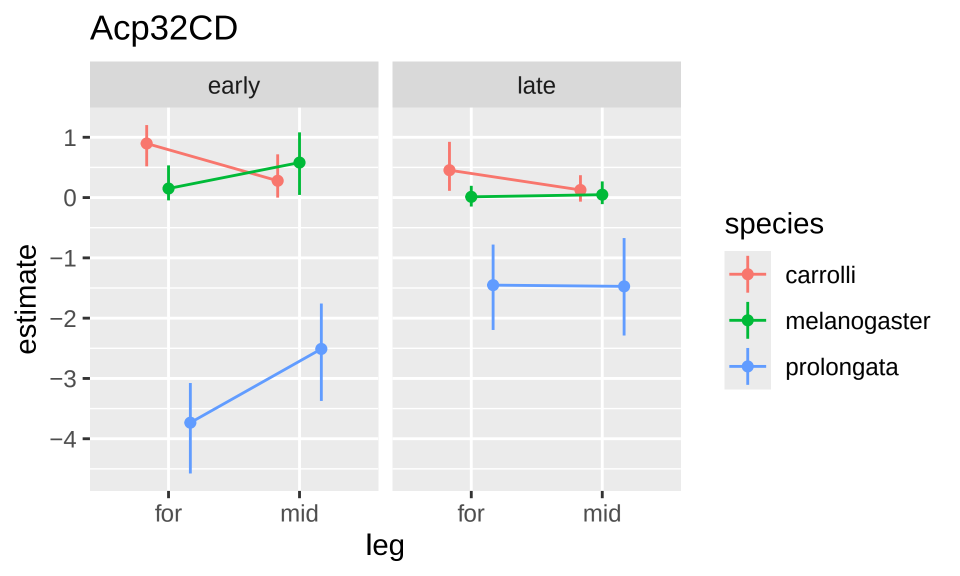

Acp32CD
early
late
estimate
1
0
−1
−2
−3
−4
for
mid
for
mid
leg
species
carrolli
melanogaster
prolongata

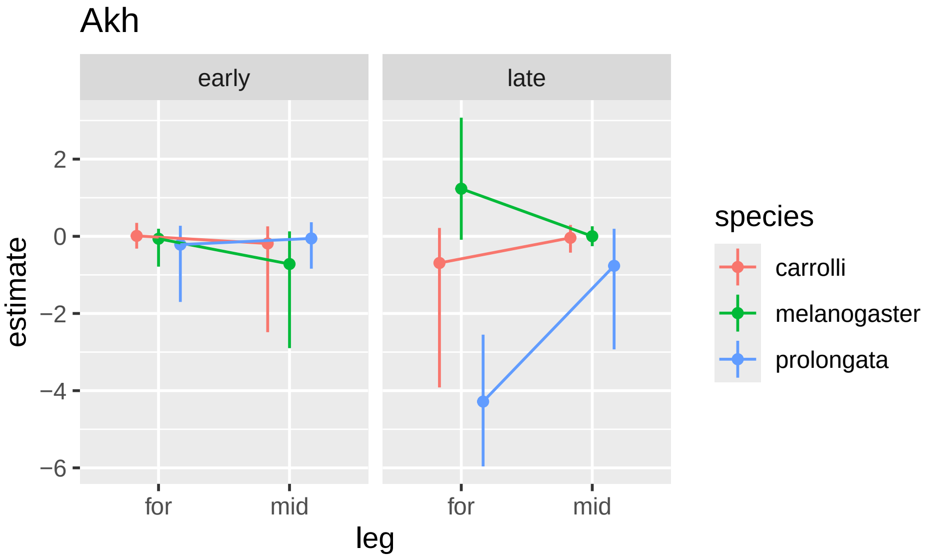

Akh
early
late
2
0
−2
−4
−6
estimate
for
mid
for
mid
leg
species
carrolli
melanogaster
prolongata

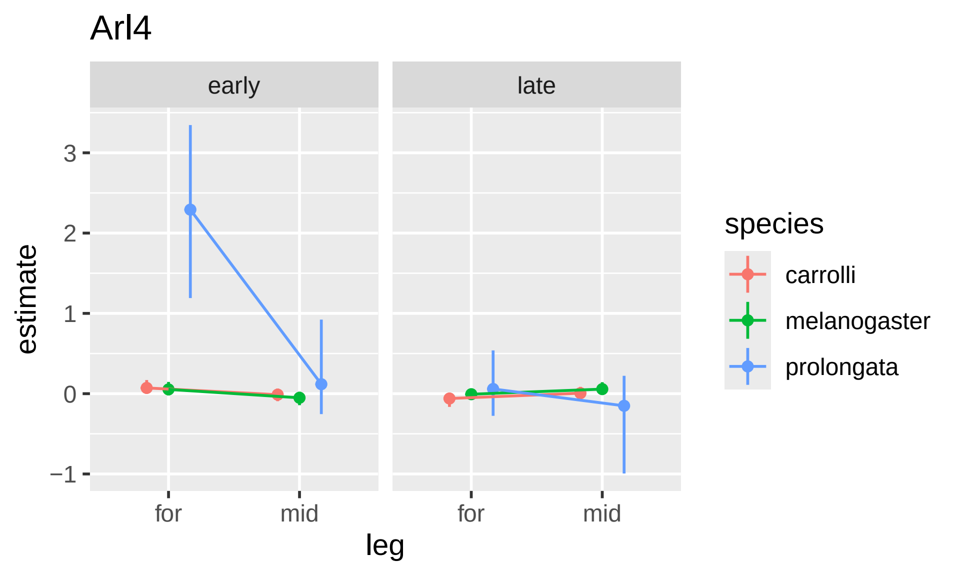

Arl4
early
late
estimate
3
2
1
0
−1
for
mid
for
mid
leg
species
carrolli
melanogaster
prolongata

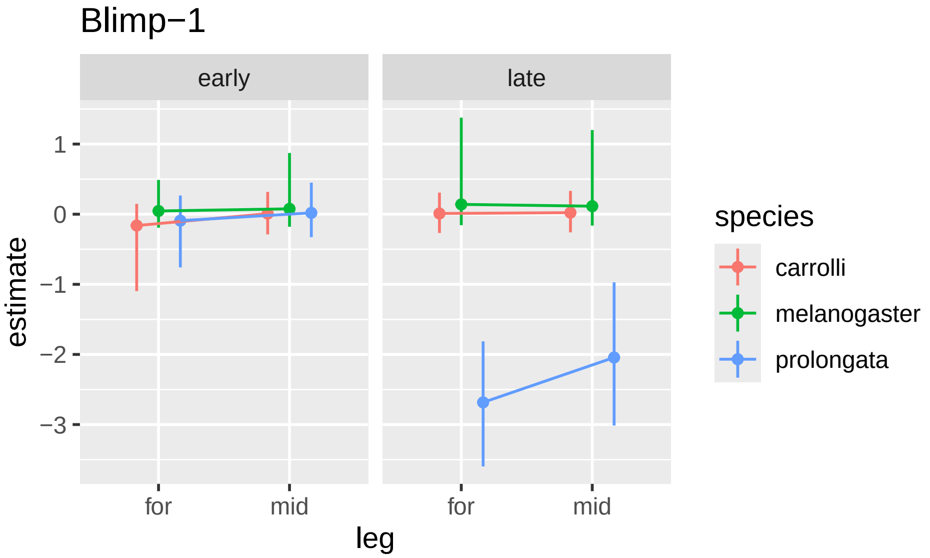

Blimp−1
early
late
1
0
−1
−2
−3
estimate
for
mid
for
mid
leg
species
carrolli
melanogaster
prolongata

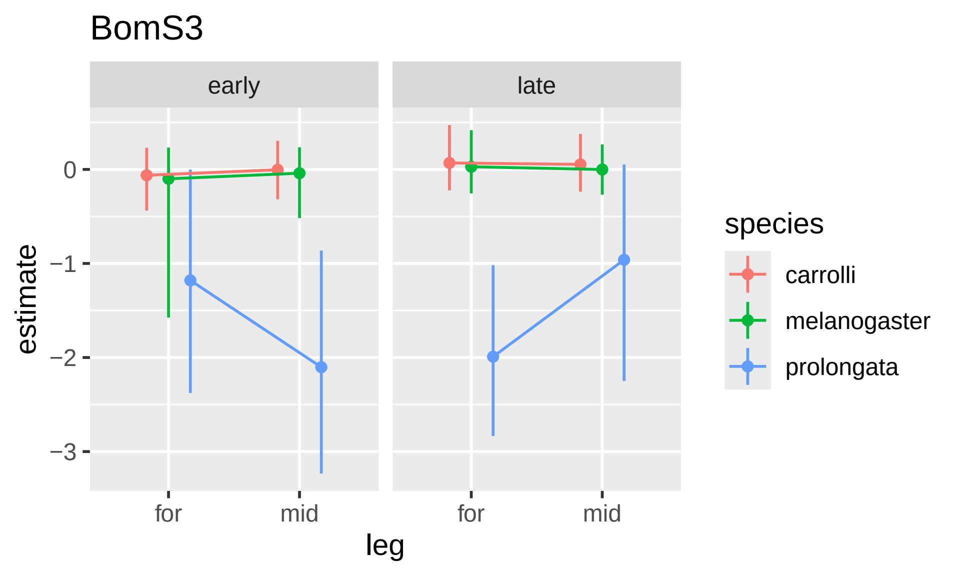
BomS3
early
late
0
−1
−2
−3
estimate
for
mid
for
mid
leg
species
carrolli
melanogaster
prolongata

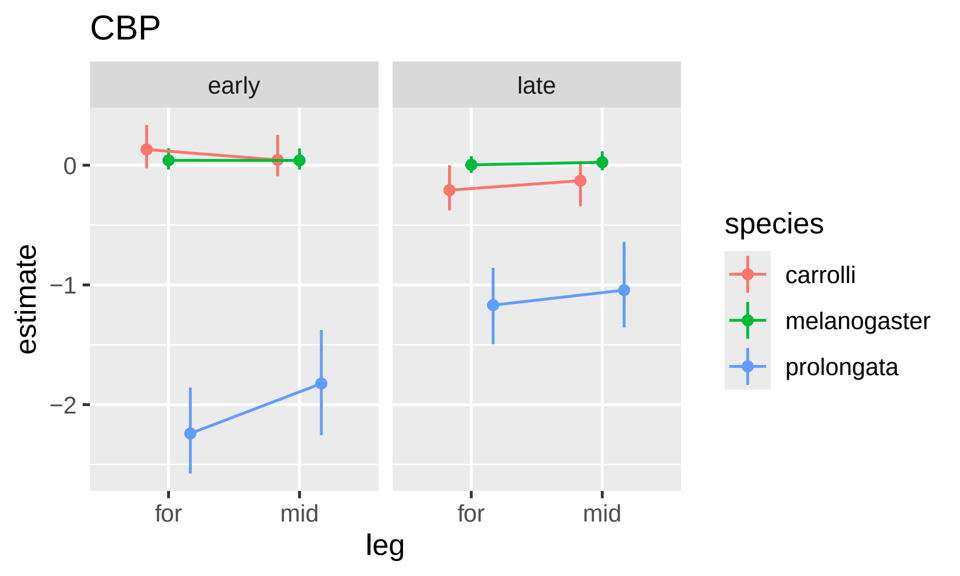

CBP
early
late
0
−1
−2
estimate
for
mid
for
mid
leg
species
carrolli
melanogaster
prolongata

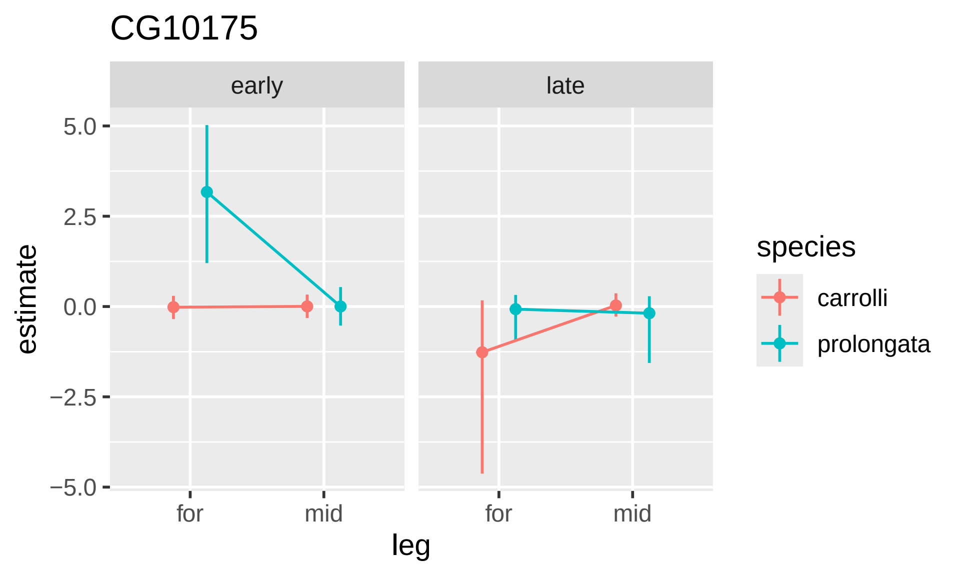
CG10175
early
late
5.0
2.5
0.0
−2.5
−5.0
estimate
for
mid
for
mid
leg
species
carrolli
prolongata

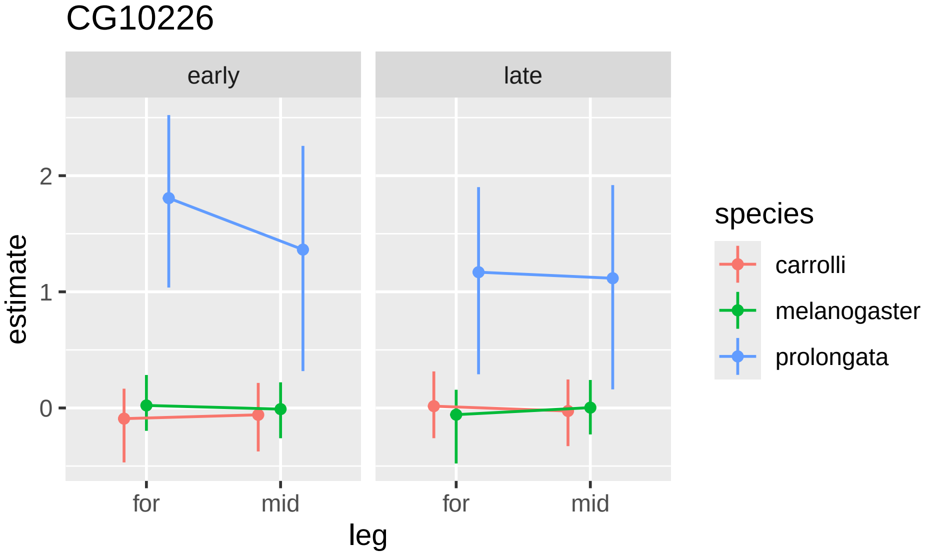
CG10226
early
late
estimate
2
1
0
for
mid
for
mid
leg
species
carrolli
melanogaster
prolongata

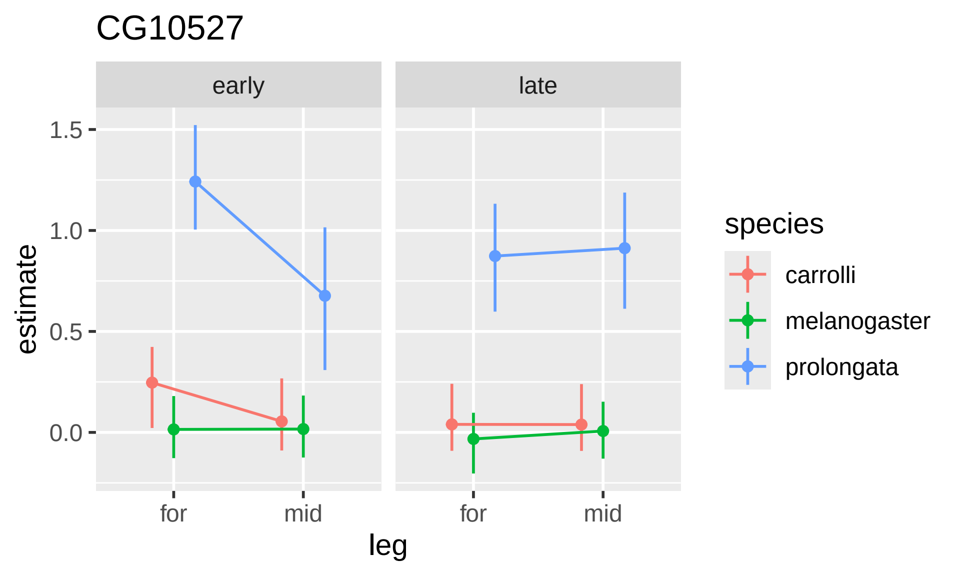

CG10527
early
late
1.5
1.0
0.5
0.0
estimate
for
mid
for
mid
leg
species
carrolli
melanogaster
prolongata

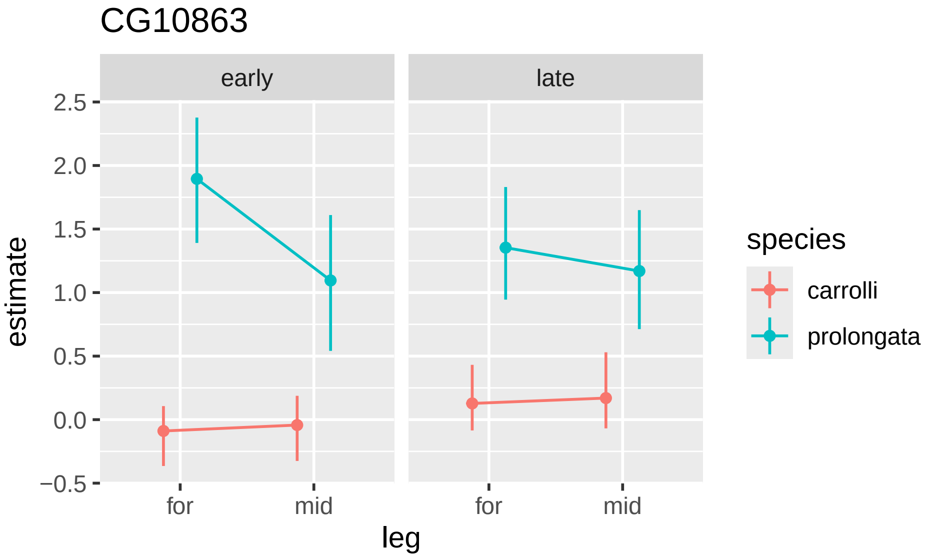
CG10863
early
late
2.5
2.0
1.5
1.0
0.5
0.0
−0.5
estimate
for
mid
for
mid
leg
species
carrolli
prolongata

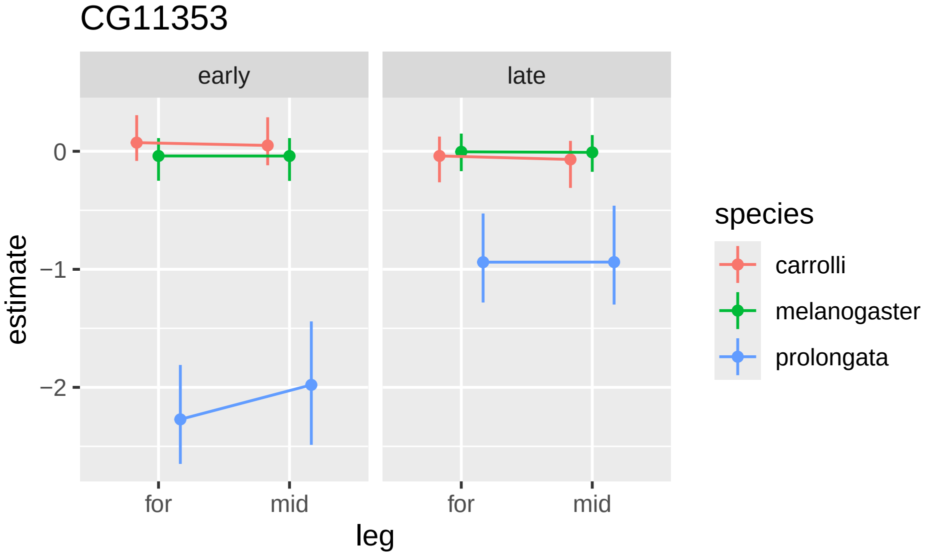
CG11353
early
late
0
−1
−2
estimate
for
mid
for
mid
leg
species
carrolli
melanogaster
prolongata

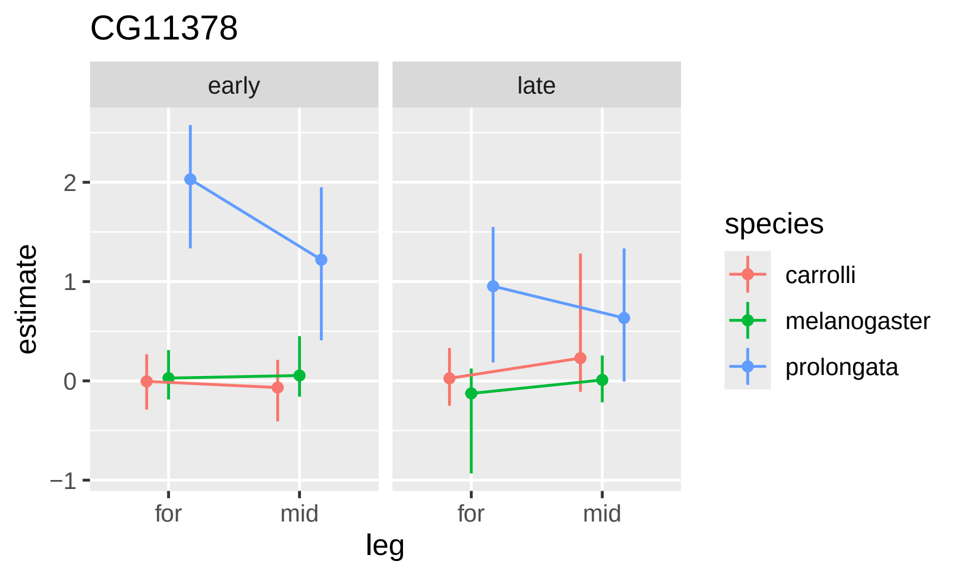

CG11378
early
late
estimate
2
1
0
−1
for
mid
for
mid
leg
species
carrolli
melanogaster
prolongata

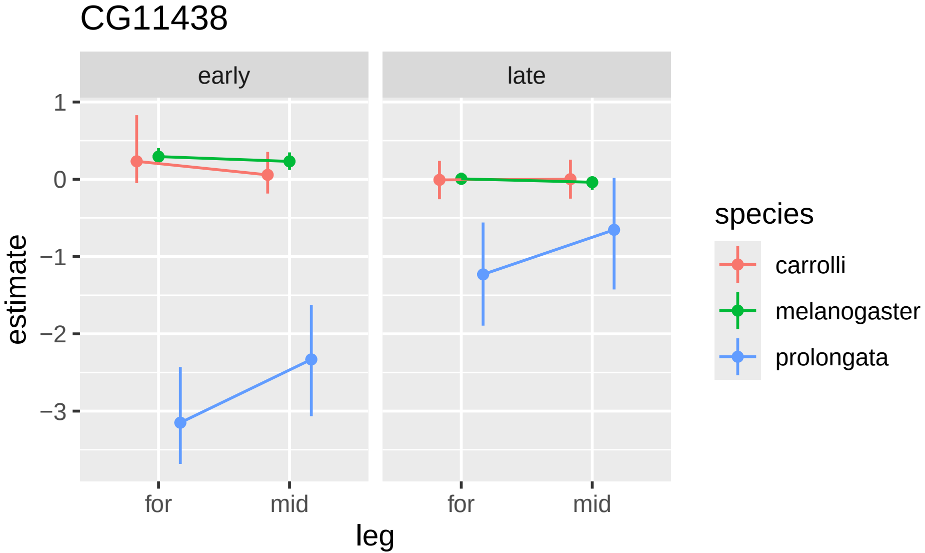
CG11438
early
late
1
0
−1
−2
−3
estimate
for
mid
for
mid
leg
species
carrolli
melanogaster
prolongata

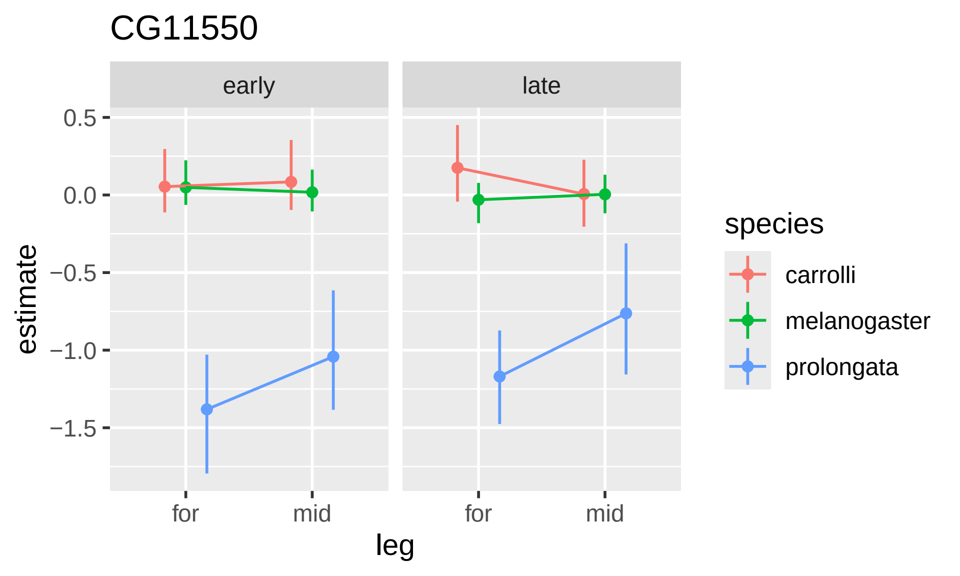

CG11550
early
late
0.5
0.0
−0.5
−1.0
−1.5
estimate
for
mid
for
mid
leg
species
carrolli
melanogaster
prolongata

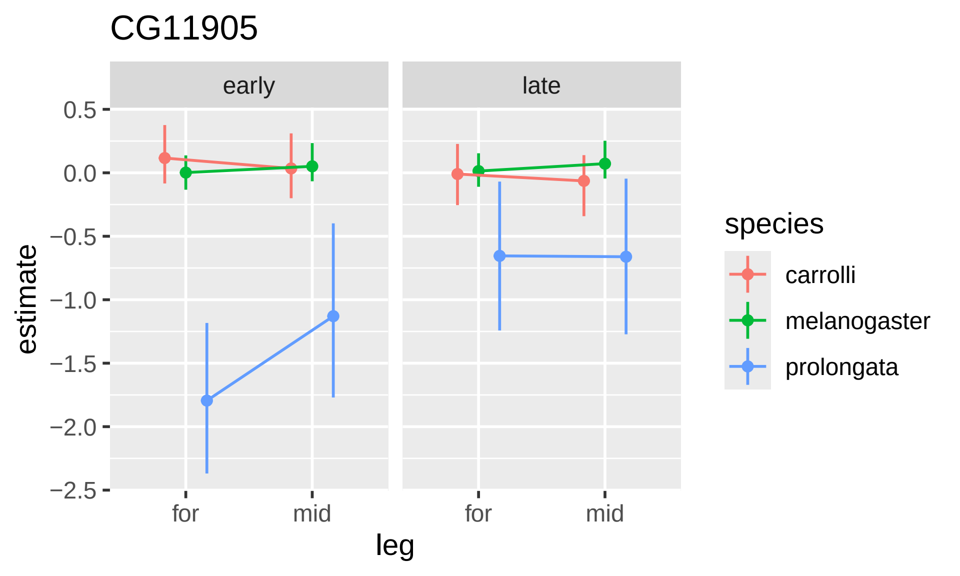

CG11905
early
late
0.5
0.0
−0.5
−1.0
−1.5
−2.0
−2.5
estimate
for
mid
for
mid
leg
species
carrolli
melanogaster
prolongata

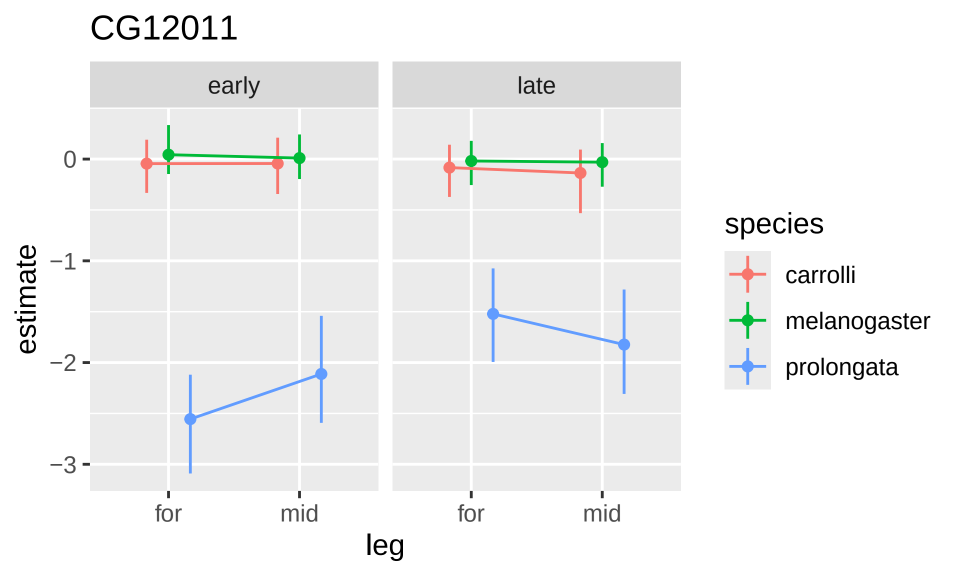

CG12011
early
late
0
−1
−2
−3
estimate
for
mid
for
mid
leg
species
carrolli
melanogaster
prolongata

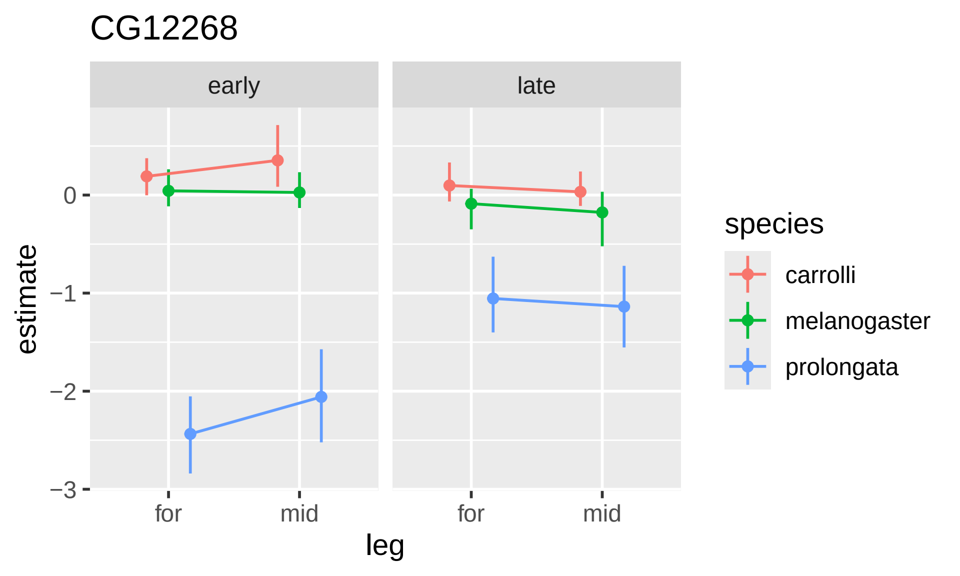

CG12268
early
late
estimate
0
−1
−2
−3
for
mid
for
mid
leg
species
carrolli
melanogaster
prolongata

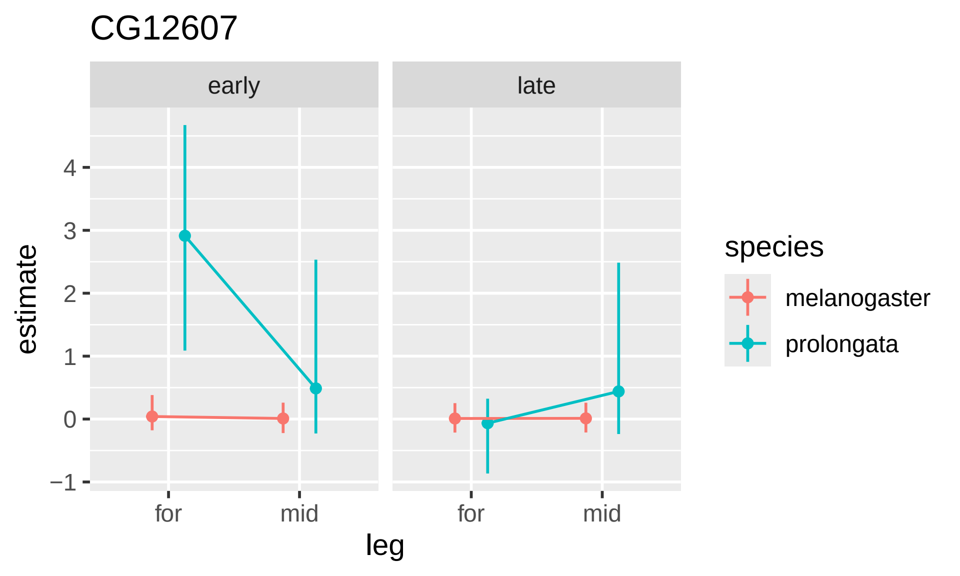

CG12607
early
late
estimate
4
3
2
1
0
−1
for
mid
for
mid
leg
species
melanogaster
prolongata

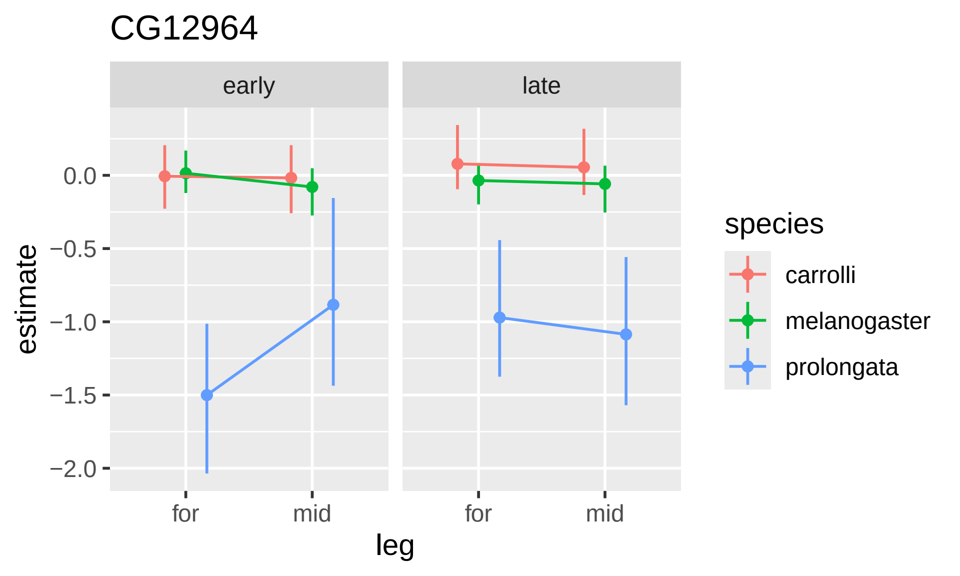

CG12964
early
late
0.0
−0.5
−1.0
−1.5
−2.0
estimate
for
mid
for
mid
leg
species
carrolli
melanogaster
prolongata

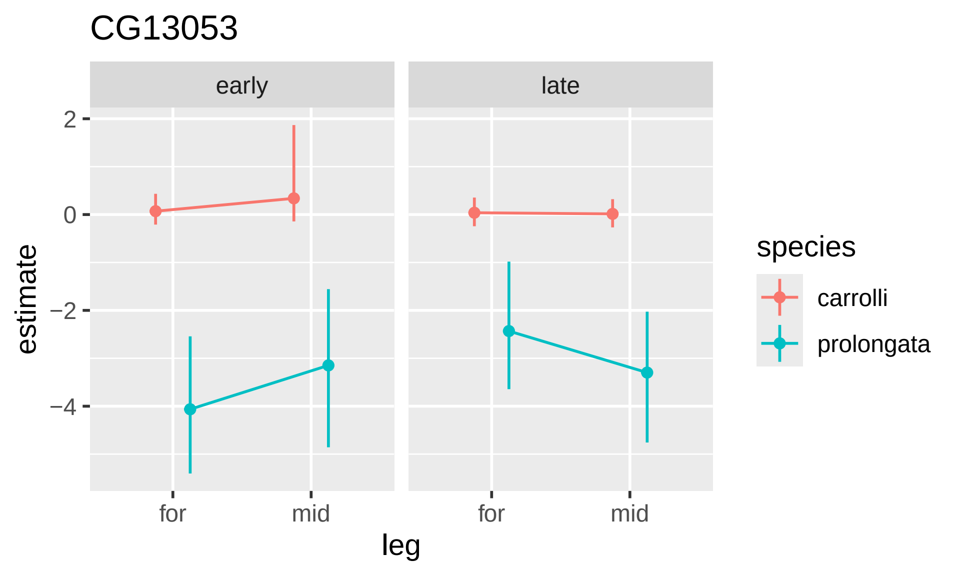

CG13053
early
late
2
0
−2
−4
estimate
for
mid
for
mid
leg
species
carrolli
prolongata

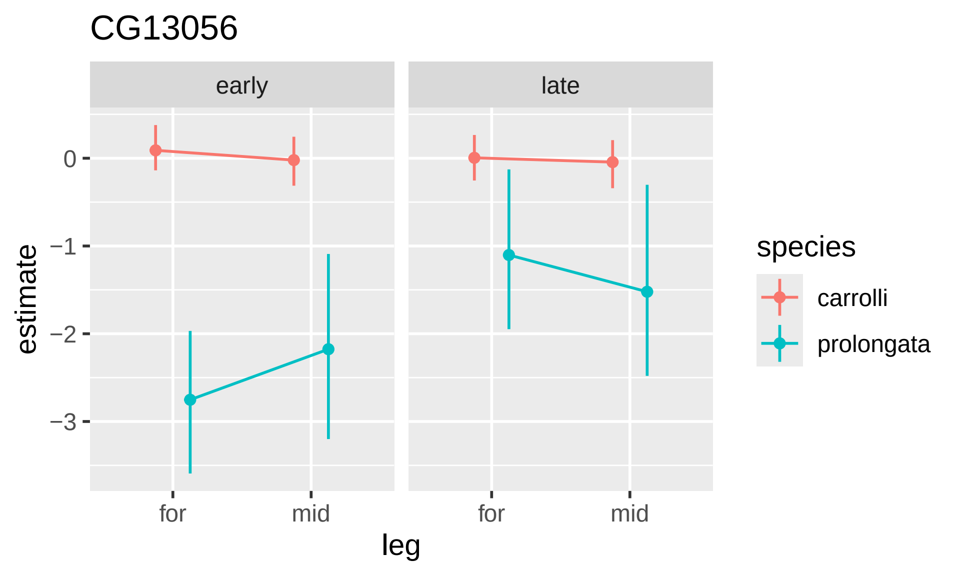

CG13056
early
late
0
−1
−2
−3
estimate
for
mid
for
mid
leg
species
carrolli
prolongata

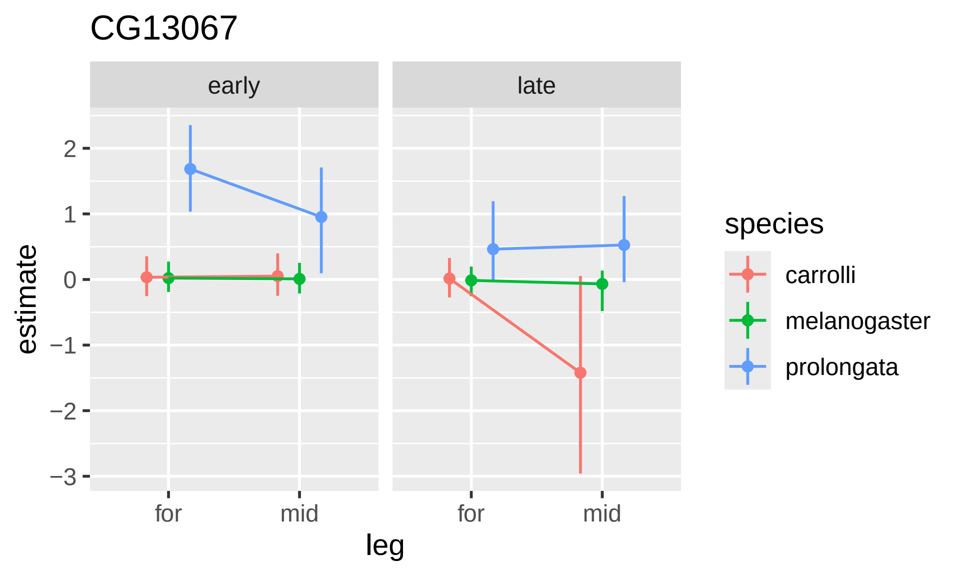

CG13067
early
late
estimate
2
1
0
−1
−2
−3
for
mid
for
mid
leg
species
carrolli
melanogaster
prolongata

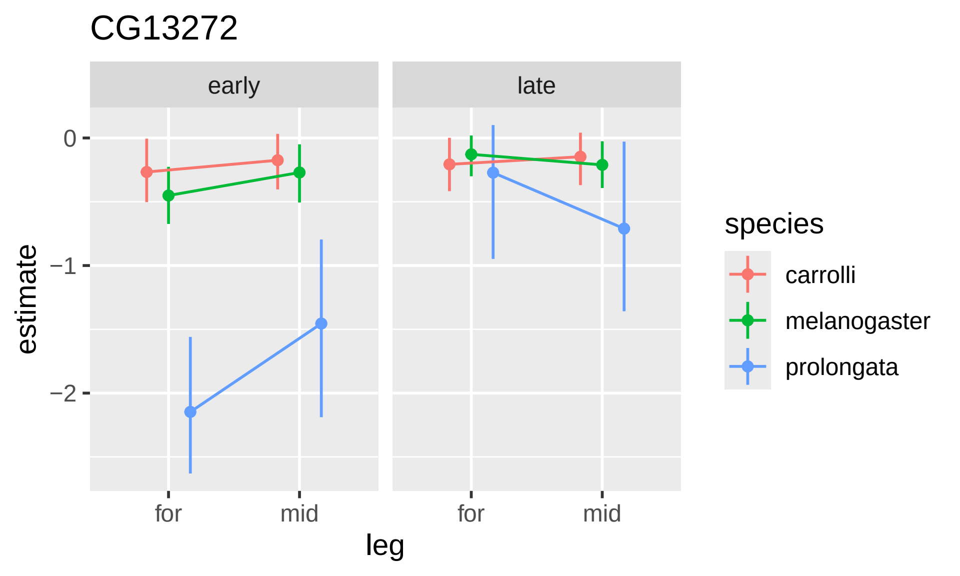
CG13272
early
late
estimate
0
−1
−2
for
mid
for
mid
leg
species
carrolli
melanogaster
prolongata

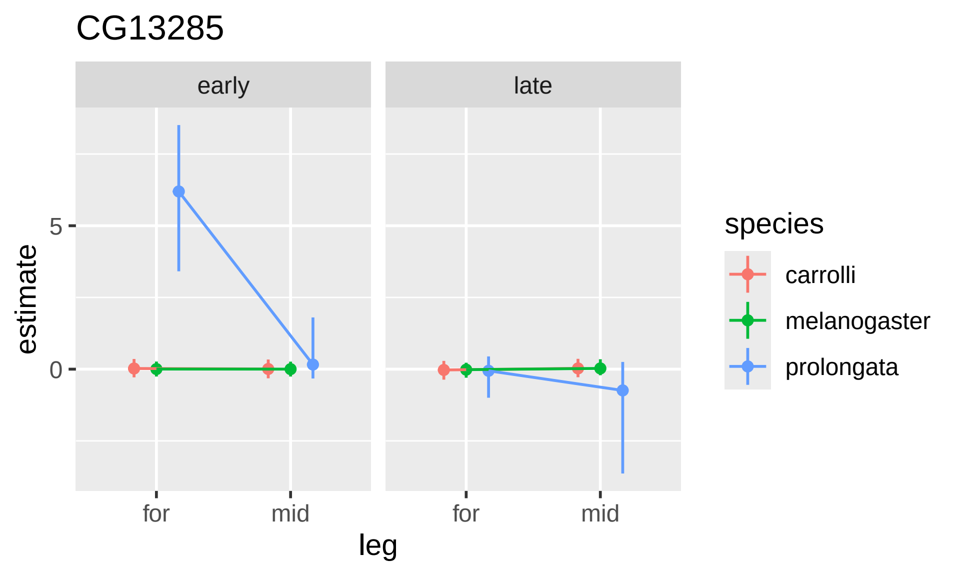
CG13285
early
late
estimate
5
0
for
mid
for
mid
leg
species
carrolli
melanogaster
prolongata

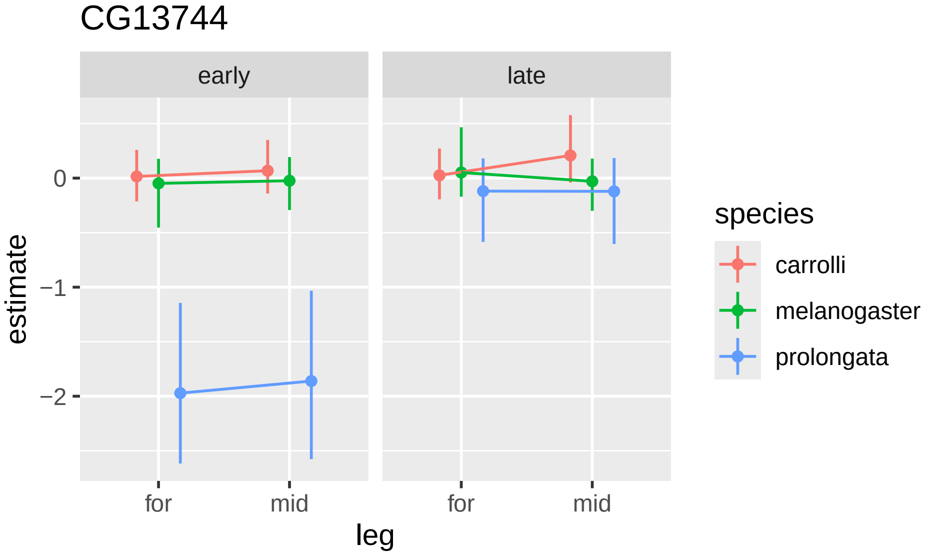

CG13744
early
late
estimate
0
−1
−2
for
mid
for
mid
leg
species
carrolli
melanogaster
prolongata

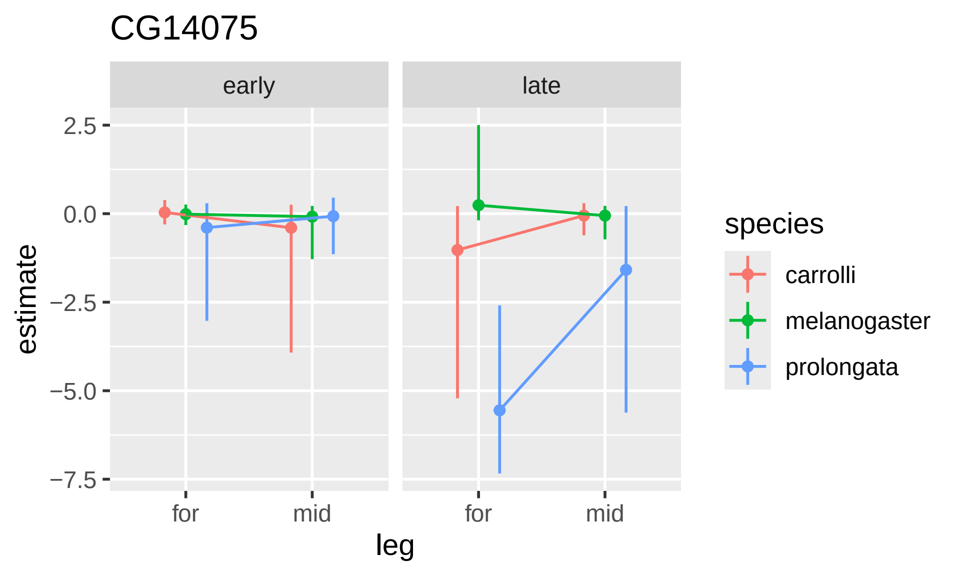

CG14075
early
late
2.5
0.0
−2.5
−5.0
−7.5
estimate
for
mid
for
mid
leg
species
carrolli
melanogaster
prolongata

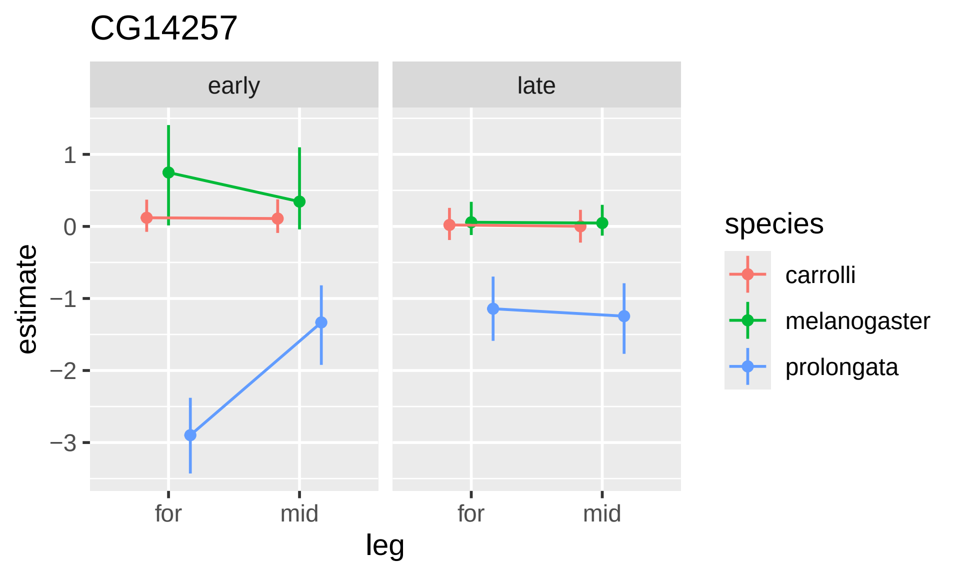

CG14257
early
late
1
0
−1
−2
−3
estimate
for
mid
for
mid
leg
species
carrolli
melanogaster
prolongata

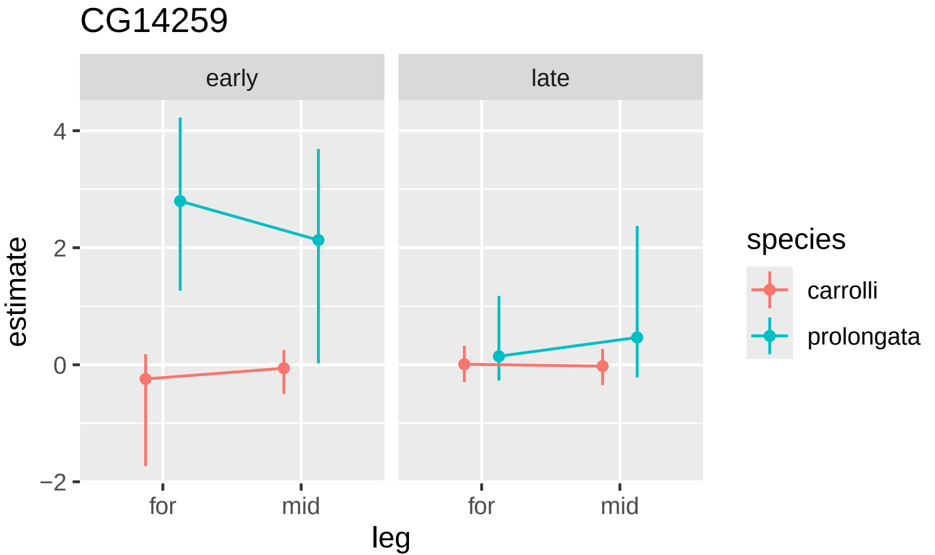

CG14259
early
late
estimate
4
2
0
−2
for
mid
for
mid
leg
species
carrolli
prolongata

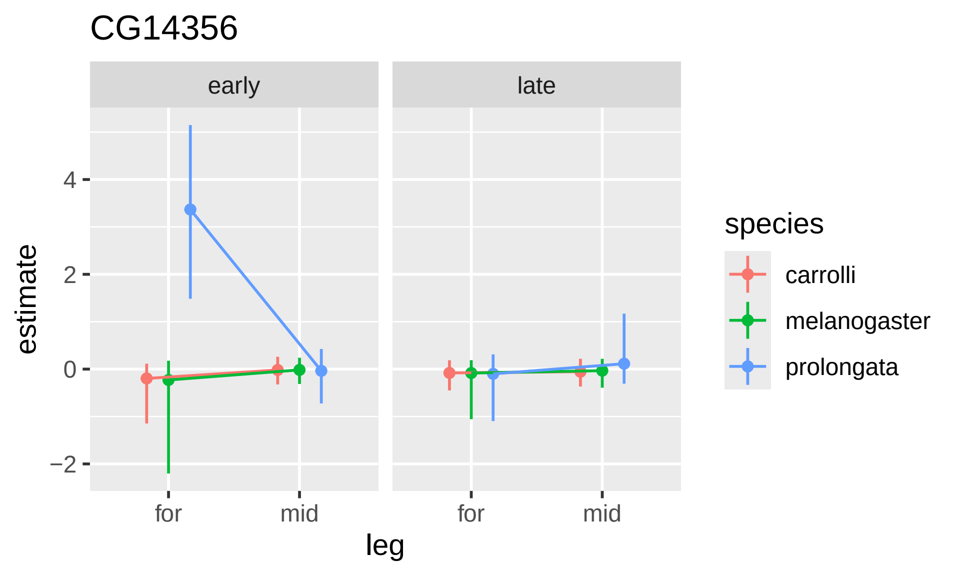
CG14356
early
late
4
2
0
−2
estimate
for
mid
for
mid
leg
species
carrolli
melanogaster
prolongata

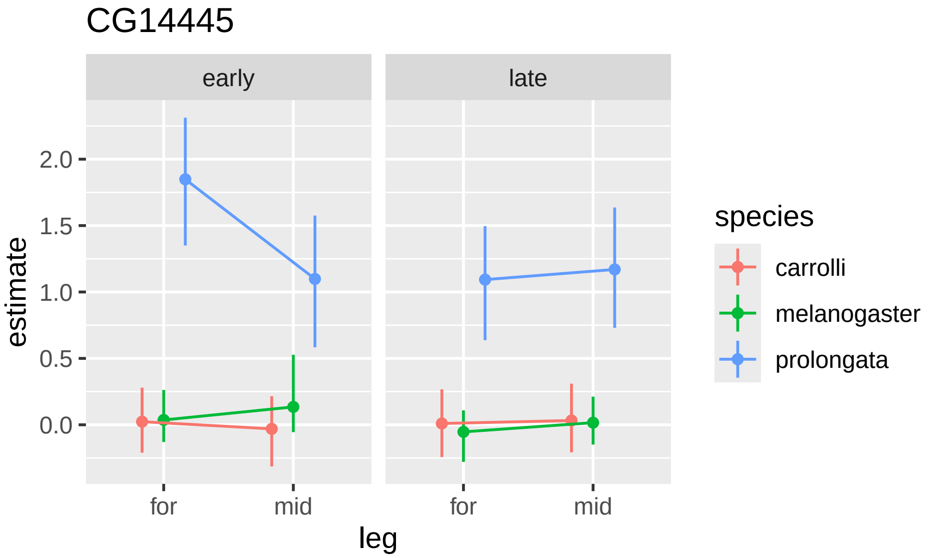
CG14445
early
late
2.0
1.5
1.0
0.5
0.0
estimate
for
mid
for
mid
leg
species
carrolli
melanogaster
prolongata

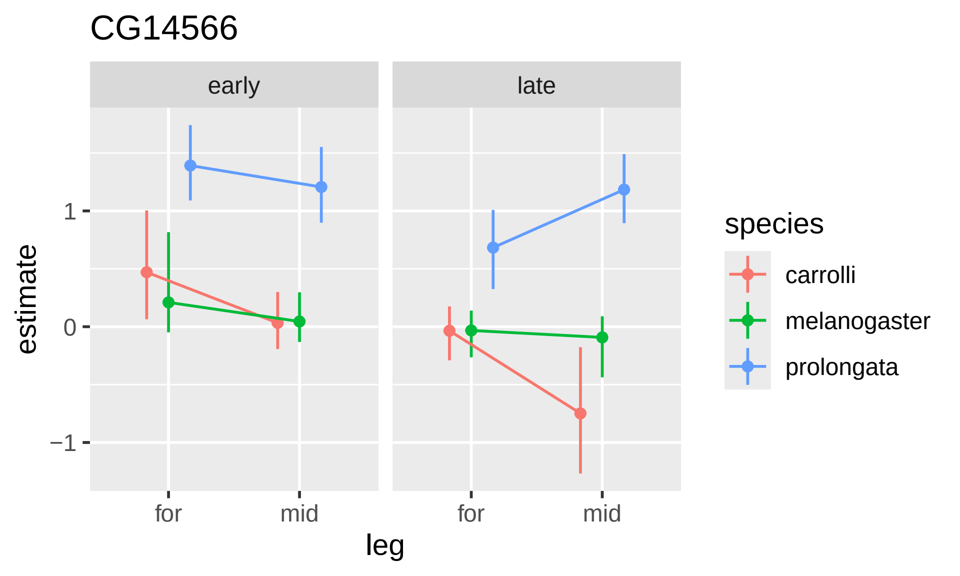
CG14566
early
late
1
0
−1
estimate
for
mid
for
mid
leg
species
carrolli
melanogaster
prolongata

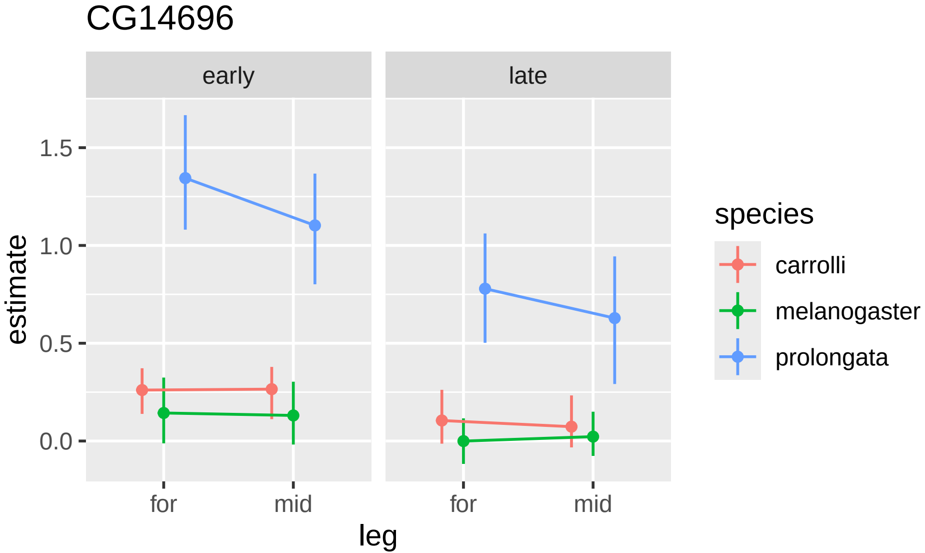

CG14696
early
late
1.5
1.0
0.5
0.0
estimate
for
mid
for
mid
leg
species
carrolli
melanogaster
prolongata

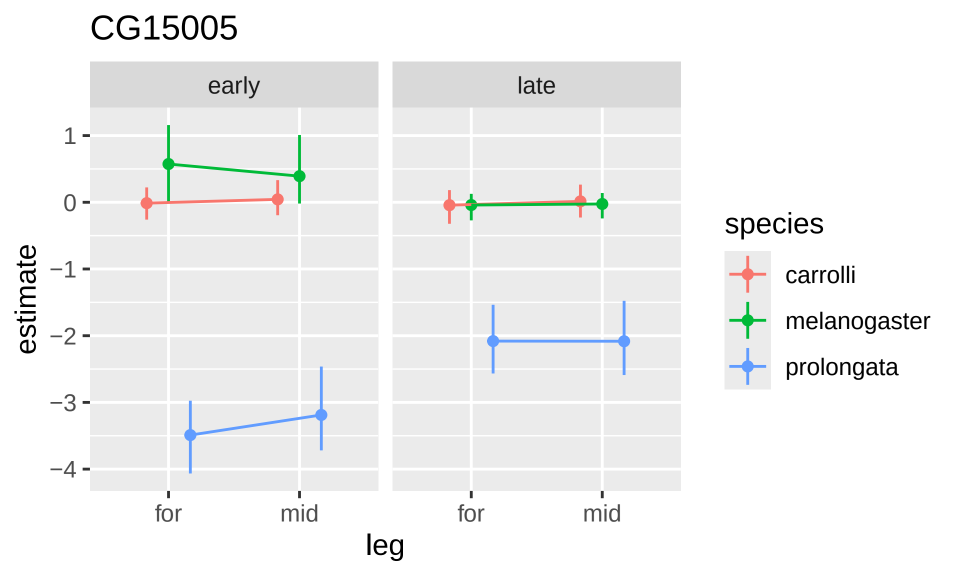

CG15005
early
late
1
0
−1
−2
−3
−4
estimate
for
mid
for
mid
leg
species
carrolli
melanogaster
prolongata

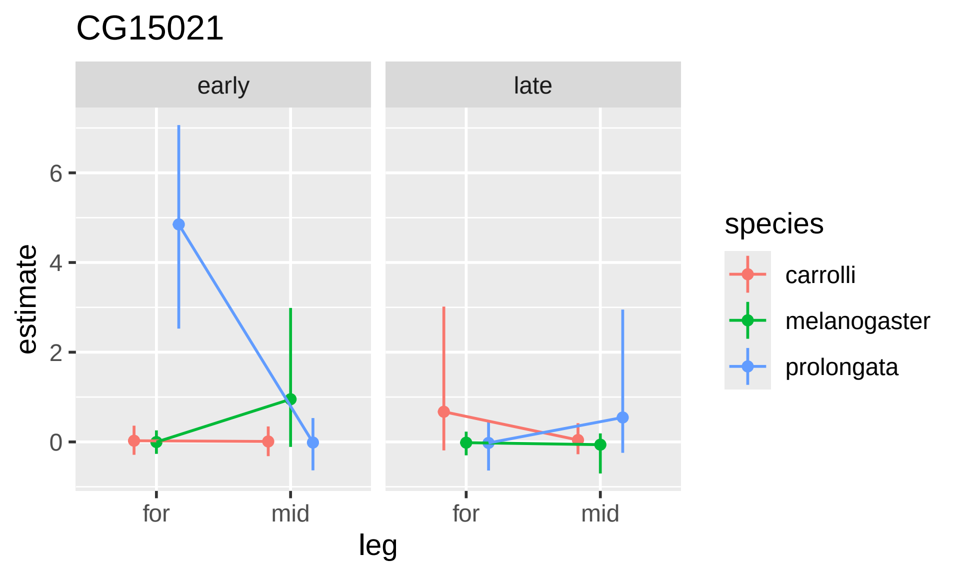

CG15021
early
late
6
4
2
0
estimate
for
mid
for
mid
leg
species
carrolli
melanogaster
prolongata

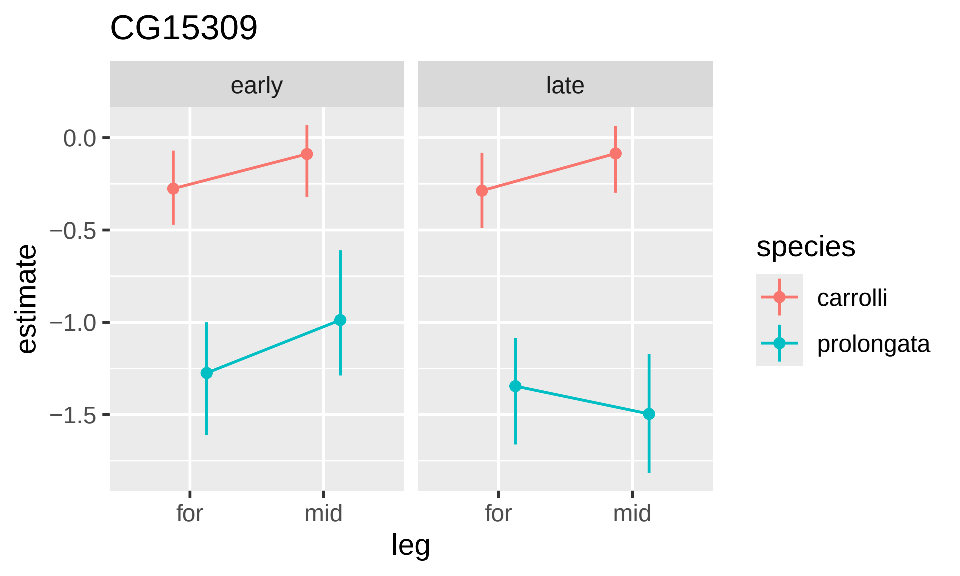
CG15309
early
late
0.0
−0.5
−1.0
−1.5
estimate
for
mid
for
mid
leg
species
carrolli
prolongata

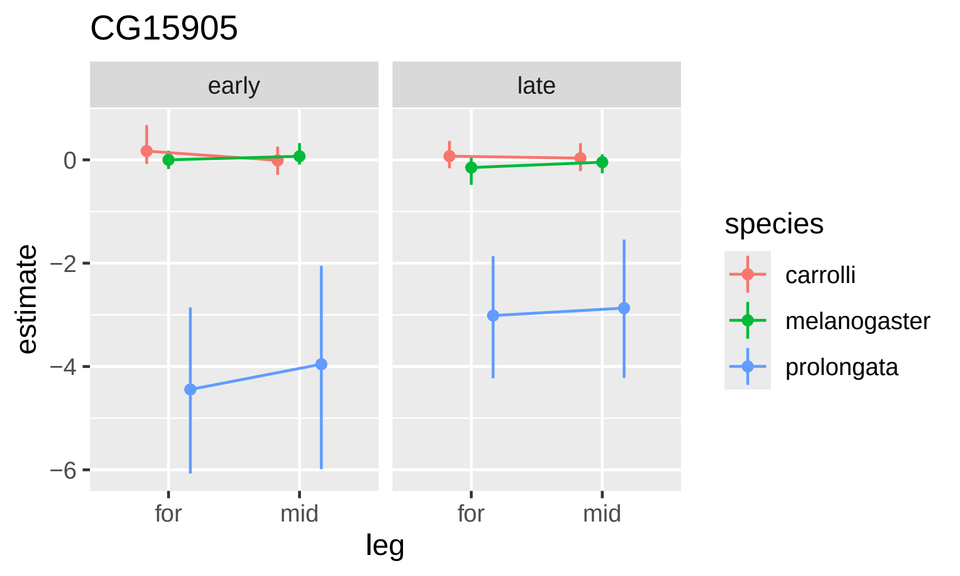

CG15905
early
late
0
−2
−4
−6
estimate
for
mid
for
mid
leg
species
carrolli
melanogaster
prolongata

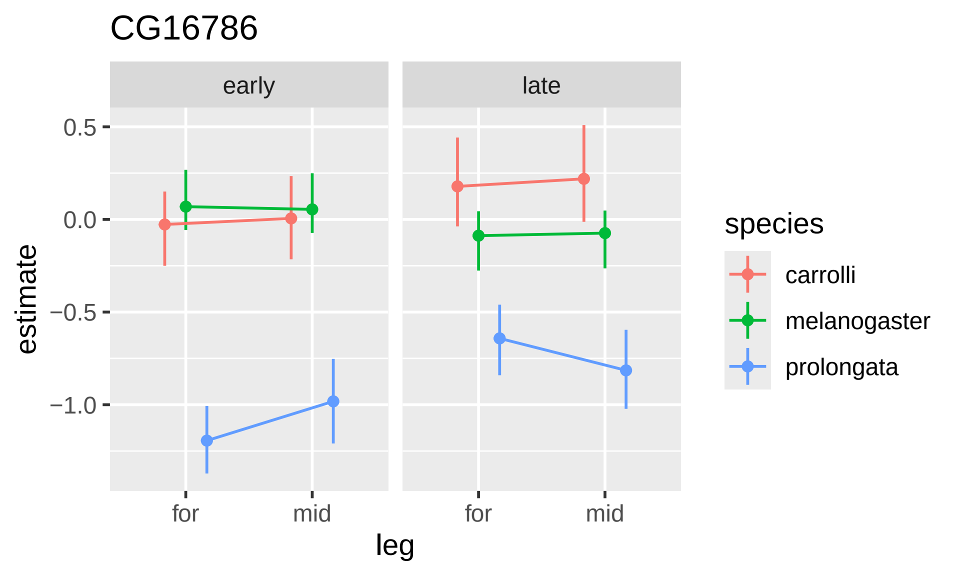

CG16786
early
late
0.5
0.0
−0.5
−1.0
estimate
for
mid
for
mid
leg
species
carrolli
melanogaster
prolongata

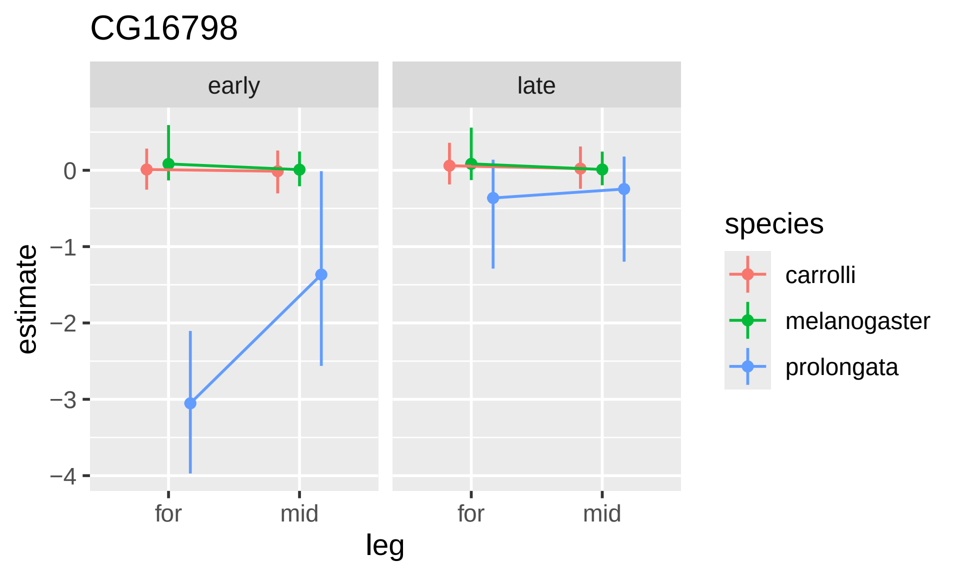

CG16798
early
late
estimate
0
−1
−2
−3
−4
for
mid
for
mid
leg
species
carrolli
melanogaster
prolongata

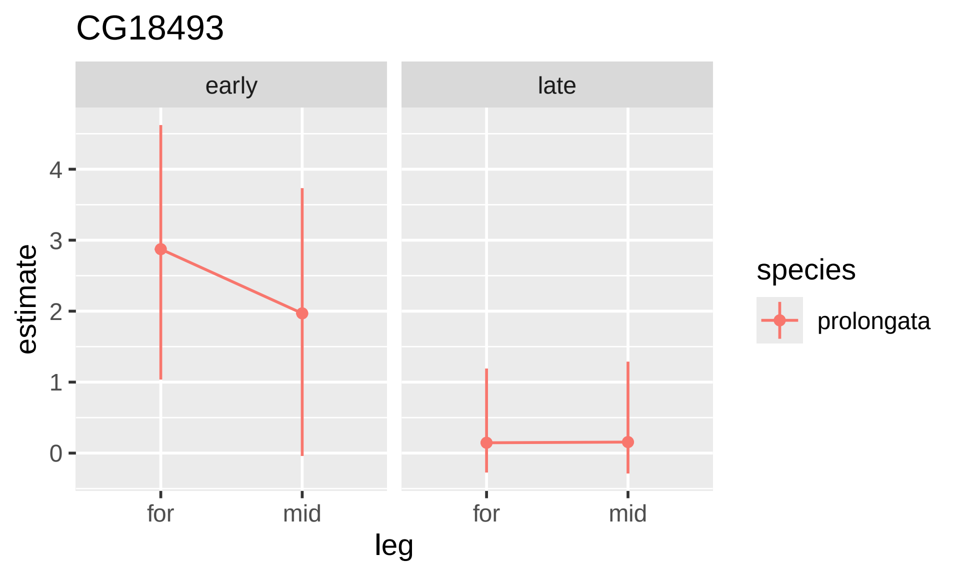
CG18493
early
late
estimate
0
1
2
3
4
for
mid
for
mid
leg
species
prolongata

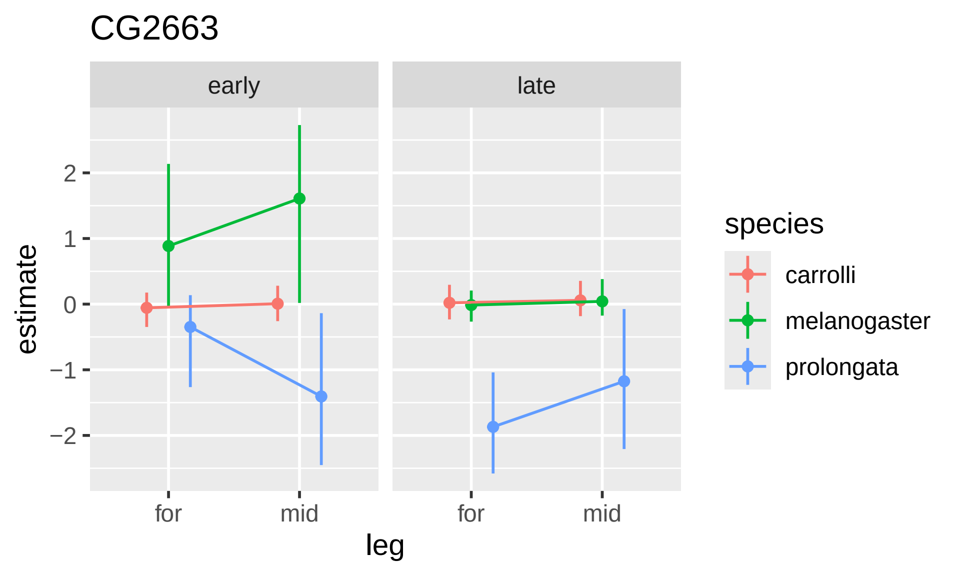
CG2663
early
late
2
1
0
−1
−2
estimate
for
mid
for
mid
leg
species
carrolli
melanogaster
prolongata

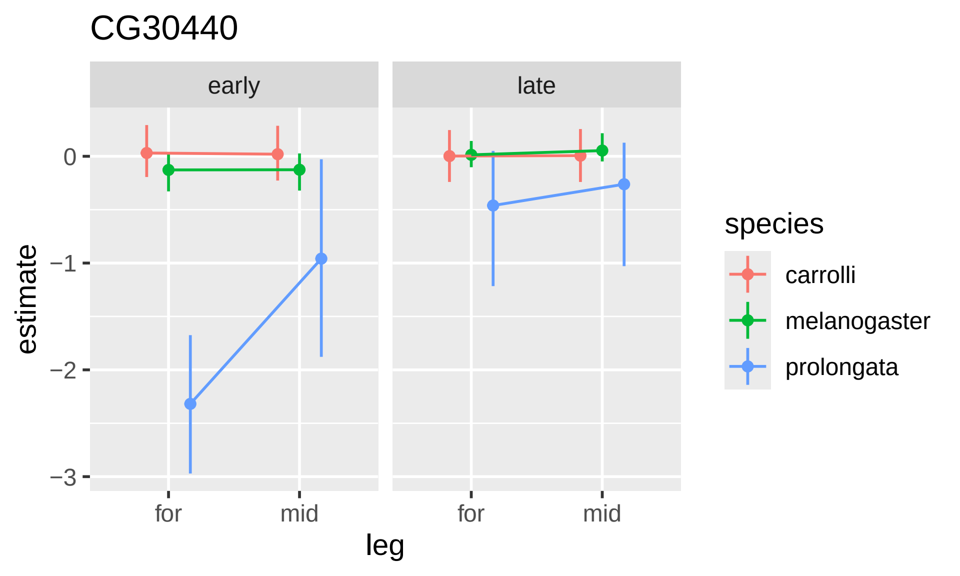
CG30440
early
late
estimate
0
−1
−2
−3
for
mid
for
mid
leg
species
carrolli
melanogaster
prolongata

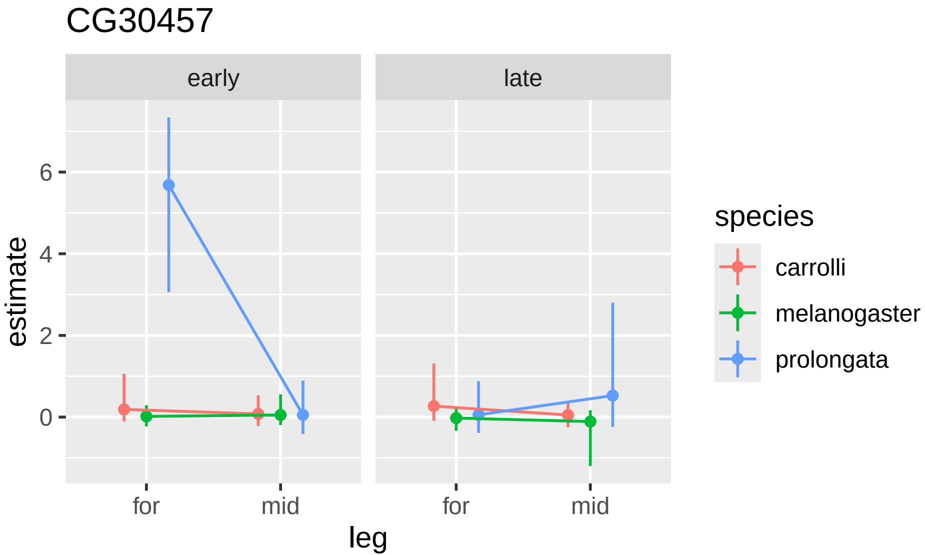

CG30457
early
late
estimate
6
4
2
0
for
mid
for
mid
leg
species
carrolli
melanogaster
prolongata

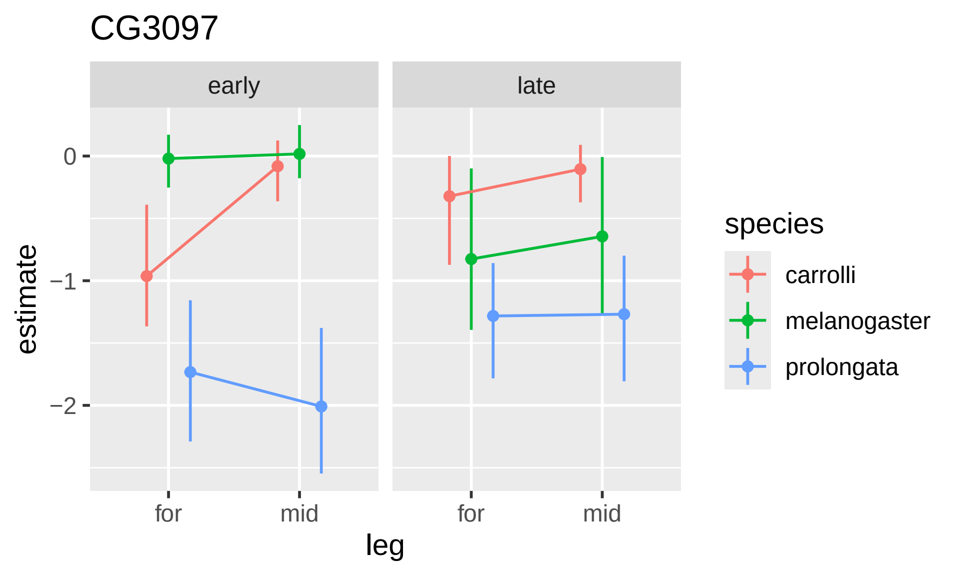

CG3097
early
late
estimate
0
−1
−2
for
mid
for
mid
leg
species
carrolli
melanogaster
prolongata

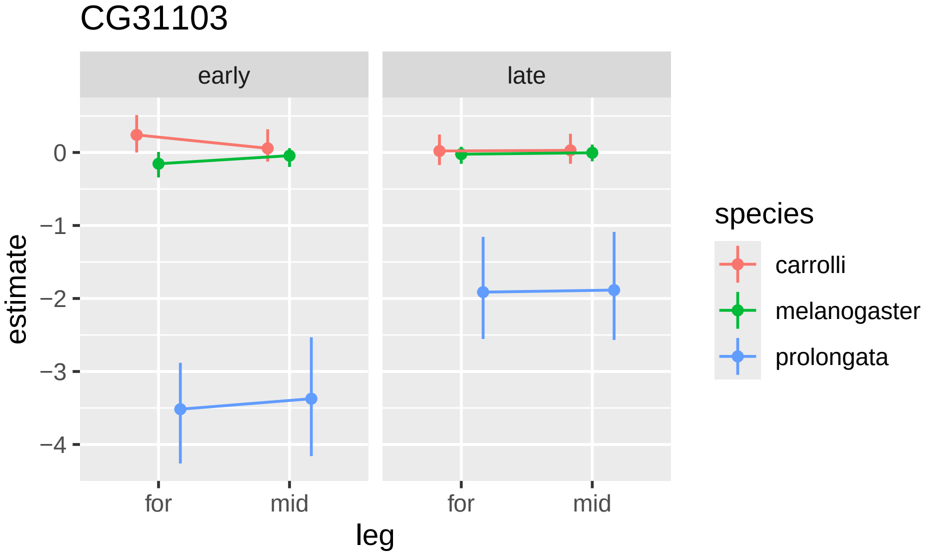
CG31103
early
late
0
−1
−2
−3
−4
estimate
for
mid
for
mid
leg
species
carrolli
melanogaster
prolongata

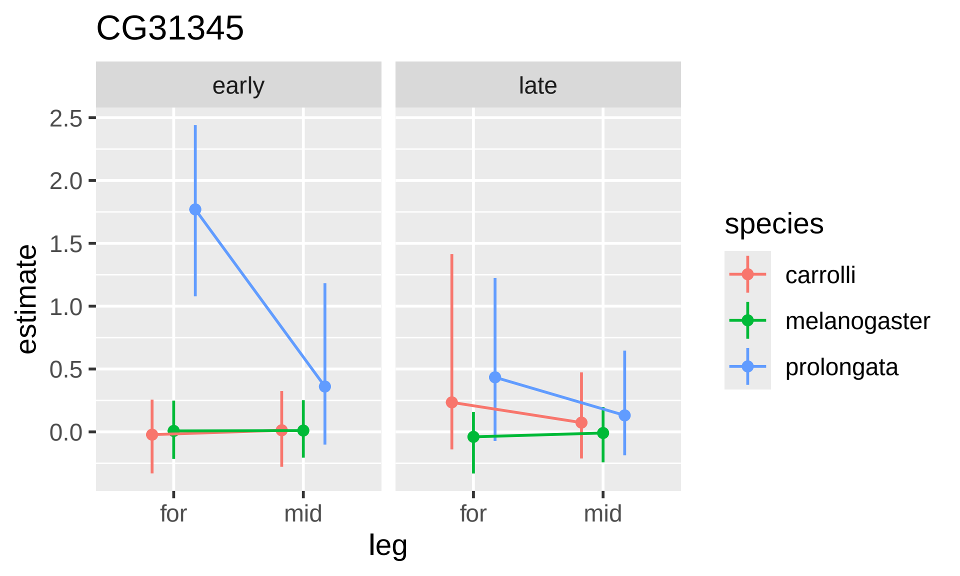

CG31345
early
late
estimate
2.5
2.0
1.5
1.0
0.5
0.0
for
mid
for
mid
leg
species
carrolli
melanogaster
prolongata

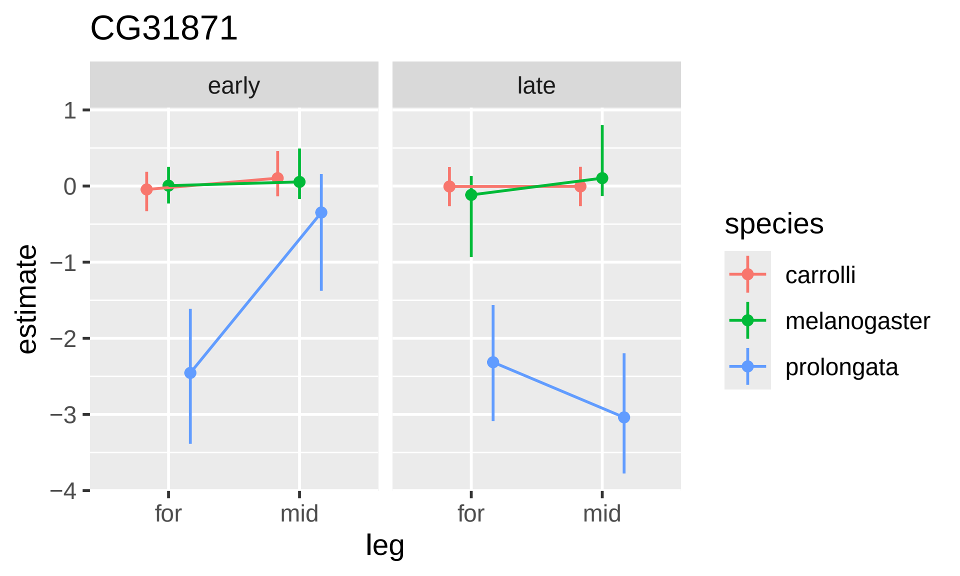
CG31871
early
late
1
0
−1
−2
−3
−4
estimate
for
mid
for
mid
leg
species
carrolli
melanogaster
prolongata

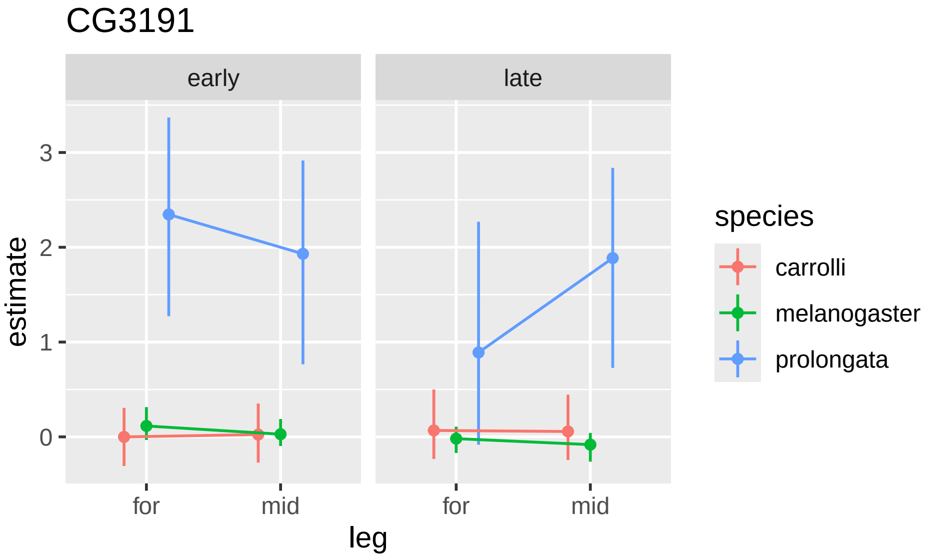
CG3191
early
late
3
2
1
0
estimate
for
mid
for
mid
leg
species
carrolli
melanogaster
prolongata

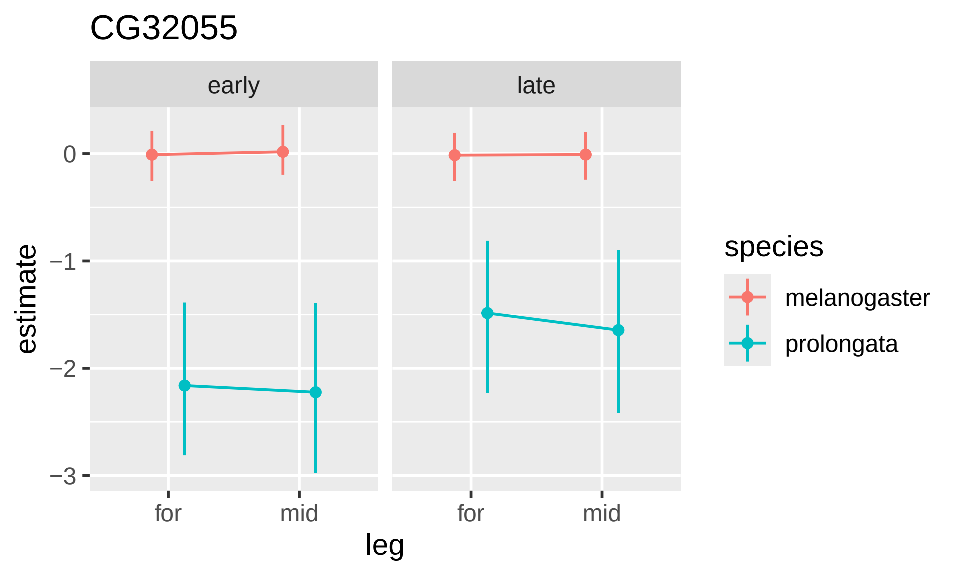

CG32055
early
late
0
−1
−2
−3
estimate
for
mid
for
mid
leg
species
melanogaster
prolongata

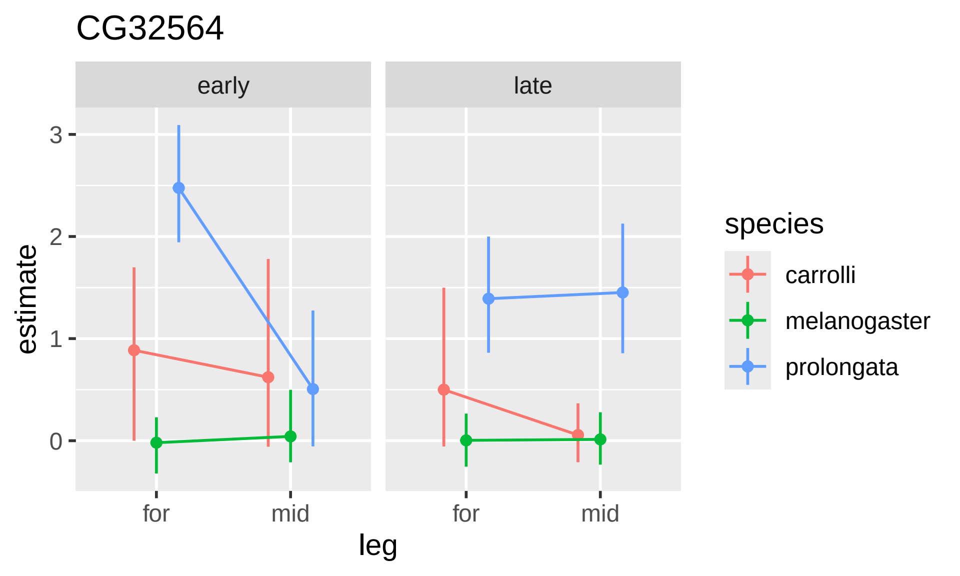

CG32564
early
late
estimate
3
2
1
0
for
mid
for
mid
leg
species
carrolli
melanogaster
prolongata

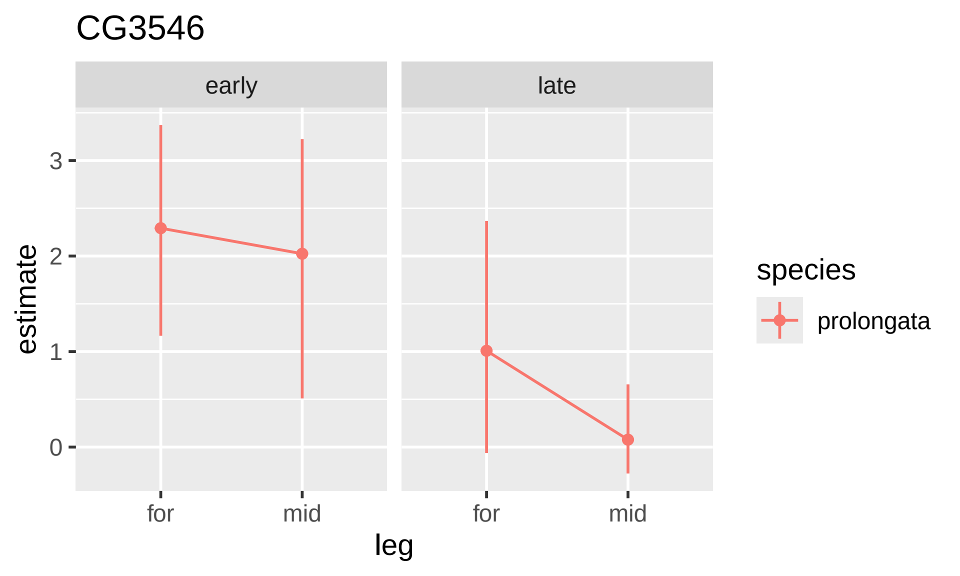

CG3546
early
late
estimate
3
2
1
0
for
mid
for
mid
leg
species
prolongata

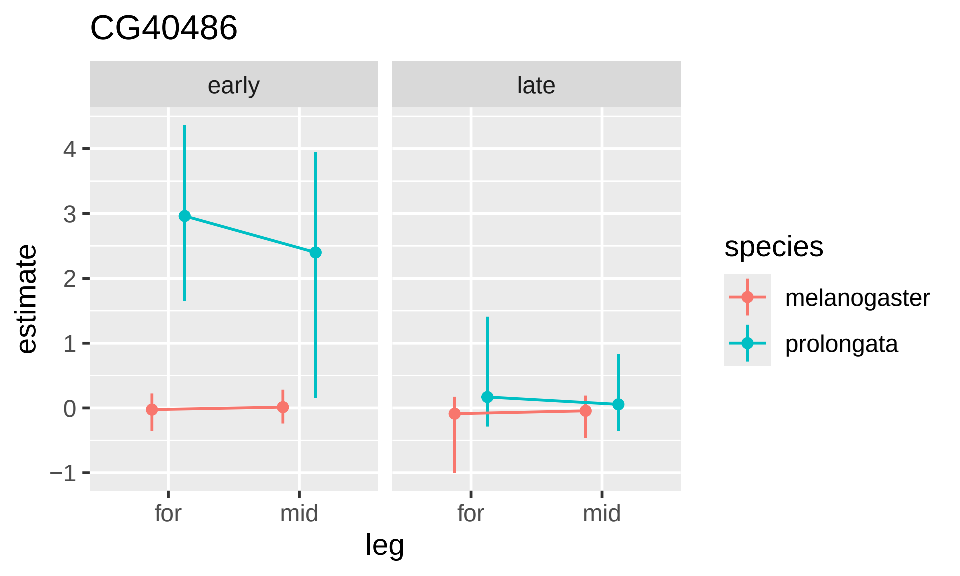
CG40486
early
late
estimate
4
3
2
1
0
−1
for
mid
for
mid
leg
species
melanogaster
prolongata

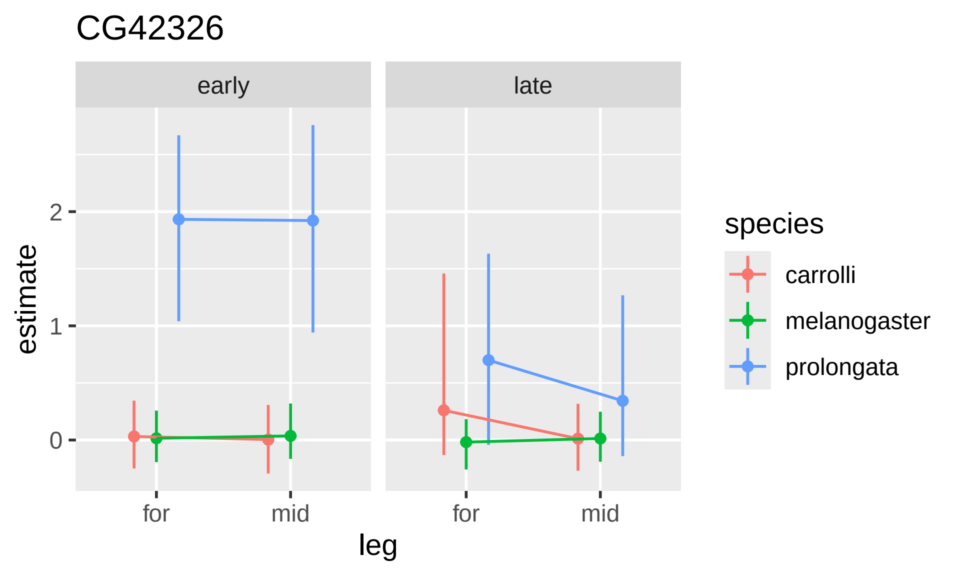

CG42326
early
late
estimate
2
1
0
for
mid
for
mid
leg
species
carrolli
melanogaster
prolongata

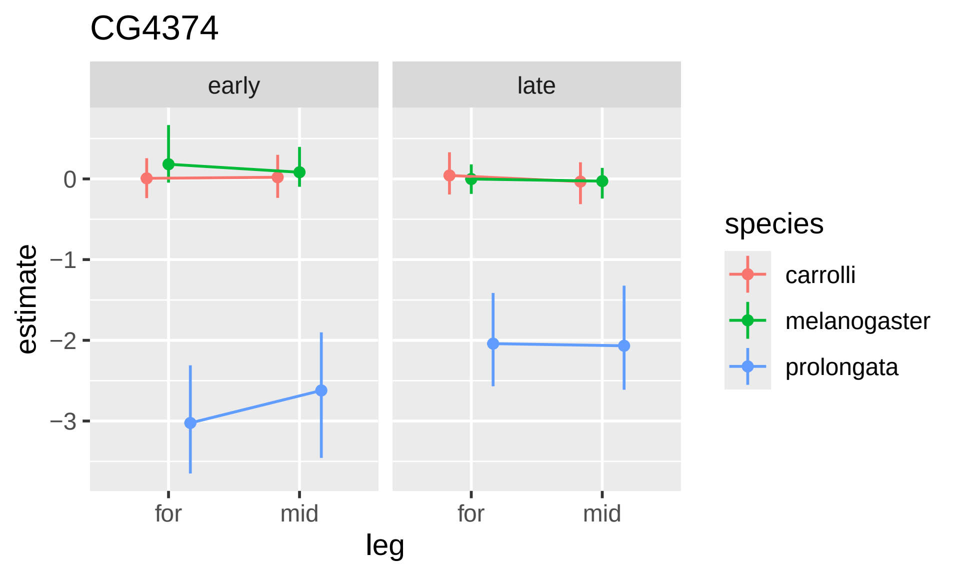
CG4374
early
late
estimate
0
−1
−2
−3
for
mid
for
mid
leg
species
carrolli
melanogaster
prolongata

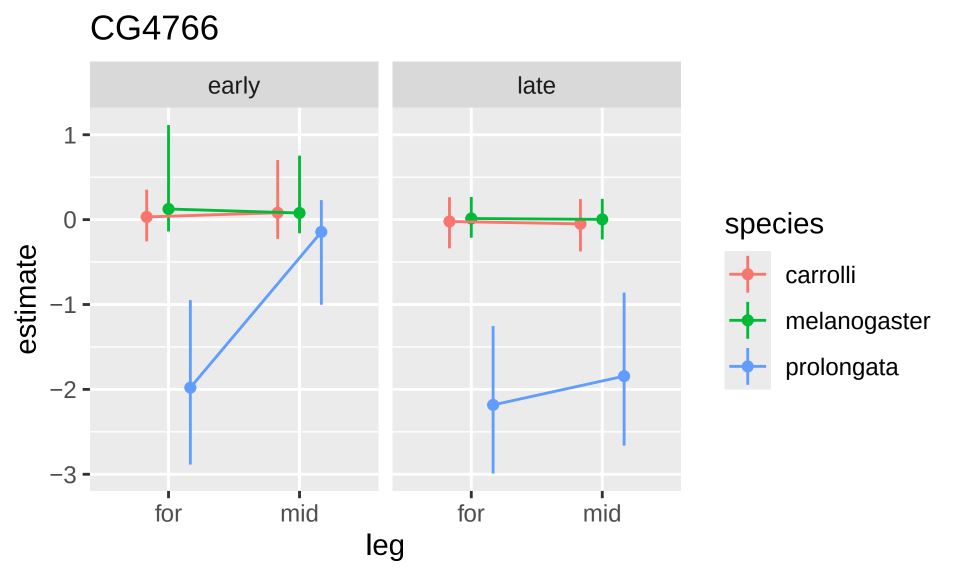
CG4766
early
late
1
0
−1
−2
−3
estimate
for
mid
for
mid
leg
species
carrolli
melanogaster
prolongata

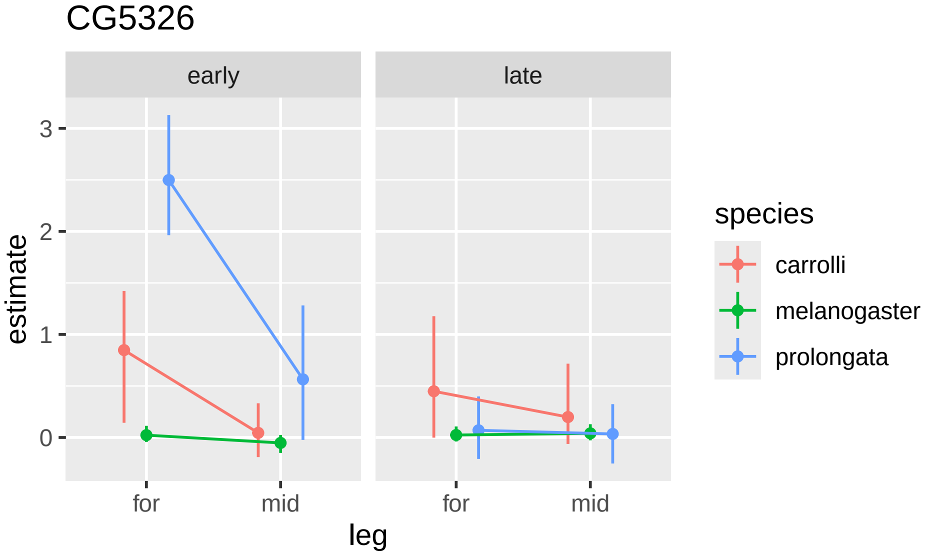
CG5326
early
late
3
2
1
0
estimate
for
mid
for
mid
leg
species
carrolli
melanogaster
prolongata

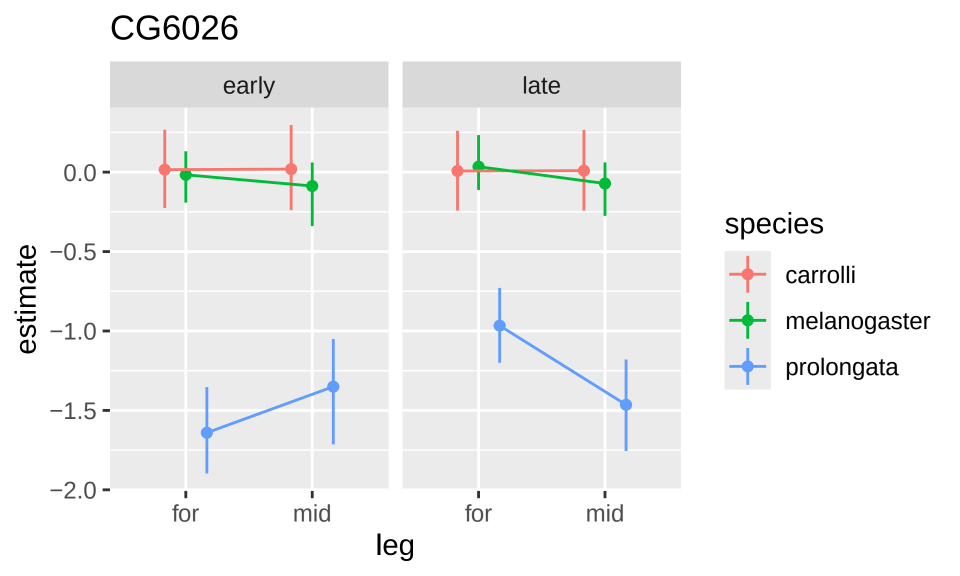

CG6026
early
late
0.0
−0.5
−1.0
−1.5
−2.0
estimate
for
mid
for
mid
leg
species
carrolli
melanogaster
prolongata

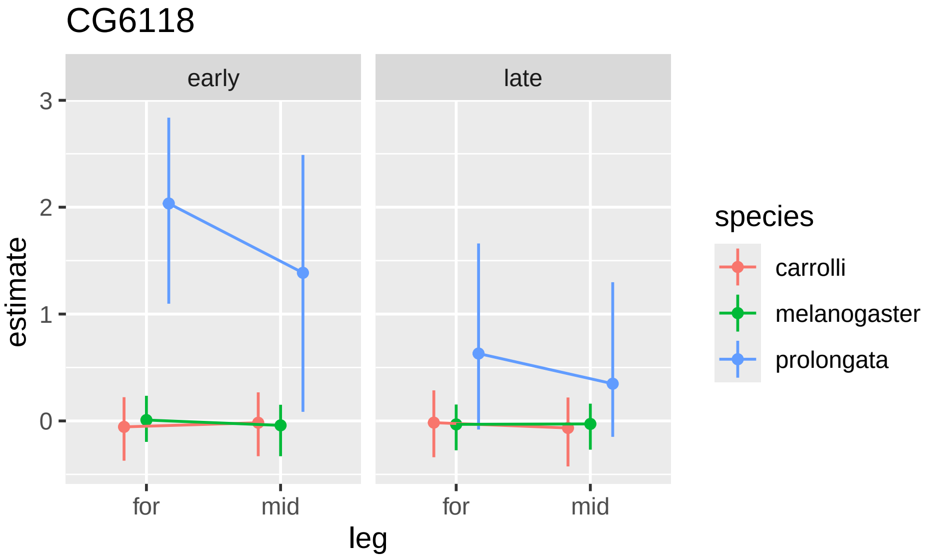

CG6118
early
late
estimate
3
2
1
0
for
mid
for
mid
leg
species
carrolli
melanogaster
prolongata

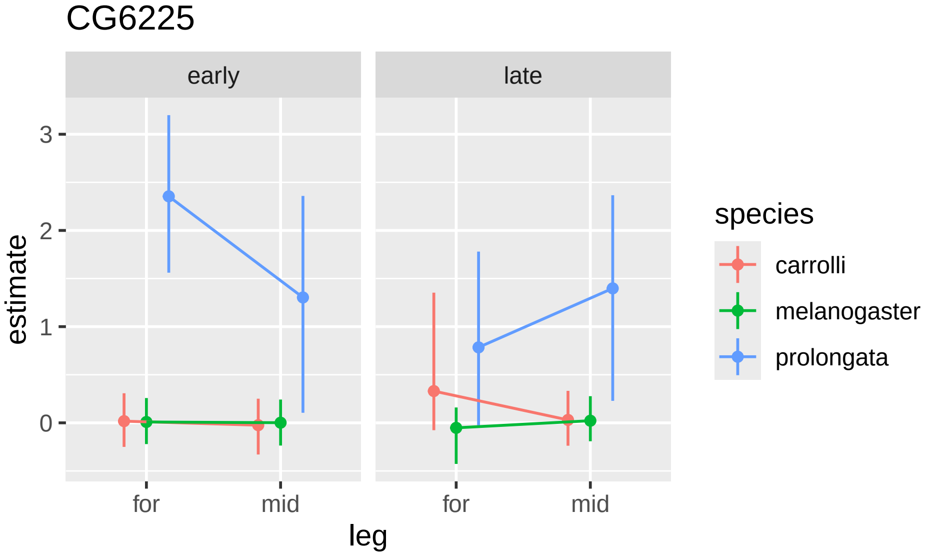
CG6225
early
late
3
2
1
0
estimate
for
mid
for
mid
leg
species
carrolli
melanogaster
prolongata

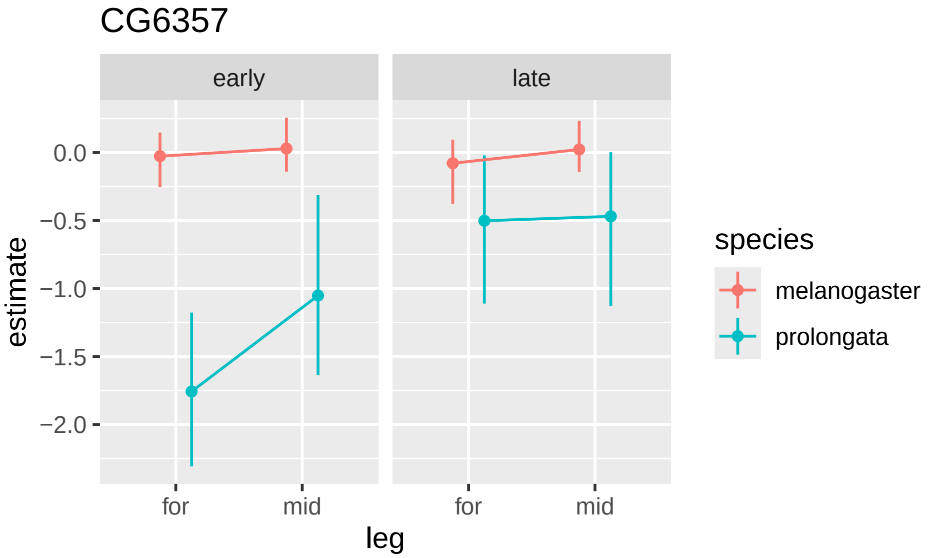

CG6357
early
late
0.0
−0.5
−1.0
−1.5
−2.0
estimate
for
mid
for
mid
leg
species
melanogaster
prolongata

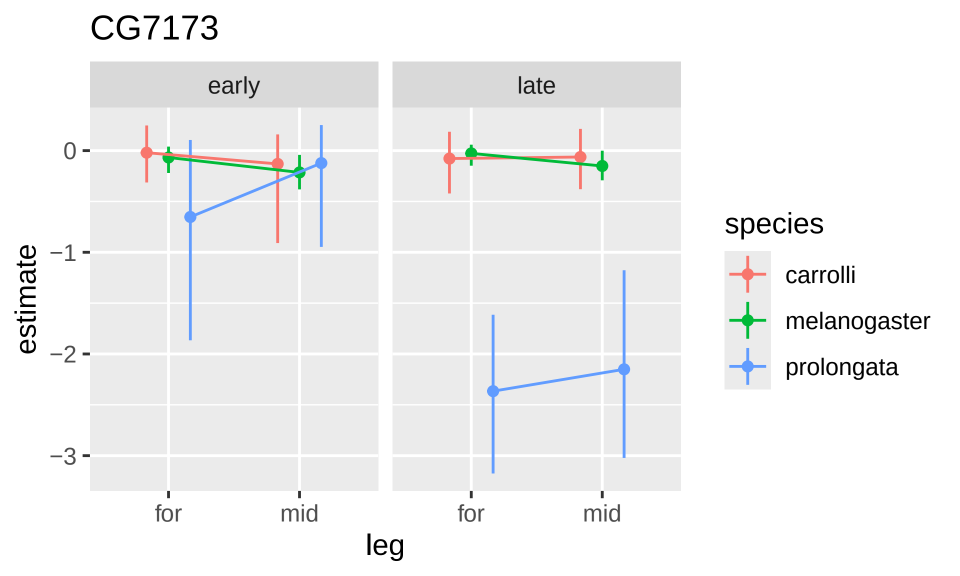

CG7173
early
late
estimate
0
−1
−2
−3
for
mid
for
mid
leg
species
carrolli
melanogaster
prolongata

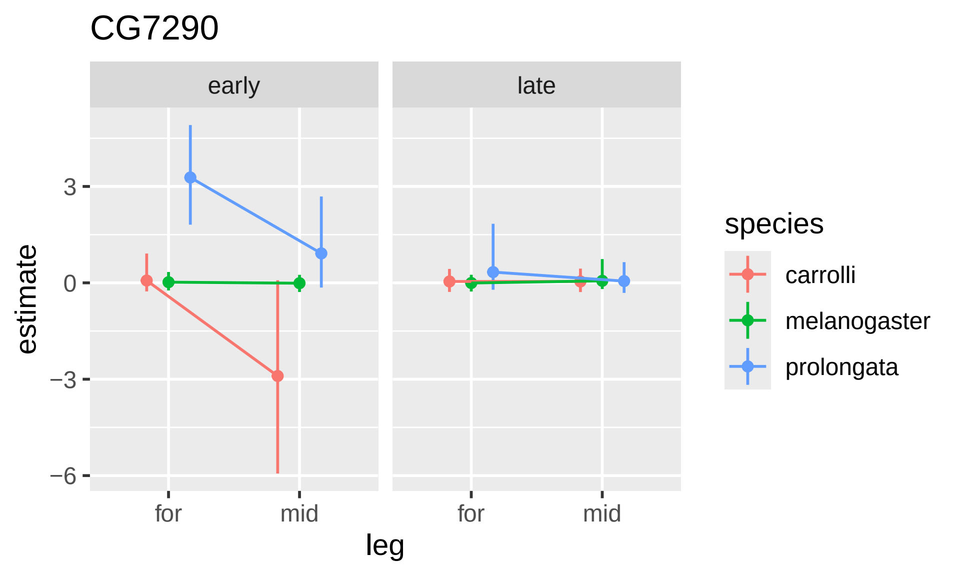

CG7290
early
late
estimate
3
0
−3
−6
for
mid
for
mid
leg
species
carrolli
melanogaster
prolongata

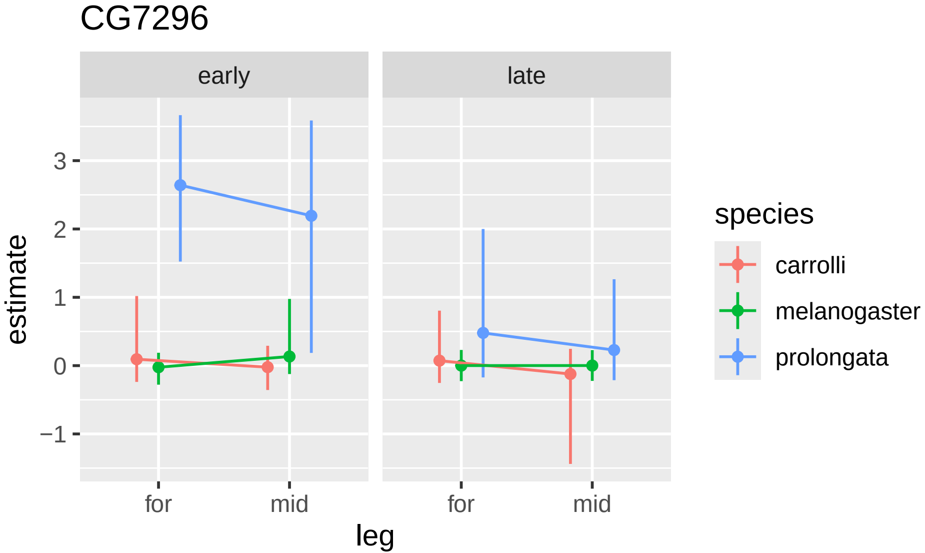
CG7296
early
late
estimate
3
2
1
0
−1
for
mid
for
mid
leg
species
carrolli
melanogaster
prolongata

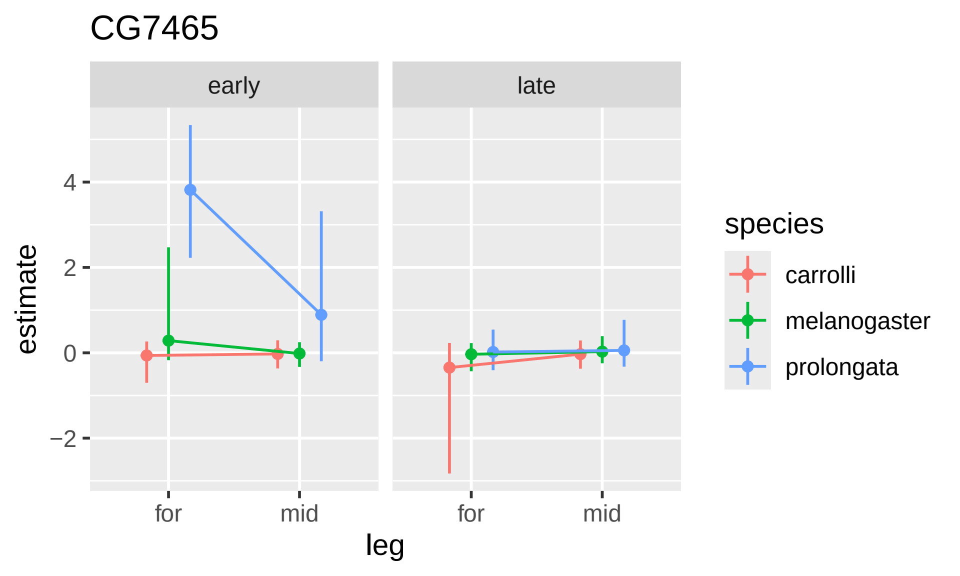
CG7465
early
late
estimate
4
2
0
−2
for
mid
for
mid
leg
species
carrolli
melanogaster
prolongata

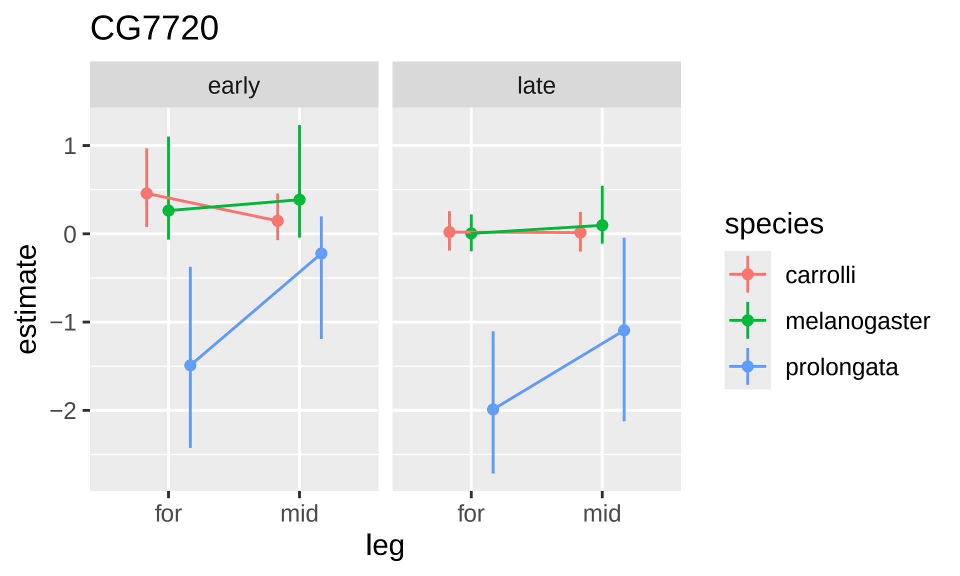
CG7720
early
late
estimate
1
0
−1
−2
for
mid
for
mid
leg
species
carrolli
melanogaster
prolongata

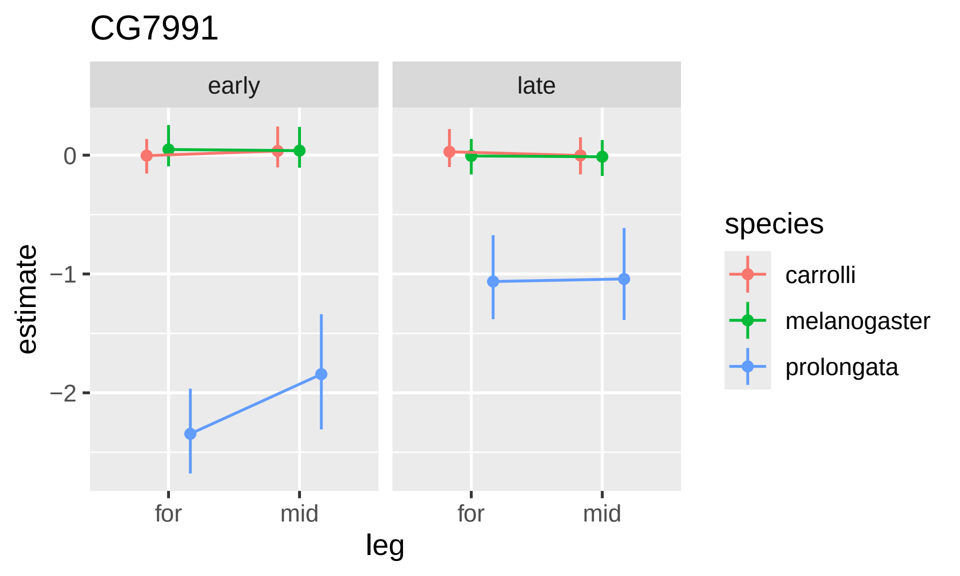

CG7991
early
late
0
−1
−2
estimate
for
mid
for
mid
leg
species
carrolli
melanogaster
prolongata

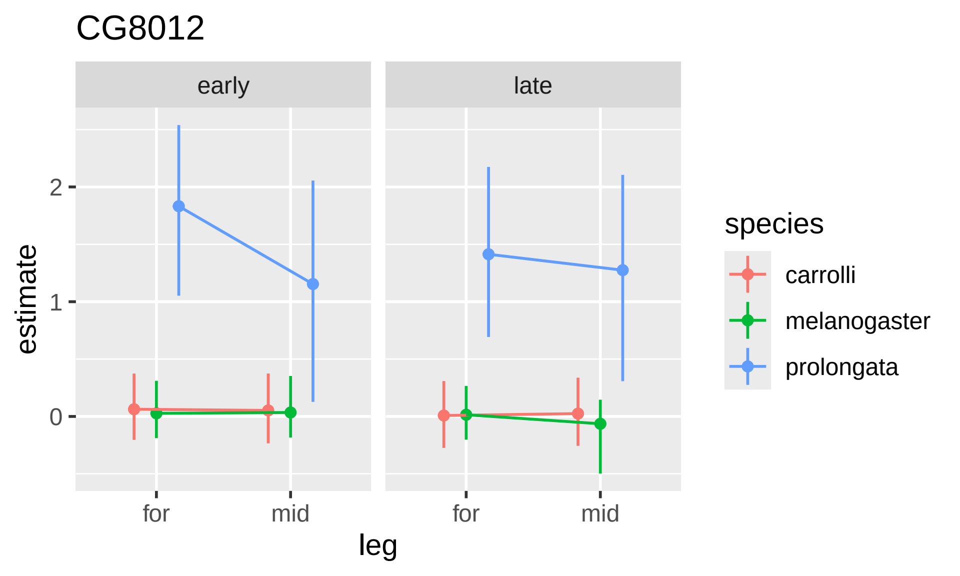

CG8012
early
late
estimate
2
1
0
for
mid
for
mid
leg
species
carrolli
melanogaster
prolongata

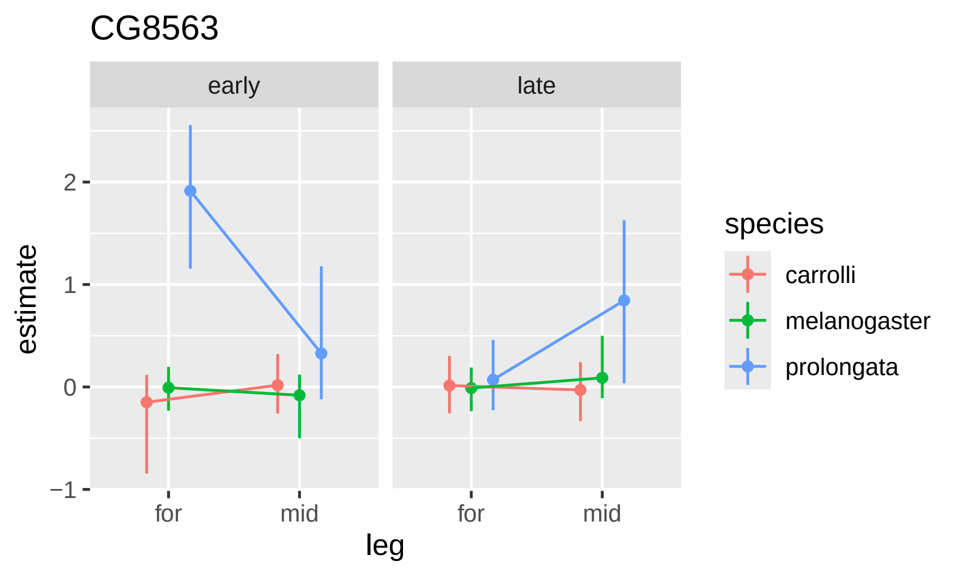

CG8563
early
late
estimate
2
1
0
−1
for
mid
for
mid
leg
species
carrolli
melanogaster
prolongata

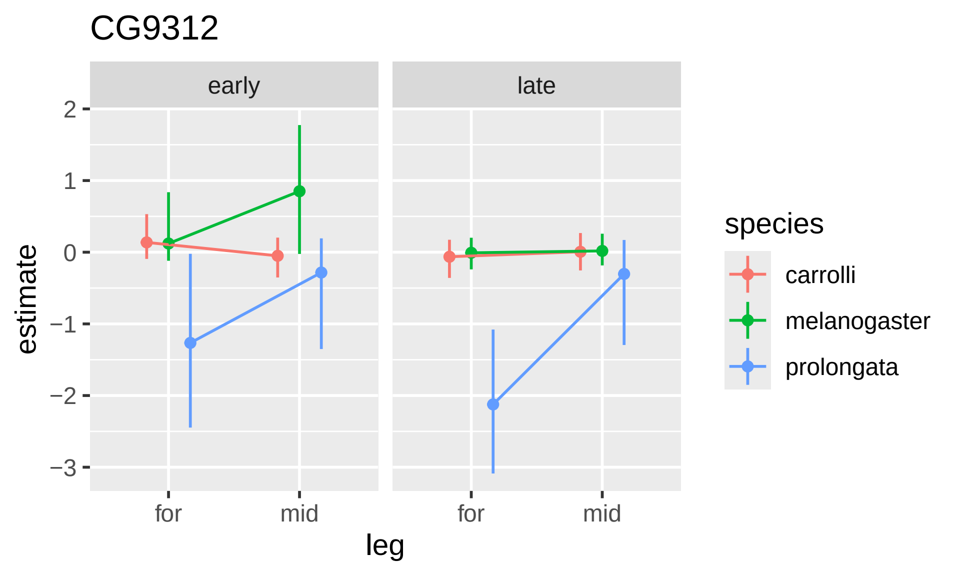

CG9312
early
late
2
1
0
−1
−2
−3
estimate
for
mid
for
mid
leg
species
carrolli
melanogaster
prolongata

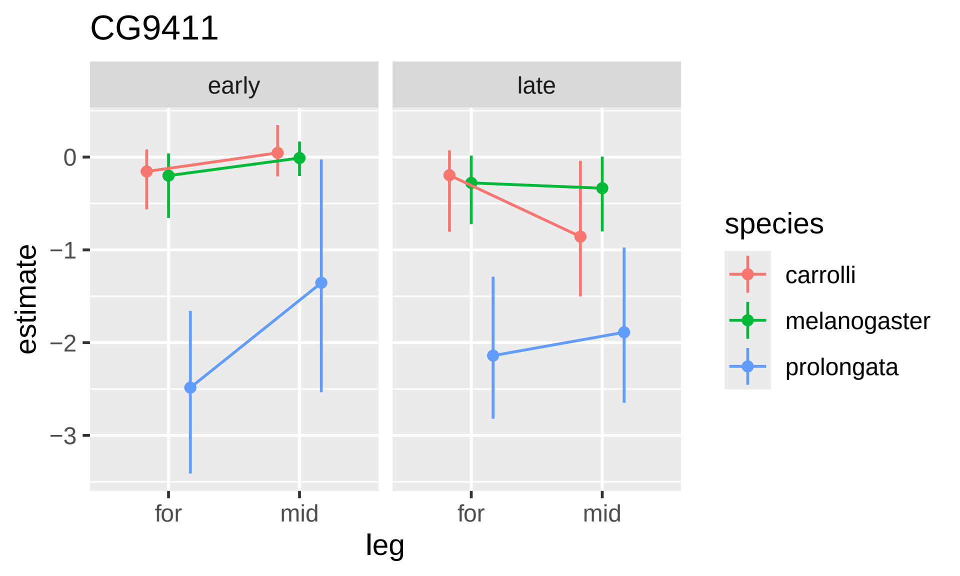

CG9411
early
late
estimate
0
−1
−2
−3
for
mid
for
mid
leg
species
carrolli
melanogaster
prolongata

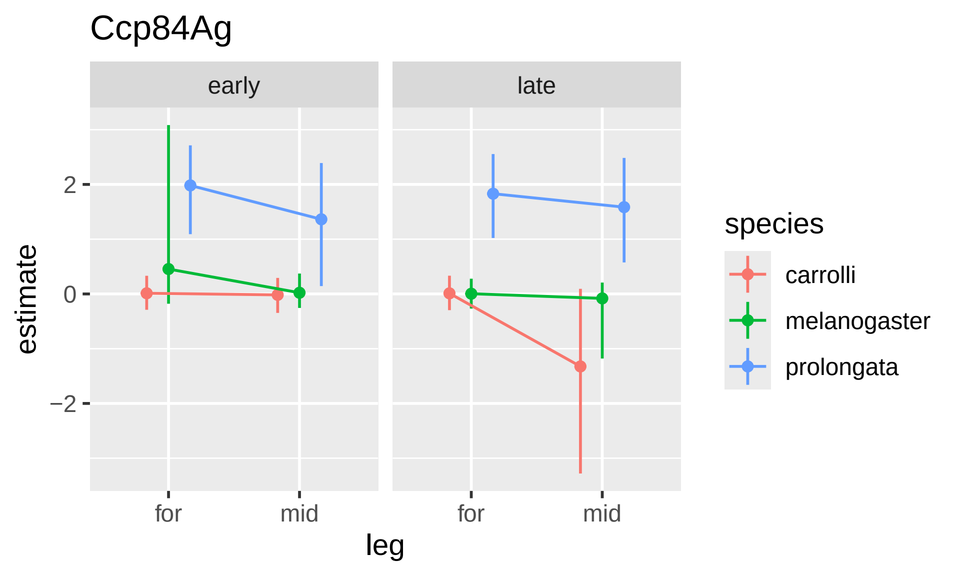

Ccp84Ag
early
late
2
0
−2
estimate
for
mid
for
mid
leg
species
carrolli
melanogaster
prolongata

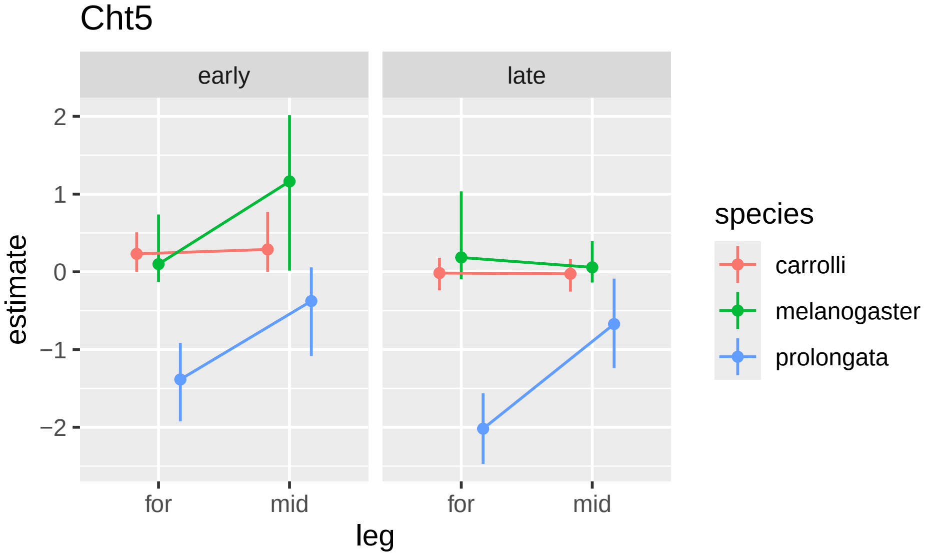

Cht5
early
late
estimate
2
1
0
−1
−2
for
mid
leg
species
carrolli
melanogaster
prolongata

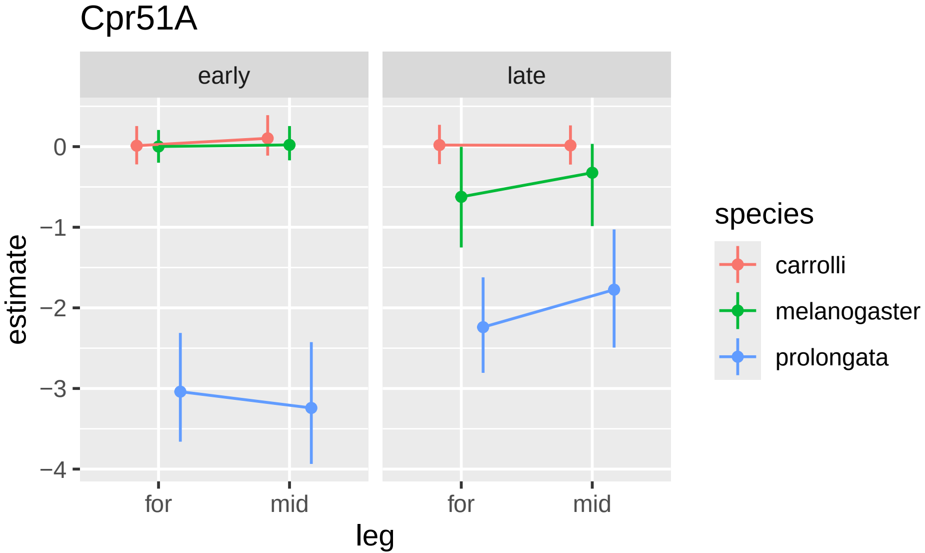

Cpr51A
early
late
0
−1
−2
−3
−4
estimate
for
mid
for
mid
leg
species
carrolli
melanogaster
prolongata

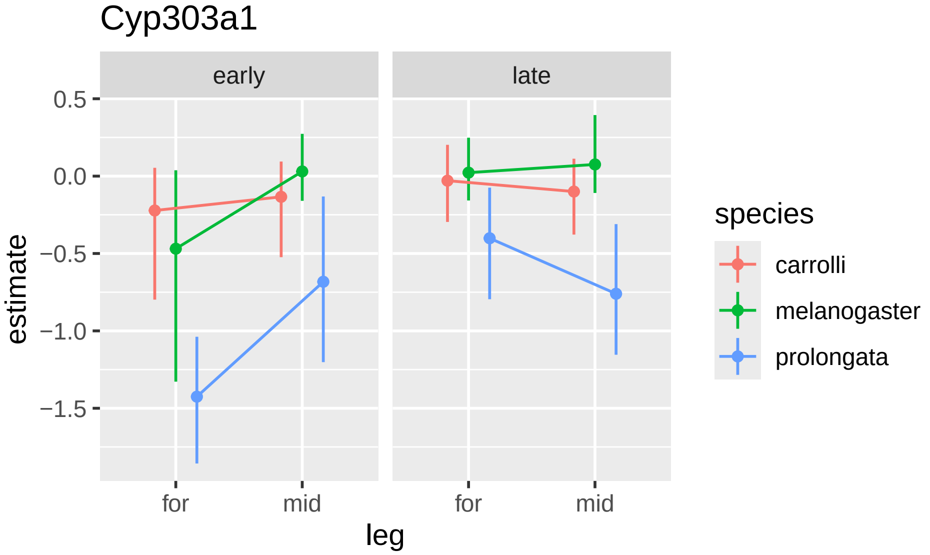
Cyp303a1
early
late
0.5
0.0
−0.5
−1.0
−1.5
estimate
for
mid
for
mid
leg
species
carrolli
melanogaster
prolongata

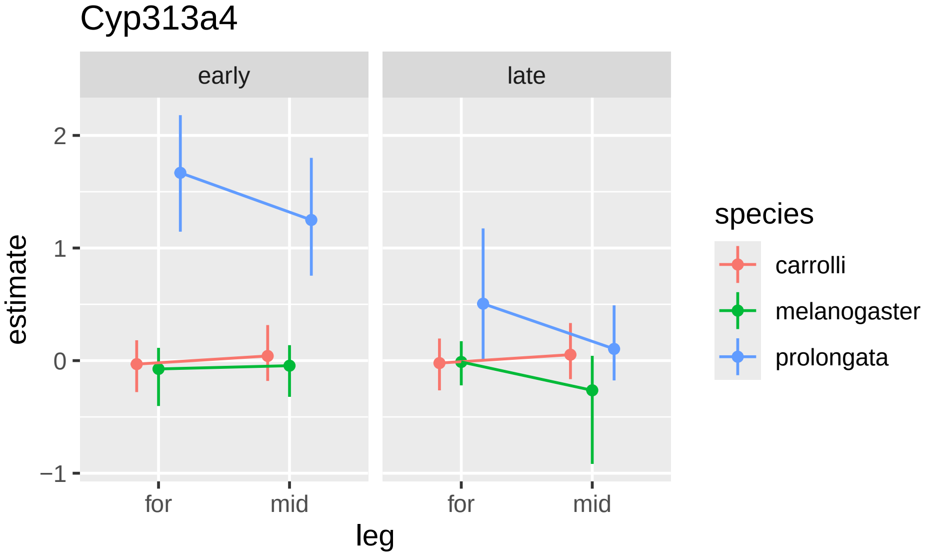

Cyp313a4
early
late
2
1
0
−1
estimate
for
mid
for
mid
leg
species
carrolli
melanogaster
prolongata

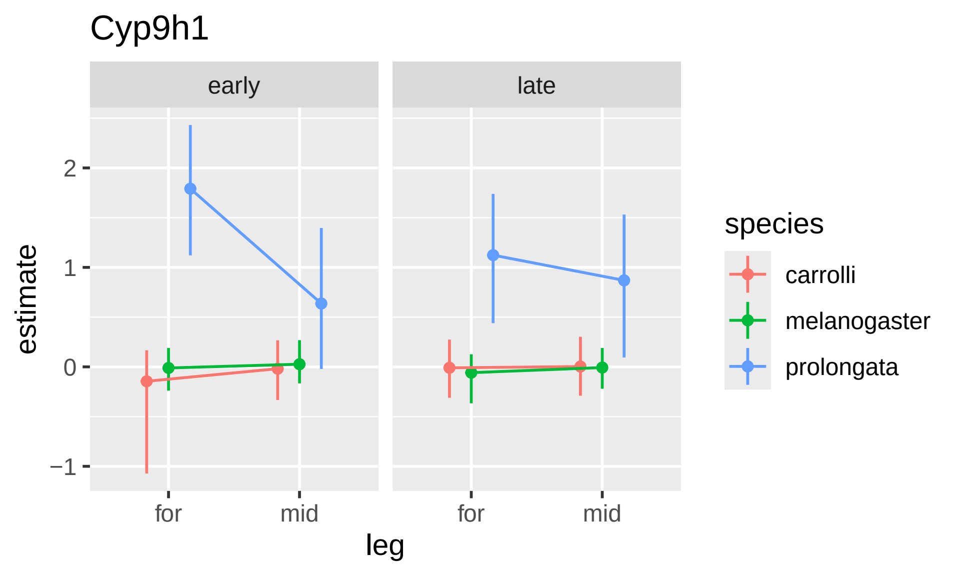

Cyp9h1
early
late
2
1
0
−1
estimate
for
mid
for
mid
leg
species
carrolli
melanogaster
prolongata

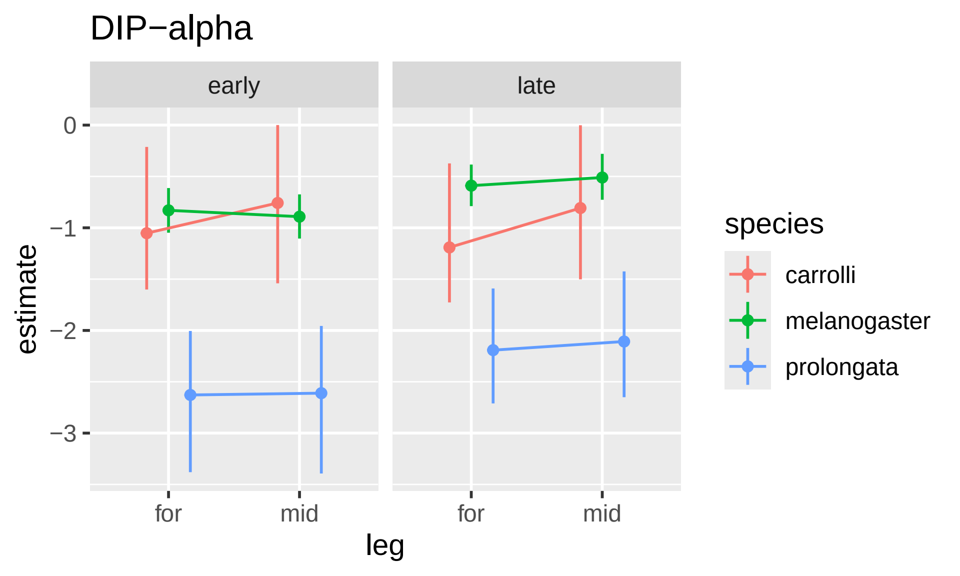

DIP−alpha
early
late
0
−1
−2
−3
estimate
for
mid
for
mid
leg
species
carrolli
melanogaster
prolongata

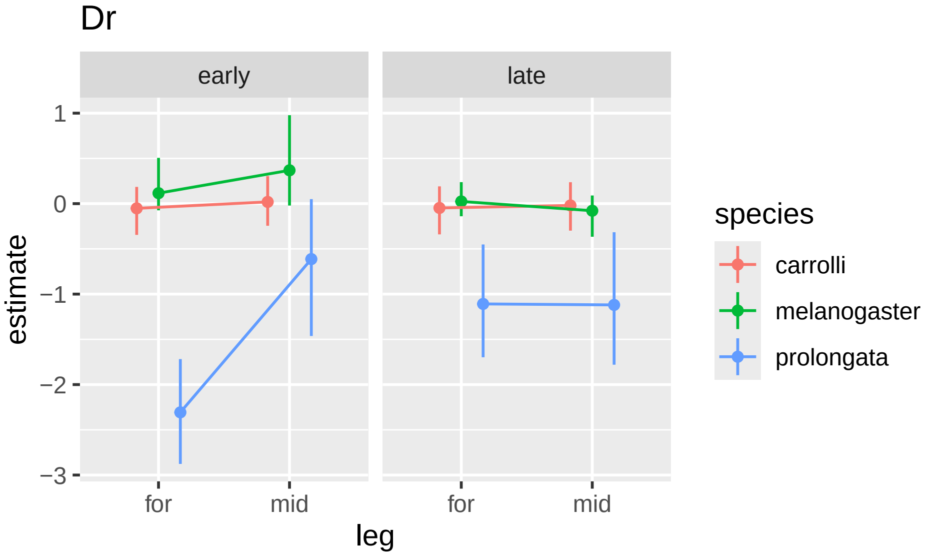

Dr
early
late
1
0
−1
−2
−3
estimate
for
mid
for
mid
leg
species
carrolli
melanogaster
prolongata

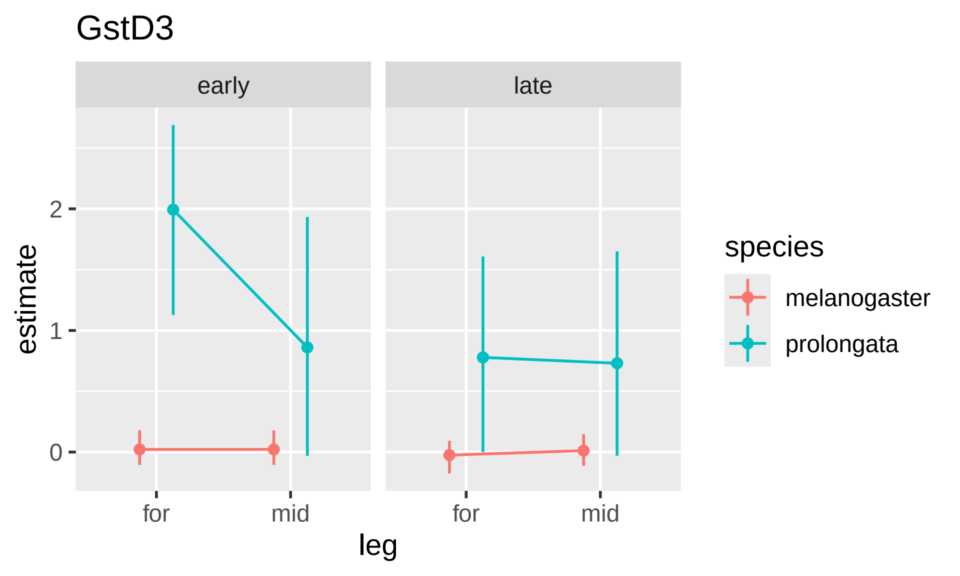
GstD3
early
late
estimate
2
1
0
for
mid
for
mid
leg
species
melanogaster
prolongata

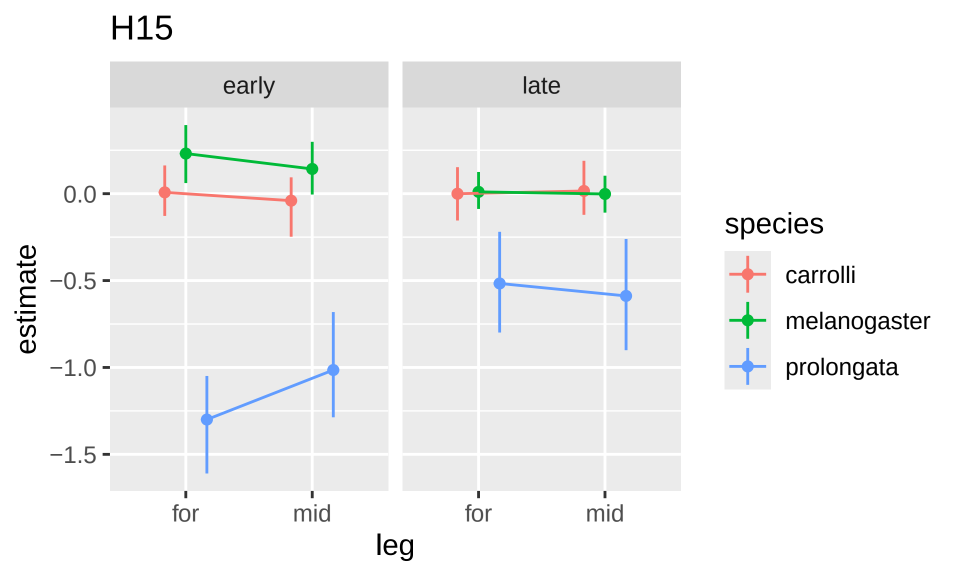

H15
early
late
0.0
−0.5
−1.0
−1.5
estimate
for
mid
for
mid
leg
species
carrolli
melanogaster
prolongata

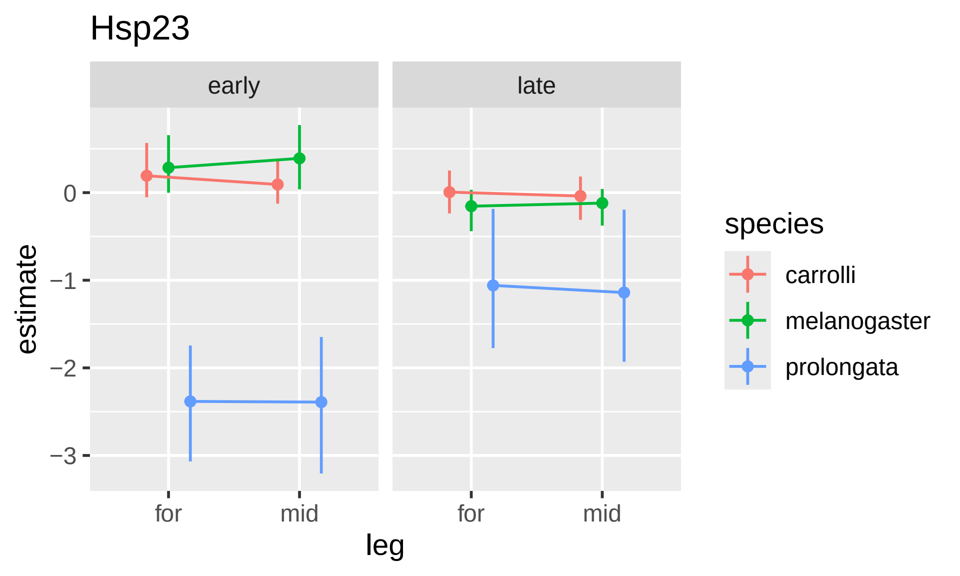

Hsp23
early
late
0
−1
−2
−3
estimate
for
mid
for
mid
leg
species
carrolli
melanogaster
prolongata

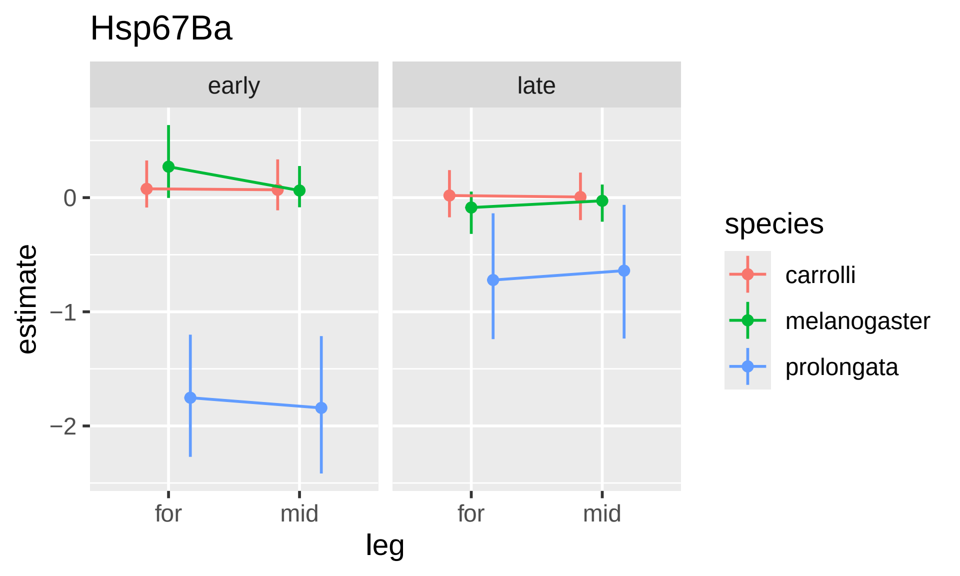
Hsp67Ba
early
late
estimate
0
−1
−2
for
mid
for
mid
leg
species
carrolli
melanogaster
prolongata

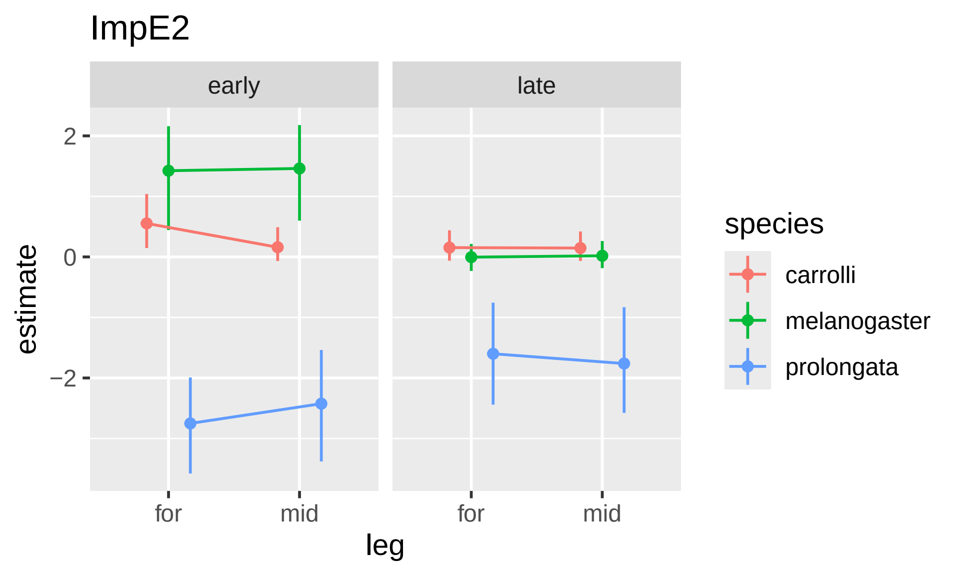

ImpE2
early
late
2
0
−2
estimate
for
mid
for
mid
leg
species
carrolli
melanogaster
prolongata

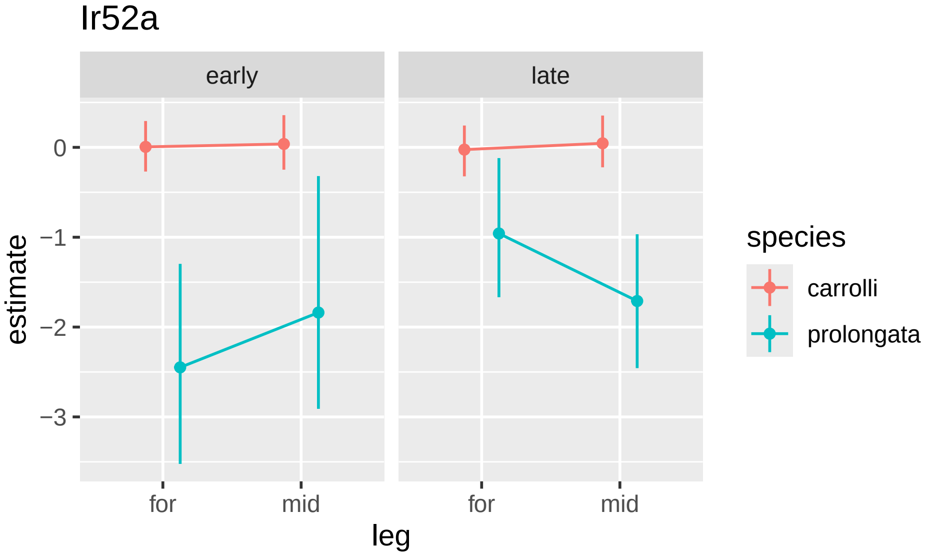
Ir52a
early
late
0
−1
−2
−3
estimate
for
mid
for
mid
leg
species
carrolli
prolongata

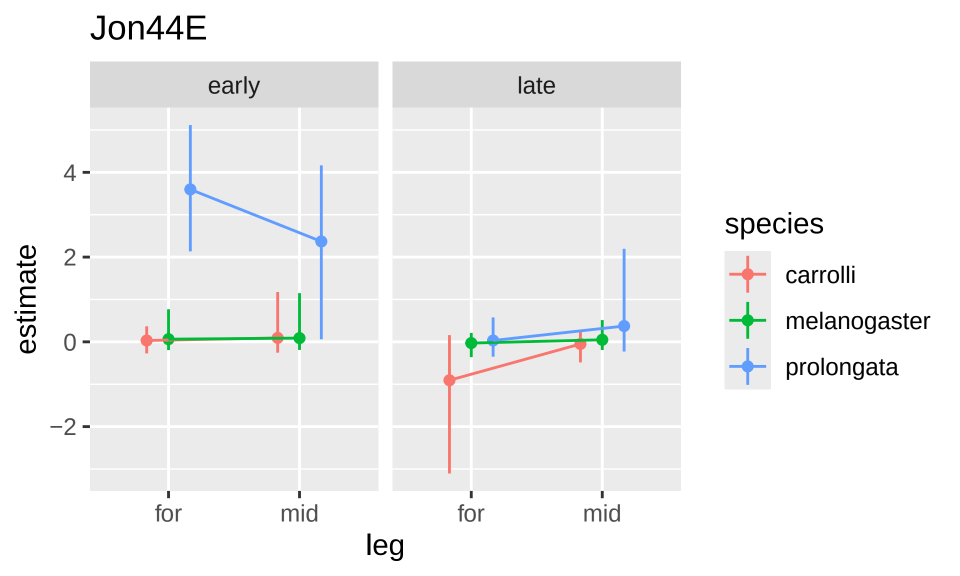

Jon44E
early
late
estimate
4
2
0
−2
for
mid
for
mid
leg
species
carrolli
melanogaster
prolongata

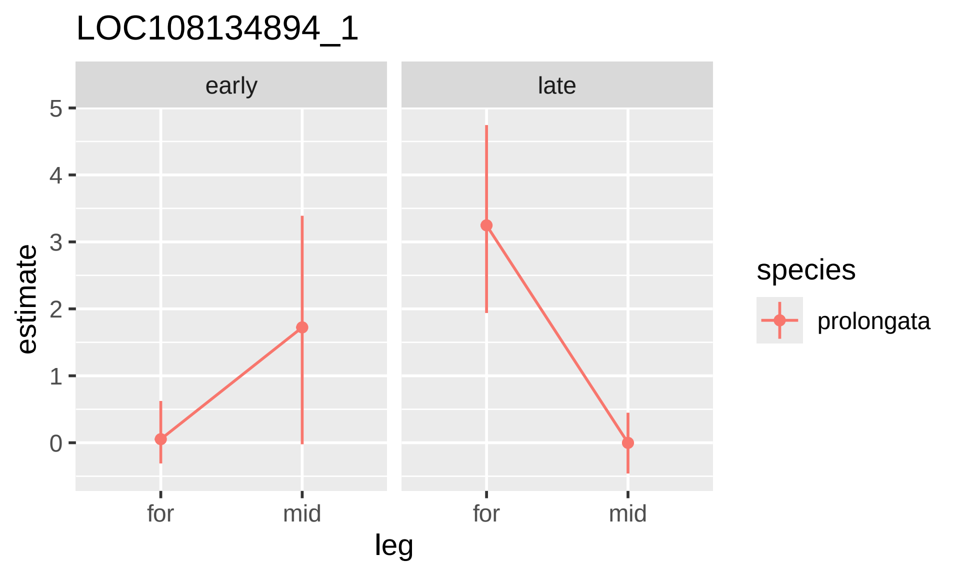

LOC108134894_1
early
late
estimate
5
4
3
2
1
0
for
mid
for
mid
leg
species
prolongata

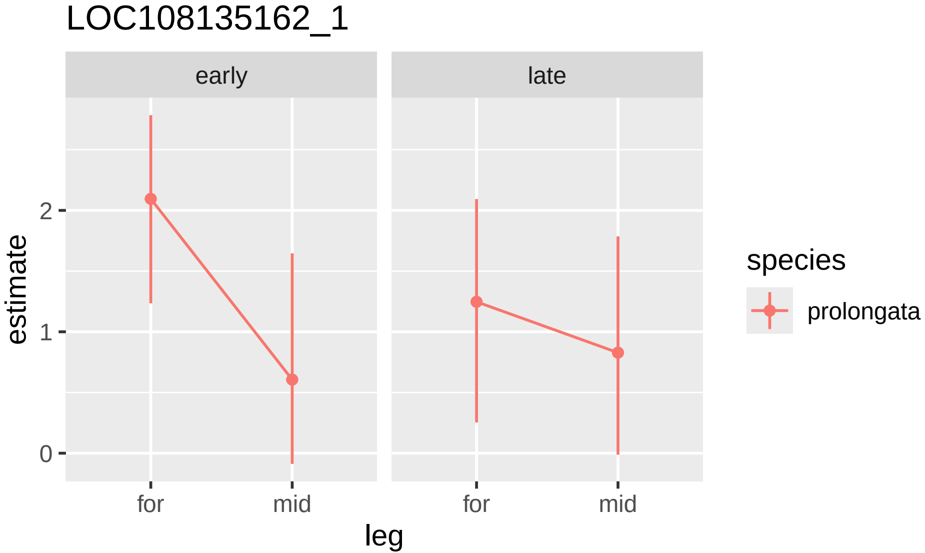

LOC108135162_1
early
late
estimate
2
1
0
for
mid
for
mid
leg
species
prolongata

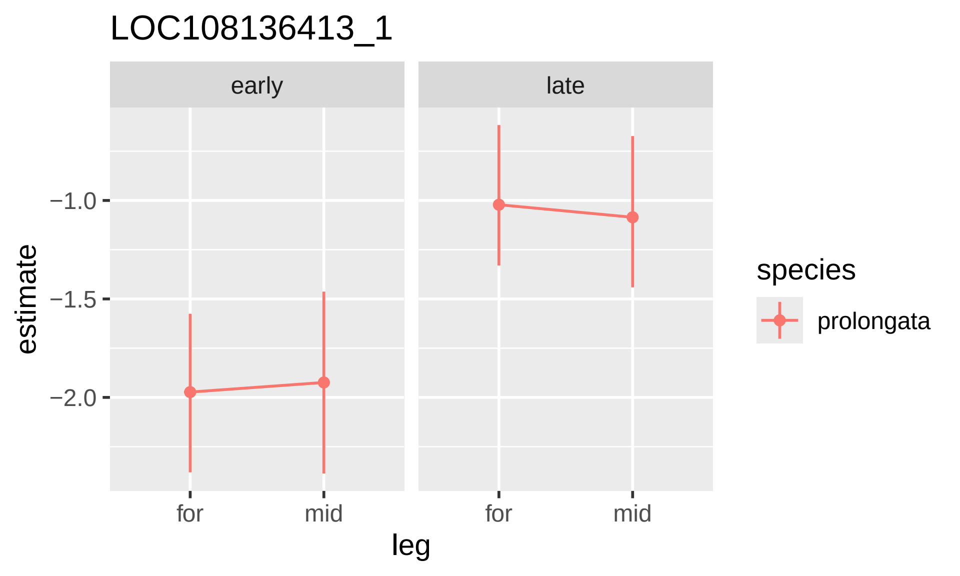

LOC108136413_1
early
late
−1.0
−1.5
−2.0
estimate
for
mid
for
mid
leg
species
prolongata

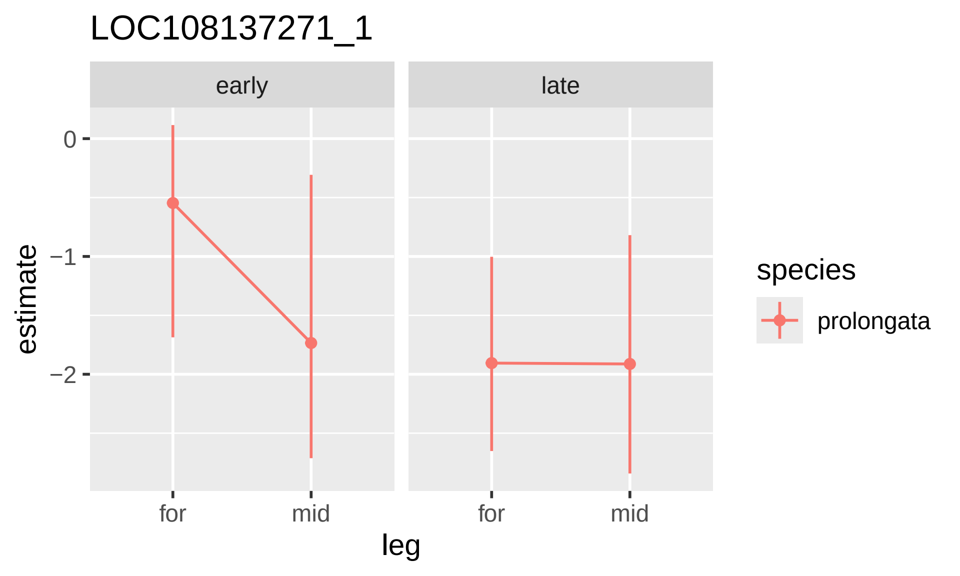

LOC108137271_1
early
late
0
−1
−2
estimate
for
mid
for
mid
leg
species
prolongata

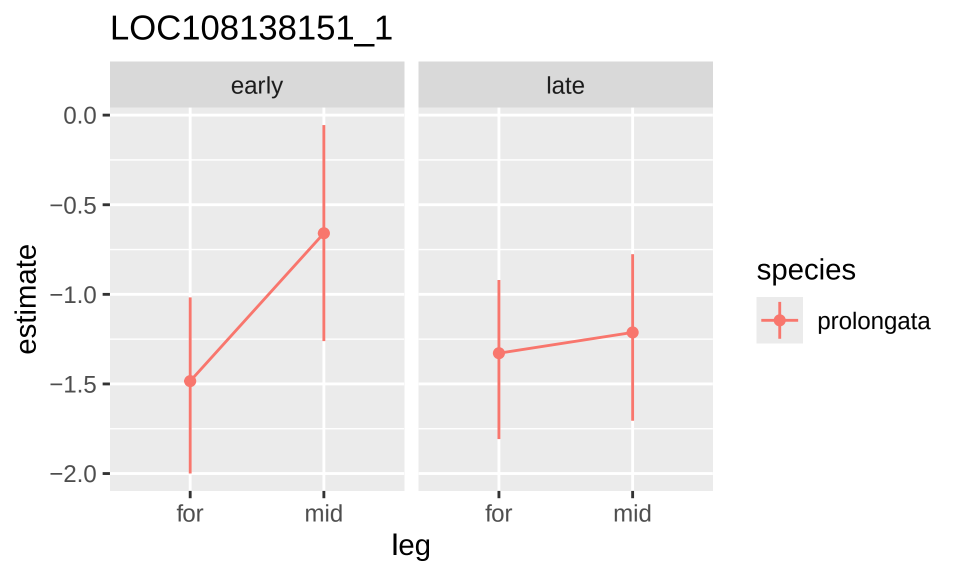

LOC108138151_1
early
late
0.0
−0.5
−1.0
−1.5
−2.0
estimate
for
mid
for
mid
leg
species
prolongata

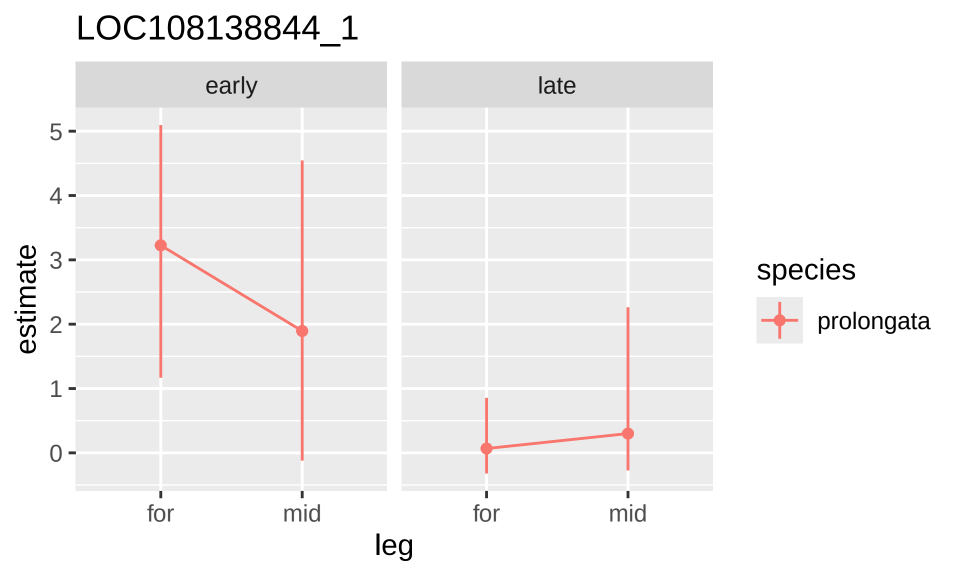

LOC108138844_1
early
late
5
4
3
2
1
0
estimate
for
mid
for
mid
leg
species
prolongata

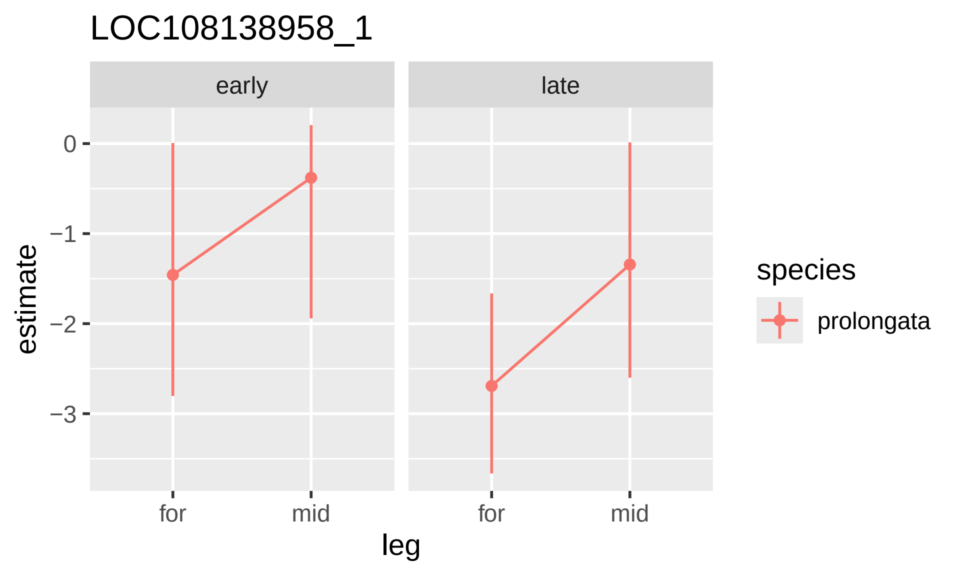

LOC108138958_1
early
late
0
−1
−2
−3
estimate
for
mid
for
mid
leg
species
prolongata

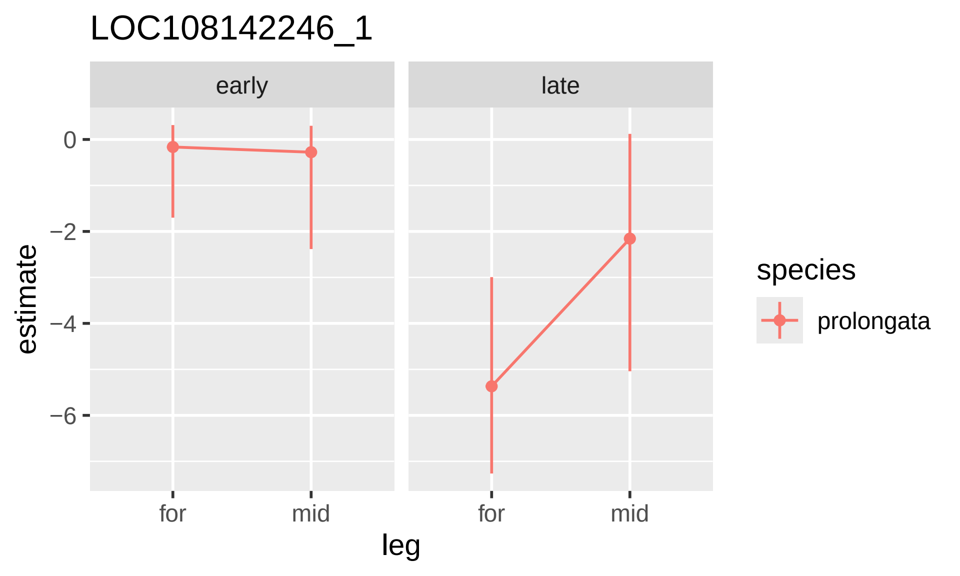
LOC108142246_1
early
late
0
−2
−4
−6
estimate
for
mid
for
mid
leg
species
prolongata

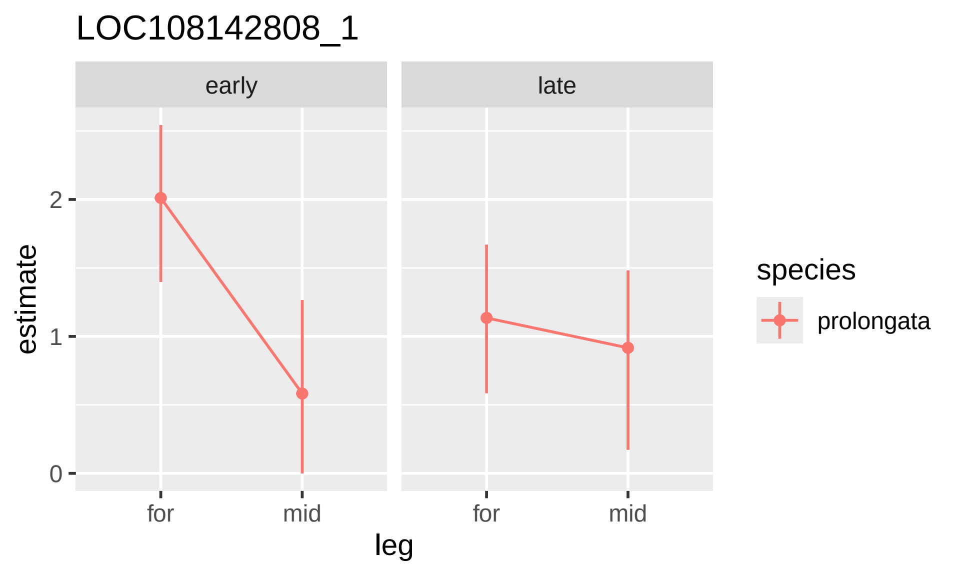
LOC108142808_1
early
late
estimate
2
1
0
for
mid
for
mid
leg
species
prolongata

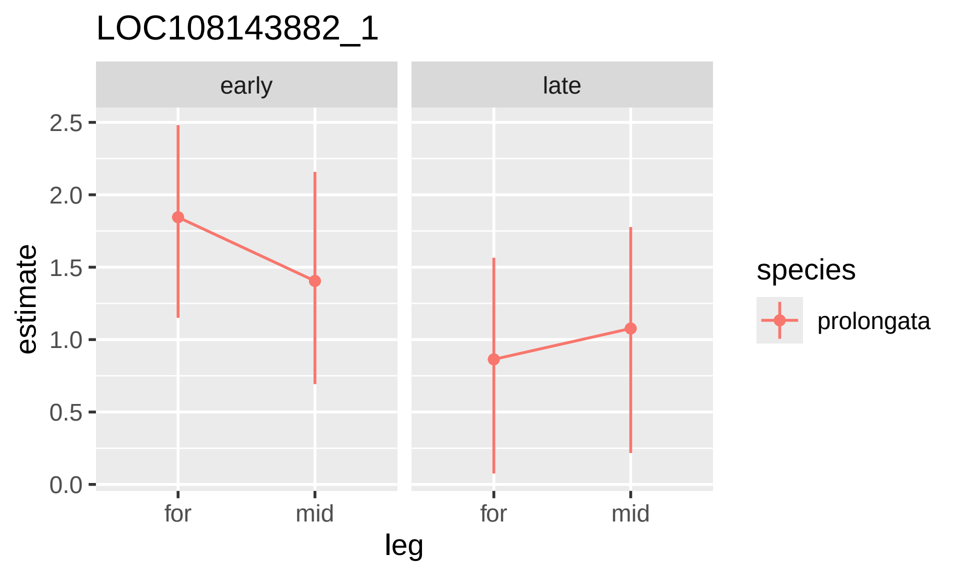

LOC108143882_1
early
late
estimate
2.5
2.0
1.5
1.0
0.5
0.0
for
mid
for
mid
leg
species
prolongata

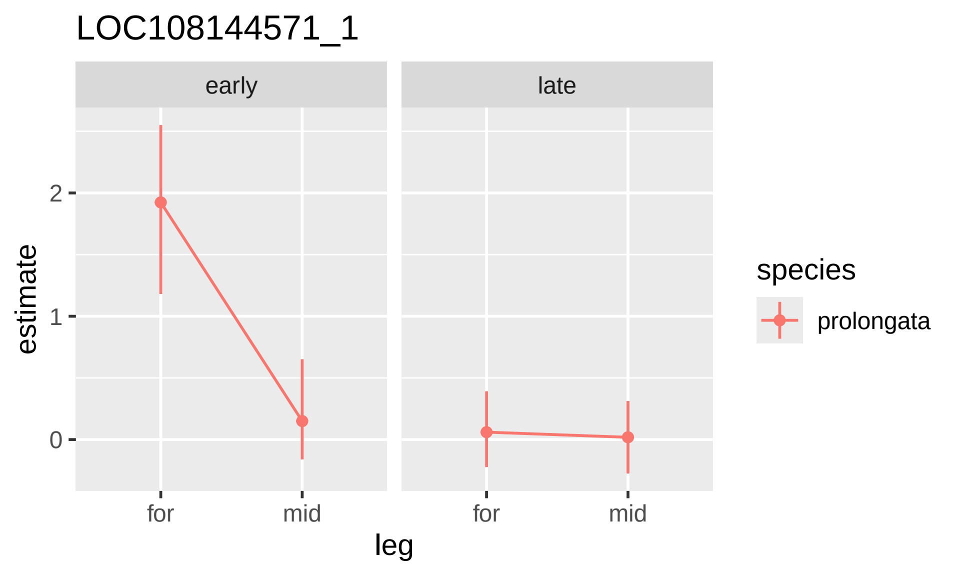
LOC108144571_1
early
late
estimate
2
1
0
for
mid
for
mid
leg
species
prolongata

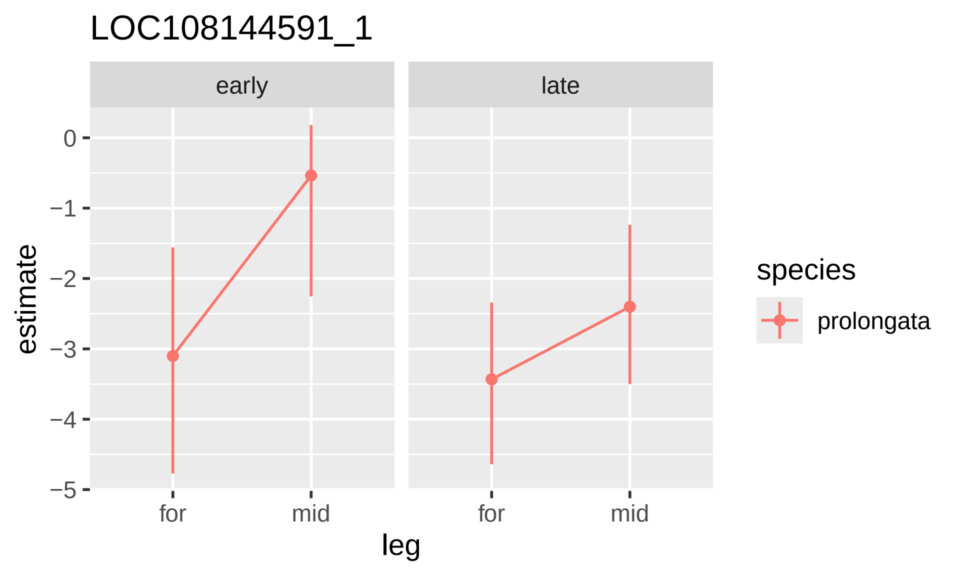

LOC108144591_1
early
late
0
−1
−2
−3
−4
−5
estimate
for
mid
for
mid
leg
species
prolongata

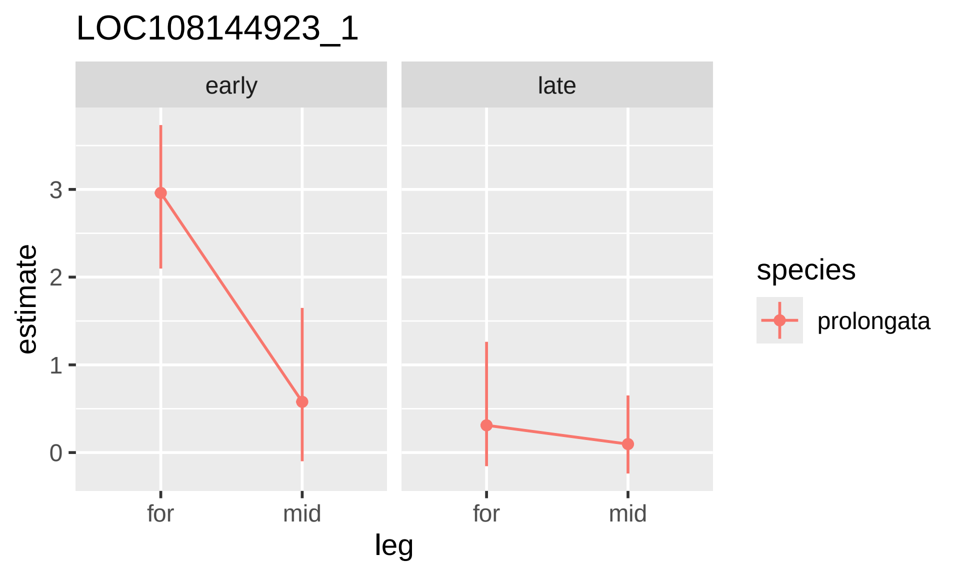

LOC108144923_1
early
late
estimate
3
2
1
0
for
mid
for
mid
leg
species
prolongata

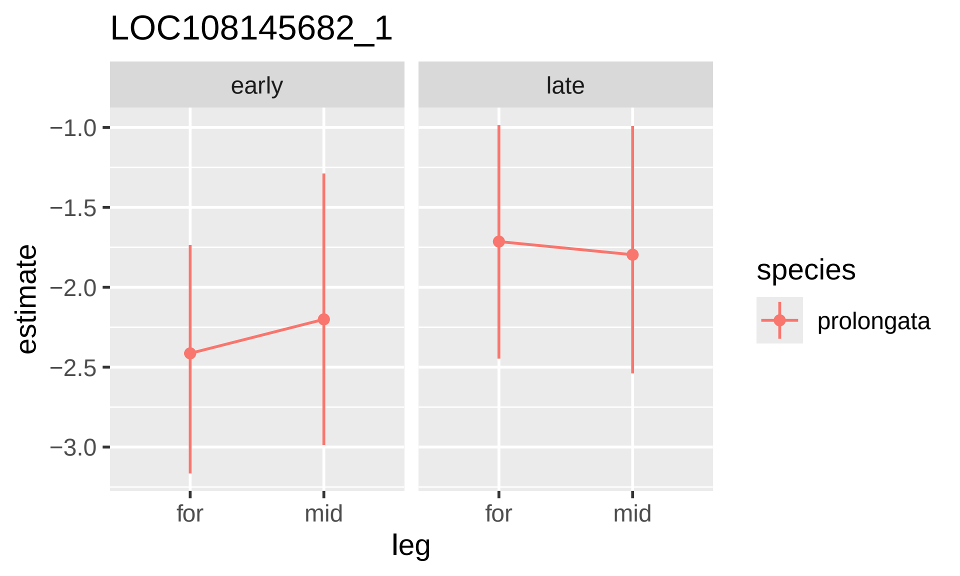

LOC108145682_1
early
late
−1.0
−1.5
−2.0
−2.5
−3.0
estimate
for
mid
for
mid
leg
species
prolongata

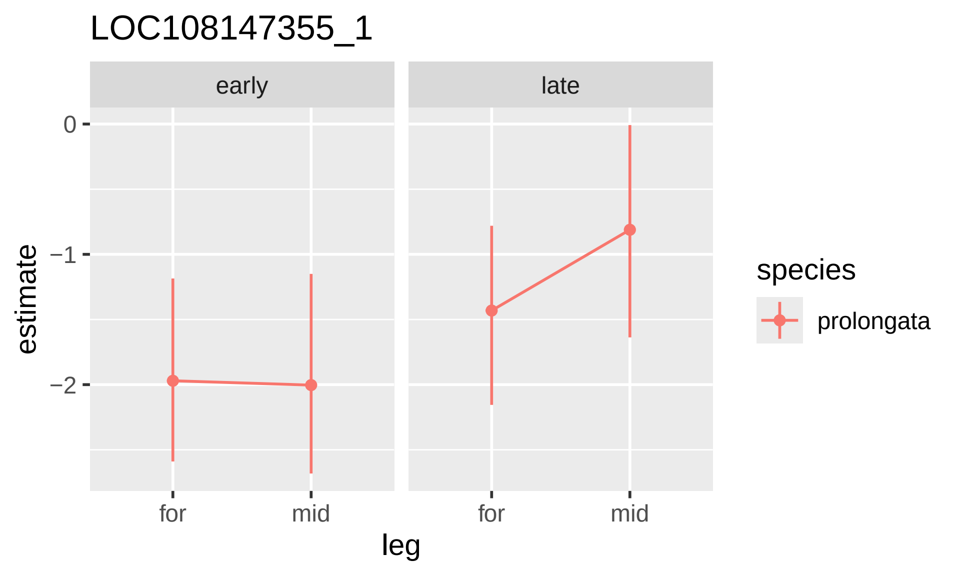

LOC108147355_1
early
late
0
−1
−2
estimate
for
mid
for
mid
leg
species
prolongata

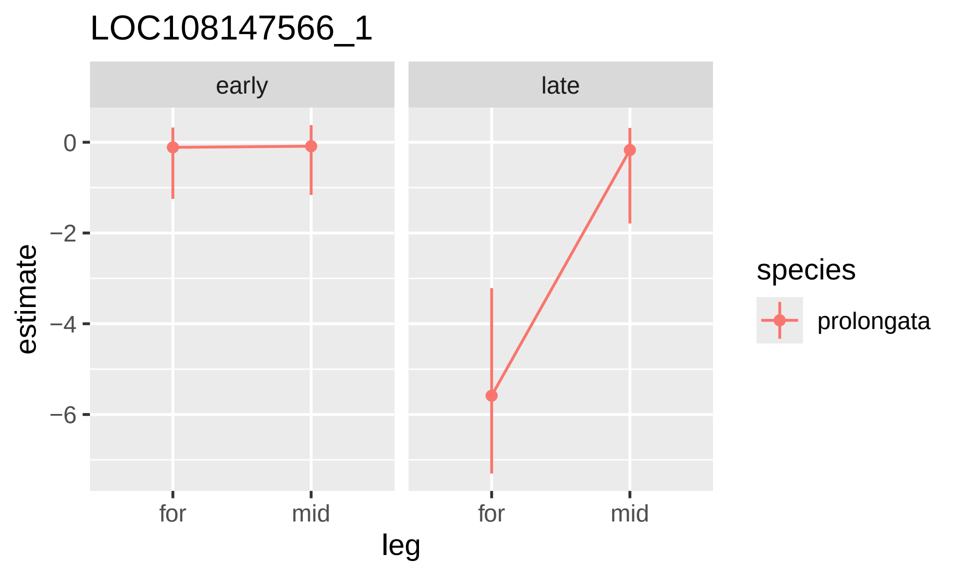
LOC108147566_1
early
late
0
−2
−4
−6
estimate
for
mid
for
mid
leg
species
prolongata

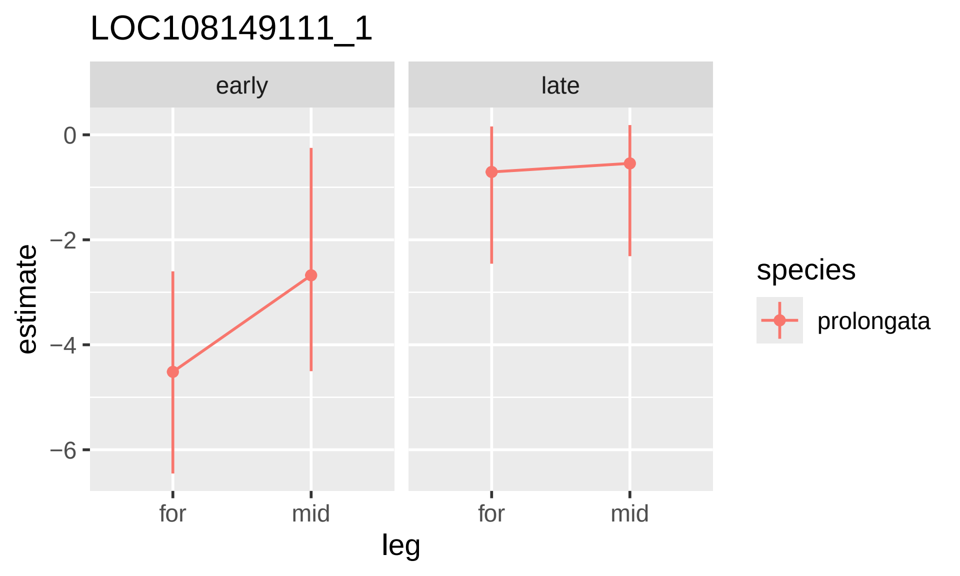

LOC108149111_1
early
late
0
−2
−4
−6
estimate
for
mid
for
mid
leg
species
prolongata

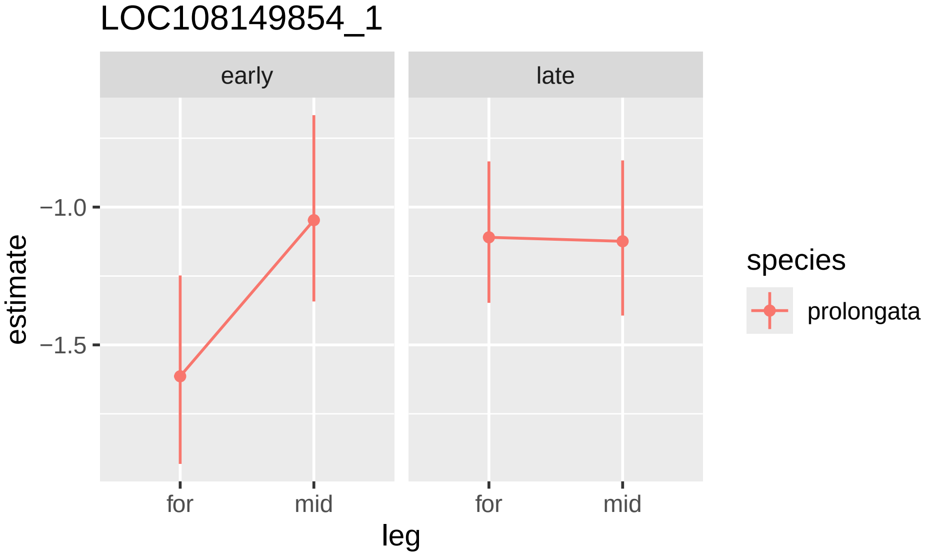

LOC108149854_1
early
late
estimate
−1.0
−1.5
for
mid
for
mid
leg
species
prolongata

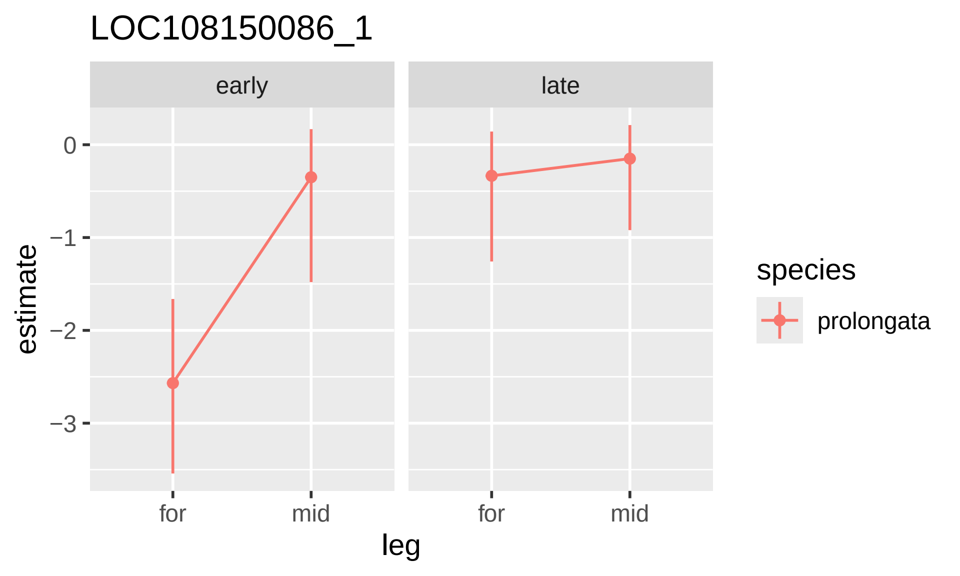

LOC108150086_1
early
late
estimate
0
−1
−2
−3
for
mid
for
mid
leg
species
prolongata

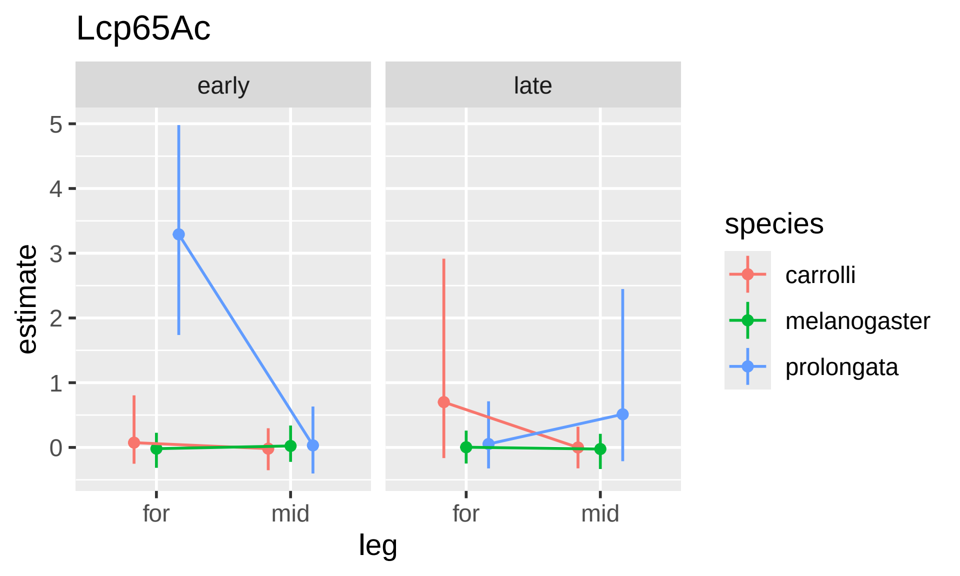

Lcp65Ac
early
late
5
4
3
2
1
0
estimate
for
mid
for
mid
leg
species
carrolli
melanogaster
prolongata

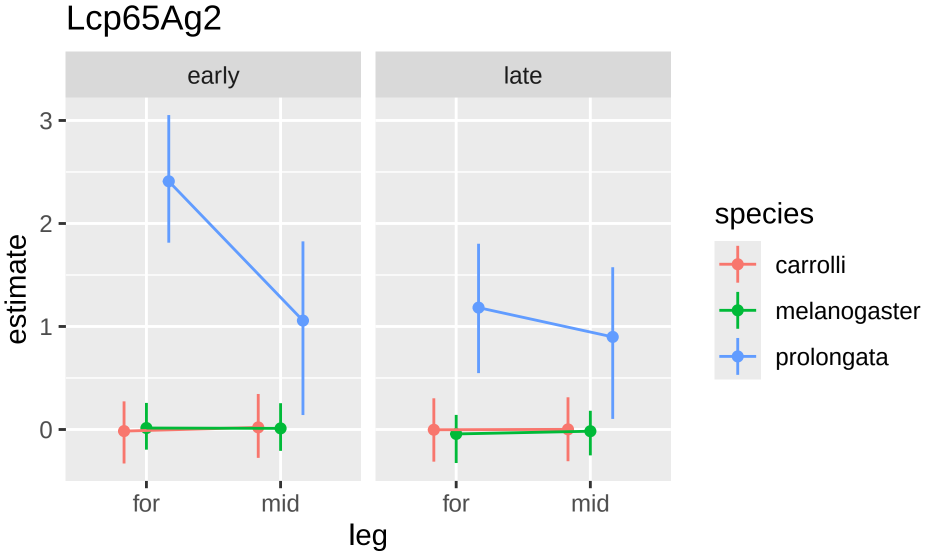

Lcp65Ag2
early
late
3
2
1
0
estimate
for
mid
for
mid
leg
species
carrolli
melanogaster
prolongata

### List

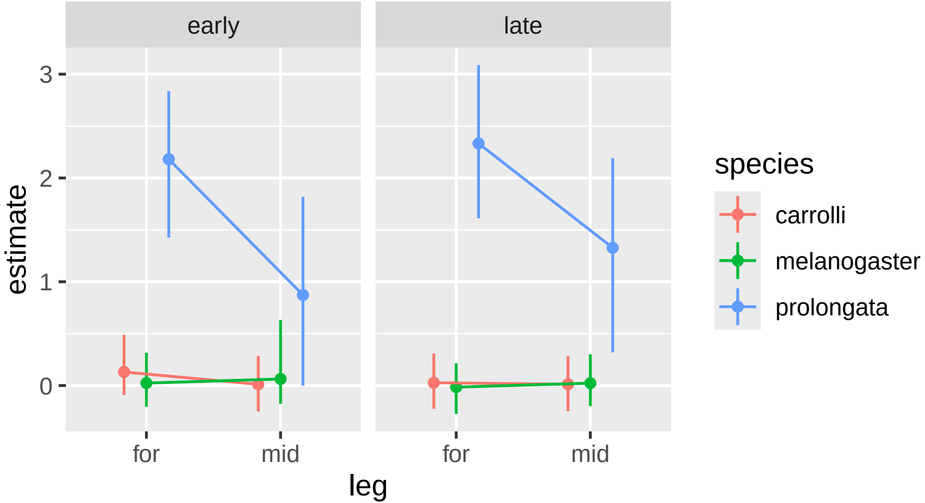

early
late
3
2
1
0
estimate
for
mid
for
mid
leg
species
carrolli
melanogaster
prolongata

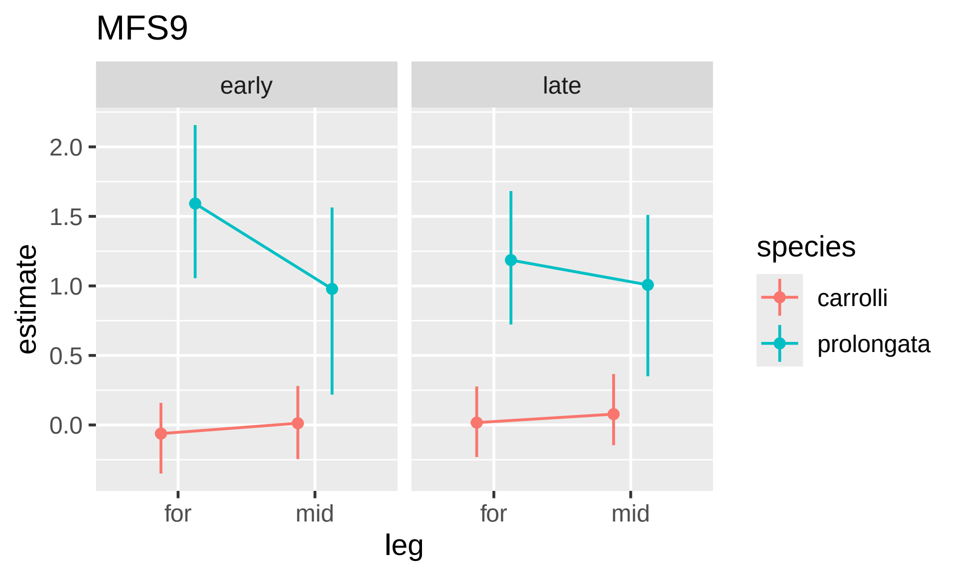

MFS9
early
late
2.0
1.5
1.0
0.5
0.0
estimate
for
mid
for
mid
leg
species
carrolli
prolongata

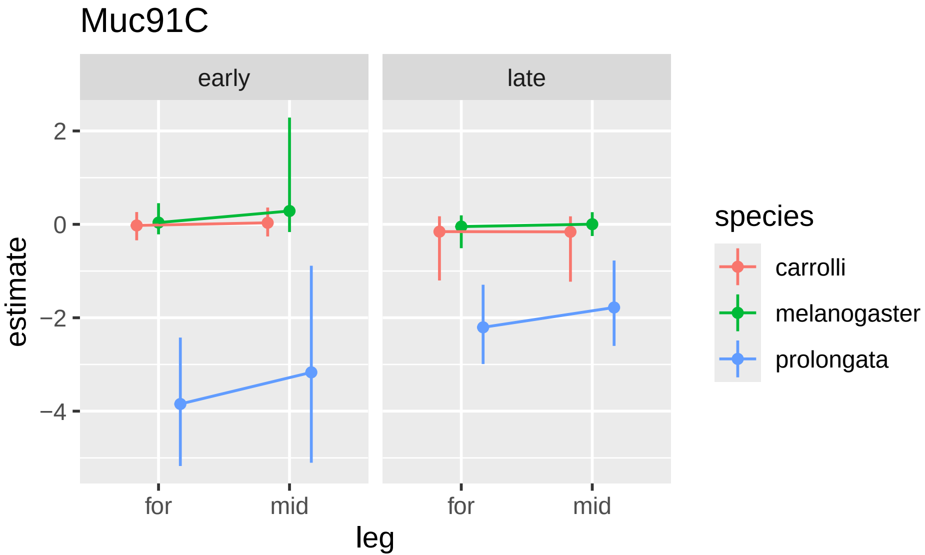

Muc91C
early
late
estimate
2
0
−2
−4
for
mid
for
mid
leg
species
carrolli
melanogaster
prolongata

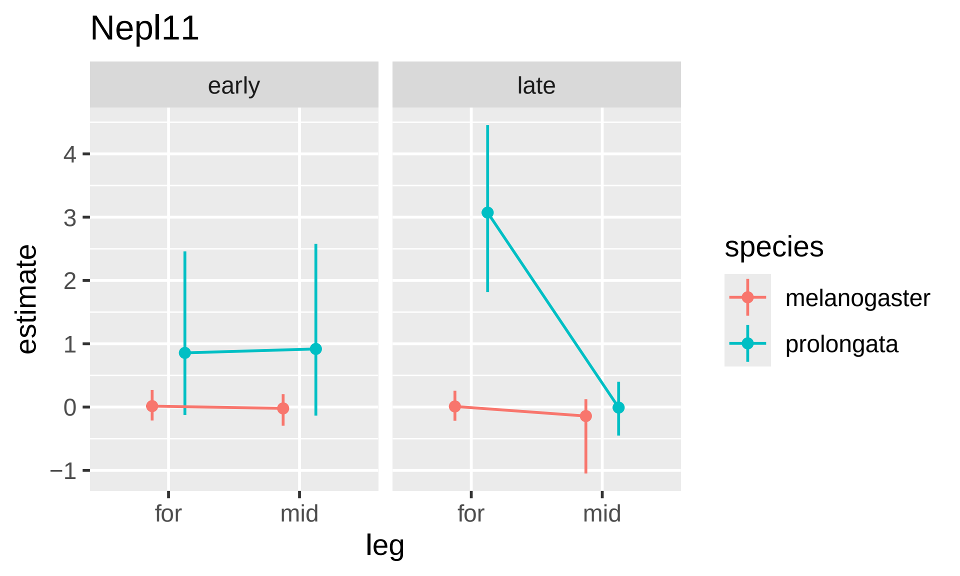

Nepl11
early
late
estimate
4
3
2
1
0
−1
for
mid
for
mid
leg
species
melanogaster
prolongata

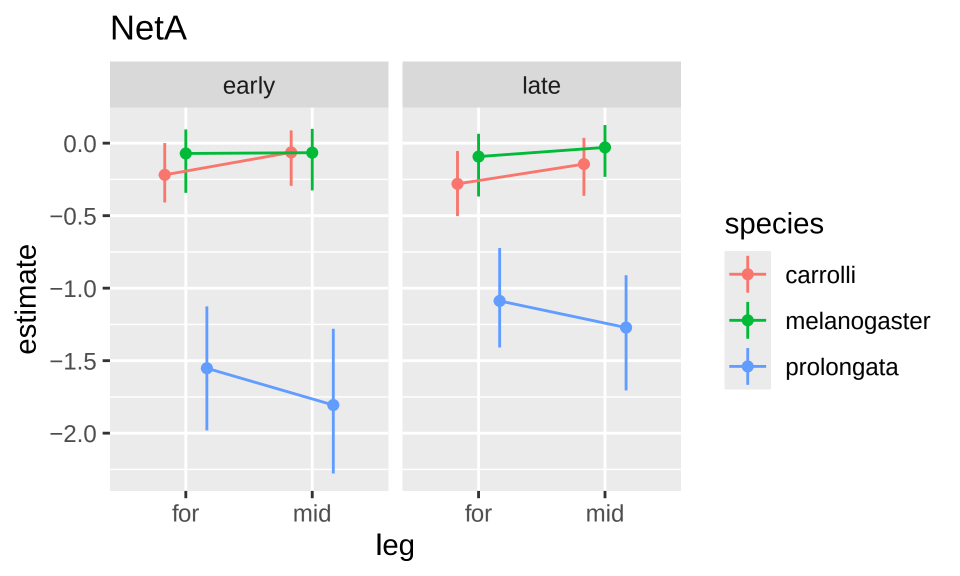
NetA
early
late
0.0
−0.5
−1.0
−1.5
−2.0
estimate
for
mid
for
mid
leg
species
carrolli
melanogaster
prolongata

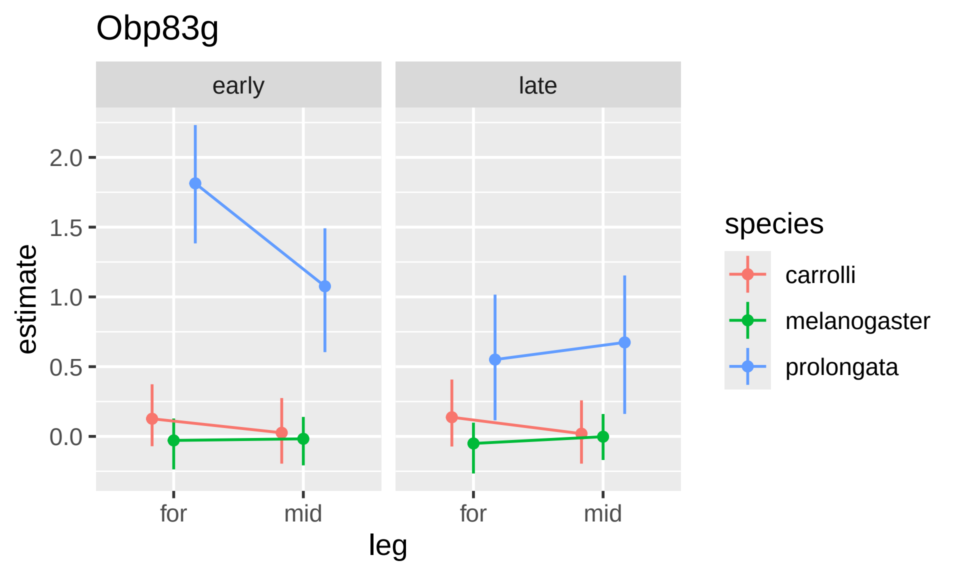

Obp83g
early
late
2.0
1.5
1.0
0.5
0.0
estimate
for
mid
for
mid
leg
species
carrolli
melanogaster
prolongata

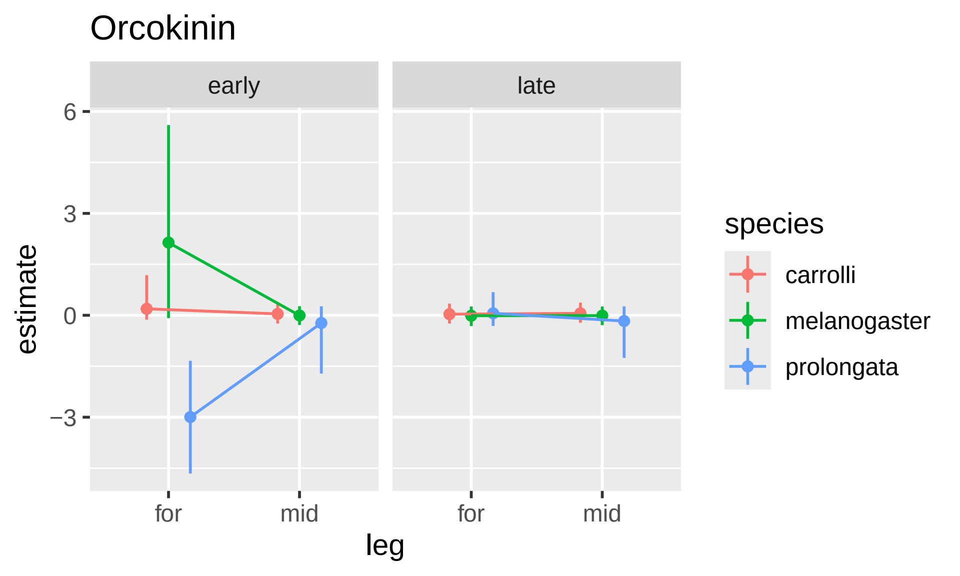

Orcokinin
early
late
6
3
0
−3
estimate
for
mid
for
mid
leg
species
carrolli
melanogaster
prolongata

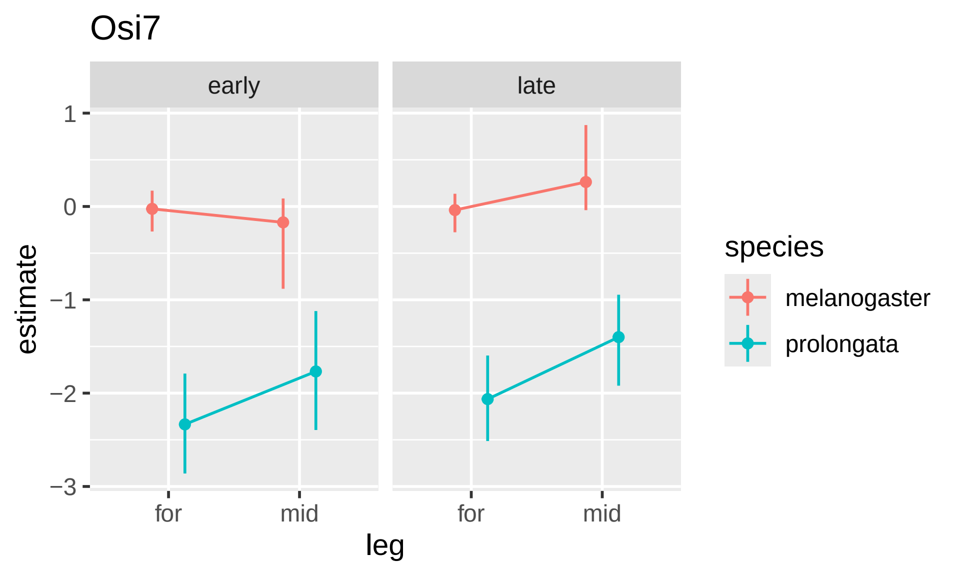

Osi7
early
late
1
0
−1
−2
−3
estimate
for
mid
for
mid
leg
species
melanogaster
prolongata

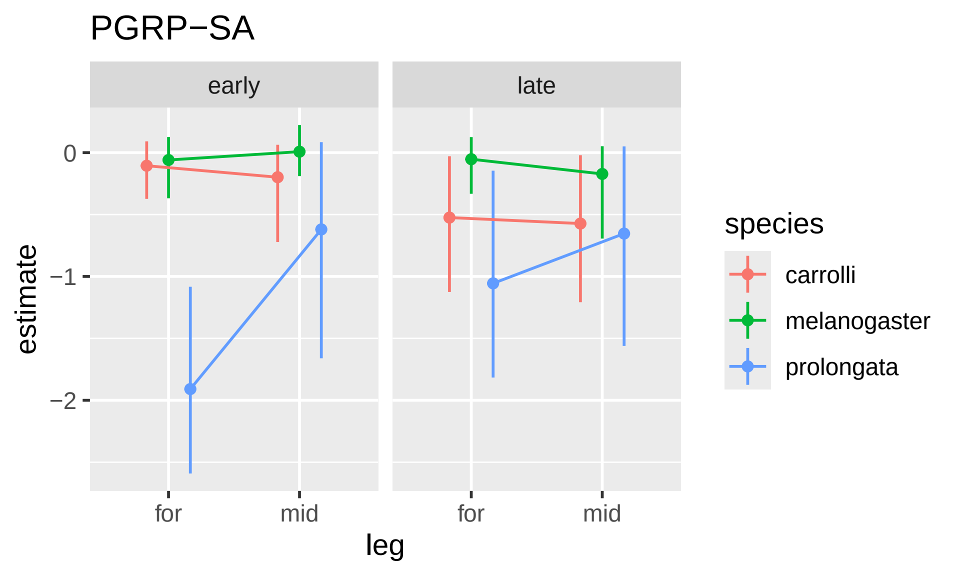

PGRP−SA
early
late
0
−1
−2
estimate
for
mid
for
mid
leg
species
carrolli
melanogaster
prolongata

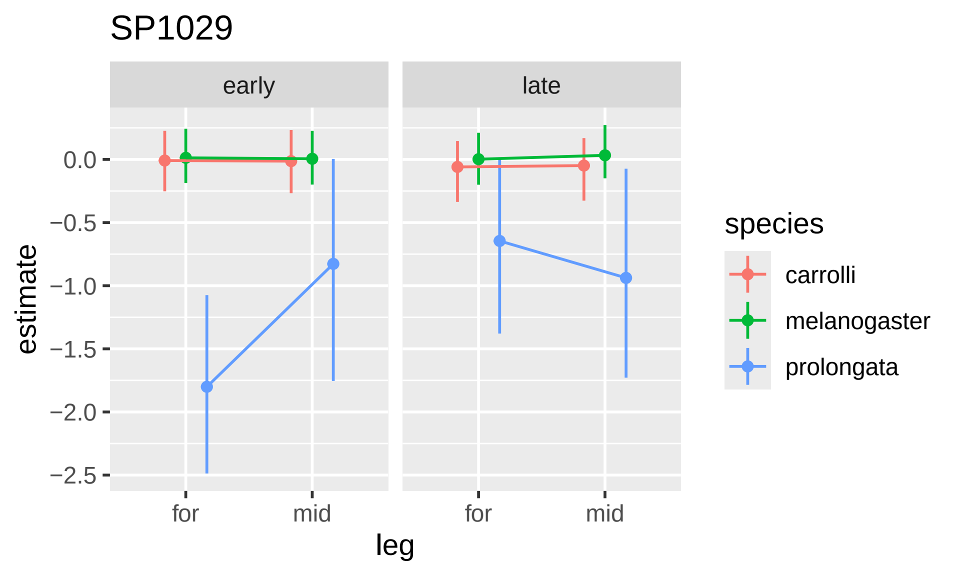

SP1029
early
late
estimate
0.0
−0.5
−1.0
−1.5
−2.0
−2.5
for
mid
for
mid
leg
species
carrolli
melanogaster
prolongata

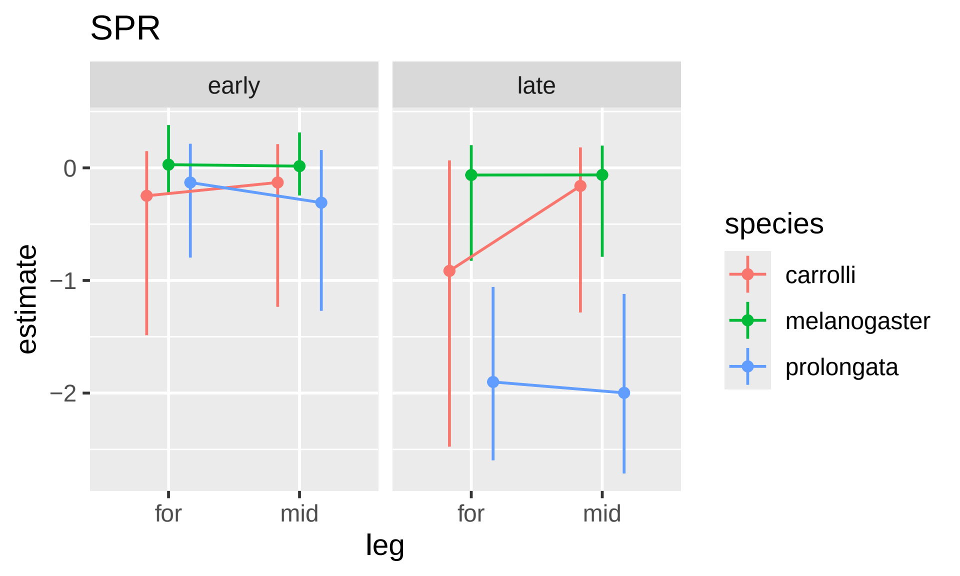

SPR
early
late
0
−1
−2
estimate
for
mid
for
mid
leg
species
carrolli
melanogaster
prolongata

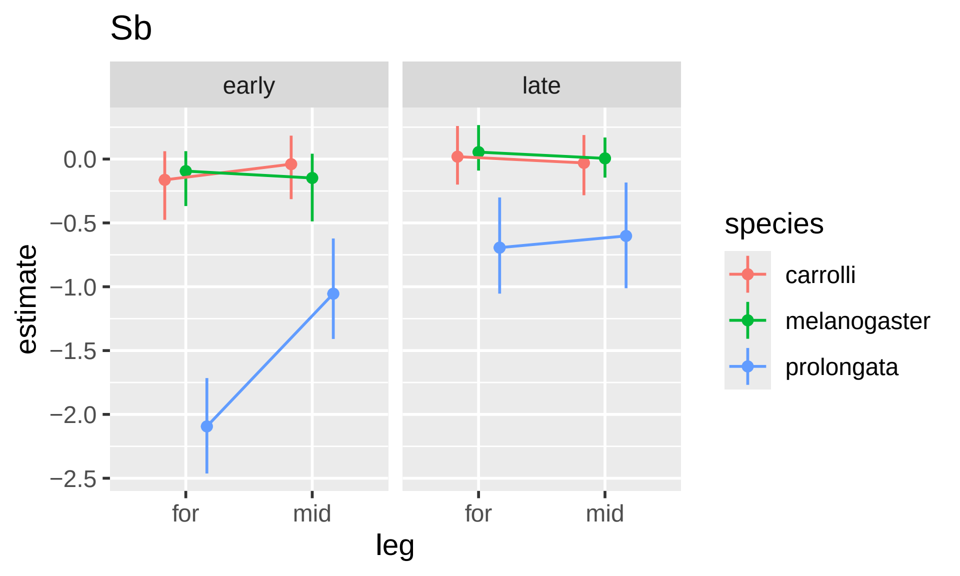

Sb
early
late
0.0
−0.5
−1.0
−1.5
−2.0
−2.5
estimate
for
mid
for
mid
leg
species
carrolli
melanogaster
prolongata

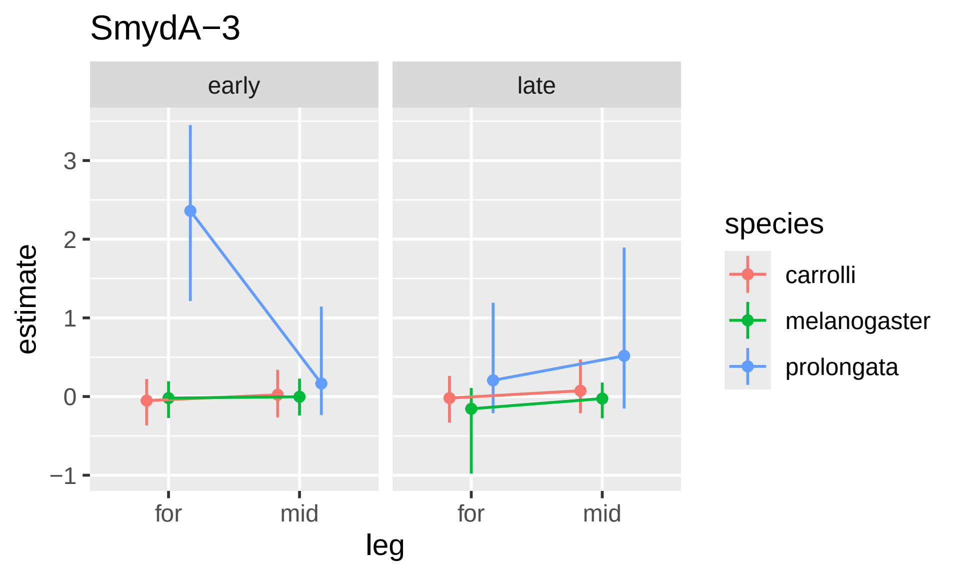

SmydA−3
early
late
estimate
3
2
1
0
−1
for
mid
for
mid
leg
species
carrolli
melanogaster
prolongata

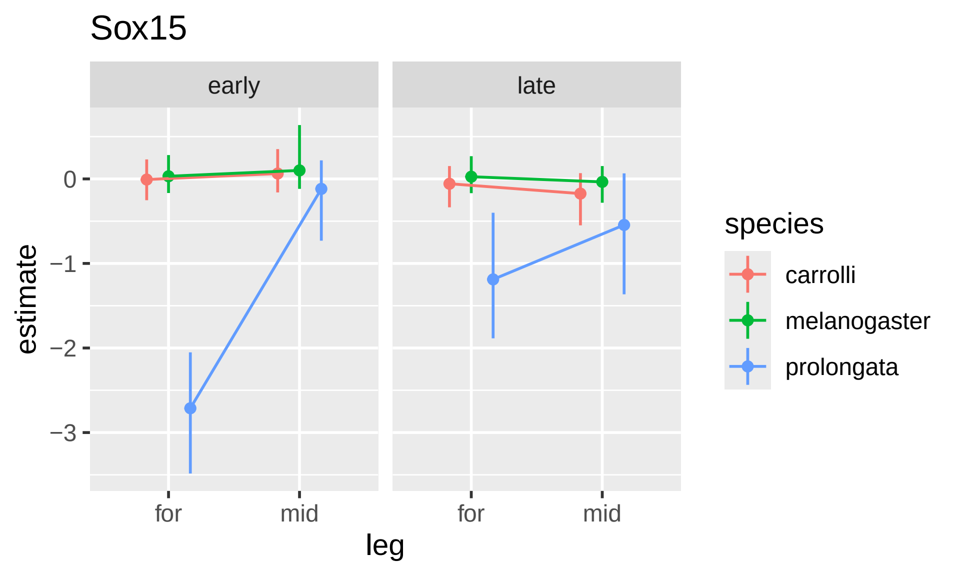

Sox15
early
late
0
−1
−2
−3
estimate
for
mid
for
mid
leg
species
carrolli
melanogaster
prolongata

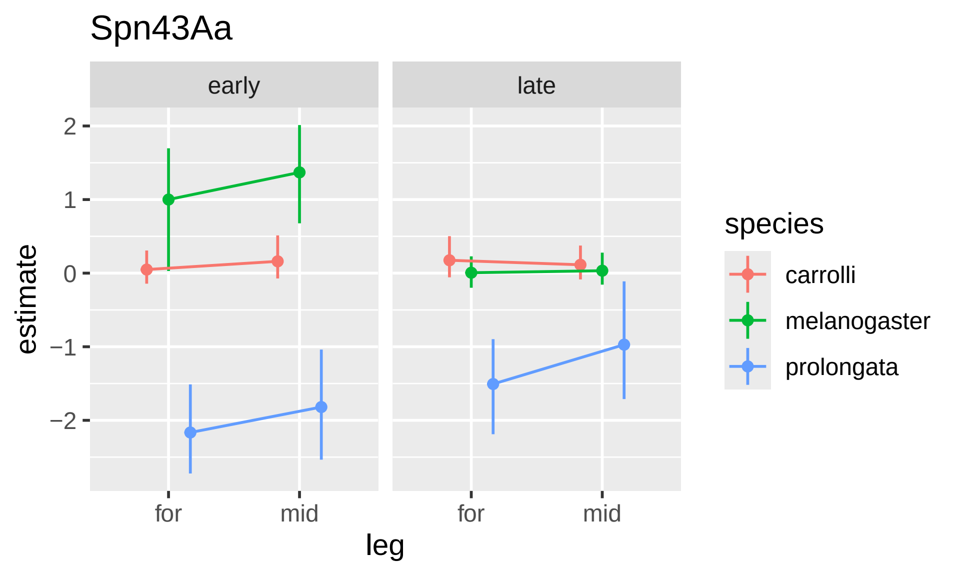

Spn43Aa
early
late
2
1
0
−1
−2
estimate
for
mid
for
mid
leg
species
carrolli
melanogaster
prolongata

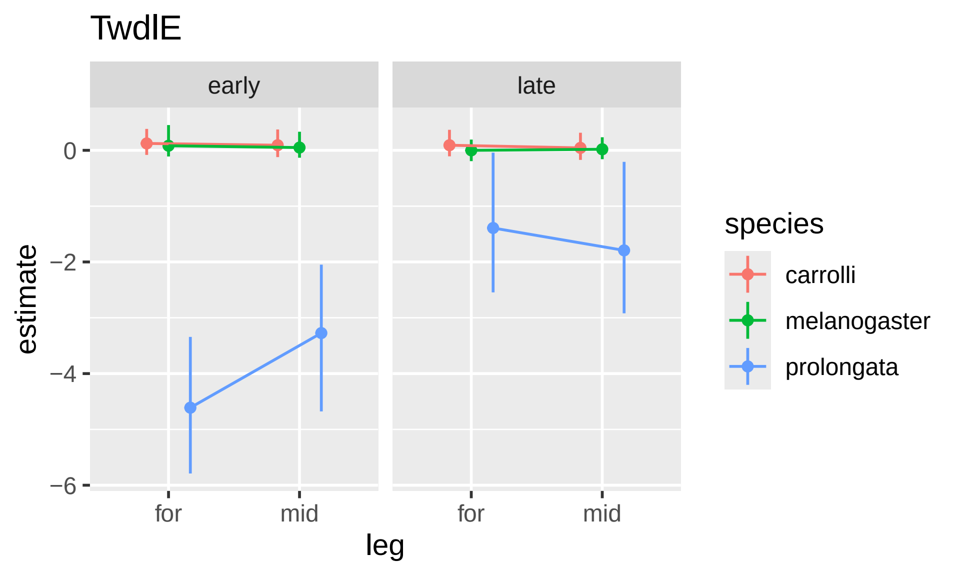

TwdlE
early
late
0
−2
−4
−6
estimate
for
mid
for
mid
leg
species
carrolli
melanogaster
prolongata

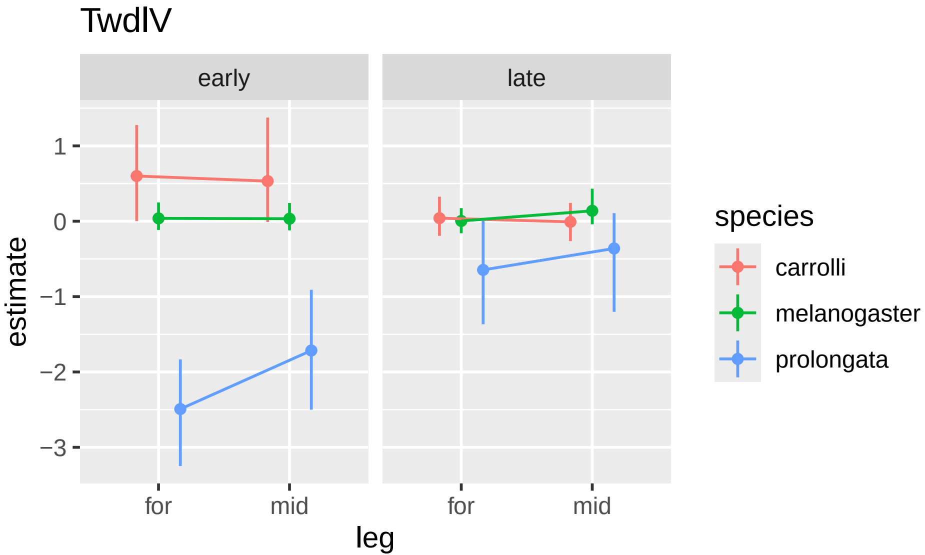
TwdlV
early
late
1
0
−1
−2
−3
estimate
for
mid
for
mid
leg
species
carrolli
melanogaster
prolongata

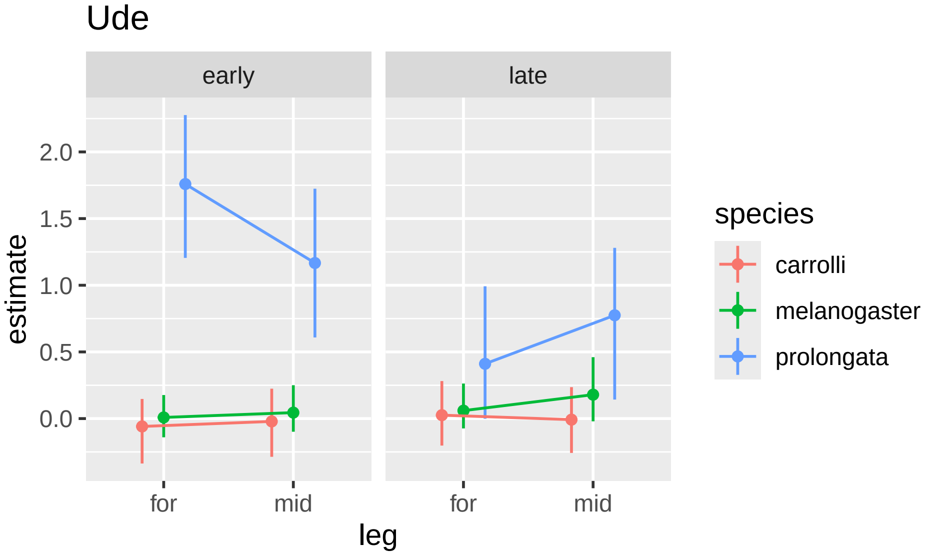

Ude
early
late
2.0
1.5
1.0
0.5
0.0
estimate
for
mid
for
mid
leg
species
carrolli
melanogaster
prolongata

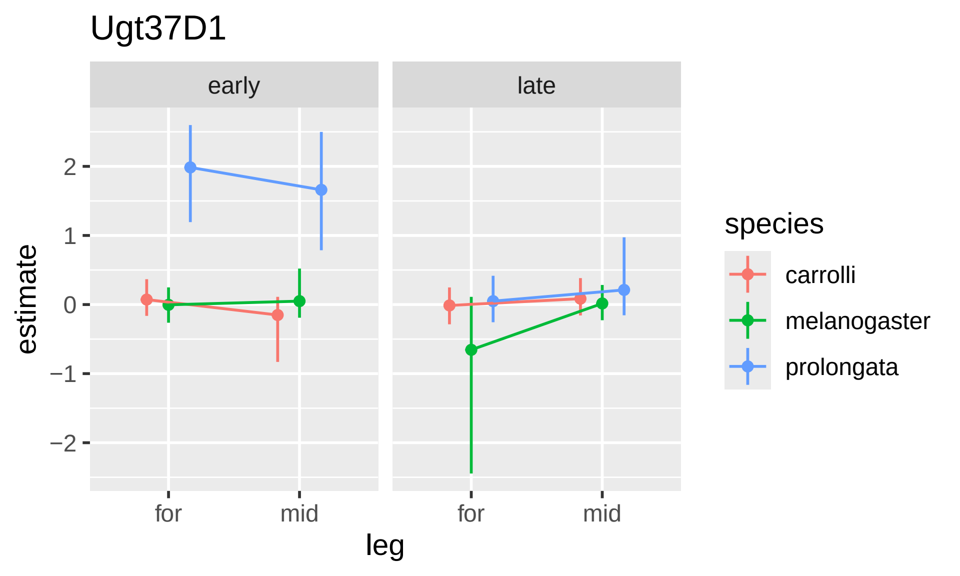
Ugt37D1
early
late
2
1
0
−1
−2
estimate
for
mid
for
mid
leg
species
carrolli
melanogaster
prolongata

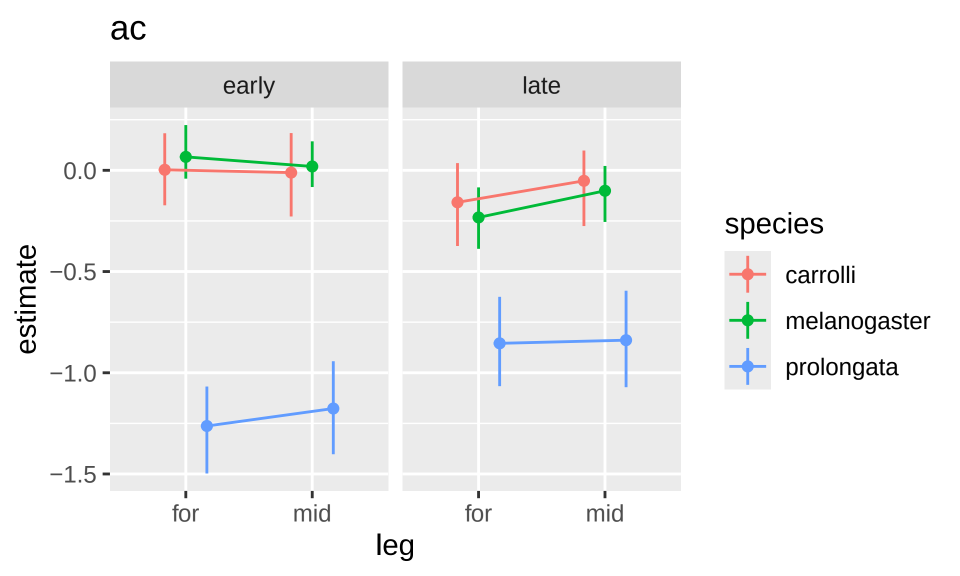

ac
early
late
0.0
−0.5
−1.0
−1.5
estimate
for
mid
for
mid
leg
species
carrolli
melanogaster
prolongata

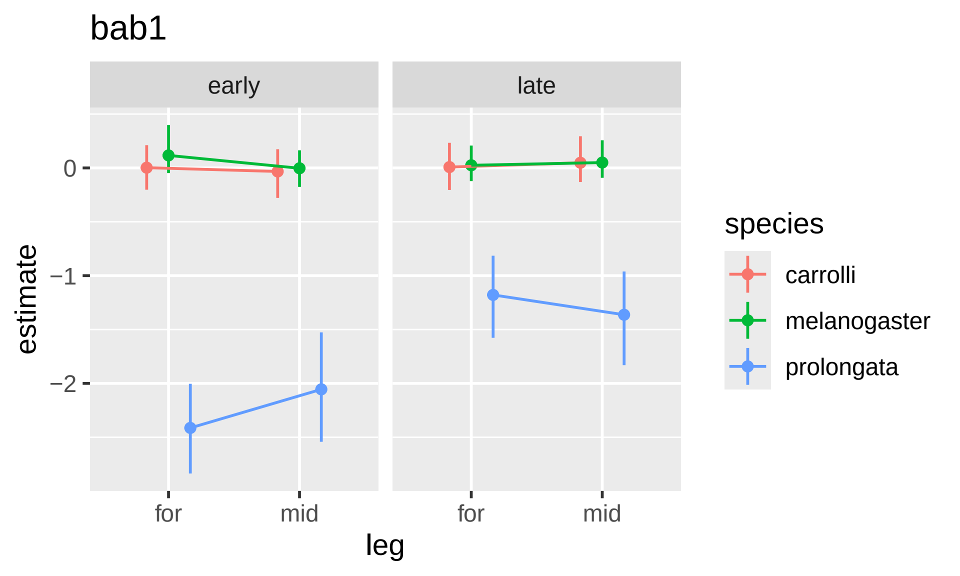
bab1
early
late
0
−1
−2
estimate
for
mid
for
mid
leg
species
carrolli
melanogaster
prolongata

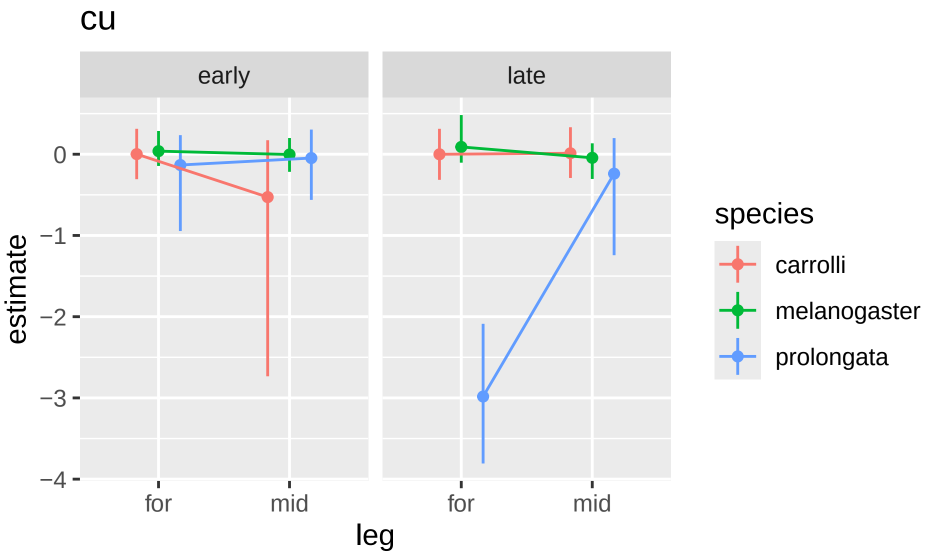
cu
early
late
0
−1
−2
−3
−4
estimate
for
mid
for
mid
leg
species
carrolli
melanogaster
prolongata

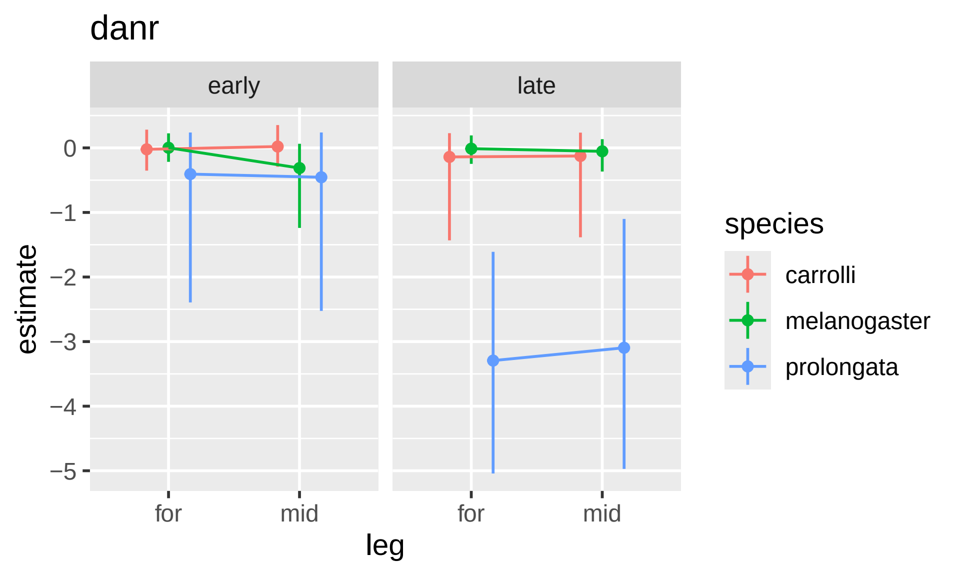
danr
early
late
0
−1
−2
−3
−4
−5
estimate
for
mid
for
mid
leg
species
carrolli
melanogaster
prolongata

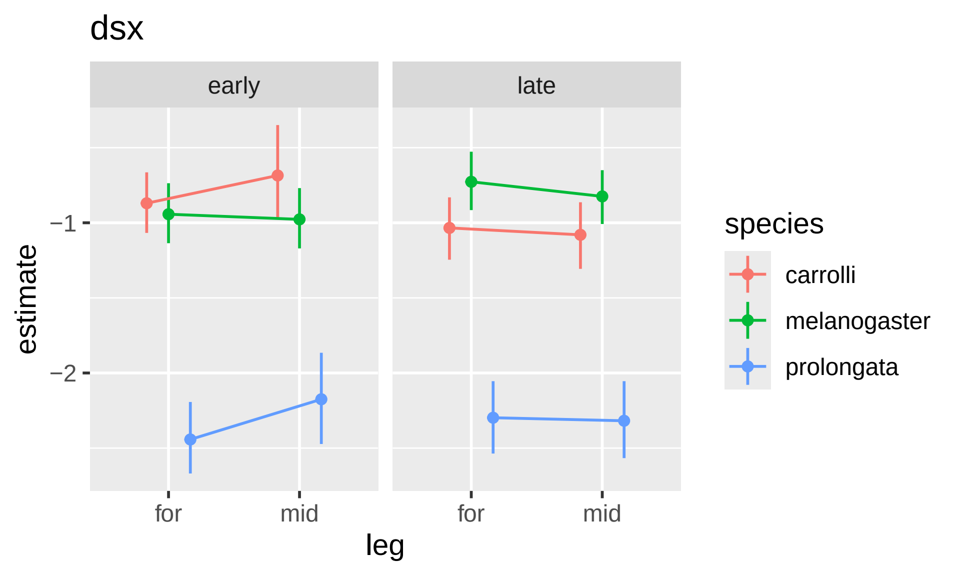

dsx
early
late
−1
−2
estimate
for
mid
for
mid
leg
species
carrolli
melanogaster
prolongata

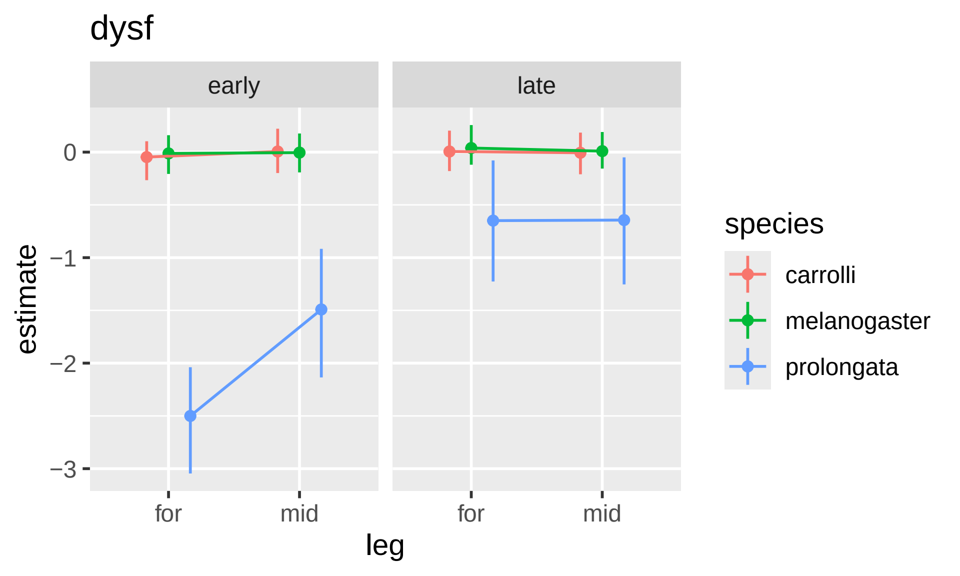
dysf
early
late
estimate
0
−1
−2
−3
for
mid
for
mid
leg
species
carrolli
melanogaster
prolongata

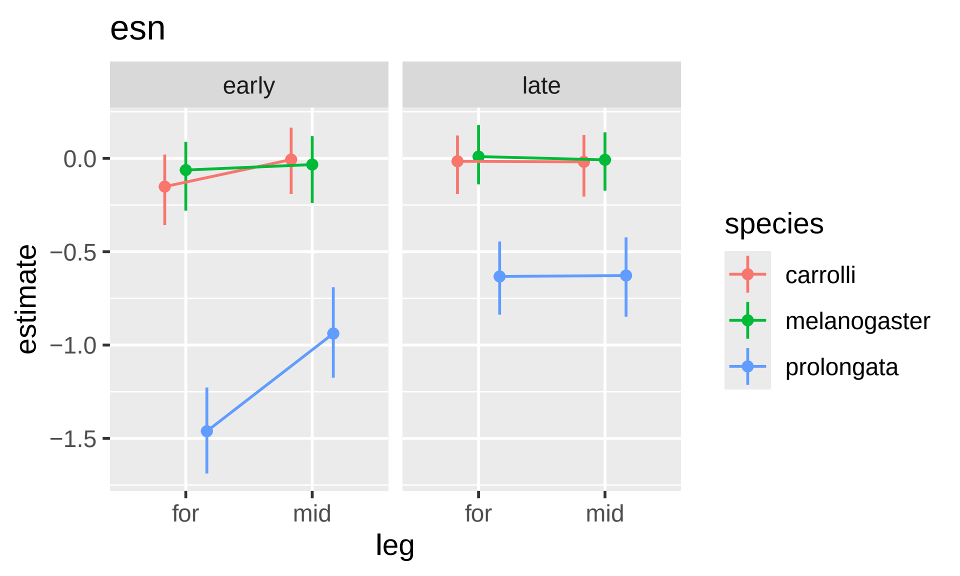
esn
early
late
0.0
−0.5
−1.0
−1.5
estimate
for
mid
for
mid
leg
species
carrolli
melanogaster
prolongata

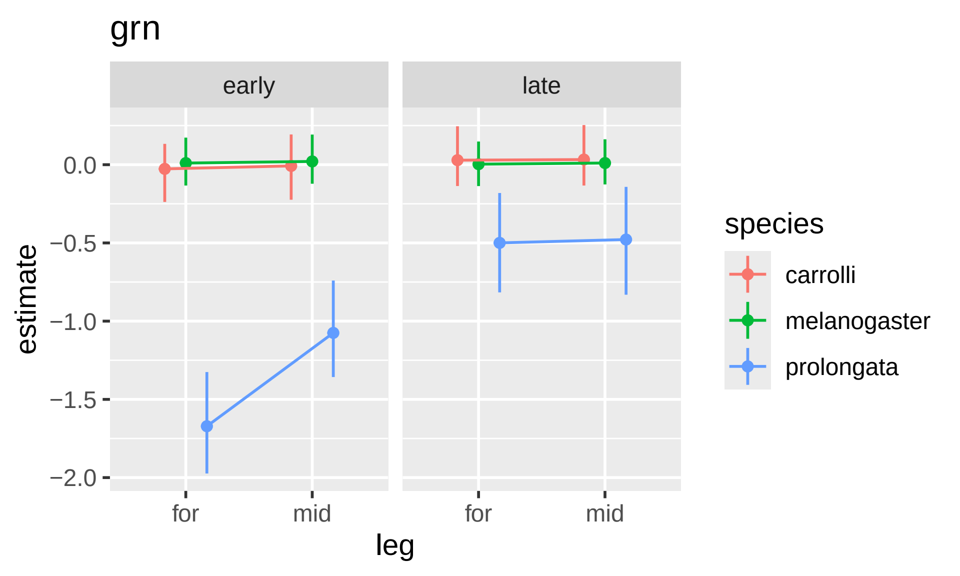

grn
early
late
0.0
−0.5
−1.0
−1.5
−2.0
estimate
for
mid
for
mid
leg
species
carrolli
melanogaster
prolongata

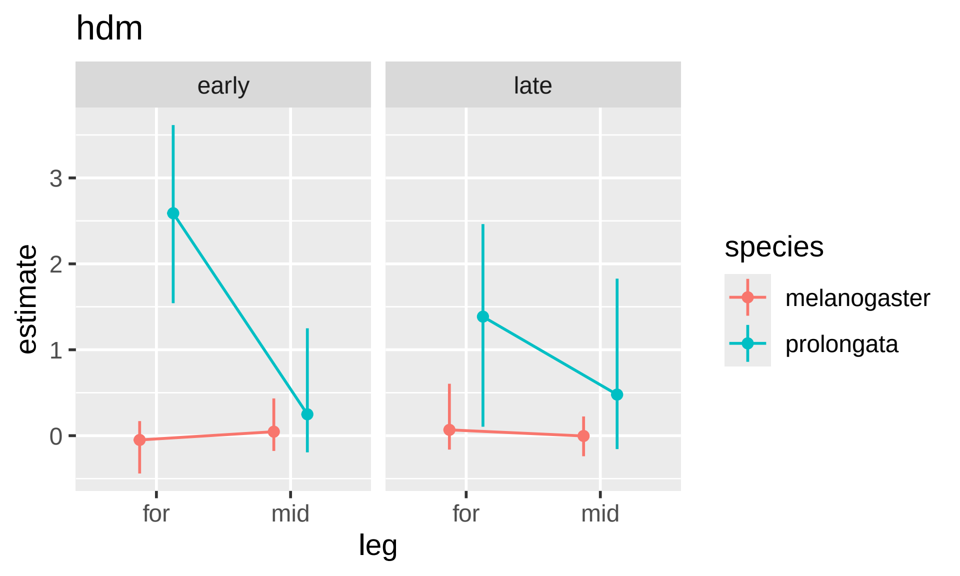

hdm
early
late
estimate
3
2
1
0
for
mid
for
mid
leg
species
melanogaster
prolongata

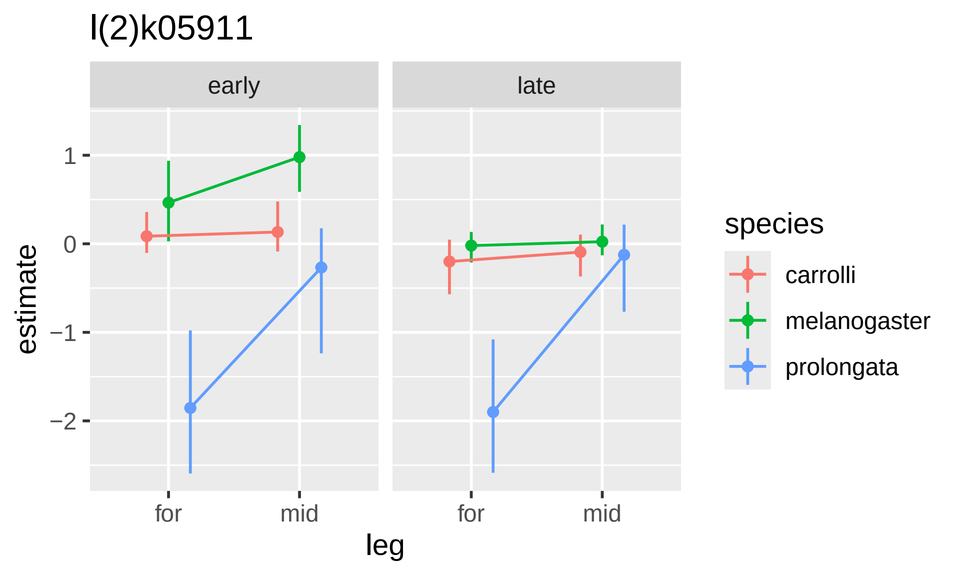

l(2)k05911
early
late
1
0
−1
−2
estimate
for
mid
for
mid
leg
species
carrolli
melanogaster
prolongata

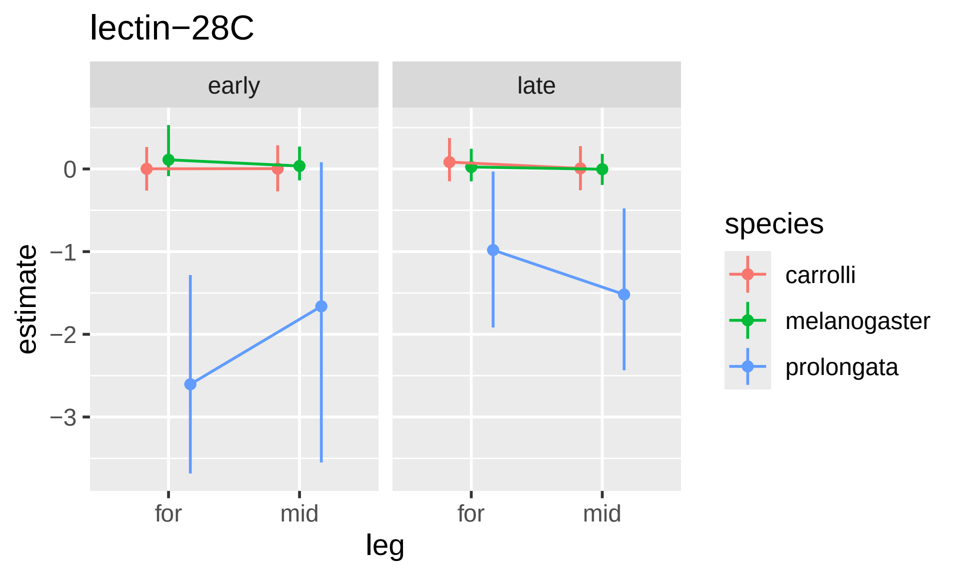
lectin−28C
early
late
estimate
0
−1
−2
−3
for
mid
for
mid
leg
species
carrolli
melanogaster
prolongata

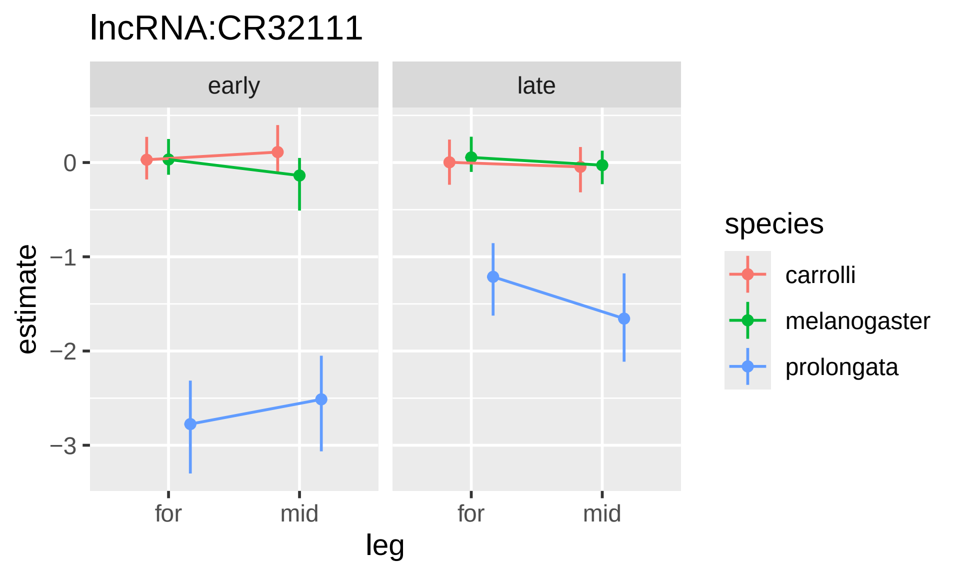

lncRNA:CR32111
early
late
estimate
0
−1
−2
−3
for
mid
for
mid
leg
species
carrolli
melanogaster
prolongata

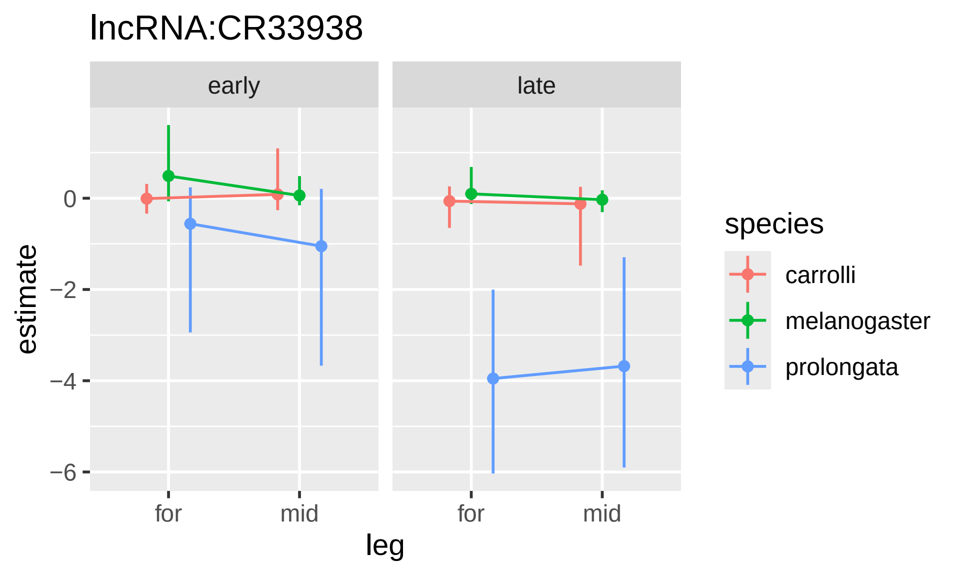

lncRNA:CR33938
early
late
estimate
0
−2
−4
−6
for
mid
for
mid
leg
species
carrolli
melanogaster
prolongata

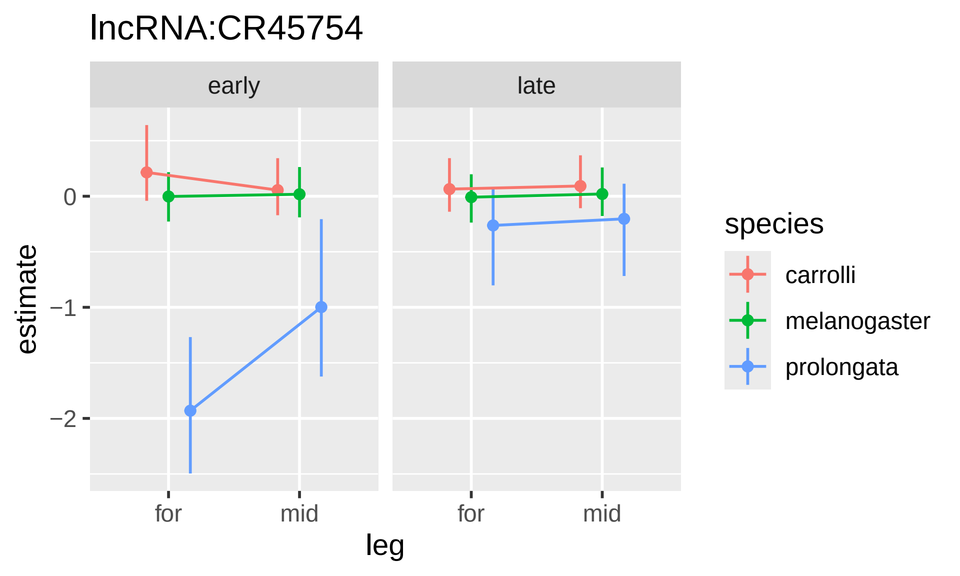

lncRNA:CR45754
early
late
estimate
0
−1
−2
for
mid
for
mid
leg
species
carrolli
melanogaster
prolongata

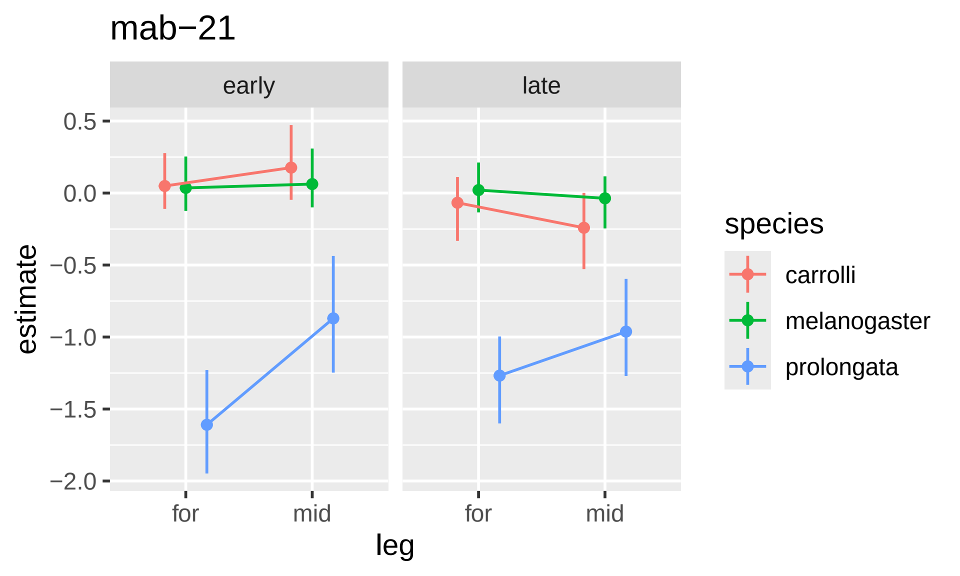

mab−21
early
late
0.5
0.0
−0.5
−1.0
−1.5
−2.0
estimate
for
mid
for
mid
leg
species
carrolli
melanogaster
prolongata

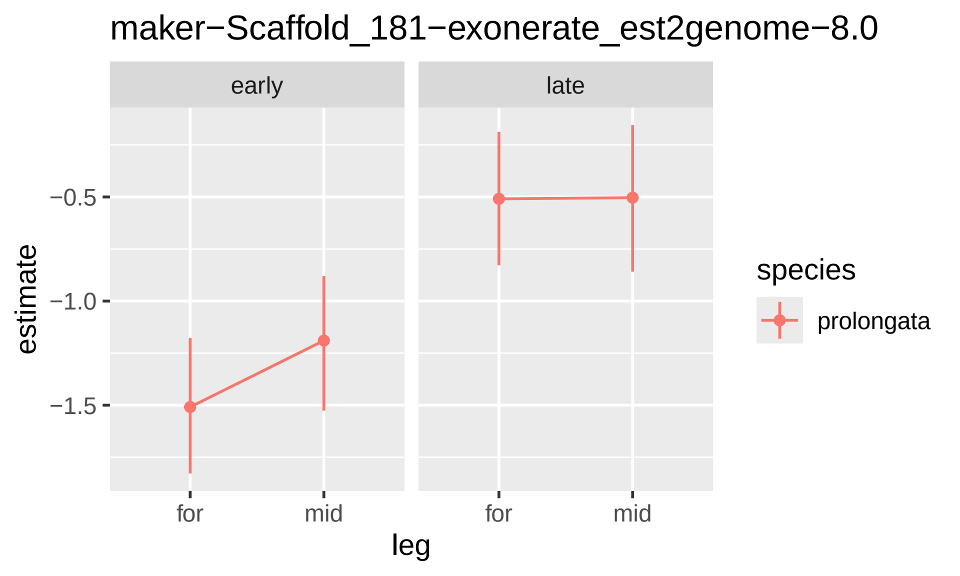
maker−Scaffold_181−exonerate_est2genome−8.0
early
late
−0.5
−1.0
−1.5
estimate
for
mid
for
mid
leg
species
prolongata

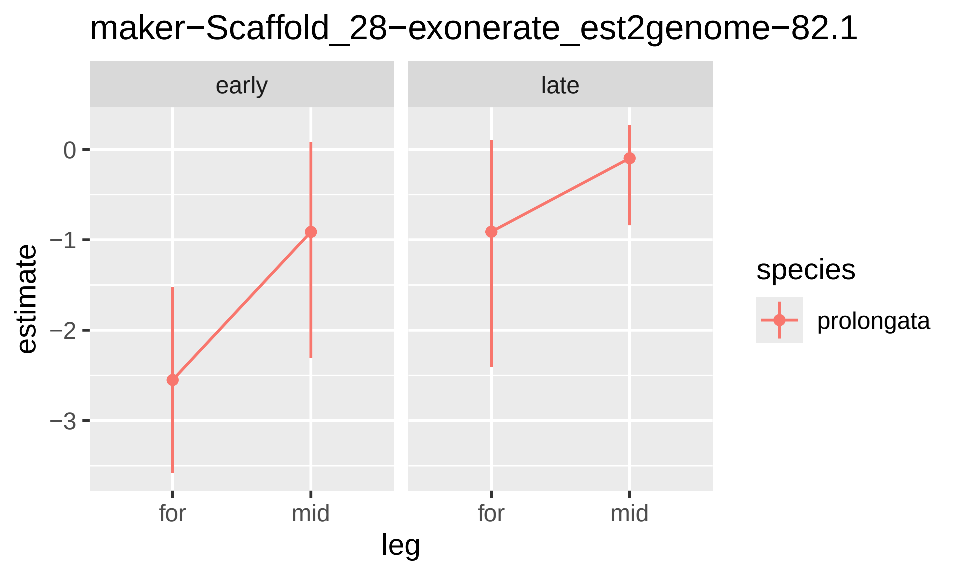

maker−Scaffold_28−exonerate_est2genome−82.1
early
late
0
−1
−2
−3
estimate
for
mid
for
mid
leg
species
prolongata

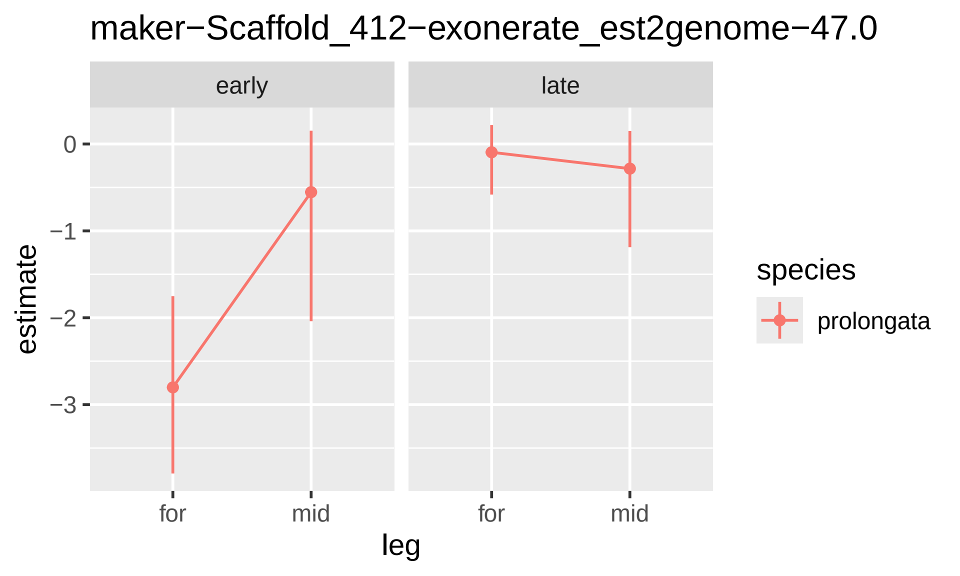
maker−Scaffold_412−exonerate_est2genome−47.0
early
late
0
−1
−2
−3
estimate
for
mid
for
mid
leg
species
prolongata

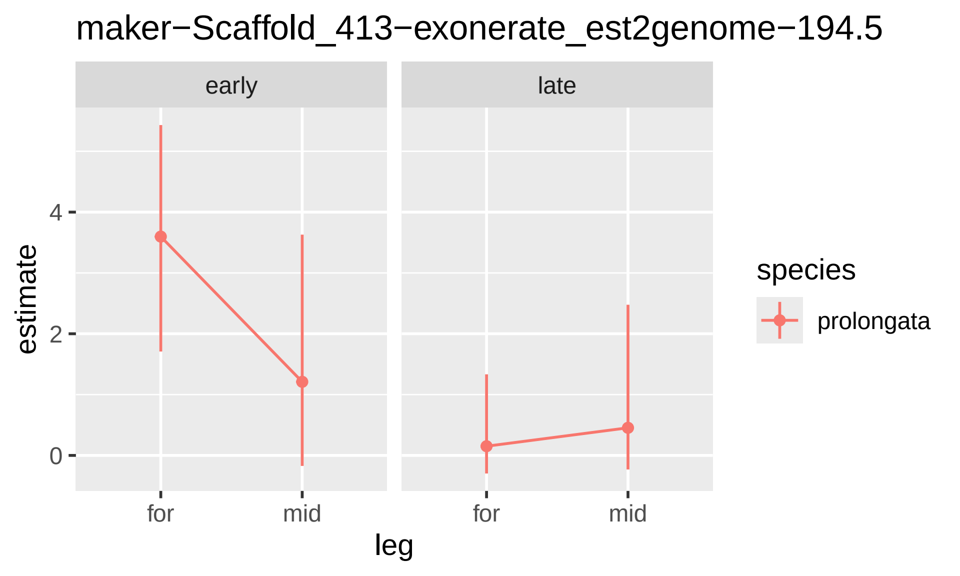
maker−Scaffold_413−exonerate_est2genome−194.5
early
late
estimate
4
2
0
for
mid
for
mid
leg
species
prolongata

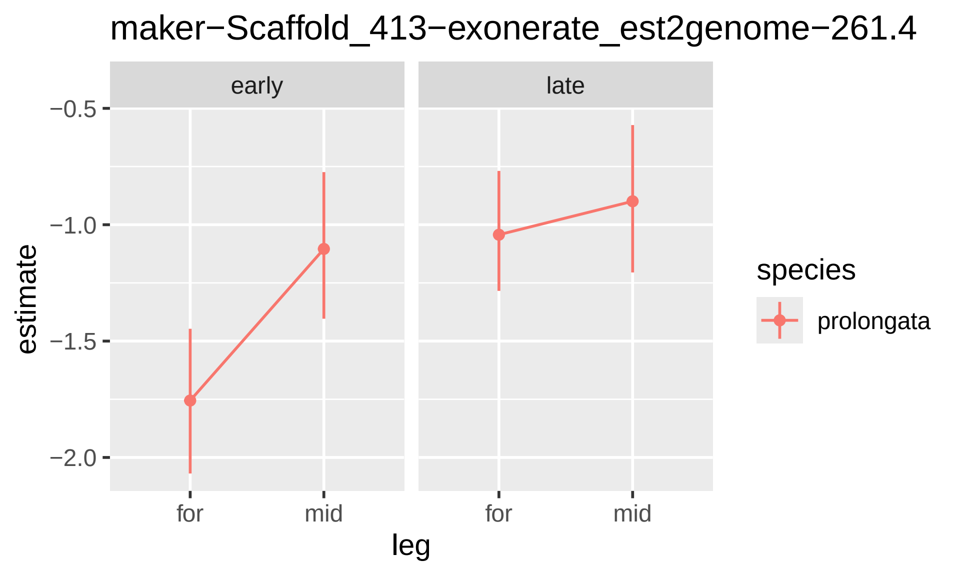

maker−Scaffold_413−exonerate_est2genome−261.4
early
late
−0.5
−1.0
−1.5
−2.0
estimate
for
mid
for
mid
leg
species
prolongata

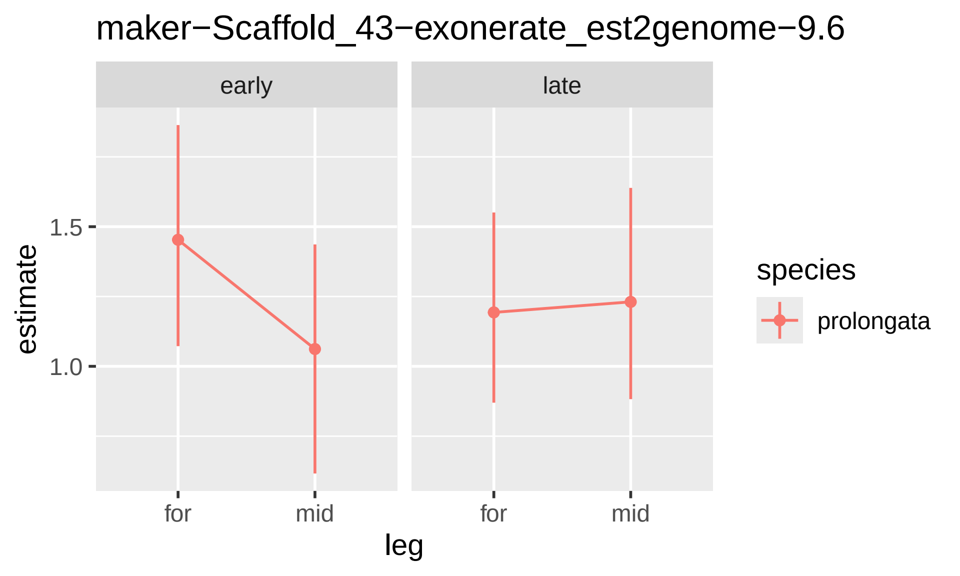

maker−Scaffold_43−exonerate_est2genome−9.6
early
late
1.5
1.0
estimate
for
mid
for
mid
leg
species
prolongata

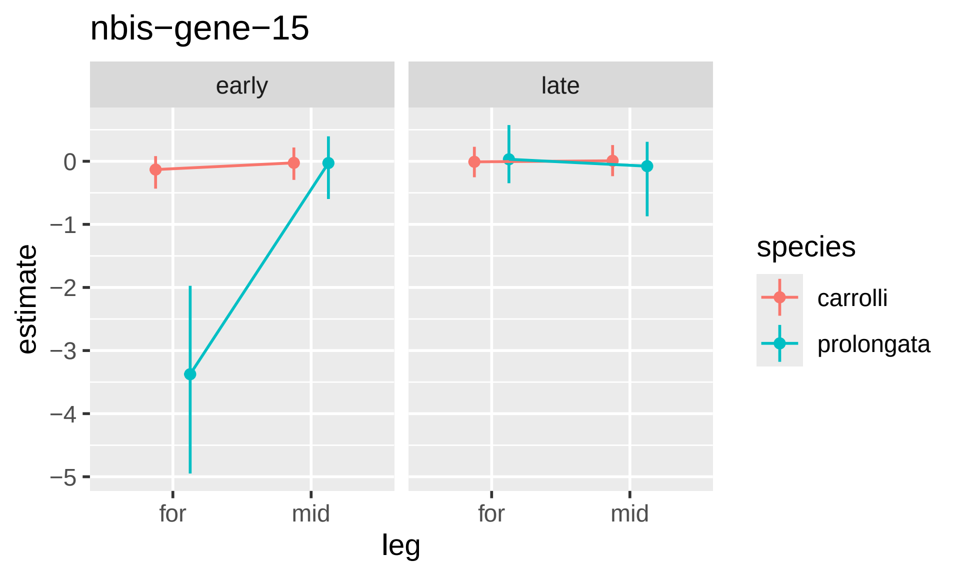
nbis−gene−15
early
late
estimate
0
−1
−2
−3
−4
−5
for
mid
for
mid
leg
species
carrolli
prolongata

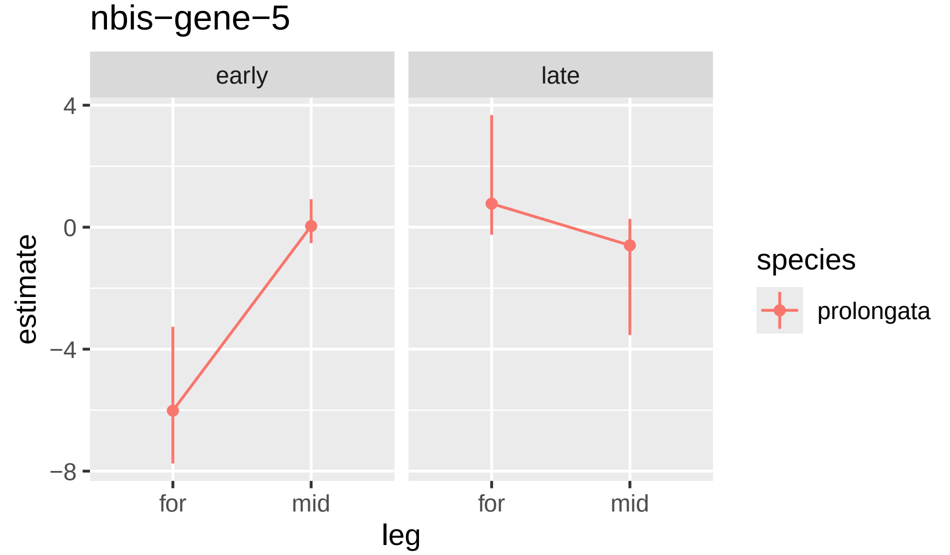

nbis−gene−5
early
late
4
0
−4
−8
estimate
for
mid
for
mid
leg
species
prolongata

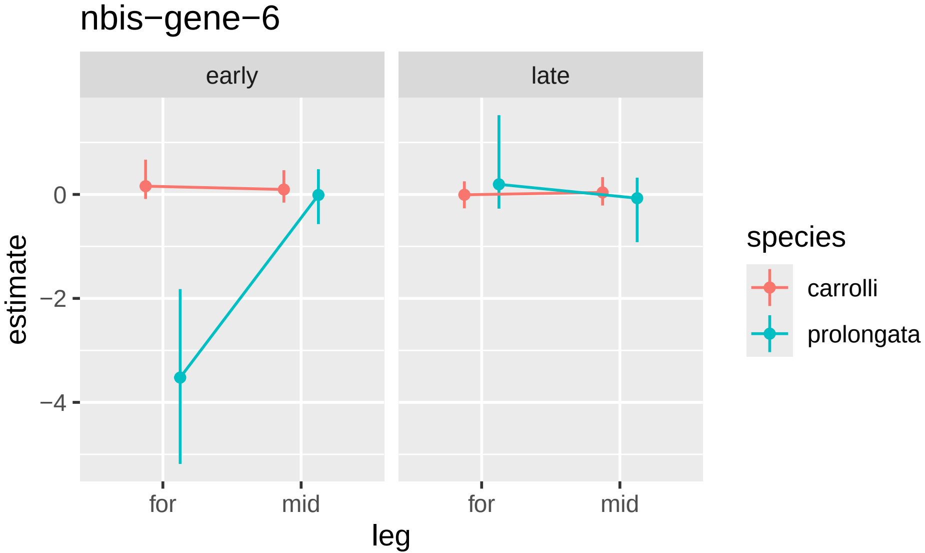

nbis−gene−6
early
late
0
−2
−4
estimate
for
mid
for
mid
leg
species
carrolli
prolongata

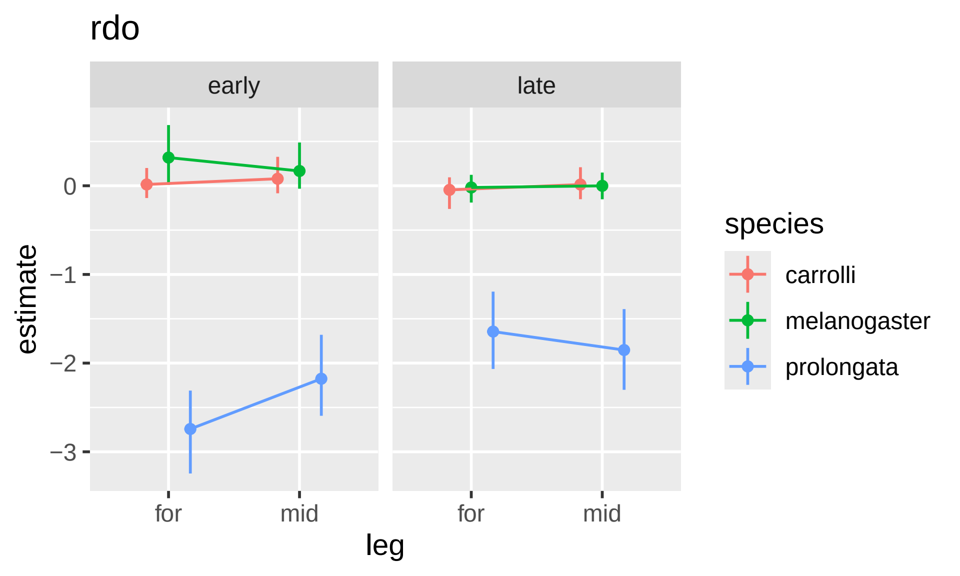

rdo
early
late
0
−1
−2
−3
estimate
for
mid
for
mid
leg
species
carrolli
melanogaster
prolongata

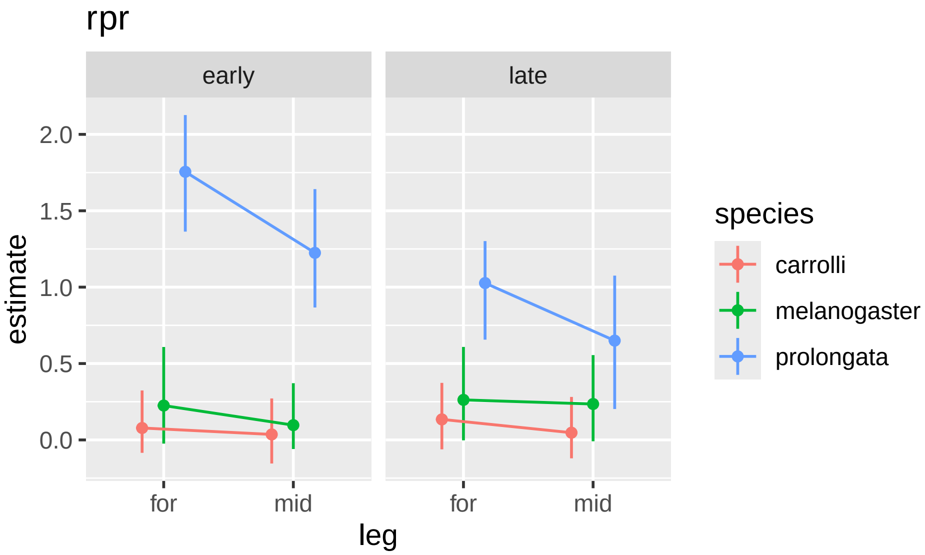

rpr
early
late
estimate
2.0
1.5
1.0
0.5
0.0
for
mid
for
mid
leg
species
carrolli
melanogaster
prolongata

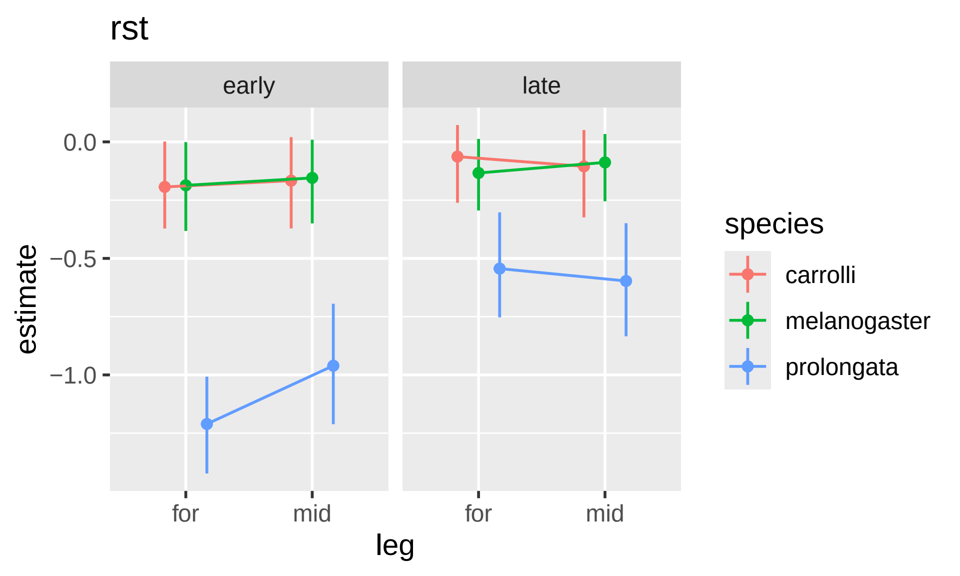

rst
early
late
0.0
−0.5
−1.0
estimate
for
mid
for
mid
leg
species
carrolli
melanogaster
prolongata

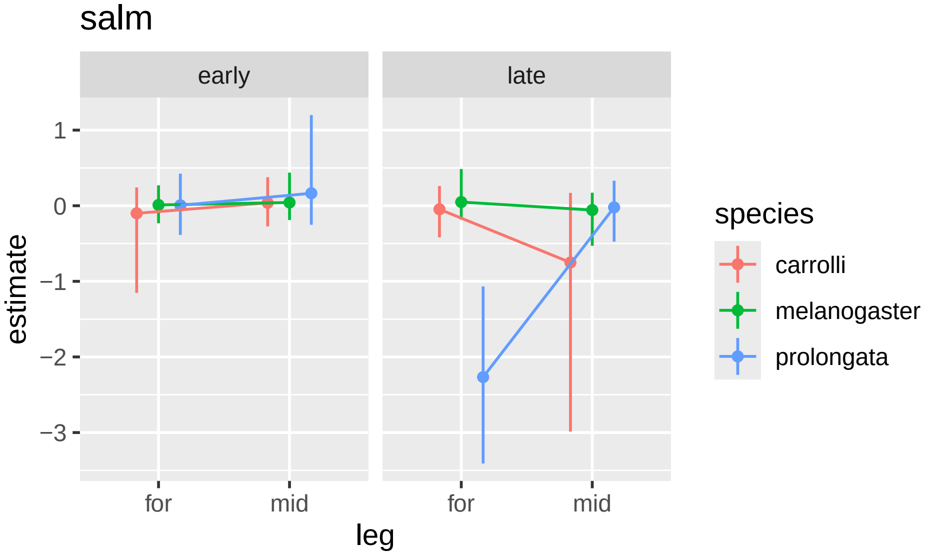
salm
early
late
1
0
−1
−2
−3
estimate
for
mid
for
mid
leg
species
carrolli
melanogaster
prolongata

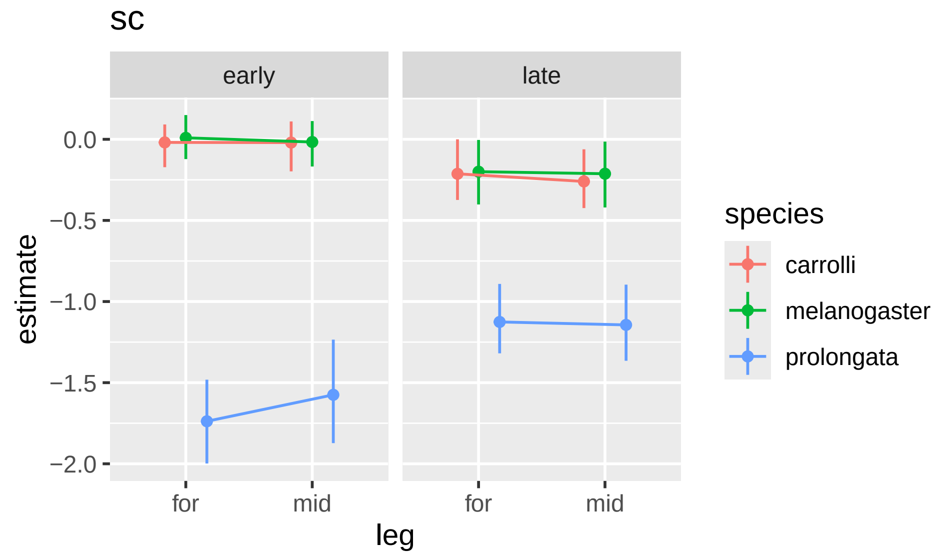
sc
early
late
0.0
−0.5
−1.0
−1.5
−2.0
estimate
for
mid
for
mid
leg
species
carrolli
melanogaster
prolongata

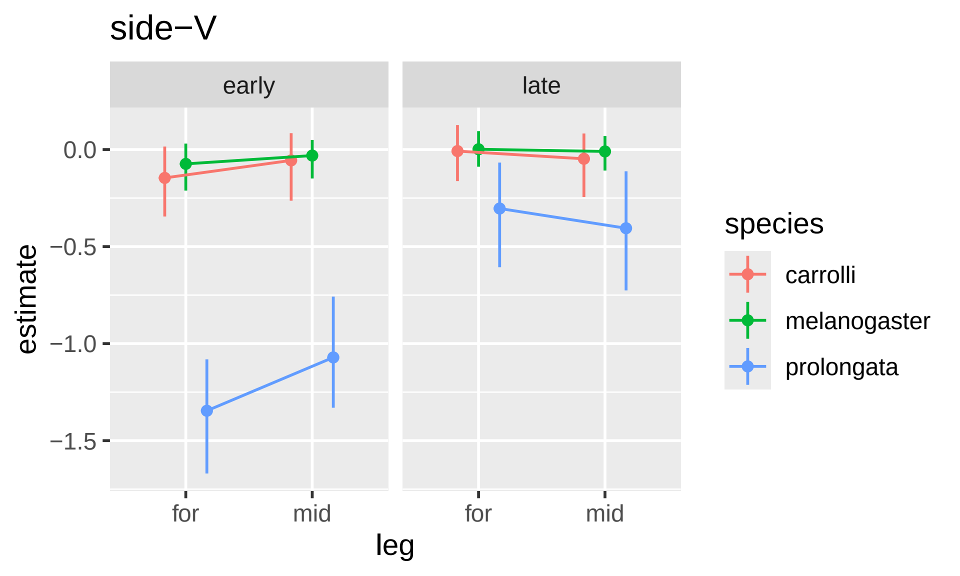
side−V
early
late
estimate
0.0
−0.5
−1.0
−1.5
for
mid
for
mid
leg
species
carrolli
melanogaster
prolongata

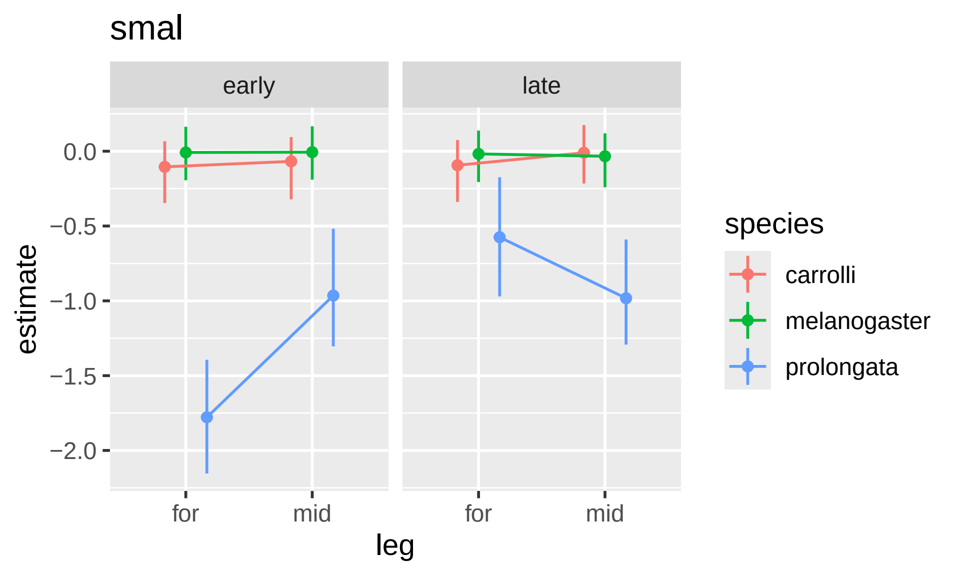

smal
early
late
0.0
−0.5
−1.0
−1.5
−2.0
estimate
for
mid
for
mid
leg
species
carrolli
melanogaster
prolongata

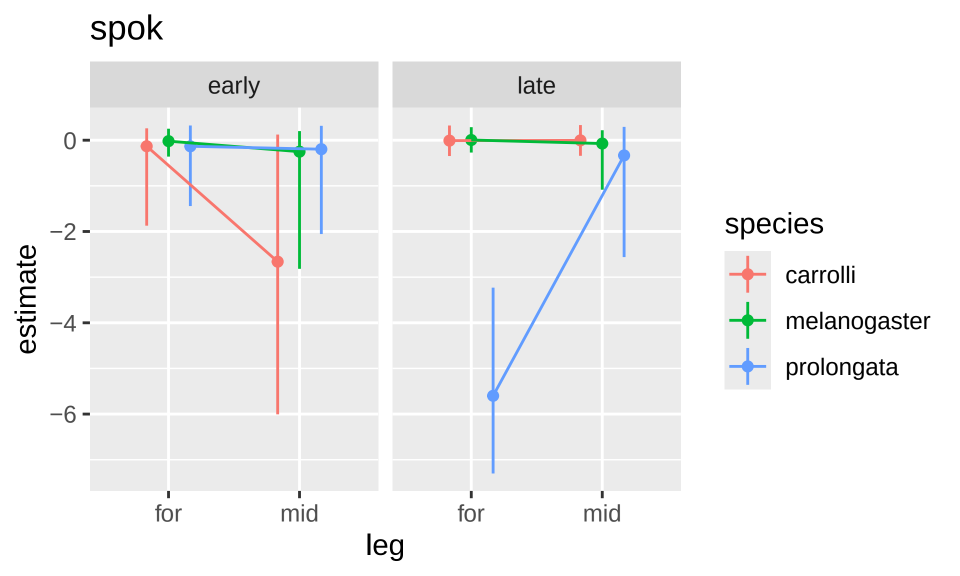

spok
early
late
0
−2
−4
−6
estimate
for
mid
for
mid
leg
species
carrolli
melanogaster
prolongata

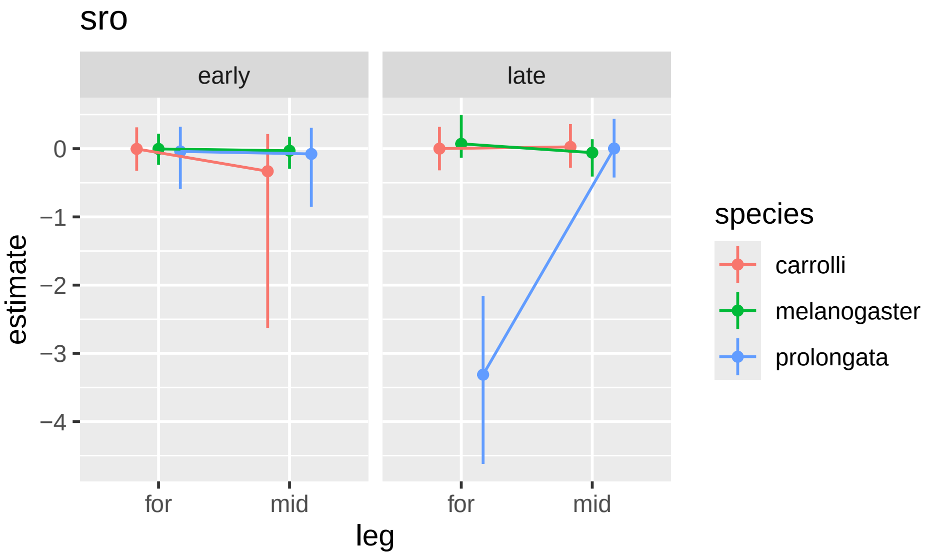

sro
early
late
0
−1
−2
−3
−4
estimate
for
mid
for
mid
leg
species
carrolli
melanogaster
prolongata

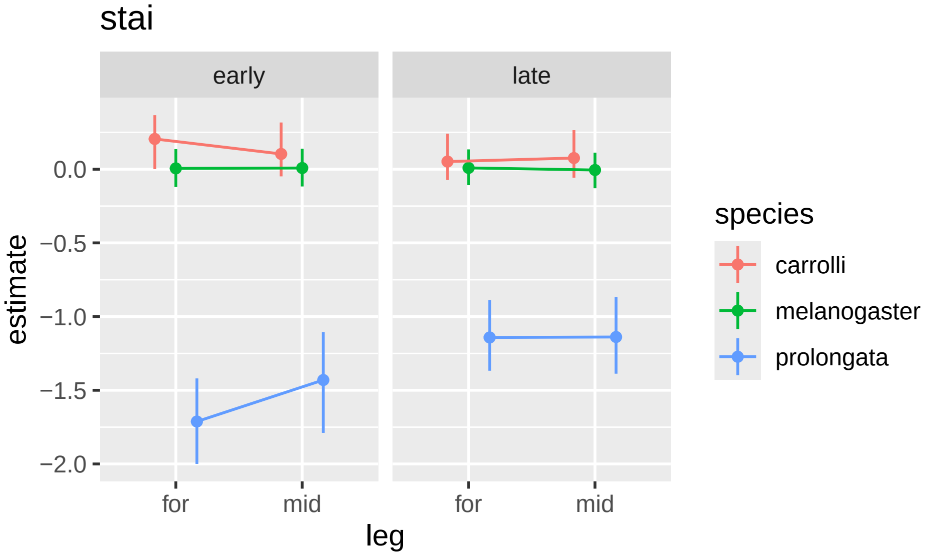
stai
early
late
0.0
−0.5
−1.0
−1.5
−2.0
estimate
for
mid
for
mid
leg
species
carrolli
melanogaster
prolongata

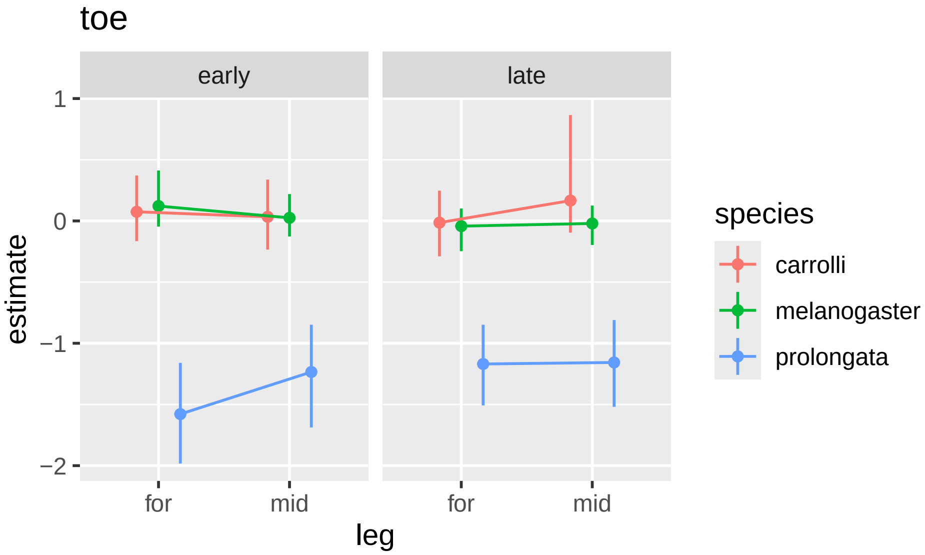
toe
early
late
1
0
−1
−2
estimate
for
mid
for
mid
leg
species
carrolli
melanogaster
prolongata

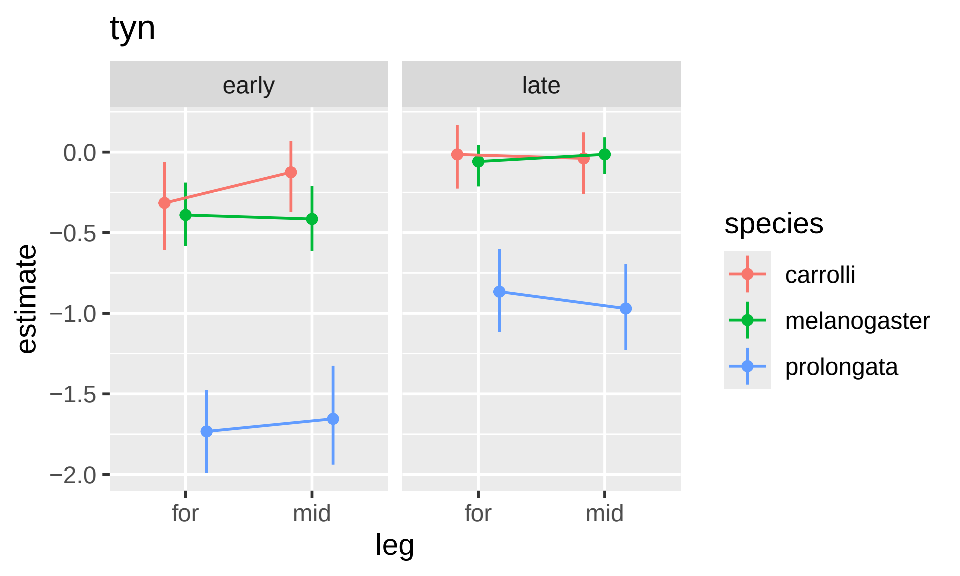
tyn
early
late
0.0
−0.5
−1.0
−1.5
−2.0
estimate
for
mid
for
mid
leg
species
carrolli
melanogaster
prolongata
